# Low E2F2 activity is associated with high genomic instability and PARPi resistance

**DOI:** 10.1101/777870

**Authors:** Jonathan P Rennhack, Eran R. Andrechek

## Abstract

The E2F family, classically known for a central role in cell cycle, has a number of emerging roles in cancer including angiogenesis, metabolic reprogramming, metastasis and DNA repair. E2F1 specifically has been shown to be a critical mediator of DNA repair; however, little is known about DNA repair and other E2F family members. Here we present an integrative bioinformatic and high throughput drug screening study to define the role of E2F2 in maintaining genomic integrity in breast cancer. We utilized *in vitro* E2F2 ChIP-chip and over expression data to identify transcriptional targets of E2F2. This data was integrated with gene expression from E2F2 knockout tumors in an MMTV-Neu background. Finally, this data was compared to human datasets to identify conserved roles of E2F2 in human breast cancer through the TCGA breast cancer, Cancer Cell Line Encyclopedia, and CancerRx datasets. Here we have computationally predicted that E2F2 transcriptionally regulates key mediators of DNA repair. Our gene expression data supports this hypothesis and low E2F2 activity is associated with a highly unstable tumor. In human breast cancer E2F2, status was also correlated with a patient’s response to PARP inhibition therapy. Taken together this manuscript defines a novel role of E2F2 in cancer progression beyond cell cycle and could be therapeutically relevant.

**Author Summary:** The E2F family of proteins have been known to regulate cell cycle and have recently been shown to have a number of roles in tumor progression. Here we use a combination of computational techniques and high-throughput drug screening data to establish a novel role of E2F2 in maintaining genomic integrity. We have shown that a number of direct and indirect target genes of E2F2 are involved in multiple classical DNA repair pathways. Importantly, this was shown to be unique to E2F2 and not present with other activator E2Fs like E2F1. We have also shown that E2F2 activity is positively correlated with PARP inhibitor sensitivity regardless of BRCA1/2 status. This is important due to the recent approval of PARP inhibitor therapy in the clinic. Based on our work E2F2 activity could serve as a novel biomarker of response and may identify a new cohort of patients which could benefit from PARPi therapy.

## Introduction

Breast cancer remains the leading cause of cancer related deaths in women. This is largely due to two factors, metastasis and the heterogeneity of breast cancer. While metastasis to distal sites is responsible for mortality, the difficulty of treating a heterogeneous disease is one of the primary factors allowing that progression to occur. The heterogeneity of breast cancer is evident in several facets, including histological subtypes, progression and response to treatment. Underlying this diversity are the unique genomic alterations, methylation patterns and the resulting gene expression differences that are recognized in the PAM50 classification system (1, 2). Each of the subtypes present (Luminal A/B, Basal, Claudin Low, HER2+ve and normal like) have a unique transcriptomic profile, resulting in the dysregulation in key proteins in breast cancer, including the alteration of the E2F family of transcription factors (3–7).

The E2F family of transcription factors is composed of nine unique family members (E2F1, E2F2, E2F3a, E2F3b, E2F4, E2F5, E2F6, E2F7, and E2F8) (8–10). These family members bind a conserved motif with gene specificity contributed by cofactors (11–13). As a result, the E2Fs have been shown to directly regulate many downstream genes. Classically they have been divided into family members that activate transcription (E2F1, E2F2, and E2F3a) and repressors of transcription (E2F3b, E2F4, E2F5, E2F6, E2F7, and E2F8). These definitions have recently become less clear, with each family member functioning in both activating and repressing roles depending on the tissue and developmental context (11, 14).

The role of the E2F family has widely been described in cell cycle where the members regulate the G1/S checkpoint in response to Cyclin D levels (15, 16). However, beyond the G1/S checkpoint de-regulation E2Fs have a number of emerging roles in cancer (17). This includes roles in other aspects of cancer progression including angiogenesis (18), metabolic reprogramming (19), and apoptosis (20, 21). Indeed, numerous accounts detail the role of the activators in metastasis of human breast cancer as well as mouse models of the disease (3, 4, 6, 22–24).

An additional emerging role for the E2Fs has been in the regulation of genomic stability. Specifically, the role of E2F1 has been well defined with both transcriptional and non-transcriptional roles in DNA repair (25). In response to DNA damage, E2F1 undergoes post translational phosphorylation by ATM (26), leading to protein stabilization and increased expression of repair proteins. In addition to the transcriptional role in DNA repair, E2F1 is physically recruited to sites of damage. During cases of double stranded breaks (27) or UV damage (28) it was observed that E2F1 formed foci with other damage induced proteins at the site of DNA damage. It has been shown the E2F1 is required for the efficient recruitment of other repair proteins including XPA/XPC (28) and NBS1 (29). It has also been shown that E2F2 is transcriptionally upregulated in response to DNA damage and has been shown to complex with Rad51 and sites of DNA damage in neuronal cells (30).

The amplification of the centrosome within a cell leads to defects in cellular segregation and DNA replication, which in turn leads to the single nucleotide variants, copy number alterations, and translocations characteristics. Importantly, activator E2Fs have been shown to be associated with centrosome amplification (31). It is through this amplification of the centrosome that it is believed E2Fs contribute to the DNA instability associated with their misregulation. However, the mechanism and specific E2Fs involved in this process remain unclear.

Together, there is an emerging role for the activator E2Fs role in maintaining genomic integrity, but only the role of E2F1 has been well defined. Here we present a key role for E2F2 in maintaining genomic integrity. Through the use of cell lines, mouse models, and human samples, we have identified that low E2F2 activity level is associated with tumors containing high levels of genomic instability. Furthermore, the levels of E2F2 have direct impact on therapeutic response in clinical data. Indeed, tumors with high E2F2 activity have an increased sensitivity to cell cycle targeted chemotherapy as well as targeted PARP inhibitors.

## Results

Based on the published literature for E2Fs in non-cell cycle roles, we hypothesized that E2F2 had key activities other than the traditional role in cell cycle. To test this hypothesis in large transcriptomic datasets, we used principle components analysis on gene expression data from cells infected with adenoviral delivered GFP compared with adenoviral delivered E2F2 (Figure 1A). This analysis revealed a consistent gene expression profile associated with over expression of E2F2 (Figure 1B). We used Significance Analysis of Microarrays (SAM) (32) analysis to identify consistently overexpressed genes with the infection of Ad-E2F2 relative to GFP. E2F2 induction allowed for the identification of overrepresented gene ontology groups through the use of PANTHER analysis (Figure 1C). As expected, this uncovered over-representation of cell cycle proteins. Interestingly we also identified a number of repair associated gene ontology groups, including double-stranded break repair and non-recombinational repair.

**Figure 1:**
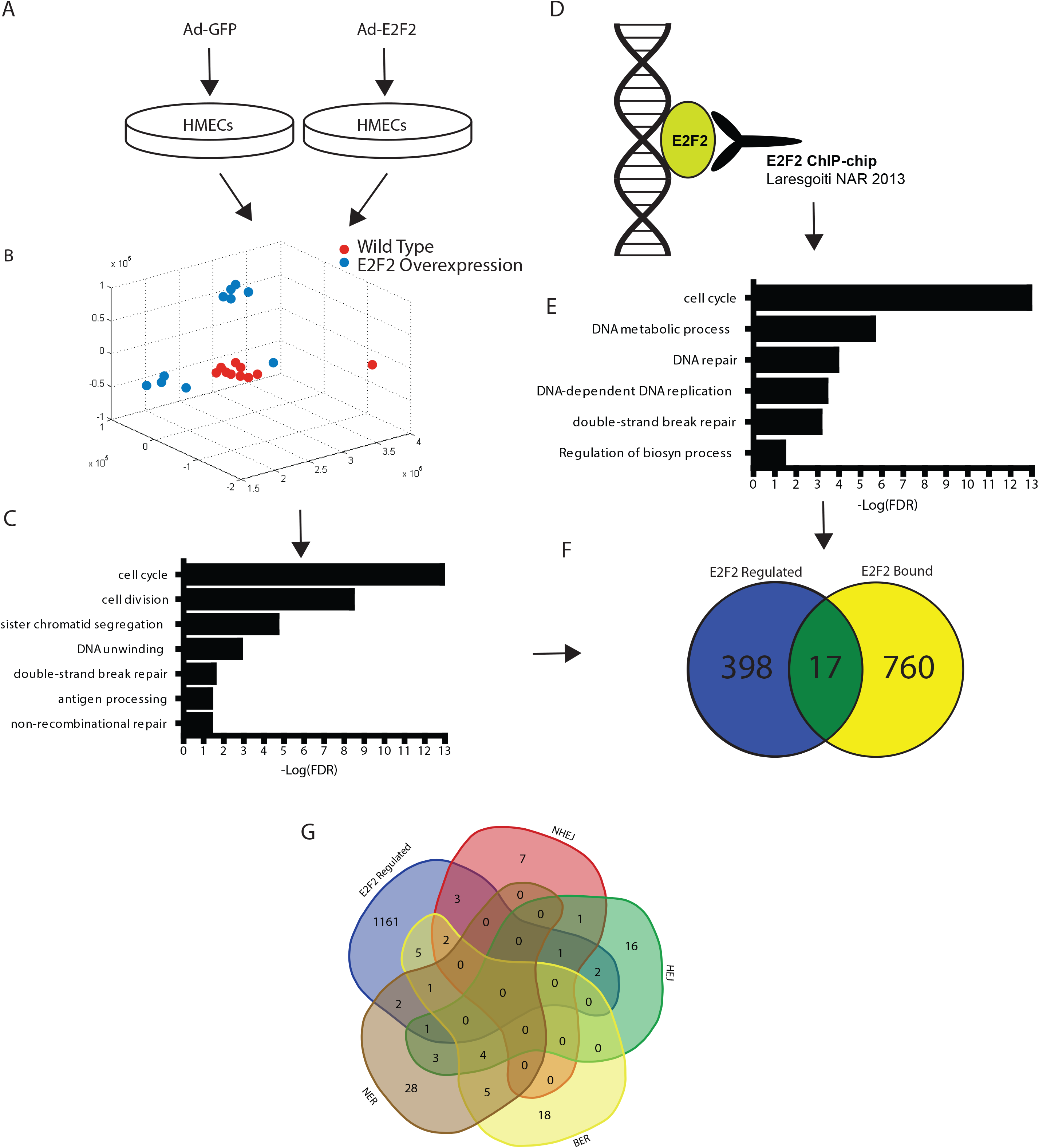
E2F2 target genes are enriched for DNA repair associated proteins. To understand genes regulated by E2F2 we utilized E2F2 overexpression data to compare the expression profile of HMECs infected with Ad-GFP and Ad-E2F2 (A). A graphic representation of the first three principle components reveals a consistent transcriptional response associated with E2F2 overexpression through displaying MEFs plus Ad-GFP (blue) and MEFs plus Ad-E2F2 (red) (B). Genes overexpressed with the addition of E2F2 as identified by SAM show a significant (FDR<.05) overrepresentation in key gene groups including cell cycle and repair (C). ChIP-CHIP of E2F2 binding genes show binding across the genome (D). The binding targets show a significant (FDR <.05) overrepresentation in a cancer related gene groups (E). A Venn Diagram showing genes predicted to be regulated by E2F2 from ChIP-Chip binding and over expression analysis show a small overlap in genes (F). E2F2 overexpressed or bound genes (E2F2 regulated) are shown to play a role in major repair pathways including Non-Homologous End Joining, Homologous End Joining, Base Excision Repair, and Nucleotide Excision Repair (G).

To identify genes and pathways directly regulated by E2F2, we utilized publicly available E2F2 ChIP-Chip data (Figure 1D) (33). This revealed numerous genes bound by E2F2 across the genome. When target genes were analyzed with PANTHER and GATHER, the ontologies were consistent with the E2F2 overexpression data. Indeed, cell cycle and DNA damage repair gene ontology groups were overrepresented, including DNA repair and double-stranded break repair (Figure 1E). Given the similar pathways it was surprising that when we compared predicted E2F2 targets obtained through ChIP-Chip and gene expression analysis that we identified little overlap in the gene lists (Figure 1F). This may indicate that the pathways are regulated by both direct and indirect / downstream targets of E2F2. To determine if there was bias towards one particular repair pathway, we identified overlap between E2F2 regulated genes (gene expression and ChIP-chip) and each of the major repair pathways including Non-Homologous End Joining, Homologous End Joining, Base Excision repair, and nucleotide excision repair. This analysis illustrated that E2F2 regulates key proteins in each repair pathway (Figure 1G).

To determine if there is a role for E2F2 in DNA repair processes in the *in vivo* setting, we utilized publicly available E2F2 knockout transcriptome data within the MMTV-Neu mouse model mammary tumors (3, 34). Unsupervised clustering identified a consistent transcriptional profile with E2F2 loss (Figure 2A). This revealed five major clusters, three primarily composed of MMTV-Neu E2F2 knockout samples and two clustered primary populated with MMTV-Neu E2F wildtype samples. This demonstrated a unique gene expression profile associated with the loss of E2F2, which was unique relative to the MMTV-Neu E2F wildtype background. To explore the enriched cellular processes in this data, we used Gene Set Enrichment Analysis (GSEA) comparing MMTV-Neu tumors with and without E2F2. This revealed, similar to the *in vitro* data, that E2F2 null samples were enriched for both the instability gene set (Figure 2B) and the repair gene set (Figure 2C). We do not believe this to be in conflict with the *in vitro* data with an upregulated gene expression signature for repair not necessarily correlating with an increase in the repair process.

**Figure 2:**
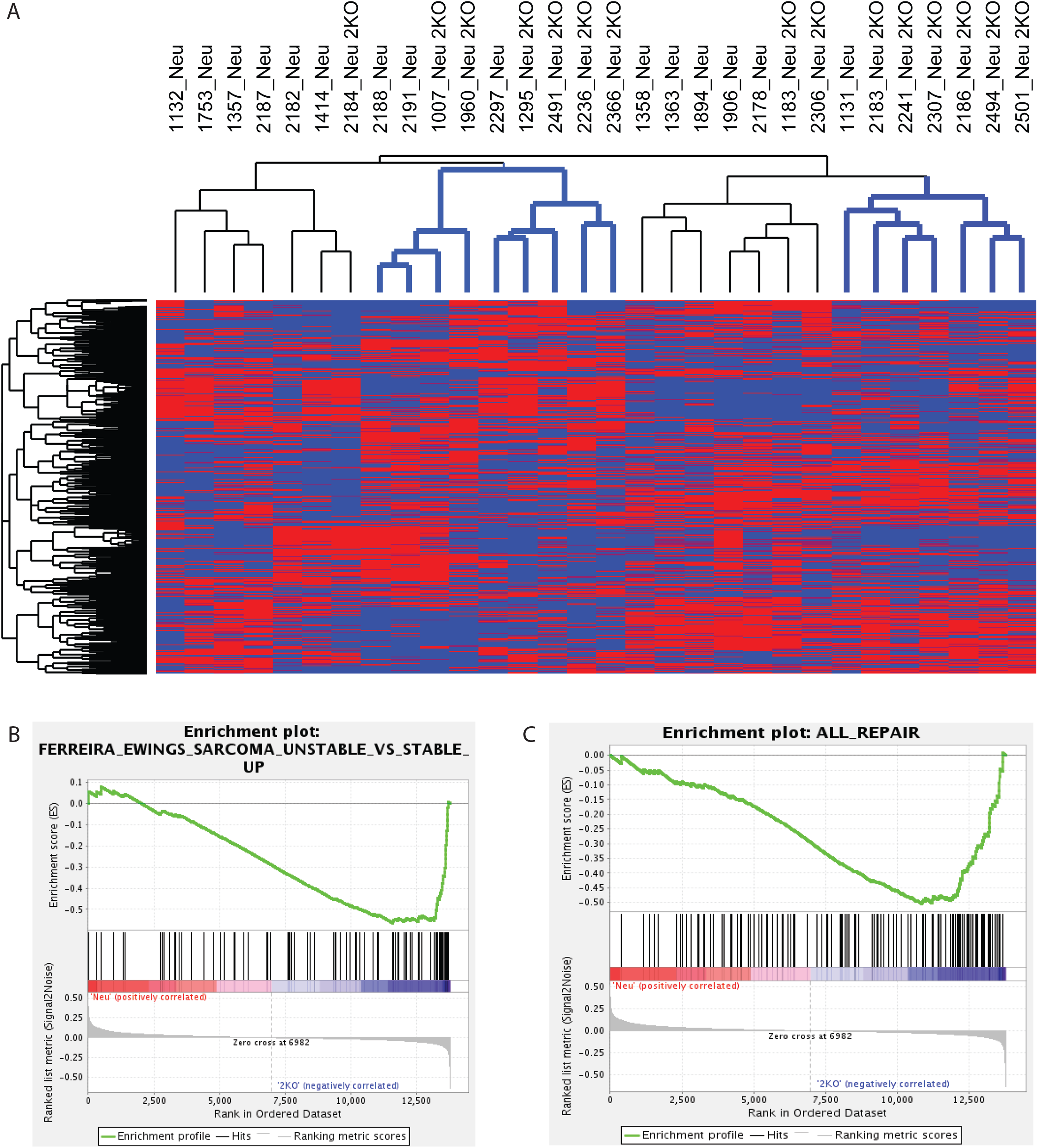
Loss of E2F2 is associated with an enrichment of genomic instability markers. E2F2 loss is associated with consistent transcriptional responses as revealed by unsupervised hierarchical clustering. MMTV-Neu or MMTV-Neu E2F2 knockout samples are arranged by column and unique genes by row. Gene expression values are represented by color from low (blue) to high (red) as indicated by the color bar. It is revealed that 2 main clusters contain an overrepresentation of MMTV-Neu, E2F2 KO samples (blue) (A). Consistent gene expression sets are enriched with the loss of E2F2 as identified by gene set enrichment analysis. This includes instability gene sets (B) and repair gene sets (C).

To test whether misregulation of E2F2 associated repair and instability functions were associated with a more unstable tumor, we predicted gene copy number alterations across the genome. We utilized the Analysis of CNAs by Expression data (ACE) algorithm (35) to predict copy number profiles of control FVB wildtype mammary glands (Figure 3A), MMTV-Neu E2F2 wildtype mammary tumors (Figure 3B), and MMTV-Neu E2F2 knockout mammary tumors (Figure 3C). Examining a portion of chromosome 4 as a case study, we observed an increased number of significant (p<.05) amplification and deletion events in the MMTV-Neu E2F2 knockout samples relative to both wild FVB mammary controls and MMTV-Neu E2F2 wildtype tumors. Expanding to the entire genome, this was a consistent across genotypes with the E2F2 knockout samples being the most unstable (Figure 3D). Interestingly, when this was compared to the E2F1 knockout samples we also saw the E2F2 null samples to be increasingly unstable indicating a specific role of E2F2 in genomic stability (Figure S1).

**Figure 3:**
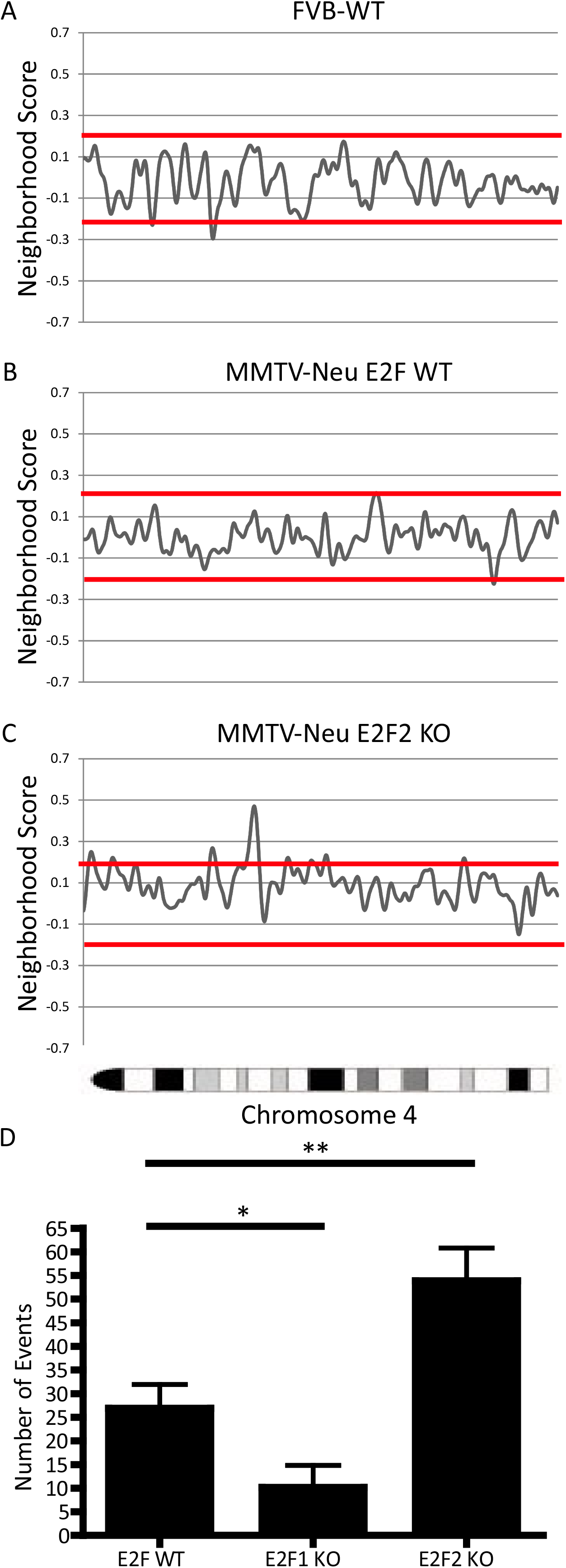
E2F2 loss is associated with higher number of copy number alterations in the MMTV-Neu mouse model. Copy number analysis obtained through the ACE algorithm for FVB tail DNA (A), MMTV-Neu tumor (B), and MMTV-Neu E2F2 knockout tumor (C) shows increased copy number variants in chromosome 4 in the E2F2 knockout sample compared to the normal DNA and MMTV-Neu tumor. Neighborhood score is graphed along the y-axis and location along the x axis. Significant (p<.05) amplification or deletion points are signified by the neighborhood score above or below the red line. This is shown to be significant across the entire genome with the E2F2 null samples having a significant increase in number of copy number event and the E2F1 null samples having a decrease in the number of copy number alterations (*=p<.05, **=P<.01).

While we have identified a potential role for E2F2 associated instability in mouse tumors, this role has not previously been examined in human breast cancer. To address a potential role for E2F2 in breast cancer, an E2F2 activity signature was used to divide the TCGA breast cohort into low / high E2F2 activity groups. As predicted from the mouse mammary tumor data, human breast cancer with low E2F2 activity contained significantly more copy number variants than those with high E2F2 activity (p<.05) (Figure 4A). Furthermore, tumors with low predicted E2F2 activity were observed to have enrichment for genomic instability in gene set enrichment analysis. Specifically, there was a significant enrichment of genes involved with the response to UV induced damage (Figure 4B).

**Figure 4:**
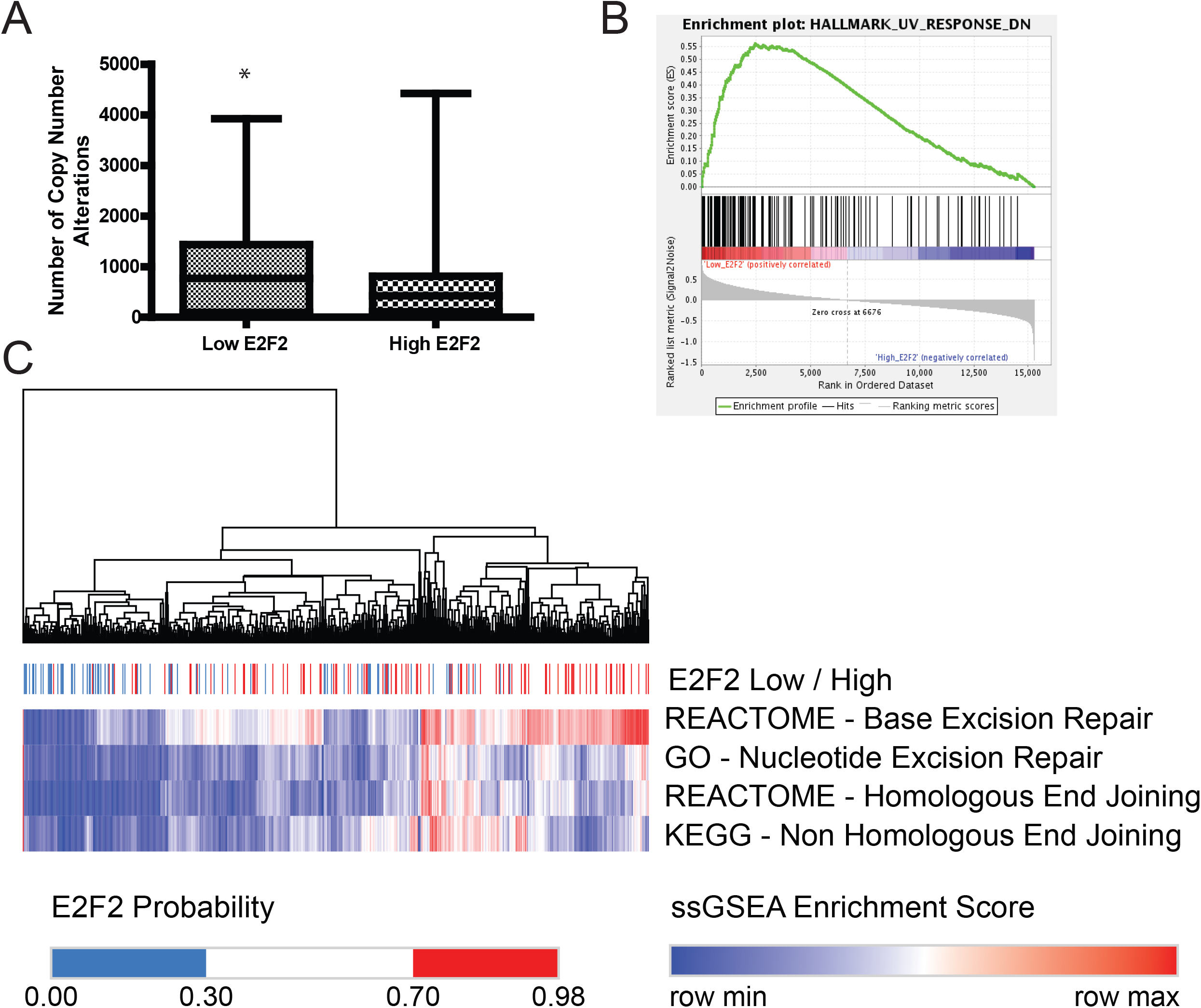
Low E2F2 activity is associated with decrease repair gene expression in the TCGA breast cancer cohort. The TCGA cohort reveals that low E2F2 activity is associated with higher number of genes with copy number alterations (A)

In order to determine if E2F2 preferentially regulated a specific repair pathway, we utilized single sample gene set enrichment analysis (ssGSEA) (36). The resulting scores were low/high normalized to return an activity score between 0 and 1 for the four major repair pathways. Low E2F2 activity resulted in significantly lower activity in each repair pathway including: Base Excision Repair, Nucleotide Excision Repair, Homologous End Joining, and Non-Homologous End joining (Figure S2). Unsupervised hierarchical clustering of this data revealed that regardless of the pathway, low E2F2 activity was associated with low ssGSEA pathway scores (Figure 4C).

Alterations in DNA repair pathways are manifested in response to therapy. Accordingly, we sought to test whether E2F2 levels impacted therapeutic response using the CancerRx dataset. After predicting the E2F2 activity level across all breast cancer datasets we identified differentially lethal compounds between E2F2 low cell lines and E2F2 high cell lines (Table S2). This revealed a number of interesting candidate compounds. For example. tumors with low E2F2 activity responded poorly to cell cycle inhibiting compounds such as Cisplatin. However, they responded well to PIK3 targeted therapy such as PI-103. In addition, we observed that tumors with high E2F2 activity were sensitive to cell cycle compounds and resistant to other forms of therapy.

With a role for E2F2 in repair, we examined response to repair targeted therapy, including PARP inhibitors. Surprisingly we identified that high E2F2 activity was associated with a significantly higher response to common PARP inhibitors including Talazoparib (Figure 5A), Olaparib (Figure 5B), and Rucaparib (Figure 5C) as identified by a significant decrease in the IC50 of each compound on each cell line. Importantly, this correlation was independent of the BRCA1/2 status of the cell line. To test if this correlation was a function of any E2F or specific to E2F2, we performed a similar analysis for E2F1. In this analysis, we predicted E2F1 activity through the use of a gene activity signature and identified the IC50 of each cell line. Importantly, in this analysis we saw no difference in the IC50 or PARP targeted therapy in low or high E2F1 activity (Figure S3). This indicates the identified differences in response to PARP targeted therapy are specific for E2F2.

**Figure 5:**
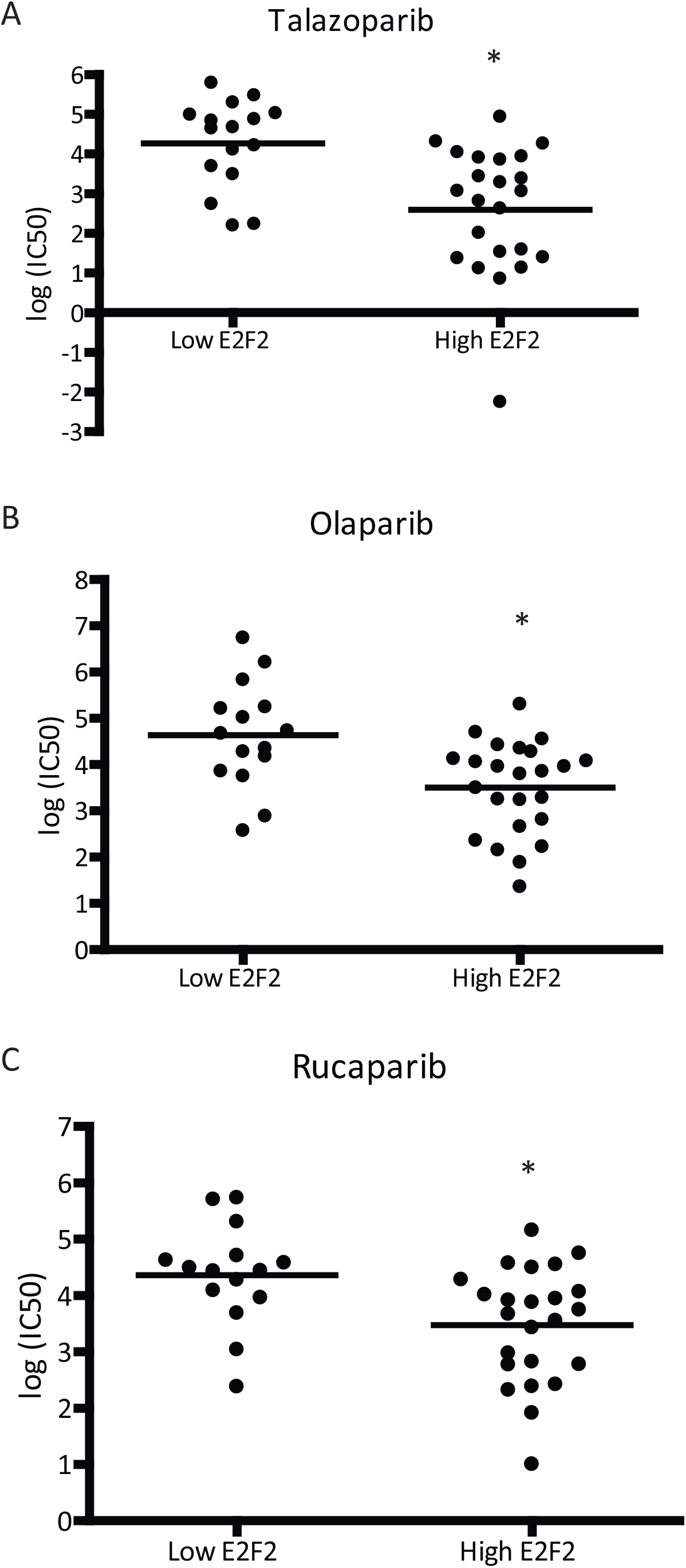
Low E2F2 activity is associated with resistance to PARP inhibitors across breast cancer cell lines. Breast cancer cell lines divided in lowly active E2F2 and highly active E2F2 show a consistent resistance to PAPRP targeted therapies associated with low E2F2 activity (p<.05) for Talazoparib (A), Olaparib (B), and Rucaparib (C).

## Discussion

Here we describe a role of E2F2 in repair and maintenance of genome integrity in both MMTV-Neu mouse model mammary tumors as well as in human breast cancer patients. We have identified this role using a combination of *in silico*, *in vitro* and *in vivo* datasets. Specifically, we observed that E2F2 controls key members of many different repair pathways including HEJ, NHEJ, BER, and NER. This involvement is associated with a genomically unstable tumors with the loss of E2F2 activity in both the mouse model and the human disease.

Furthermore, this finding has direct clinical application. We have identified E2F2 activity as a biomarker for response to cell cycle inhibition therapy. E2F2 activity, as determined by a gene expression signature, correlates with PARP inhibition therapy. Interestingly, although tumors with lowly active E2F2 are significantly more unstable than E2F2 tumors they are resistant to PARP inhibitor therapy. We have identified that this effect is independent of BRCA1/2 status and is correlated with E2F2 levels. This indicates that E2F2 has a role in PARP response and loss of E2F2 may phenocopy or directly trigger other known causes of PARP inhibitor resistance.

This manuscript serves as a proof of concept study for an expanded role of the E2Fs in DNA repair in breast cancer. The data presented here, combined with the previously established literature, show key roles of E2F1 and E2F2 as contributing factors in the genomic instability seen in breast cancer. Further research needs to be completed of the other E2F family members to identify if this role is unique to E2F1 and E2F2 or if other E2Fs may play similar roles. Indeed, while certain E2Fs have specific roles in mammary development (37, 38) and tumor biology (3, 4, 6), there is significant overlap and compensation amongst the E2Fs (38, 39). Taken together, this data shows the key role that the E2Fs play in cancer progression and heterogeneity. As central drivers of repair and due to the impact they have on the ability of a tumor to respond to key therapies, there must be more research to understand the clinical application of E2F status and the way that it should shape patient care.

## Materials and Methods

### Datasets Used

For the E2F2 overexpression data, Affymetrix cDNA microarray profiled gene expression data was downloaded from previously published data (40). For E2F2 binding analysis we utilized ChIP-Chip for data for E2F2 T lymphocytes isolated from 4-week-old C57B16:129SV mice (33). For the E2F2 knockout data in the MMTV-Neu background we utilized the dataset GSE42533. For the human patient analysis, we utilized the TCGA breast cancer cohort (41).

### Overrepresentation analysis

All overrepresentation analysis experiments were performed through the use of GATHER (42) and PANTHER bioinformatic analysis to identify overrepresented gene ontology groups. Significant groups were noted filtered by a p value of less than .05 and a Bayes factor greater than 3.

### ssGSEA and Clustering analysis

ssGSEA was performed on the Broad genepattern software with the designated genesets for each repair pathway downloaded from msig-db. These scores were normalized between 0 (low) and 1 (high). Samples were clustered through the use of Morpheus (https://software.broadinstitute.org/morpheus/). Clustering was performed using Complete linkage unsupervised hierarchical clustering. Heatmaps were visualized using MATLAB.

### ACE Analysis

ACE analysis was performed as previously described to infer copy number alterations from gene expression data (43, 44). The analysis was performed with default parameters and used a significance cutoff of q<.05.

### E2F2 Activity Levels

E2F2 activity was assayed using a gene expression signature as previously described (37, 45, 46). Briefly this method identifies differentially expressed genes between Ad-GFP and Ad-E2F2 infected cell lines and uses binary regression analysis to compare unknown samples and known controls from each group to get a score between 0 (low E2F2 activity) and 1 (high E2F2 activity).

### E2F2 Drug Sensitivity Data

Breast cancer gene expression data was downloaded from the cancer cell line encyclopedia (47). From this data E2F2 activity was predicted as described above. For drug sensitivity data, we downloaded the small molecule sensitivity dataset from CancerRx.org. Breast cancer cell lines were divided into high and low E2F2 activity groups and significantly different compounds between the two groups were identified by significantly different IC50’s as identified by a student’s T-test.

## Acknowledgements

We would like to acknowledge member of the Andrechek lab for their helpful comments and critical reading of the manuscript.

## Availability of data and material

All datasets used in this publication can be accessed on Gene Expression Omnibus through their appropriate GSE number as noted in the manuscript.

## Supplemental Legends

**Supplemental Figure 1.**
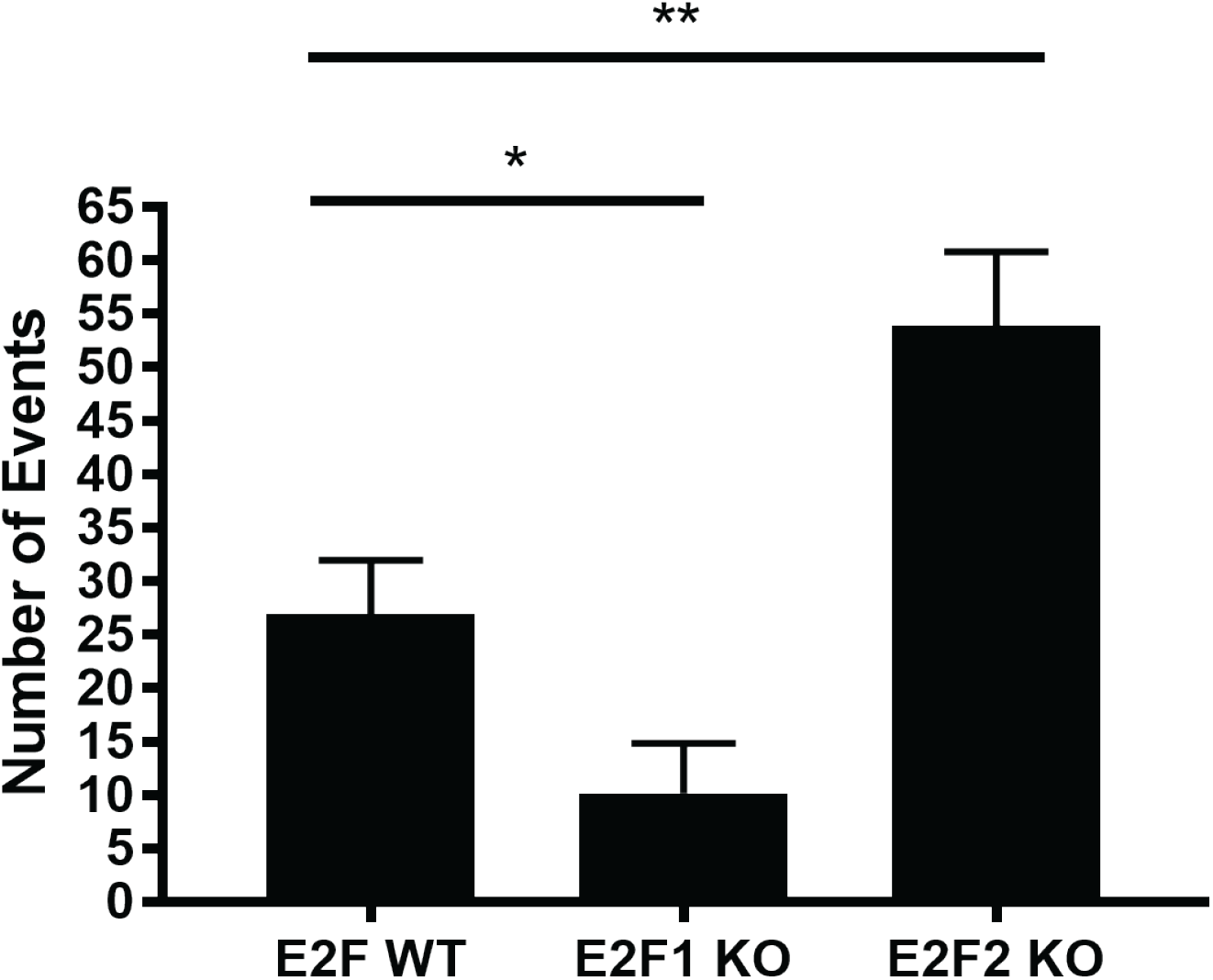
Copy number analysis obtained through the ACE algorithm for MMTV-Neu tumors, and MMTV-Neu E2F1 and E2F2 knockout tumors shows increased copy number variants in the E2F2 knockout sample compared to the normal DNA and MMTV-Neu tumors and decreased variants in the E2F1 knockout background. This is shown to be significant across the entire genome with the E2F2 null samples having a significant increase in number of copy number event and the E2F1 null samples having a decrease in the number of copy number alterations (*=p<.05, **=P<.01).

**Supplemental Figure 2.**
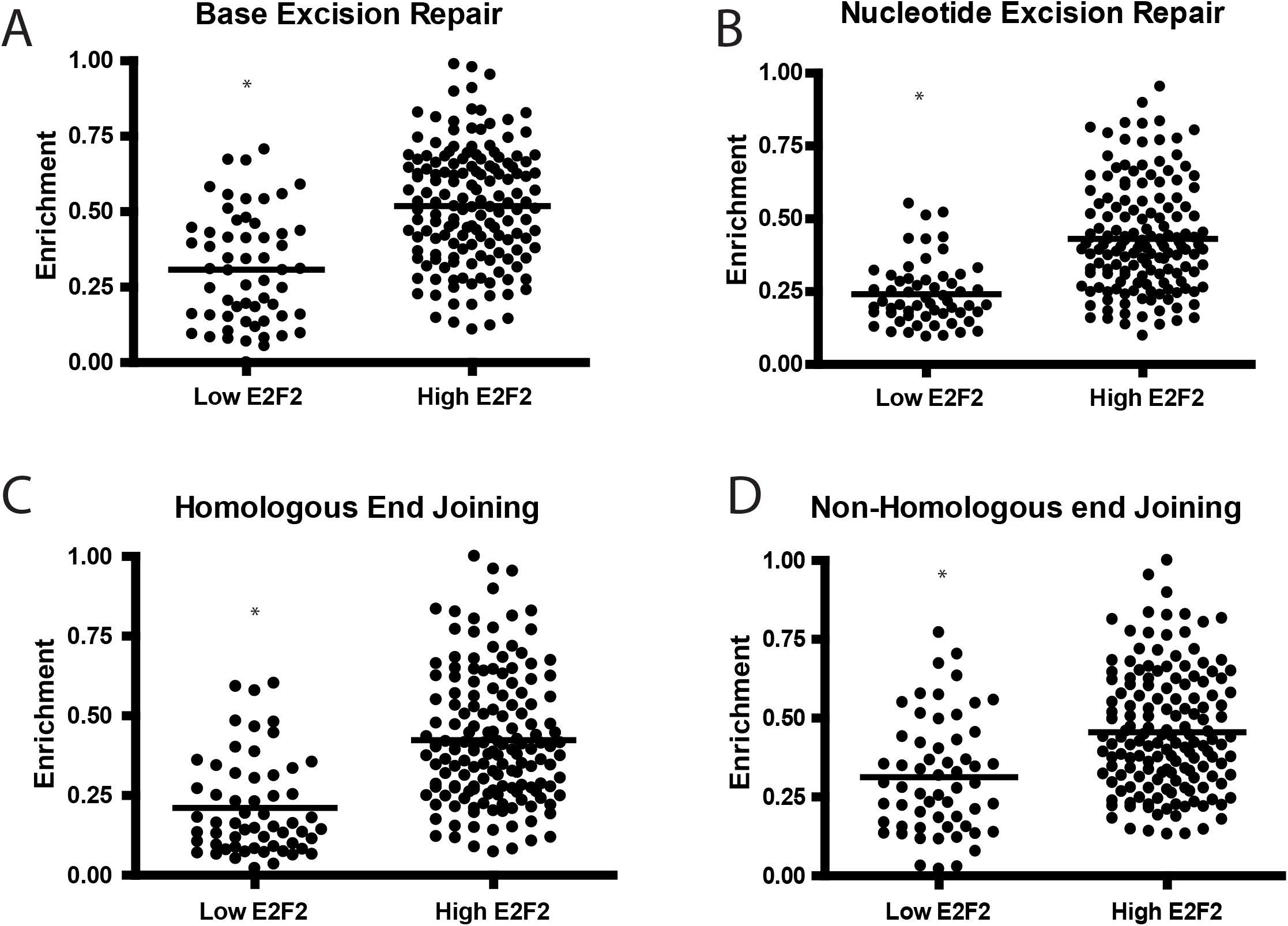
Dotplot representation of figure 4, split by major repair pathway. ssGSEA shows low E2F2 activity is associated with lower gene expression enrichment in a number of different repair pathways including Base Excision Repair (A), Nucleotide Excision Repair (B), and Homologous End Joining (C), Non-Homologous End Joining (D).

**Supplemental Figure 3.**
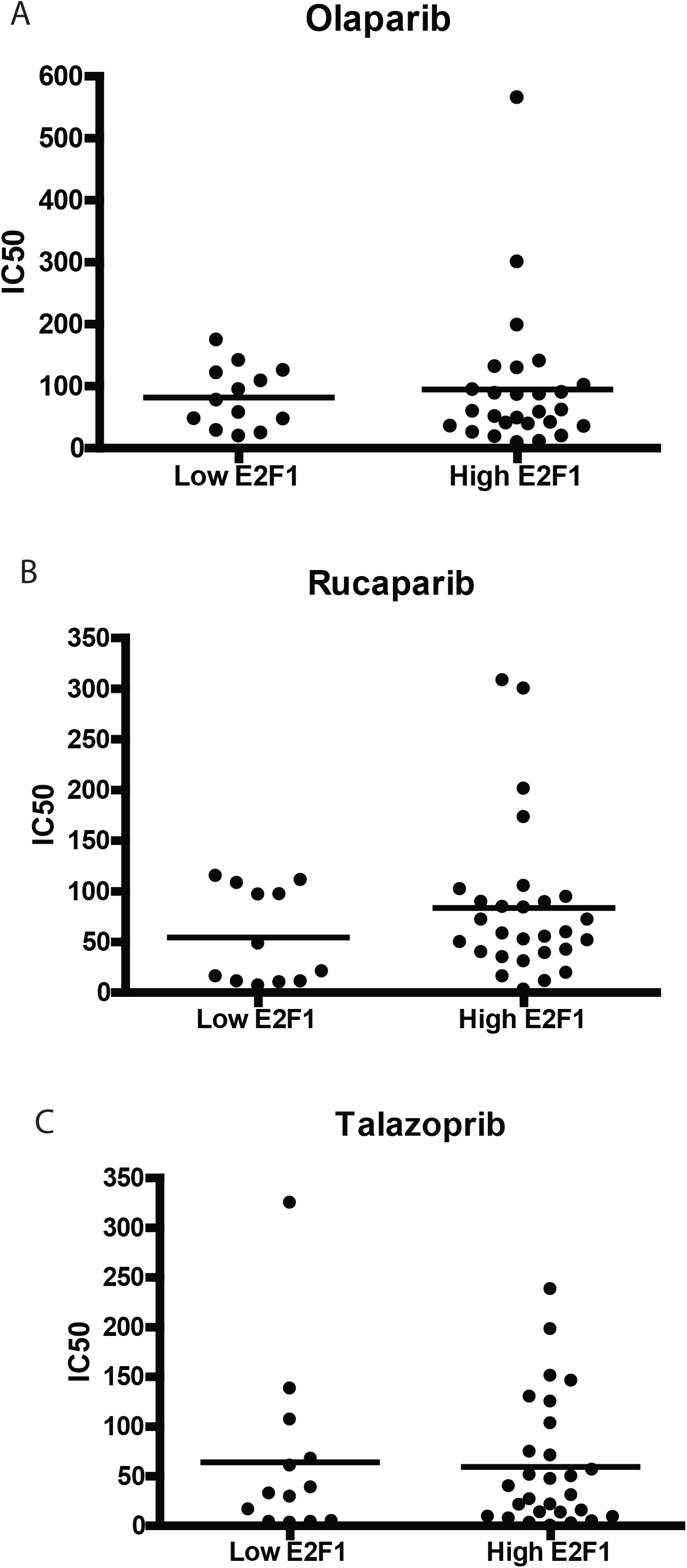
Breast cancer cell lines divided in lowly active E2F1 and highly active E2F1 show no consistent resistance to PAPRP targeted therapies associated with low E2F1 activity for Olaparib (A), Rucaparib (B) or Talazoparib (C).

**Supplemental Table 1**

Table One includes a list of of E2F2 bound or co-expressed genes included in each of the major repair pathways.

**Supplemental Table 2**

Table Two includes the IC50 for various breast cancer cell lines ordered by E2F2 pathway signature activity. The last row shows significance through a t-test for low (<0.5) and high (>0.5) E2F2 predictions. Significant compounds (p<0.05) are shown in bold.

