## Supplementary material for "Low E2F2 activity is associated with high genomic instability and PARPi resistance": Table 1 and 2

| Cell Line | E2F2_Activ | BMN-673 | CHIR-9902 | Olaparib (r | Bleomycin | AG-01469 | AS601245 | AZD7762 | CHIR-9902 |
|---|---|---|---|---|---|---|---|---|---|
| ZR-75-30 | 0.172674 | 4.831413 | 5.778228 | 6.210492 | 7.017756 | 5.7039 | NA | 2.933205 | NA |
| HCC1599 | 0.19589 | NA | NA | NA | 1.878536 | NA | 2.196232 | NA | 3.44923 |
| DU-4475 | 0.221544 | 3.482276 | 3.170432 | NA | 2.756884 | 2.377256 | 1.766916 | -0.63141 | 2.838778 |
| HCC2157 | 0.229377 | 4.984005 | 4.616089 | 4.727899 | 4.325277 | 4.084896 | 2.114977 | -1.05479 | 3.73798 |
| HCC2218 | 0.232926 | 4.670262 | 4.710469 | 5.207943 | 5.927842 | 4.706033 | 3.635666 | 3.684096 | 3.388665 |
| HCC1428 | 0.246971 | 3.686001 | 4.629534 | 4.351636 | 5.377754 | 4.489309 | 2.667916 | 2.058364 | 3.58686 |
| HCC1500 | 0.248124 | 5.018679 | 5.018679 | 5.018679 | 4.823976 | 4.435623 | 2.30015 | 2.047044 | 3.526322 |
| BT-483 | 0.272263 | 2.731514 | 5.131225 | 6.734945 | 6.274419 | 5.731183 | 6.079238 | 3.818782 | 5.606226 |
| EVSA-T | 0.321515 | 2.232709 | 4.080782 | 2.882333 | 1.464536 | 3.685837 | 2.761081 | 0.661062 | 3.024469 |
| HCC1419 | 0.347955 | 5.288988 | 4.903418 | 5.831973 | 7.094178 | 5.305562 | 3.305839 | 3.967624 | 4.507715 |
| T47D | 0.355446 | 5.782737 | 5.785629 | 5.240223 | 4.551292 | 3.033465 | 3.222415 | 4.40608 | 4.593205 |
| CAMA-1 | 0.390238 | 5.470893 | 5.170566 | 3.859481 | 6.161734 | 4.429987 | 3.329246 | NA | 4.202409 |
| MDA-MB-4 | 0.406452 | 4.212353 | 4.61121 | 4.670301 | 6.221267 | NA | 2.631692 | 1.950864 | 4.425192 |
| EFM-19 | 0.424525 | 4.102543 | 5.747966 | 3.750004 | 4.780249 | 4.575343 | 2.222148 | 0.939586 | 1.94289 |
| UACC-812 | 0.448281 | NA | NA | NA | NA | NA | 4.695599 | NA | 3.694253 |
| UACC-893 | 0.463831 | 4.635757 | 5.628592 | 2.569415 | 2.986994 | 4.622383 | 4.012544 | 4.016693 | 4.4424 |
| MDA-MB-3 | 0.488162 | 4.868625 | 4.629582 | 4.173797 | 5.304552 | 4.274598 | 1.421386 | -0.52255 | 3.491228 |
| HCC38 | 0.495164 | 2.195339 | 3.449611 | 4.276386 | 2.142495 | 3.958404 | 1.248765 | -0.22068 | 3.711218 |
| CAL-148 | 0.507011 | NA | NA | NA | NA | NA | 4.346315 | NA | 3.319813 |
| MCF7 | 0.508007 | 1.110639 | 2.17295 | 3.252239 | 2.138198 | 1.907525 | 0.222062 | 1.620276 | 2.275748 |
| AU565 | 0.517117 | 1.372234 | NA | NA | 4.926217 | 2.38322 | 0.784133 | -0.52949 | 3.587735 |
| HCC202 | 0.542512 | 3.28477 | 5.447247 | 4.425001 | 5.610512 | 5.155564 | 2.41998 | 2.048066 | NA |
| BT-474 | 0.542742 | 4.035084 | 4.379491 | 3.851281 | 5.65449 | 2.769537 | 2.254004 | 2.73895 | 3.939417 |
| BT-20 | 0.567886 | 3.065859 | 4.468621 | 4.550445 | 2.543184 | 3.66445 | 3.688836 | 0.937712 | 4.061699 |
| CAL-51 | 0.576739 | -2.25266 | 3.24197 | 1.353893 | -1.58008 | 1.000823 | 1.041737 | -0.58322 | 3.03941 |
| HCC1954 | 0.579405 | 3.933652 | 4.129473 | 4.352374 | 1.67788 | 3.908693 | 1.821509 | 1.056772 | 3.951843 |
| HCC1806 | 0.589735 | NA | 3.861043 | 2.222954 | -1.20819 | 2.819255 | 0.217407 | NA | 2.498027 |
| HCC1569 | 0.610772 | 1.3908 | 3.642967 | 3.282759 | 4.254627 | 2.775353 | 0.675198 | 1.005155 | 1.064706 |
| MDA-MB-4 | 0.614431 | NA | 4.663064 | 2.356764 | 1.379961 | 4.01002 | 2.581057 | -0.25101 | 3.39764 |
| HDQ-P1 | 0.615508 | 4.927595 | 3.345958 | 4.057441 | 3.251652 | 4.747196 | 2.062223 | 0.703034 | 2.763785 |
| MDA-MB-1 | 0.623478 | 2.620001 | 3.495571 | 4.076912 | 3.507223 | 2.970848 | 1.289365 | NA | 2.738685 |
| MDA-MB-4 | 0.631151 | 3.053612 | 4.839225 | 3.954624 | 4.836065 | NA | 3.032996 | 0.656869 | 3.785517 |
| HCC70 | 0.653552 | 3.851926 | 4.373748 | 4.273914 | 4.045316 | 3.939267 | 1.718007 | -0.89108 | 3.524932 |
| HS-578-T | 0.679204 | 2.806696 | 2.465456 | 4.121294 | 2.319519 | 3.87678 | 2.086645 | -0.68102 | 3.137159 |
| HCC1143 | 0.687603 | 3.379658 | 5.592451 | 5.301073 | 3.545506 | 4.57136 | 1.652607 | -1.28675 | 2.165547 |
| EFM-192A | 0.69005 | 4.310257 | 3.485003 | 4.69476 | 6.57658 | 4.546893 | -0.52607 | 2.378049 | 1.550724 |
| MDA-MB-4 | 0.695288 | 1.522867 | 4.925697 | 1.881495 | 2.175016 | 4.062379 | 2.751514 | 0.266813 | 3.907934 |
| HCC1187 | 0.697293 | 3.904177 | 1.72901 | 2.655109 | 0.829458 | 2.419654 | 0.720429 | -1.73671 | 2.519043 |
| JIMT-1 | 0.710134 | 1.130782 | 3.771768 | 3.236867 | 2.982429 | 4.277253 | 1.322525 | -0.92191 | 3.158416 |
| CAL-120 | 0.712178 | 2.006479 | 3.791532 | 2.152588 | 1.949369 | 3.548974 | 3.435589 | 1.506538 | 3.558759 |
| HCC1395 | 0.715104 | 0.849773 | 3.265475 | 3.79661 | 1.950418 | 4.494484 | 2.794891 | -1.32203 | 1.500492 |
| HCC1937 | 0.772489 | 3.428751 | 3.29749 | 3.955583 | 3.280989 | 3.429646 | 1.561914 | -1.1185 | 3.8994 |
| MDA-MB-2 | 0.77948 | 1.585491 | 3.961346 | 2.807325 | 3.400769 | 2.318364 | 2.113155 | 0.740676 | 3.715641 |
| BT-549 | 0.798877 | 4.257894 | 3.302338 | 3.494637 | 2.215068 | 3.743025 | NA | -0.81413 | NA |
| | Significanc | 0.000473 | 0.000803 | 0.004025 | 0.004676 | 0.007299 | 0.007331 | 0.008144 | 0.009225 |
| | 0.331185 | 4.262131 | 4.816376 | 4.6337 | 4.652338 | 4.360919 | 2.918342 | 1.870265 | 3.774649 |

| | | | | | | | | |
|---|---|---|---|---|---|---|---|---|
| 0.639144 | 2.590276 | 3.818704 | 3.504498 | 2.890487 | 3.472524 | 1.842721 | 0.240133 | 3.044253 |

| JNK-9L | PD-173074 | piperlongu | YK 4-279 | AP-24534 | PI-103 | Cetuximab | OSU-03012 | BX-795 | XAV 939 |
|---|---|---|---|---|---|---|---|---|---|
| NA | 4.787609 | 3.703994 | NA | NA | NA | NA | NA | 5.7039 | 5.509821 |
| -0.938 | NA | NA | NA | 1.821199 | -0.95642 | NA | 1.899682 | NA | NA |
| -0.55346 | 2.521195 | 1.688269 | 2.111051 | -0.77018 | 0.884285 | 5.96462 | 1.215319 | 0.681465 | 3.253538 |
| -0.5311 | 2.172291 | 1.326911 | 4.451779 | 2.126142 | -2.14396 | 6.86804 | 2.777007 | 1.918954 | 4.265257 |
| 0.376655 | 3.41178 | 1.8891 | 3.921407 | 2.312513 | -0.31806 | 7.334622 | 2.722441 | 4.801432 | 4.700037 |
| 0.181863 | 3.75099 | 2.474611 | 2.50065 | 1.193737 | 2.22469 | 6.686492 | 2.827933 | 3.61804 | 4.524115 |
| 0.299339 | 2.919339 | 1.93084 | 3.101283 | 2.148859 | 0.313379 | 6.902713 | 1.948533 | 3.757579 | 4.03214 |
| 1.895405 | 4.845739 | 3.805655 | 6.227111 | 3.663774 | 0.028027 | 6.695621 | 5.534807 | 4.730662 | 6.086337 |
| 0.327754 | 2.766194 | 1.515154 | 0.848933 | 1.4444 | -1.21686 | 6.229232 | 2.741296 | 1.182538 | 3.011549 |
| 0.031596 | 4.31411 | 4.545497 | 4.120854 | 2.982722 | 1.221674 | 6.969105 | 3.253287 | 3.970879 | 5.13618 |
| 2.162903 | 4.381591 | 1.646189 | 3.019059 | 2.89162 | -0.22888 | 7.526436 | 2.193505 | 4.306391 | 2.622104 |
| 2.736632 | NA | 1.768772 | 1.864271 | 2.355593 | 0.499753 | 7.335912 | 1.456718 | NA | 3.877476 |
| 0.446427 | 3.512774 | 2.23172 | NA | 1.90298 | 4.102549 | 5.448285 | 2.569253 | 5.157172 | 3.893437 |
| -0.3513 | NA | 1.852814 | 1.725506 | 1.984582 | 0.785622 | 7.545996 | 1.840563 | NA | 5.020224 |
| 0.381588 | NA | NA | NA | 2.076867 | -0.04077 | NA | 4.809792 | NA | NA |
| 2.055885 | 3.332611 | 1.052854 | NA | 2.836663 | 0.400145 | 7.036776 | 4.270798 | 4.783339 | 3.535975 |
| -0.17418 | 3.358308 | 1.133875 | 1.172244 | 1.99573 | 3.361363 | 6.179747 | 2.327666 | 3.504123 | 3.191741 |
| 0.560344 | 2.657742 | 2.303666 | 0.613259 | 0.221772 | -0.02184 | 5.092236 | 2.503469 | 2.161797 | 3.045332 |
| -0.30614 | NA | NA | NA | 1.313892 | 1.90321 | NA | 2.408915 | NA | NA |
| -1.15627 | 1.992114 | 1.469146 | 1.498999 | -0.08893 | 5.025787 | 6.509872 | 0.166764 | 2.450228 | 3.67738 |
| -0.71806 | 3.528464 | 1.546895 | NA | 1.930314 | 3.156637 | NA | 1.879458 | 3.210517 | 4.167042 |
| NA | 3.50545 | 2.003444 | 2.760535 | NA | NA | 6.707138 | 1.762575 | 5.022792 | 3.302903 |
| 0.046684 | 3.60418 | 1.803576 | NA | 2.262848 | -1.0665 | 6.934045 | 1.389069 | 3.881383 | 3.715876 |
| -0.46732 | 3.969455 | 1.212435 | 2.036492 | 2.152389 | 0.011318 | 4.343809 | 2.973814 | 4.157475 | 3.85098 |
| -1.58597 | 2.555578 | -0.01934 | 0.411372 | -0.21384 | 1.818811 | 6.011129 | -1.14196 | 0.277188 | 2.265792 |
| -0.64908 | 2.859551 | 1.248529 | 1.570377 | 1.998639 | 3.61239 | 6.176905 | 2.165298 | 3.03619 | 3.963975 |
| -1.08867 | 2.754734 | 1.249743 | 0.548604 | 0.127155 | 3.404958 | 5.374206 | 1.473585 | 2.328539 | 3.763769 |
| -0.85706 | 1.315301 | 2.786822 | 1.534797 | 2.63963 | -1.68361 | 6.504863 | 1.865003 | 2.985002 | 3.320823 |
| -0.24506 | 2.8531 | 0.899973 | 1.150922 | 1.21062 | 0.560949 | 3.262926 | 2.38835 | 0.482869 | 3.45601 |
| 0.693783 | 3.872027 | 2.241135 | 2.858375 | 0.556506 | 1.671566 | 5.447712 | 1.885759 | 3.381025 | 3.434491 |
| -0.32713 | 2.375876 | 1.508482 | 2.014975 | 0.122561 | 2.714923 | 6.662682 | 2.583569 | 0.440382 | 3.962528 |
| 1.116874 | 2.10435 | 1.587741 | 1.390712 | 2.05369 | -0.85255 | 6.430256 | 2.214842 | 3.785884 | 4.41652 |
| -1.13379 | NA | 1.334229 | 2.991041 | 1.47399 | 0.934097 | 5.621678 | 2.969636 | NA | 4.095263 |
| 2.401047 | 0.804292 | 1.777242 | 1.049827 | 0.579837 | 5.215496 | 5.431279 | 2.877632 | 1.176654 | 3.844619 |
| -0.02957 | 2.292319 | 1.754166 | 1.961077 | 1.813774 | 1.670133 | 7.180265 | 3.203218 | 2.298404 | 4.259126 |
| -2.63471 | 3.036574 | 2.233502 | 1.36467 | 2.276722 | 5.218006 | 5.978362 | 0.138459 | 2.734774 | 4.418756 |
| 0.026148 | 3.990774 | 0.691005 | 1.964103 | 2.596665 | 3.139002 | 6.902495 | 3.159428 | 3.210166 | 2.666365 |
| -0.34014 | 2.278415 | 1.419962 | -0.19666 | 1.773636 | 0.8498 | 6.507622 | 3.851266 | 1.169795 | 0.766339 |
| -0.68402 | 2.666726 | 0.826698 | 0.286883 | 0.26435 | 0.3568 | 5.325417 | 1.829899 | 1.301006 | 2.298762 |
| 0.535509 | 1.65718 | 0.907023 | 1.655311 | -0.70274 | -0.79875 | 6.588385 | 2.575128 | 1.608756 | 3.296968 |
| -0.51175 | 0.995574 | 2.809949 | 1.801346 | -0.47606 | 2.544552 | 6.3332 | 1.810206 | 3.299755 | 4.026918 |
| 0.14384 | 3.759464 | 1.891547 | 2.707559 | 2.390322 | 0.329766 | 6.127254 | 1.551405 | 3.936617 | 3.385279 |
| 0.059729 | 2.594906 | 0.608196 | 1.825202 | 0.964871 | 2.369128 | 5.995082 | 1.1604 | 1.772682 | 3.065537 |
| NA | 3.997544 | 0.65667 | 1.35037 | NA | NA | 7.162445 | NA | 2.102697 | 2.973059 |
| 0.013687 | 0.016344 | 0.018518 | 0.028952 | 0.031318 | 0.031927 | 0.032056 | 0.032295 | 0.035333 | 0.037209 |
| 0.52402 | 3.480877 | 2.17937 | 2.744416 | 1.952292 | 0.523218 | 6.654389 | 2.758357 | 3.591305 | 4.106579 |

| | | | | | | | | | |
|---|---|---|---|---|---|---|---|---|---|
| -0.3213 | 2.723498 | 1.457951 | 1.58856 | 1.209202 | 1.754413 | 6.063293 | 1.965669 | 2.502116 | 3.455803 |

| PF-562271 | Cisplatin | (5Z)-7-Oxo | Gefitinib | SN-38 | I-BET | Bryostatin | BX-912 | Olaparib | GSK-19045 |
|---|---|---|---|---|---|---|---|---|---|
| NA | 5.231838 | 3.453564 | 3.178167 | 0.912175 | NA | NA | NA | 5.7039 | NA |
| 2.199525 | NA | NA | NA | -6.16386 | 2.557942 | -2.30597 | 1.680101 | NA | 2.851873 |
| 0.782466 | 2.516856 | -2.08125 | 0.12721 | -4.55076 | 1.990062 | -4.48863 | 1.36964 | 2.964822 | 1.672763 |
| 1.313927 | 3.2593 | 1.60199 | 0.57964 | -1.26079 | 2.966649 | -2.19357 | 4.673059 | 4.110472 | 0.607605 |
| 3.355113 | 5.053287 | 2.599752 | 1.891081 | -2.03051 | 3.51772 | -1.57484 | 5.618227 | 4.344174 | 3.178425 |
| NA | 5.323404 | 2.554607 | 1.798277 | -2.46545 | 3.582308 | -2.98932 | 4.393342 | 3.926668 | 1.794169 |
| 3.003786 | 5.037177 | 2.181913 | 1.103776 | -3.69554 | 2.420157 | -2.07751 | 5.099389 | 3.223817 | 2.740959 |
| 4.932546 | 6.579369 | 1.831044 | 3.050475 | -0.39304 | 4.350356 | -0.41632 | 7.033593 | 6.336712 | 4.643651 |
| 2.826884 | 2.300026 | 1.033165 | 1.420694 | -4.65128 | 0.968587 | -2.89165 | 3.900131 | 3.723279 | 2.08341 |
| 4.509163 | 2.081667 | 3.328108 | 1.821528 | 0.466268 | 4.864732 | -1.60356 | 5.903836 | 5.288459 | 3.71926 |
| 2.795079 | 4.831335 | 2.078189 | 3.020034 | -2.98958 | 3.787999 | -3.01962 | 4.095058 | 4.8246 | 3.114878 |
| 1.440215 | NA | 2.199292 | NA | -2.14565 | 3.953649 | -2.1003 | 4.241084 | NA | 1.964253 |
| 3.207342 | 4.03393 | 1.896552 | 2.854587 | -2.91778 | 6.537287 | -2.01413 | 5.742345 | 5.157172 | 2.394196 |
| NA | 4.836621 | 2.028932 | 2.85788 | -2.92163 | 3.322145 | -4.41835 | 3.309159 | 4.048679 | 1.661685 |
| 3.464285 | NA | NA | NA | NA | 3.655914 | -2.71786 | 5.0797 | NA | 2.778427 |
| 4.295862 | 3.335112 | 1.298917 | 2.516015 | -3.29192 | 5.286219 | -1.72313 | 5.827983 | 4.939071 | 2.802522 |
| 2.372267 | 2.696525 | 1.537264 | 0.124397 | -1.90032 | 2.611188 | -2.88658 | 4.972763 | 4.057999 | 2.140955 |
| 3.237375 | 4.321389 | 2.375457 | 1.553316 | -5.03191 | 3.508861 | -2.3785 | 3.366132 | 4.461596 | 2.293921 |
| 2.015123 | NA | NA | NA | NA | 2.753456 | -2.44765 | 4.291339 | NA | 2.271076 |
| 1.137707 | 3.119642 | 1.187821 | 1.02863 | -6.23815 | 5.69418 | -3.90286 | 4.994247 | 3.193926 | 2.17257 |
| 3.05057 | 2.135266 | NA | -0.33061 | -4.75278 | 4.39388 | -2.68825 | 4.530978 | 3.856696 | 0.631467 |
| NA | 3.462991 | 2.737893 | 1.83246 | -1.76723 | 3.076802 | -2.96003 | 3.290842 | 3.656634 | 2.186926 |
| 3.856008 | 5.308559 | 0.913706 | -0.17784 | -1.54471 | 4.661485 | -1.91598 | 4.19899 | 4.47975 | 1.945236 |
| 2.227746 | 3.406734 | 1.131463 | 0.127861 | -3.36746 | 6.040444 | -2.30809 | 4.514487 | 4.873831 | 2.166279 |
| 0.459596 | 2.106029 | -0.26207 | 0.595923 | -6.08969 | 4.549189 | -4.28304 | 3.408025 | 2.378359 | 2.057661 |
| 3.39982 | 4.211371 | 1.50594 | 1.257925 | -3.51418 | 5.97823 | -3.88769 | 5.256772 | 3.877398 | 2.910936 |
| 1.719302 | NA | 1.315145 | NA | -6.24572 | 4.268179 | -3.78104 | 3.369416 | NA | 1.242122 |
| 3.099463 | 4.963899 | 1.750524 | 2.550116 | -2.81545 | 3.673603 | -3.32617 | 2.440822 | 4.947107 | 2.020151 |
| 0.850336 | 1.831789 | 0.227915 | -0.51033 | -5.90304 | 4.375705 | -4.94708 | 0.085619 | 3.539518 | 0.887086 |
| 2.208718 | 3.688831 | 0.439281 | -0.21888 | -2.41156 | 6.144067 | -3.24923 | 4.284119 | 4.792145 | 3.05562 |
| 3.639437 | NA | 1.587809 | NA | -4.44819 | 4.899137 | -3.38254 | 2.795835 | NA | 2.174391 |
| 2.362586 | 4.880537 | 1.886043 | 2.155044 | -4.35585 | 1.244565 | -1.99456 | 2.246323 | 4.497219 | 1.843668 |
| 0.508189 | 2.288884 | 1.57784 | -0.04509 | -2.70774 | 2.481503 | -4.05936 | 1.912853 | 4.080932 | 2.537613 |
| 3.81394 | 2.531229 | 1.063642 | 1.822326 | -3.2724 | 5.737388 | -2.01086 | 3.89586 | 3.840579 | 2.910971 |
| 1.417619 | 3.926546 | 1.382298 | 0.940189 | -2.82331 | 6.153162 | -3.59976 | 4.402599 | 4.545143 | 2.571314 |
| -0.50974 | 4.105288 | 1.529494 | 1.665995 | -1.74368 | 3.111689 | -5.36102 | 3.983597 | 4.458873 | 0.649039 |
| 3.490238 | 1.533546 | 0.53645 | 1.940762 | -5.20307 | 4.844691 | -2.17725 | 5.514689 | 3.694957 | 0.307983 |
| 1.937517 | 2.441117 | 1.543672 | 1.669912 | -3.33733 | 3.128707 | -2.82832 | -0.07339 | 2.148416 | 2.463241 |
| 1.981048 | 2.615458 | 0.545187 | 0.223566 | -2.86477 | 5.570133 | -3.59914 | 4.371779 | 2.945602 | 1.576236 |
| 0.645555 | 3.880586 | 0.055861 | 1.244093 | -4.36203 | 2.660354 | -1.90879 | 2.479855 | 4.544478 | 2.499183 |
| 3.412051 | 2.30756 | 2.059545 | 2.501775 | -3.20869 | 5.42906 | -2.56531 | 4.249172 | 2.905373 | 1.223475 |
| 2.30426 | 3.027183 | 1.963418 | 1.36901 | -3.04026 | 3.779169 | -2.27671 | 3.440672 | 3.566916 | 2.859113 |
| 0.357546 | 2.928112 | -0.22819 | 1.845289 | -5.00218 | 4.987353 | -2.10909 | 4.626004 | 3.3429 | 1.177055 |
| NA | 3.390256 | 0.443387 | 2.712876 | -3.72654 | NA | NA | NA | 4.860366 | NA |
| 0.03829 | 0.041537 | 0.041559 | 0.044115 | 0.044413 | 0.044888 | 0.044946 | 0.045763 | 0.046044 | 0.046192 |
| 2.915722 | 4.095856 | 1.869843 | 1.859805 | -2.64892 | 3.522457 | -2.45881 | 4.488561 | 4.474095 | 2.496644 |

| | | | | | | | | | |
|---|---|---|---|---|---|---|---|---|---|
| 2.057693 | 3.221366 | 1.120586 | 1.139174 | -3.78984 | 4.385445 | -3.10279 | 3.54046 | 3.870744 | 1.933616 |

| Midostauri | Vinorelbin | Epothilone | IPA-3 | Trametinib | Pazopanib | BMS-7548 | BAY 61-36 | KIN001-10 | JW-7-24-1 |
|---|---|---|---|---|---|---|---|---|---|
| NA | NA | NA | NA | 3.907907 | NA | NA | NA | NA | NA |
| -0.14243 | -4.99671 | -4.73721 | 4.212156 | NA | 3.932494 | 0.647959 | 1.148779 | 0.456112 | 0.355777 |
| -0.89089 | -3.15388 | -6.12673 | 3.070265 | -6.58334 | 3.421375 | -0.01638 | 0.74988 | 1.915194 | 1.1853 |
| -0.24411 | -3.88879 | -6.44712 | 4.391752 | 2.681126 | 4.543178 | -1.14051 | 2.240086 | 2.58928 | 0.445482 |
| 2.403972 | -3.20976 | -4.26745 | 5.309546 | 3.148003 | 2.51542 | 3.562362 | 2.563088 | 0.278426 | 1.304272 |
| 0.630848 | -1.73096 | -3.11524 | 6.483796 | 2.4843 | 2.680371 | 2.787622 | 5.132913 | 2.602823 | 2.004268 |
| 1.609742 | -3.84247 | -1.81519 | 4.754986 | 1.085616 | 3.166231 | 2.307829 | 3.821011 | 2.644524 | 1.340893 |
| 2.452228 | 0.294838 | 0.64183 | 7.200604 | 4.427726 | 6.414244 | 4.805036 | 6.050956 | 2.688934 | 2.438359 |
| 1.249444 | -2.50253 | -3.80399 | 4.978838 | 1.468879 | 3.850818 | 2.788086 | 3.496604 | 0.403005 | 1.50878 |
| 2.915077 | -2.49072 | -2.45743 | 4.56018 | 2.474246 | 2.729854 | 4.526719 | 4.905304 | 1.326838 | 2.586034 |
| 1.480253 | -1.51449 | -2.07905 | 3.732737 | 1.987017 | 4.315208 | 1.132843 | 3.560336 | -0.37954 | -0.4002 |
| 1.413943 | -4.41947 | -4.0142 | 4.524951 | 2.540606 | 5.407815 | 2.752889 | 4.245394 | 2.484314 | 2.000756 |
| 2.758034 | -3.93377 | -3.624 | 6.301144 | 0.48846 | 2.526502 | 3.101285 | 4.866008 | 2.628592 | 2.984989 |
| -1.9097 | -5.15749 | -5.02544 | 3.075905 | 2.439089 | 1.306493 | 2.150106 | 4.844345 | 0.210545 | 1.684403 |
| 2.075821 | -0.05708 | -1.35676 | 6.185236 | NA | 4.512387 | 3.69415 | 4.830704 | 3.115092 | 1.887878 |
| 2.756789 | -0.84771 | -3.29694 | 6.78664 | -1.23688 | 5.081244 | 4.448339 | 5.197304 | 0.761124 | 0.804296 |
| 1.839627 | -4.68376 | -4.25682 | NA | 1.476383 | 2.264301 | 1.551989 | 1.450701 | 0.810148 | 2.070056 |
| -0.77899 | -5.24977 | -5.67 | 5.70463 | 1.14597 | 4.306564 | 1.32688 | 3.974566 | 2.509642 | 1.523724 |
| -0.14554 | -4.38193 | -5.15059 | 5.418199 | NA | 3.993312 | 1.249043 | 3.18646 | 2.444577 | 2.086841 |
| -0.89767 | -5.04208 | -7.94039 | 1.202687 | 2.37559 | 2.811065 | -2.33244 | 0.150967 | 5.025787 | 5.002746 |
| 0.78833 | -5.54595 | -5.65883 | 2.774839 | NA | 2.429474 | 2.722045 | 1.961254 | 1.288038 | 1.862823 |
| NA | NA | -3.62612 | 4.910921 | 2.688847 | 1.958305 | 2.858412 | 1.957558 | 1.47201 | NA |
| 2.284232 | -3.32961 | -4.25465 | 2.152238 | 2.865382 | 3.655207 | 3.190994 | 2.413563 | -0.05838 | 2.261426 |
| 0.844491 | -4.1076 | -5.2181 | 6.67437 | -0.75056 | 3.818017 | 1.904965 | 4.922946 | 2.286607 | 1.005325 |
| -2.30819 | -4.73884 | -6.53411 | 1.252308 | 0.546741 | 2.017476 | -0.85717 | 1.247448 | 3.131692 | 2.129099 |
| 2.075844 | -4.73424 | -4.69911 | 3.285981 | -1.80299 | 3.310153 | 3.647489 | 3.711561 | 1.210711 | 0.551362 |
| 0.141963 | -5.42332 | -6.5739 | 3.419227 | 0.398291 | 3.863923 | 2.361713 | 2.420504 | 2.55085 | 1.317001 |
| 0.199283 | -5.15239 | -4.63472 | 4.97628 | -0.67939 | 2.449736 | 1.54669 | 1.961914 | 0.370328 | 1.081228 |
| -1.24321 | -5.37744 | -6.23354 | 5.829653 | 0.100559 | 3.189537 | 0.248932 | 4.44898 | 2.217929 | 1.365689 |
| 0.268912 | -3.40812 | -4.59206 | 3.766438 | 0.390292 | 4.30568 | 1.774849 | 2.504765 | 4.516814 | 2.917543 |
| -1.33725 | -2.76805 | -1.75164 | 5.449754 | 2.681984 | 3.203466 | 1.478456 | 4.840384 | 2.987451 | 2.717674 |
| 2.128894 | -2.75133 | -2.82661 | 4.146647 | 0.740011 | 3.735937 | 2.184215 | 5.020102 | -0.19831 | 1.509786 |
| 0.327094 | -4.59125 | -5.39732 | 6.05066 | -0.52183 | 2.882352 | -0.33781 | 0.898401 | 0.333848 | 1.629414 |
| 0.530427 | -0.38199 | -2.99062 | 4.587478 | -0.59095 | 2.345194 | 1.829729 | 3.636253 | 5.218975 | 5.217906 |
| 2.493454 | -1.96981 | -3.03242 | 3.638469 | -2.6093 | 3.400361 | 0.926384 | 4.252522 | 2.737528 | 1.113575 |
| -1.22195 | -5.15673 | -5.77797 | 2.630944 | 1.54984 | 0.436813 | -1.85391 | -0.64234 | 1.757487 | 4.999977 |
| 2.478773 | -3.3037 | -4.04053 | 5.981389 | -0.67969 | 4.369952 | 3.267049 | 3.975247 | 4.020539 | 3.095327 |
| -0.09466 | -5.04887 | -4.31801 | 3.94661 | 1.795029 | 3.01523 | 0.589311 | 2.346433 | 1.568754 | 1.45652 |
| 0.361242 | -5.34738 | -7.10612 | 3.332216 | -1.60126 | 2.124655 | 0.932177 | 1.506473 | 3.287061 | 1.833053 |
| -1.41118 | -3.42406 | -4.37887 | 6.426475 | -1.49568 | 3.028693 | 0.982828 | 2.325654 | 2.969804 | 1.166043 |
| -1.30941 | -4.98083 | -2.74682 | 5.516876 | -0.12214 | 0.873905 | 2.930613 | 1.863746 | 2.819224 | 2.166094 |
| 1.803888 | -2.42298 | -3.45671 | 3.104516 | 0.024956 | 3.850392 | 3.109904 | 3.765882 | 1.869097 | 1.001303 |
| -1.38864 | -3.51626 | -4.1886 | 4.732643 | -2.80238 | 3.577239 | 0.955915 | 4.836068 | 2.9665 | 1.971958 |
| NA | NA | NA | NA | -0.38889 | NA | NA | NA | NA | NA |
| 0.051139 | 0.05385 | 0.054633 | 0.056934 | 0.061153 | 0.064404 | 0.065554 | 0.065939 | 0.066009 | 0.068976 |
| 1.154097 | -3.02262 | -3.61481 | 5.079585 | 1.495944 | 3.704382 | 2.378071 | 3.710469 | 1.590885 | 1.513239 |

| | | | | | | | | | |
|---|---|---|---|---|---|---|---|---|---|
| 0.223714 | -4.0377 | -4.68513 | 4.208313 | 0.088019 | 2.985843 | 1.412415 | 2.78051 | 2.351797 | 2.144155 |

| Thapsigarg | Obatoclax | LAQ824 | ZM-44743! | XL-880 | GSK212645 | AZD6244 | CGP-0829! | Bosutinib | rTRAIL |
|---|---|---|---|---|---|---|---|---|---|
| NA | NA | NA | 5.480756 | NA | NA | 5.480756 | NA | 2.656999 | 1.500469 |
| -2.13324 | -1.17849 | -3.91763 | NA | 0.217238 | -4.00231 | NA | 3.060186 | NA | NA |
| -5.11575 | -1.78434 | -4.14376 | 1.562851 | -0.49023 | -3.1499 | 2.454812 | 2.128681 | 1.72868 | -1.15679 |
| -5.31169 | 0.135899 | -3.88531 | 1.009134 | 4.431127 | -3.63716 | 3.44023 | 3.564874 | 2.590433 | -2.04108 |
| -2.8268 | 0.871306 | -4.06745 | 4.568157 | 1.857114 | -4.49834 | 4.394458 | 3.238629 | 2.892677 | -1.19175 |
| -4.89544 | -1.0485 | -4.34486 | 4.311142 | 4.518124 | -1.3743 | 3.871579 | NA | 2.722894 | 0.443602 |
| -5.08611 | -0.60592 | -3.13865 | 3.763563 | 2.279205 | -3.06323 | 3.484443 | NA | 1.9985 | -0.06873 |
| 2.683596 | 1.248586 | -1.96697 | 5.36306 | 5.268187 | -2.72276 | 4.217242 | NA | 4.967891 | 2.174314 |
| -2.46336 | -0.26642 | -2.81172 | 1.359806 | 0.713231 | -4.01734 | 3.464542 | 1.775457 | 2.472247 | -0.28626 |
| -0.13376 | 1.253129 | -3.32931 | 3.55059 | 5.109895 | -3.52109 | 2.808105 | NA | 4.193449 | 1.226803 |
| 0.073659 | 0.185653 | -3.5759 | 3.072209 | 2.047729 | -6.37929 | 4.425886 | NA | 3.743738 | 1.134091 |
| -4.27621 | -0.9229 | -3.14377 | NA | 2.735684 | -2.72756 | NA | NA | NA | -0.8235 |
| -2.03654 | 0.908128 | -1.98248 | 4.934029 | 5.180889 | -1.27341 | 4.934029 | NA | 4.240882 | 0.646657 |
| NA | -2.23457 | -3.47334 | NA | 0.69211 | -3.21534 | NA | NA | 2.898111 | 1.131631 |
| 1.931399 | 2.597472 | 0.492632 | NA | 1.63411 | -2.59163 | NA | NA | NA | NA |
| -2.33625 | 1.115656 | -2.1538 | 3.290221 | 3.018819 | -4.40351 | NA | NA | 2.58059 | 0.361799 |
| -5.72532 | -0.26727 | -3.75082 | 2.686139 | 2.635083 | -5.31429 | 3.102471 | NA | 1.399532 | -0.36964 |
| -5.03841 | 0.811372 | -3.6602 | 3.158159 | -0.30959 | -2.81245 | 4.159882 | NA | 3.5632 | -0.1625 |
| -0.42917 | 0.116684 | -3.17141 | NA | 0.57152 | -2.70315 | NA | 3.227284 | NA | NA |
| -7.54894 | -4.45886 | -3.7321 | 2.883385 | 4.33264 | 0.64376 | 3.736568 | NA | 3.165394 | 0.279185 |
| -4.74173 | -0.88955 | -3.94866 | 2.760714 | 1.677758 | -3.24442 | 4.221612 | NA | 2.632323 | 0.341073 |
| -3.71359 | -0.30854 | -3.8313 | 4.93871 | 1.791367 | NA | 4.845031 | NA | 4.270215 | 0.692748 |
| -1.91729 | -1.3882 | -3.11027 | 4.027359 | 1.228759 | -5.01855 | 4.224651 | 3.682369 | 3.210853 | 0.57111 |
| -1.82405 | 1.282379 | -1.16918 | 2.754761 | 2.752267 | -3.9266 | 2.870975 | NA | 2.792236 | -0.09795 |
| -5.631 | -2.59965 | -2.767 | 0.542423 | 1.351984 | -2.30461 | 3.02632 | NA | 2.816245 | -0.31175 |
| -4.19703 | 0.809731 | -2.27477 | 2.545258 | 4.098559 | -5.1764 | 1.991434 | NA | 1.829475 | 0.48992 |
| -5.05049 | -2.14404 | -2.41035 | 1.472728 | 0.592483 | -2.50046 | 1.762746 | NA | NA | -4.16736 |
| -5.28083 | -2.79105 | -3.12194 | 3.206764 | 0.599672 | -4.81334 | 4.010753 | NA | 3.80076 | 0.643499 |
| -5.77074 | -0.06454 | -1.71324 | 1.641224 | 0.584112 | -3.80542 | 2.700076 | NA | 1.547244 | -1.19018 |
| -6.12274 | 0.366343 | -1.21646 | 3.143651 | 1.006063 | -2.3313 | 1.377238 | NA | 1.415012 | -0.858 |
| -0.32256 | -0.12634 | -4.0994 | 0.924759 | 0.865793 | -0.81243 | 3.176425 | NA | NA | -1.74202 |
| -0.60006 | 0.466845 | -3.02268 | 3.729705 | 1.941375 | -5.32867 | 4.282518 | NA | 2.850938 | 0.502662 |
| -3.71768 | 0.066243 | -2.76868 | NA | 0.966941 | -3.79764 | NA | NA | 3.194034 | -1.84244 |
| NA | -1.15925 | 2.768001 | NA | 0.724067 | 0.743892 | 4.174237 | NA | 2.562327 | 0.473654 |
| NA | -1.51049 | -3.20572 | 2.275664 | -0.12677 | -2.452 | 4.240978 | NA | 3.016 | 0.943282 |
| -5.8761 | -2.04845 | -3.47357 | NA | 2.277107 | -1.57383 | 2.603273 | NA | 2.403321 | 0.510539 |
| -4.99304 | 0.058235 | -0.825 | NA | 2.019785 | -0.45294 | 4.693503 | NA | 2.392762 | -2.14159 |
| -1.52222 | 0.76214 | 0.652347 | 0.81349 | -0.23737 | -3.48295 | 2.608428 | 3.610325 | 1.865993 | -1.68965 |
| -5.08034 | -0.94566 | -2.95894 | NA | 1.4883 | -3.28904 | 3.257606 | NA | 0.57157 | -0.43037 |
| -3.36416 | -1.48423 | -2.39339 | 1.39531 | 0.243846 | -3.8235 | 2.23605 | NA | 2.711429 | -1.57557 |
| -5.02913 | 0.837056 | -1.97923 | 2.942808 | 0.845742 | 0.933966 | 3.847504 | NA | 2.196235 | -2.06341 |
| -7.2067 | -0.46614 | -2.64445 | 4.148706 | 1.487981 | -2.74082 | 4.14023 | NA | 0.644639 | -0.07163 |
| -6.00022 | -0.13306 | -2.62992 | 2.450001 | 4.101236 | -1.27819 | 1.144875 | NA | 2.615692 | -0.47689 |
| NA | NA | NA | 3.268785 | NA | NA | 4.180838 | NA | 1.305018 | 0.900967 |
| 0.070695 | 0.071947 | 0.082062 | 0.082706 | 0.083374 | 0.089742 | 0.093223 | 0.09526 | 0.09545 | 0.09648 |
| -2.66814 | 0.048164 | -3.10902 | 3.436415 | 2.443454 | -3.45317 | 3.864495 | 2.753565 | 2.976655 | 0.157446 |

| | | | | | | | | | |
|---|---|---|---|---|---|---|---|---|---|
| -4.1713 | -0.7101 | -2.36189 | 2.59331 | 1.487409 | -2.60561 | 3.306411 | 3.50666 | 2.426509 | -0.49241 |

| CCT01815 | HG-5-113- | HG-6-64-1 | FH535 | KIN001-24 | OSI-906 | Docetaxel | AZD-0530 | SB 216763 | JNJ-26854 |
|---|---|---|---|---|---|---|---|---|---|
| 3.918395 | 3.542845 | NA | NA | NA | NA | -3.83588 | NA | 4.404266 | 4.681742 |
| NA | NA | 1.79406 | 2.55494 | 1.695807 | 3.661304 | NA | 1.851124 | NA | NA |
| 2.572675 | NA | 0.314659 | 1.863958 | 2.646711 | 1.366059 | -4.09655 | 2.540592 | 4.201356 | 1.909191 |
| 1.770152 | 1.472246 | 1.608242 | 2.309332 | 4.265847 | 1.689439 | -5.48222 | 3.166796 | 5.099654 | 2.772485 |
| 2.453471 | 2.130509 | 3.478511 | 1.384061 | 2.615076 | 4.250691 | -2.17493 | 2.44106 | 4.562455 | 4.504446 |
| 3.283986 | 3.276492 | 3.801642 | 1.673764 | 4.303559 | 3.135243 | -3.65992 | NA | 5.261353 | 3.598651 |
| 5.053628 | NA | 1.350589 | 0.228277 | 3.667181 | 3.396382 | -2.40918 | 3.259376 | 4.733403 | 3.753397 |
| 6.816611 | 5.932732 | 6.056788 | 6.648677 | 4.224653 | 4.305548 | 0.252336 | NA | 7.029859 | 3.897835 |
| 2.440111 | 1.59133 | 3.209683 | 2.639138 | 1.497584 | 3.053821 | -6.13827 | 1.95984 | NA | 2.781531 |
| 5.210782 | 4.24451 | 3.54544 | 1.462474 | 4.628305 | 4.401204 | -2.19718 | NA | 6.020256 | 2.838202 |
| 3.664499 | NA | 3.461654 | 3.062713 | 2.172093 | 3.794077 | -5.46532 | NA | 5.088006 | 2.539164 |
| 4.295374 | NA | -0.38787 | 0.086225 | 3.291395 | 3.558849 | NA | NA | NA | 2.84018 |
| NA | 3.852632 | 1.914388 | 1.611409 | 3.903246 | 4.136907 | -5.62306 | NA | NA | NA |
| 4.903527 | NA | 0.668381 | 1.74551 | 2.601435 | 2.132381 | -6.62905 | NA | 5.965988 | 4.513746 |
| NA | NA | 3.949226 | 3.371274 | 3.635612 | 3.694717 | NA | 1.244721 | NA | NA |
| 4.092588 | 4.328073 | 4.991147 | 0.869543 | 3.687774 | 4.052268 | -3.42996 | NA | 5.524027 | 2.282531 |
| 2.775558 | NA | 1.49365 | 0.993528 | 3.558056 | 3.032584 | -5.49929 | NA | 4.804017 | 1.83097 |
| 3.516382 | 2.270182 | 1.501389 | 2.374012 | 3.339724 | 2.973036 | -4.80044 | NA | 5.169576 | 2.43479 |
| NA | NA | 2.829438 | 2.147825 | 2.937106 | 3.320672 | NA | 2.906149 | NA | NA |
| 2.128981 | NA | 0.949836 | 1.021694 | 4.894026 | 0.448191 | -6.40977 | NA | 2.906915 | 2.704935 |
| 1.70806 | 1.671553 | 1.99253 | 0.104195 | 2.917469 | 2.789401 | -6.02519 | NA | 4.18593 | 1.846832 |
| 5.671195 | 3.039057 | NA | 0.610859 | 3.471968 | 2.326066 | -1.47089 | NA | 5.879653 | 4.465345 |
| 2.213715 | 2.771177 | 2.737666 | 1.669397 | 3.117458 | 3.911251 | -3.83553 | 2.888081 | 5.00926 | 3.061318 |
| 2.93279 | 2.728544 | 2.751498 | 2.101645 | 4.391499 | 3.816043 | -5.56278 | NA | 5.55528 | 4.02703 |
| 1.400475 | NA | -1.13169 | 1.208764 | 4.963725 | 2.739712 | -5.6823 | NA | 3.070165 | 2.247961 |
| 2.564539 | 2.708636 | 4.047373 | 0.778678 | 5.92171 | 3.929662 | -4.26491 | NA | 5.362344 | 2.222434 |
| 2.641337 | NA | 0.473206 | 2.977856 | 3.479389 | 2.051525 | NA | NA | NA | 1.711343 |
| 2.58948 | 2.633306 | 0.686756 | 1.407174 | 3.992307 | 3.752135 | -2.19152 | NA | 4.909582 | 2.504819 |
| 3.232116 | 0.551386 | 2.528249 | 0.30526 | 2.475908 | 3.325888 | -5.16536 | NA | 4.760217 | 2.68385 |
| 3.028873 | NA | 0.992723 | 2.238043 | 4.24065 | 0.772633 | -4.65038 | NA | 5.48798 | 4.061459 |
| 3.015463 | NA | 0.993576 | 1.945519 | 4.760607 | 3.342162 | NA | NA | NA | 2.614476 |
| NA | NA | 3.631269 | 2.206704 | 2.82643 | 3.691203 | -5.378 | NA | 4.974592 | NA |
| 3.338919 | NA | 2.160284 | 1.701771 | 2.489552 | 3.52281 | -6.07003 | NA | 4.840127 | 2.666907 |
| 3.745105 | 2.693497 | 2.121468 | 2.069653 | 5.020028 | 3.112799 | -5.61662 | NA | 4.684878 | 2.515175 |
| 4.443201 | NA | NA | 0.378993 | 4.172925 | 0.641903 | -5.46326 | NA | NA | 3.198552 |
| 3.60148 | 2.117514 | 1.333259 | -0.9453 | 3.715569 | 1.213337 | -4.00633 | NA | 4.858717 | 3.108527 |
| 4.994376 | NA | 1.898484 | 0.09397 | 3.145345 | 3.862595 | -4.77482 | NA | NA | 2.629314 |
| 2.22274 | NA | 3.703008 | 2.545915 | 3.147812 | 2.095443 | -6.60204 | 2.797581 | 4.665233 | 2.768269 |
| 3.221702 | 2.089646 | -0.63374 | 1.127993 | 3.542556 | 3.232956 | -6.96885 | NA | 4.36378 | 2.317601 |
| 3.00285 | NA | -0.00976 | 2.036691 | 2.507093 | 3.183724 | -5.54507 | NA | 5.234254 | 2.535295 |
| 4.725347 | 2.798055 | 0.299582 | -0.58874 | 3.415816 | 3.872751 | -3.89421 | NA | 5.245707 | 2.605861 |
| 3.338595 | 2.996711 | 3.318293 | 2.044354 | 3.663978 | 3.32569 | -4.40804 | NA | NA | 2.755637 |
| 1.236454 | NA | 0.373792 | 2.595457 | 4.809667 | 2.378074 | -5.16325 | NA | 3.495015 | 1.411001 |
| 3.016594 | NA | NA | NA | NA | NA | -5.83927 | NA | 5.726514 | 1.566562 |
| 0.099364 | 0.102334 | 0.103794 | 0.104776 | 0.107541 | 0.107727 | 0.111103 | 0.112012 | 0.114303 | 0.115563 |
| 3.784516 | 3.264155 | 2.514799 | 2.051696 | 3.278474 | 3.331442 | -4.07926 | 2.35193 | 5.220324 | 3.145258 |

| | | | | | | | | | |
|---|---|---|---|---|---|---|---|---|---|
| 3.083933 | 2.399924 | 1.654222 | 1.351375 | 3.760824 | 2.826345 | -4.9995 | 2.863937 | 4.760807 | 2.676271 |

| MLN4924 | BMS-70816 | Tipifarnib | ABT-888 | CEP-701 | RDEA119 (( | CMK | QL-XII-61 | Cyclopamir | JW-7-52-1 |
|---|---|---|---|---|---|---|---|---|---|
| NA | 5.599381 | NA | 5.695832 | 1.946481 | NA | NA | 6.210492 | NA | NA |
| NA | NA | 2.104874 | NA | NA | NA | 0.592935 | NA | 4.033746 | -2.89888 |
| -2.613 | 3.361825 | 0.694831 | 3.508208 | -1.67239 | -2.66685 | 2.579957 | NA | 4.667482 | -2.3126 |
| 2.481303 | 3.679676 | -0.78586 | 4.407411 | -0.96178 | 4.278093 | 3.066445 | NA | 4.767863 | -1.51082 |
| 2.910458 | 4.670892 | 2.314874 | 4.777435 | 2.905273 | 4.51629 | 2.24144 | NA | 4.486272 | -1.3157 |
| -2.88626 | 4.466691 | 2.800532 | 4.60205 | 2.959007 | 4.522936 | NA | 4.65419 | NA | NA |
| 2.716094 | 4.057597 | 1.744822 | 4.003889 | 2.337575 | 2.830235 | NA | NA | NA | NA |
| 4.476899 | 6.179653 | 6.89163 | 6.336712 | 2.796737 | 5.863028 | NA | 6.174516 | NA | NA |
| NA | 3.172789 | 2.281768 | 3.723257 | -0.60253 | 3.233239 | 0.720767 | 2.077513 | 2.555326 | -4.45874 |
| NA | 3.138265 | 1.692912 | 4.143298 | 3.582932 | 4.329572 | NA | 5.117352 | NA | NA |
| 2.199484 | 3.208302 | 2.478212 | 5.270697 | 1.535883 | 4.376772 | NA | NA | NA | NA |
| 0.423758 | 4.168237 | 3.148643 | NA | NA | 4.531278 | NA | NA | NA | NA |
| NA | 3.368235 | 1.511306 | 5.156107 | 1.167005 | 2.725961 | NA | 5.374866 | NA | NA |
| 3.040382 | 4.906138 | 0.423744 | 5.257239 | -1.78106 | 5.054819 | NA | NA | NA | NA |
| NA | NA | 4.900214 | NA | NA | NA | NA | NA | 5.526151 | NA |
| NA | 5.071027 | 1.886254 | 4.951282 | 2.841741 | 2.130337 | NA | 5.633429 | NA | NA |
| -0.29369 | 2.593747 | 0.690137 | 3.742485 | -0.49785 | 2.717748 | NA | NA | NA | NA |
| -0.37193 | 2.032903 | 1.803873 | 4.44312 | 0.740594 | 3.661828 | NA | 4.046917 | NA | NA |
| NA | NA | 0.170803 | NA | NA | NA | 2.39832 | NA | 5.021899 | -2.10056 |
| -1.29545 | 3.214564 | -1.62865 | 4.308923 | 0.151008 | 3.775071 | NA | NA | NA | NA |
| NA | 1.790222 | 0.490158 | 4.444755 | 0.22069 | NA | NA | NA | NA | NA |
| 1.386132 | 4.469956 | 0.793659 | 4.84285 | 3.676188 | 4.407459 | NA | 4.810359 | NA | NA |
| NA | 2.41159 | 3.07137 | 4.615412 | 0.508437 | 4.118347 | 4.292005 | 0.570398 | 5.568235 | -1.14216 |
| 2.305944 | 3.197845 | 1.958622 | 4.220982 | 0.490009 | 3.398638 | NA | 5.414475 | NA | NA |
| -1.54572 | 3.429015 | 0.739831 | 3.279957 | -1.34564 | 2.754388 | NA | NA | NA | NA |
| NA | 3.336689 | 1.686713 | 4.591148 | 1.673511 | 1.685757 | NA | 4.867473 | NA | NA |
| -3.50973 | 3.908938 | 2.265627 | NA | NA | 2.48813 | NA | NA | NA | NA |
| -1.58073 | 2.724011 | 0.86475 | 4.895682 | 2.420272 | 2.179113 | NA | 4.413981 | NA | NA |
| -0.99383 | 3.020614 | 2.035276 | 4.06707 | -0.703 | 2.320777 | NA | 4.540751 | NA | NA |
| 1.550549 | 4.001189 | 1.986907 | 4.789686 | -0.15531 | 2.97531 | NA | NA | NA | NA |
| -0.49993 | 1.805924 | 2.069291 | NA | NA | 4.123087 | NA | NA | NA | NA |
| 0.865289 | 2.435318 | 3.649794 | 4.550098 | 0.473524 | 3.081593 | NA | NA | NA | NA |
| -1.34044 | 2.747917 | 0.086372 | 3.080514 | 0.484247 | 1.623372 | NA | NA | NA | NA |
| -0.02142 | 4.502111 | 2.447402 | 4.481857 | -0.13238 | 1.402852 | NA | 4.81176 | NA | NA |
| 0.982418 | 3.879385 | 2.225496 | 5.076394 | 0.342908 | 0.70186 | NA | NA | NA | NA |
| 1.329763 | 3.828285 | -0.59248 | 3.928542 | 1.570387 | 4.231164 | NA | 2.593285 | NA | NA |
| 0.133281 | 3.774845 | 1.620049 | 4.916647 | 0.228733 | 2.302148 | NA | NA | NA | NA |
| -1.50888 | 2.41319 | 1.644382 | 3.723075 | -1.51661 | NA | 2.902331 | NA | 4.785403 | -0.84077 |
| NA | 4.217604 | -0.61262 | 3.722447 | 0.265137 | 1.337081 | NA | 2.898506 | NA | NA |
| -0.44625 | 4.744778 | 1.960682 | 4.805004 | -1.31539 | 2.302913 | NA | NA | NA | NA |
| NA | 4.289936 | 2.024524 | 3.97146 | 0.824123 | 2.270231 | NA | 4.545949 | NA | NA |
| 0.193685 | 3.295456 | 1.723601 | 3.751069 | 0.658549 | 2.608789 | NA | 3.429358 | NA | NA |
| 0.087059 | 4.100888 | 1.776776 | 3.69588 | -0.52073 | 0.731456 | NA | NA | NA | NA |
| -0.39846 | 4.814345 | NA | 4.962313 | -1.28433 | 2.9555 | NA | NA | NA | NA |
| 0.119966 | 0.12303 | 0.123973 | 0.125532 | 0.129035 | 0.133285 | 0.134198 | 0.13524 | 0.138428 | 0.146197 |
| 1.0985 | 3.97971 | 2.151928 | 4.667935 | 1.153175 | 3.473686 | 1.840309 | 4.911159 | 4.339473 | -2.49935 |

| | | | | | | | | | |
|---|---|---|---|---|---|---|---|---|---|
| -0.21534 | 3.454185 | 1.378333 | 4.292251 | 0.304971 | 2.598915 | 3.197552 | 3.899663 | 5.125179 | -1.36116 |

| JQ12 | 681640 | DMOG | SL 0101-1 | ABT-869 | ATRA | Gemcitabir | Vismodegil | OSI-027 | Embelin |
|---|---|---|---|---|---|---|---|---|---|
| NA | 4.787609 | NA | 6.326907 | NA | 5.913542 | NA | 6.397047 | NA | NA |
| 0.658535 | NA | 4.800609 | NA | 2.853722 | NA | -3.09324 | NA | -0.71624 | 2.236298 |
| 1.157995 | 1.509024 | 6.805193 | 3.001991 | 1.711145 | 3.413325 | -5.01156 | 3.734374 | 1.086309 | 2.245511 |
| 0.582376 | 3.49112 | 8.302976 | 4.701431 | 2.821689 | 2.335241 | 0.195478 | 3.242529 | 1.052841 | 3.772285 |
| 2.017919 | 3.552097 | 9.81835 | 5.563018 | 2.953605 | 4.596423 | 1.311052 | 5.407425 | 0.680917 | 2.845983 |
| -0.04119 | 2.591429 | 9.89317 | 4.979725 | 2.594997 | 2.489689 | 2.550824 | 5.36002 | 2.014411 | 3.488467 |
| 1.671681 | 3.08015 | 5.078331 | 4.694202 | 2.024091 | 2.615885 | 0.037159 | 4.20701 | -1.29442 | 3.859959 |
| 3.424973 | 3.291616 | 10.67158 | 4.779451 | 3.99531 | 4.663379 | 4.509703 | 6.319405 | 0.340746 | 5.373178 |
| 2.160317 | 2.192616 | 7.755202 | 4.416426 | 2.137558 | 2.814561 | 0.895603 | 4.416426 | 1.419357 | 3.922125 |
| -0.24063 | 1.80064 | 8.948995 | 3.669713 | 3.196481 | 2.011709 | 2.528603 | 4.785616 | 0.941729 | 3.117759 |
| 1.219361 | 4.220548 | 9.792483 | 5.90854 | 3.498132 | 4.67825 | 0.794021 | 5.744801 | -2.17417 | 5.508252 |
| 0.32941 | NA | 7.870519 | NA | 2.743201 | NA | -1.11814 | NA | 1.422614 | 1.840056 |
| 1.465682 | NA | 7.810074 | 5.819851 | 3.571451 | 5.035765 | 1.726412 | 5.85032 | 4.818514 | 3.42389 |
| -1.43618 | NA | 6.804465 | NA | 2.559363 | 3.821927 | 2.806349 | 5.524393 | 2.290799 | 1.828539 |
| 4.365008 | NA | 7.90334 | NA | 2.770801 | NA | 1.534162 | NA | 3.627549 | 3.620326 |
| 2.370684 | 3.351804 | 9.141169 | 5.810917 | 3.390467 | 3.708794 | 1.340783 | 5.360204 | -0.16867 | 3.575904 |
| -1.11214 | 2.152506 | 7.296818 | 4.903945 | 2.681425 | 3.792885 | -1.17001 | 4.585691 | 1.688642 | 2.40523 |
| 1.891398 | 1.417337 | 6.837124 | 4.734139 | 2.844467 | 5.130739 | -3.63992 | 4.62168 | 0.182161 | 2.927231 |
| 1.965852 | NA | 7.855056 | NA | 2.246354 | NA | -3.36089 | NA | -0.09429 | 3.884631 |
| -1.12287 | 1.181031 | 7.941586 | 4.270219 | 2.385033 | 2.808433 | -5.95855 | 4.99363 | 5.607905 | 1.989126 |
| 0.588562 | 3.295951 | NA | 5.137902 | 2.367553 | 2.453498 | -3.49658 | 5.137902 | 2.014428 | 2.512362 |
| NA | 4.212394 | NA | 5.444927 | NA | 5.879653 | 1.477012 | 4.041832 | NA | 2.559318 |
| 2.655405 | 3.432221 | 9.676272 | 5.308559 | 2.864646 | 5.063419 | 2.384115 | 5.173205 | -0.14762 | 2.82829 |
| 4.486033 | 2.167905 | 6.67233 | 5.300326 | 2.906387 | 4.608966 | -1.68455 | 3.65304 | 1.426915 | 2.820834 |
| 0.524545 | 1.394004 | 5.39711 | 2.777539 | 2.146779 | 2.877054 | -1.56469 | 3.017803 | 2.678976 | 1.604846 |
| 0.609689 | 0.857607 | 6.231704 | 1.904346 | 2.031507 | 4.507808 | 0.432497 | 4.653538 | -0.06372 | 2.73102 |
| 1.424879 | 0.525068 | 7.320565 | 3.112376 | 2.347617 | NA | -5.32407 | NA | 1.925877 | 3.060597 |
| 2.678611 | 0.797343 | 7.728156 | 4.724005 | 2.642446 | 4.229809 | -2.4248 | 4.368371 | -0.31203 | 2.504025 |
| 2.816709 | NA | 8.497674 | 4.116967 | 2.453384 | 4.394096 | -4.16905 | 4.348647 | -0.47723 | 3.049108 |
| 4.575116 | 2.409198 | 6.404601 | 4.598145 | 2.685941 | 5.48798 | 0.022364 | 5.429744 | 2.79892 | 2.064274 |
| -0.21974 | 0.741186 | 6.249964 | 3.216069 | 2.658272 | NA | 2.265695 | NA | 1.442087 | 2.144695 |
| 3.137281 | 3.375071 | 10.16491 | 4.648378 | 2.277055 | 5.243246 | 1.846299 | 5.173619 | -1.29671 | 5.7155 |
| 0.897811 | NA | 5.957088 | NA | 2.296925 | 4.35777 | NA | 3.338456 | 1.873987 | 4.425117 |
| 4.388568 | NA | 7.584461 | 3.875313 | 2.912099 | 2.591386 | 2.620321 | 4.614078 | 5.888406 | 2.537302 |
| 0.518362 | 4.188302 | 9.305967 | 5.552988 | 3.278764 | 5.759617 | 1.71759 | 4.359712 | 2.020064 | 2.240429 |
| -0.10398 | NA | 4.421241 | 4.647387 | 2.691097 | 1.75536 | -5.36974 | 4.815721 | 3.728964 | 1.848054 |
| 2.397083 | NA | 6.964447 | 5.609794 | 3.193624 | 5.609794 | -0.30064 | 5.609794 | 4.743981 | 2.13356 |
| 3.340449 | 2.504495 | 8.490384 | 4.052662 | 1.973638 | 4.319965 | 2.305829 | 4.453697 | 1.181945 | 3.698391 |
| 0.875417 | NA | 6.532645 | 4.437302 | 2.795194 | 4.195956 | -3.52194 | 4.410526 | 1.288736 | 2.63515 |
| 2.870613 | 1.559796 | 6.577786 | 3.723394 | 3.199276 | 4.588209 | -0.82042 | 5.515466 | -1.18433 | 3.415464 |
| 2.937933 | 2.31913 | 8.209999 | 5.50421 | 3.147182 | 5.476998 | 2.685291 | 5.378882 | 4.406863 | 3.397197 |
| 3.709914 | 3.75321 | 6.703118 | 5.261733 | 2.690671 | 5.159597 | 3.08421 | 4.691495 | 2.079166 | 2.486121 |
| 0.335558 | 1.558842 | 5.854509 | 4.7704 | 2.467907 | 3.403775 | -3.52702 | 3.719444 | 2.244415 | 3.051471 |
| NA | 3.207555 | NA | 5.726514 | NA | 5.711773 | NA | 5.67846 | NA | NA |
| 0.150386 | 0.151827 | 0.153487 | 0.153778 | 0.157032 | 0.159559 | 0.159672 | 0.163034 | 0.169818 | 0.170384 |
| 1.205012 | 2.879884 | 7.972376 | 4.950733 | 2.843994 | 3.801474 | 0.364545 | 5.037129 | 1.012535 | 3.293588 |

| | | | | | | | | | |
|---|---|---|---|---|---|---|---|---|---|
| 1.928659 | 2.288437 | 7.249634 | 4.488394 | 2.610806 | 4.368877 | -0.86174 | 4.633785 | 1.823988 | 2.853475 |

| JQ1 | MG-132 | BMS-5097 | EX-527 | THZ-2-49 | GSK-65039 | GSK26996 | ZSTK474 | Axitinib | RDEA119 |
|---|---|---|---|---|---|---|---|---|---|
| NA | NA | NA | NA | NA | NA | 3.725545 | NA | 4.337563 | 5.176746 |
| 0.355273 | 2.539951 | 3.803622 | 5.766681 | 1.181387 | 3.468804 | NA | 0.018692 | NA | NA |
| -1.05208 | -1.87229 | 2.739073 | 4.314186 | 2.626697 | 2.790786 | 1.991969 | 1.524103 | 1.163854 | -3.06867 |
| -1.38014 | -1.28936 | 4.078939 | 5.752085 | 2.918373 | 2.388889 | 4.207025 | 1.444755 | 1.163471 | 2.904582 |
| 0.475235 | 1.819109 | 3.523075 | 5.982479 | 2.134374 | 6.009931 | 1.351668 | 0.245591 | 3.545302 | 4.020655 |
| -1.589 | NA | NA | 6.030451 | 3.076633 | 5.306985 | 4.549292 | 2.927051 | 2.909833 | 4.580163 |
| 0.390088 | 1.297333 | NA | 5.633608 | 1.863802 | 5.421689 | 1.106155 | -0.35298 | 2.738465 | 2.286524 |
| 1.713639 | NA | NA | 6.974184 | 5.020603 | 7.281311 | 6.086337 | -1.03472 | 5.105473 | 4.584542 |
| -0.52487 | -1.41645 | 2.80703 | 4.799248 | 0.398588 | 4.244858 | 1.939491 | -0.60044 | 1.367242 | 3.229515 |
| -0.40986 | NA | NA | 6.713403 | 3.961187 | 6.48338 | 4.436697 | 0.881687 | 1.338609 | 4.655122 |
| 1.336326 | NA | NA | 6.705712 | 1.364992 | 5.990336 | 1.725044 | -1.6085 | 4.336429 | 4.015701 |
| -2.20126 | NA | NA | 5.130602 | 2.008503 | 2.857823 | 3.32553 | 1.307269 | NA | NA |
| 0.837796 | NA | NA | 6.261377 | 2.405436 | 3.897212 | 4.113742 | 2.49146 | 2.325264 | 2.064652 |
| -0.21511 | NA | NA | 6.643156 | -0.74553 | 2.859891 | 2.838657 | 0.120672 | 2.167109 | NA |
| 3.655574 | 0.766552 | NA | 5.75023 | 2.610738 | 5.520821 | NA | 0.73024 | NA | NA |
| 1.035515 | NA | NA | 6.503316 | 1.39849 | 6.235456 | 2.009388 | 0.206986 | 3.656045 | 2.729722 |
| -1.26942 | NA | NA | 5.327621 | 3.158388 | 2.694132 | 2.762548 | 0.061543 | 2.82464 | 1.710493 |
| -1.36519 | NA | NA | 5.151719 | 0.475768 | 4.10694 | 2.918294 | 1.493841 | 2.601906 | 3.510112 |
| -0.21002 | 0.719469 | 3.781098 | 4.858882 | 0.204901 | 3.772884 | NA | 1.797164 | NA | NA |
| -0.15027 | NA | NA | 5.36174 | 3.544424 | 0.351968 | 1.334062 | 4.325176 | 1.850646 | 3.73536 |
| 0.859036 | NA | NA | 5.830233 | 0.209811 | 3.7665 | 2.475084 | 0.967897 | 2.079958 | 3.690994 |
| NA | NA | NA | 6.216284 | 0.689641 | 3.268569 | 2.936197 | -0.56331 | 1.545527 | 5.023067 |
| 2.235082 | 1.76006 | 4.977363 | 4.900594 | 1.710247 | 5.754847 | 1.957923 | -0.66441 | 3.481672 | 2.579432 |
| 1.968401 | NA | NA | 5.869215 | 0.984528 | 4.891124 | 4.686301 | 0.587281 | 3.338919 | 2.246649 |
| 1.15553 | NA | NA | 4.724464 | 2.668392 | 1.41858 | 1.183188 | 1.692435 | 1.259479 | 1.295966 |
| 2.978862 | NA | NA | 5.907159 | 3.268233 | 3.908926 | 3.525441 | -0.48497 | 3.030144 | 2.629019 |
| 1.016607 | NA | NA | 4.890028 | 0.659282 | 4.094732 | 2.211053 | 1.876722 | NA | 2.113863 |
| -1.3984 | NA | NA | 6.13477 | 5.254831 | 3.916075 | 3.179923 | -0.5317 | 1.758864 | 2.781486 |
| -0.55859 | NA | NA | 4.523331 | 1.062751 | NA | 2.214895 | -0.37632 | 0.766138 | 1.637065 |
| 2.675998 | NA | NA | 5.553381 | 1.735295 | 4.050058 | 3.655111 | 1.481488 | 3.878542 | 0.757705 |
| 0.270872 | NA | NA | 5.566114 | 1.003842 | 5.214038 | 2.535865 | 3.749291 | NA | 2.025495 |
| -0.71409 | NA | NA | 5.165988 | 1.879239 | 2.794432 | 2.566607 | -0.8028 | 3.346557 | 4.100374 |
| -0.80312 | NA | NA | 5.468512 | -0.16681 | 5.060895 | 4.095263 | -0.1253 | 1.632337 | NA |
| 1.398676 | NA | NA | 5.452937 | 2.395207 | 4.356067 | 1.488263 | 4.477394 | 1.126848 | 0.324655 |
| 1.4439 | NA | NA | 6.337303 | 0.835363 | 6.204293 | 4.46801 | 1.122412 | 2.522587 | 0.226827 |
| -0.43224 | NA | NA | 5.94181 | 2.242307 | -0.04494 | 2.21651 | 0.045434 | 3.500746 | 3.300162 |
| 1.212375 | NA | NA | 6.049689 | 0.699771 | 5.550956 | 3.160079 | 3.081926 | 3.553313 | 1.208173 |
| 0.434376 | 1.687883 | 3.801558 | 5.1964 | 0.514671 | 4.344466 | NA | -0.6076 | 1.061314 | 2.457572 |
| 0.280793 | NA | NA | 5.538092 | 2.215089 | 3.401338 | 2.407959 | 0.801386 | 1.138446 | 1.683349 |
| 0.231339 | NA | NA | 6.213909 | 0.622118 | 3.284125 | 0.127621 | 0.483033 | 3.504139 | 1.676126 |
| -1.50429 | NA | NA | 6.185189 | 0.91476 | 5.802253 | 2.15289 | 3.1166 | 1.163125 | 2.881705 |
| 1.325434 | NA | NA | 6.050897 | 0.014127 | 4.396969 | 3.845945 | 1.009152 | 1.355 | 2.308769 |
| 0.611986 | NA | NA | 5.114497 | 3.069361 | 3.861695 | 2.063798 | 2.489333 | 2.23036 | 0.159994 |
| NA | NA | NA | NA | NA | NA | -0.8741 | NA | 2.962346 | 2.586053 |
| 0.17044 | 0.170829 | 0.173391 | 0.181008 | 0.183002 | 0.186421 | 0.187742 | 0.192555 | 0.198653 | 0.200084 |
| -0.0122 | 0.263549 | 3.390348 | 5.849415 | 2.109319 | 4.562309 | 3.067961 | 0.579839 | 2.77208 | 3.028561 |

| | | | | | | | | | |
|---|---|---|---|---|---|---|---|---|---|
| 0.59701 | 1.389137 | 4.186673 | 5.562057 | 1.529255 | 3.892535 | 2.483912 | 1.157909 | 2.264652 | 2.226244 |

| FK866 | QL-VIII-58 | Imatinib | XMD15-27 | XMD14-99 | Cytarabine | GSK10709 | QL-X-138 | PD-032590 | Methotrex |
|---|---|---|---|---|---|---|---|---|---|
| 3.902735 | 1.113515 | NA | NA | NA | 1.207208 | NA | NA | 0.174232 | 2.284752 |
| NA | NA | 2.864849 | 4.562693 | 4.402772 | NA | -2.19886 | -0.28236 | NA | NA |
| -6.74183 | NA | 2.590154 | 4.213962 | 2.303815 | -1.25668 | 0.719763 | NA | -5.70496 | -1.63267 |
| -4.11779 | -1.74632 | 3.49112 | 5.124274 | 5.124147 | 1.475061 | 2.68102 | 1.51637 | 0.80473 | 0.306654 |
| -2.26477 | -0.18452 | 2.655552 | 4.619305 | 5.542231 | 4.003831 | 3.966103 | 0.024132 | 1.924389 | -0.04077 |
| 1.817009 | 0.082566 | NA | 5.210188 | 5.384145 | 1.948403 | 3.765461 | 2.340197 | 0.013377 | 1.434191 |
| -3.1132 | NA | 2.684335 | 4.590386 | 5.133677 | 3.519332 | 3.497832 | 0.14051 | 1.370302 | -0.17385 |
| 3.416614 | -1.49821 | NA | 6.968761 | 6.643063 | 5.239041 | 5.441079 | 0.516792 | 2.408612 | 3.056676 |
| 0.722868 | -2.90103 | 2.806988 | 4.440143 | 4.424356 | -0.53475 | 2.989147 | 1.560961 | -0.53651 | -0.27324 |
| 3.326869 | -1.5966 | NA | 5.906383 | 5.905589 | 0.756129 | 3.807521 | 0.731533 | -1.98837 | 1.945574 |
| -2.00293 | NA | NA | 6.007905 | 5.177829 | 1.770175 | 2.883008 | -0.24484 | 2.083251 | 2.044191 |
| 0.487673 | NA | NA | 5.28892 | 5.092454 | NA | 2.36635 | 1.411114 | NA | NA |
| 1.975284 | -1.86136 | NA | 5.874036 | 5.766867 | 0.631588 | 4.485362 | 4.275551 | -0.53763 | 1.672719 |
| 0.122172 | NA | NA | 5.999238 | 5.958738 | 0.49257 | 2.185492 | 1.64194 | NA | 1.633584 |
| NA | NA | 1.814045 | 5.038193 | 5.074121 | NA | 2.970151 | 3.430962 | NA | NA |
| NA | -2.19507 | NA | 5.497662 | 5.575379 | 2.5027 | 4.351595 | -0.2528 | 1.48238 | -0.20442 |
| 1.61474 | NA | NA | 4.677538 | 4.981215 | 0.919788 | 3.527086 | 1.292157 | -0.9677 | 0.861523 |
| 2.744704 | -0.50775 | NA | 3.739851 | 5.189259 | 0.369613 | 2.334635 | 0.360595 | 1.169274 | 1.201637 |
| NA | NA | 2.97865 | 4.683591 | 4.70035 | NA | 2.945014 | 0.905476 | NA | NA |
| -2.523 | NA | NA | 5.025787 | 4.99735 | 0.443234 | 3.639493 | 4.877221 | -0.30279 | 1.047754 |
| NA | -1.79593 | NA | 4.649195 | 5.086311 | -1.64006 | 3.397651 | 1.109333 | -1.76983 | -2.75464 |
| 1.193695 | -0.50244 | NA | NA | NA | 1.157034 | 2.247011 | NA | 1.893057 | 1.96763 |
| 0.779299 | -1.93882 | 3.597029 | 5.332276 | 5.329311 | 2.850993 | 3.088846 | -0.87651 | 0.698121 | 1.047751 |
| 1.742538 | -0.93705 | NA | 5.456034 | 5.602609 | 3.963279 | 1.120644 | 1.303384 | -0.47801 | -0.11872 |
| -6.39981 | NA | NA | 4.449373 | 4.057369 | -1.266 | 2.462361 | 2.33895 | 0.110679 | -0.4262 |
| 1.447775 | -2.99589 | NA | 4.50865 | 3.781951 | 2.514348 | 2.713509 | -0.41866 | -1.08199 | 1.41465 |
| -3.85864 | NA | NA | 4.101478 | 3.93738 | NA | 3.051867 | 1.457783 | -1.05913 | NA |
| -2.40382 | -1.88247 | NA | 4.79076 | 4.148068 | 2.307443 | 1.060117 | 1.059062 | -0.41308 | 0.464364 |
| NA | -2.97429 | NA | 4.248815 | 3.823861 | -0.02301 | -3.81546 | -0.02068 | -2.61747 | 0.768507 |
| 1.277525 | NA | NA | 5.285857 | 5.511696 | 1.096563 | 2.939328 | 2.80606 | -1.29595 | 0.043792 |
| -1.30431 | NA | NA | 5.111679 | 5.114657 | NA | 2.747315 | 2.174989 | 1.059197 | NA |
| -1.82686 | NA | NA | 3.419607 | 3.312157 | 1.494372 | 3.153652 | -0.81352 | 1.324433 | 0.184527 |
| -1.2801 | NA | NA | 4.911226 | 4.662537 | 1.54916 | 2.302746 | 1.51135 | NA | -0.87673 |
| 2.51683 | -1.94874 | NA | 5.218975 | 5.218975 | -0.72473 | 3.824409 | 3.666889 | -4.17225 | 1.172321 |
| -3.79856 | NA | NA | 5.821456 | 5.787208 | 0.680976 | 2.347221 | 1.51321 | -1.56392 | 0.008914 |
| 0.349167 | -2.96355 | NA | 5.238793 | 4.952131 | 2.961319 | 3.156224 | 5.240533 | -0.24643 | -0.75091 |
| 0.002246 | NA | NA | 5.63351 | 5.62802 | 0.24575 | 4.125788 | 3.719665 | -2.4866 | 0.659132 |
| -0.88341 | NA | 2.827324 | 4.574921 | 4.259398 | 1.813334 | -3.67316 | 1.927467 | -0.22256 | -0.55783 |
| -3.81938 | -2.11897 | NA | 5.090405 | 5.116216 | -0.50147 | 2.687508 | 2.202634 | -1.3466 | 1.075981 |
| -4.41431 | NA | NA | 4.412676 | 5.222021 | 0.352009 | 1.764457 | 0.192957 | 0.751203 | 1.505908 |
| -0.08192 | -0.63006 | NA | 4.782848 | 5.412217 | 0.748612 | 2.953668 | 3.432527 | -0.57896 | 1.549797 |
| 1.477101 | -0.27164 | NA | 5.146472 | 5.388273 | 0.23781 | 2.54614 | 0.504071 | -1.31261 | 0.244426 |
| -2.50216 | NA | NA | 5.126458 | 4.77851 | -0.87887 | 2.914967 | 1.830378 | -2.05855 | 1.089929 |
| -1.33732 | NA | NA | NA | NA | 0.486525 | NA | NA | 1.134145 | 1.814491 |
| 0.201115 | 0.212269 | 0.212566 | 0.214351 | 0.217104 | 0.219763 | 0.224041 | 0.224969 | 0.227662 | 0.231789 |
| 0.12601 | -1.12948 | 2.701006 | 5.16232 | 5.157627 | 1.536267 | 2.927809 | 1.153926 | 0.121099 | 0.941103 |

| | | | | | | | | | |
|---|---|---|---|---|---|---|---|---|---|
| -1.11511 | -1.74665 | 3.134334 | 4.875868 | 4.826191 | 0.863854 | 2.228053 | 1.735191 | -0.66816 | 0.459776 |

| NSC-87877 | Temozolon | PD-033299 | Mitomycin | Bicalutami | FR-180204 | WZ3105 | HG-5-88-0: | AZD8055 | Bleomycin |
|---|---|---|---|---|---|---|---|---|---|
| NA | 7.309104 | 5.480756 | NA | NA | NA | NA | 5.323976 | 2.03403 | NA |
| 5.344281 | NA | NA | -0.99998 | NA | 4.753504 | -2.30257 | NA | NA | 0.769875 |
| 3.889081 | 4.739896 | 3.285065 | -2.96603 | 3.617614 | 3.676106 | -0.96485 | NA | -0.24915 | 0.3717 |
| 3.914125 | 4.931195 | 4.078938 | -0.2558 | 4.984005 | 5.793705 | -3.61515 | 3.447998 | -1.40454 | 4.233593 |
| 5.459469 | 6.5492 | 4.608763 | -0.57493 | 5.319904 | 5.529537 | -2.3017 | 4.463998 | 0.636028 | 3.868778 |
| 5.765304 | 6.341052 | 2.125334 | 1.154755 | 5.241743 | 5.762941 | -0.73901 | 5.232121 | 0.54702 | 4.565741 |
| 5.586051 | 5.993915 | 4.188253 | -0.14411 | 4.487337 | 5.775328 | -2.06106 | NA | -2.06576 | -0.5925 |
| 7.488978 | 7.878096 | 4.399261 | 2.654632 | 6.779484 | 7.065682 | 1.036819 | 6.742257 | 0.818342 | 7.399005 |
| 4.436442 | 5.453596 | 1.881867 | 0.402503 | 4.357172 | 4.743262 | -0.35185 | 4.357172 | -1.45924 | 5.15896 |
| 6.490259 | 6.930585 | 4.484216 | 1.586521 | 5.026686 | 5.7153 | -0.16091 | 5.826963 | 0.056672 | 5.207652 |
| 6.094744 | 6.627371 | 0.187503 | 2.050098 | 4.061591 | 5.765696 | -0.51578 | NA | -1.25367 | 4.945506 |
| 5.526472 | 4.857846 | NA | -0.66056 | 4.417839 | 6.249193 | -0.41802 | NA | NA | 2.339499 |
| 5.899666 | 6.614242 | 3.289634 | -0.36232 | 4.054 | 6.507319 | 1.817177 | 2.788709 | 0.257422 | 3.208609 |
| 6.146734 | 6.000286 | NA | 0.801457 | 4.810572 | 6.321762 | -0.59077 | NA | NA | -1.92782 |
| 5.370852 | NA | NA | 2.152897 | NA | 4.744038 | 2.980429 | NA | NA | 4.708179 |
| 4.541765 | 5.73113 | NA | -0.62175 | 4.059476 | 6.084106 | -0.6739 | 5.633429 | -0.16691 | 3.501145 |
| 4.092951 | 5.206669 | 1.849501 | -0.23608 | 4.656098 | 4.653694 | 0.24038 | NA | -0.77246 | -1.52022 |
| 3.942307 | 6.145901 | 3.26809 | -0.49267 | 5.047289 | 5.218955 | -0.95571 | 5.021629 | 1.34343 | -1.47818 |
| 4.363902 | NA | NA | -2.82526 | NA | 5.15138 | -0.6512 | NA | NA | 1.550322 |
| 4.895969 | 5.257403 | 0.398061 | -3.76768 | 4.56848 | 5.695218 | 2.528574 | NA | -0.72112 | 0.255555 |
| 3.522988 | NA | 2.24936 | -0.63979 | NA | 5.018204 | -1.09589 | 3.261301 | 0.273267 | 0.94687 |
| NA | 6.58615 | 4.94928 | 0.938724 | 4.969892 | 4.800626 | NA | 5.615143 | 2.293678 | 3.352587 |
| 4.772583 | 6.253399 | 3.940825 | 0.874882 | 4.714557 | 4.627764 | -0.83652 | 3.872247 | 0.463185 | 6.664339 |
| 5.636889 | 6.375606 | 4.278308 | -1.05304 | 5.418992 | 6.166347 | 0.519209 | 5.419375 | 0.084981 | 0.819087 |
| 1.778179 | 5.315239 | 0.33508 | -3.11384 | 3.642996 | 5.118979 | 1.543953 | NA | -1.57815 | -2.87135 |
| 4.715836 | 6.227005 | 2.768309 | 0.514634 | 3.932742 | 6.033484 | -1.0882 | 4.302881 | 1.381865 | 4.030649 |
| 2.614183 | 5.012518 | 1.437114 | -1.99549 | 4.334062 | 4.313825 | -1.70275 | NA | -0.1729 | -1.48852 |
| 5.639004 | 5.730452 | 4.155152 | -0.37965 | 4.057129 | 5.373552 | -1.62027 | 3.836154 | -1.04787 | -0.14984 |
| 5.173178 | 5.570383 | 3.672414 | -1.06322 | 4.623169 | 5.252213 | -1.56548 | 4.141987 | 1.034457 | 3.684741 |
| 5.950293 | 6.431368 | 4.42737 | 0.544436 | 4.631802 | 4.763617 | 1.855322 | NA | 1.014922 | 3.834665 |
| 5.465653 | 6.083182 | 2.921748 | 0.812804 | 4.692372 | 5.65828 | -0.61186 | NA | -0.38598 | 4.769807 |
| 5.604159 | 6.136512 | 4.010754 | -0.06234 | 4.464695 | 5.633746 | -1.24515 | NA | 0.99153 | 2.468775 |
| 5.349304 | 5.174188 | NA | -0.15427 | 3.989716 | 3.028609 | -2.58087 | NA | NA | 4.05367 |
| 5.5566 | 5.925798 | -0.0375 | 1.882155 | 4.221995 | 5.879915 | 3.058632 | 4.128592 | -0.45494 | 5.497327 |
| 4.258518 | 6.695671 | 0.563056 | 0.222682 | 4.699309 | 5.277314 | -1.34675 | NA | -0.1367 | 2.749578 |
| 5.700967 | 5.900039 | 2.454971 | -2.61249 | 4.637401 | 4.87216 | 2.869248 | 5.11571 | 0.312584 | -1.78645 |
| 6.079798 | 5.66016 | 3.470862 | 0.648636 | 4.385867 | 6.026169 | 2.465856 | NA | 0.25499 | 3.301251 |
| 5.009328 | 4.605749 | 3.150757 | 1.073797 | 4.032192 | 4.931318 | -2.47412 | NA | 0.880263 | 3.258369 |
| 4.882755 | 5.910104 | 2.61392 | -0.02317 | 4.135277 | 5.52331 | 2.488858 | 3.790833 | -0.67203 | 0.540316 |
| 3.659879 | 5.599893 | 1.732396 | 1.442508 | 4.116247 | 5.75948 | 0.265733 | NA | 0.052923 | 1.607026 |
| 5.931194 | 6.109092 | 2.165124 | -0.03205 | 4.974732 | 6.072383 | 2.783855 | 5.162744 | 0.704375 | -1.23755 |
| 5.406174 | 6.288177 | 3.600866 | -0.60003 | 5.227128 | 5.554408 | -1.33602 | 4.059077 | 0.110848 | -2.74336 |
| 5.420635 | 4.542579 | 0.736405 | 1.053544 | 4.777723 | 5.156306 | -0.78735 | NA | 0.666401 | -0.7747 |
| NA | 5.062426 | 4.808468 | NA | 4.156728 | NA | NA | NA | 1.471571 | NA |
| 0.232865 | 0.235166 | 0.237282 | 0.243345 | 0.250988 | 0.255111 | 0.255654 | 0.266665 | 0.274287 | 0.283994 |
| 5.293499 | 6.08188 | 3.317476 | 0.205213 | 4.728054 | 5.550596 | -0.56332 | 4.883825 | -0.11991 | 2.632913 |

| | | | | | | | | | |
|---|---|---|---|---|---|---|---|---|---|
| 4.891165 | 5.768879 | 2.700129 | -0.33254 | 4.475217 | 5.267544 | 0.059867 | 4.39217 | 0.284256 | 1.693327 |

| Sunitinib | Vinblastine | Erlotinib | UNC0638 | Z-LLNle-CH | Tubastatin | Zibotentan | Y-39983 | SGC0946 | SB-715992 |
|---|---|---|---|---|---|---|---|---|---|
| NA | -2.24725 | NA | NA | NA | NA | NA | NA | 3.21476 | NA |
| 2.283829 | NA | 3.130688 | 2.298006 | -0.70534 | 3.185976 | 5.73239 | 2.831123 | NA | -3.61751 |
| 1.119297 | -2.73519 | 2.574158 | 2.302143 | -0.16381 | 3.026766 | 4.889398 | 3.108626 | 1.157914 | -3.17974 |
| 3.802692 | -3.43696 | 1.420509 | 3.34356 | 1.126975 | 4.374183 | 5.783027 | 5.787427 | 0.828277 | -1.29728 |
| 5.111926 | -1.3089 | 3.35383 | 2.225149 | 1.227148 | 4.122428 | 5.05111 | 5.418727 | 2.363556 | -2.37114 |
| NA | -2.5992 | NA | 3.025252 | NA | 5.141228 | 5.916319 | 5.655247 | 2.223315 | 0.936572 |
| NA | -1.63691 | NA | 2.25796 | NA | 4.296697 | 5.377114 | 5.315948 | 1.85901 | -1.06496 |
| NA | -0.64984 | NA | 3.692354 | NA | 7.204119 | 7.231458 | 6.087685 | 3.160459 | 2.639187 |
| 1.867899 | -4.01776 | 2.653534 | 1.877952 | -0.04264 | 3.175661 | 5.080703 | 3.134453 | 1.286156 | -2.67998 |
| NA | -3.35705 | NA | 3.708755 | NA | 5.220067 | 6.52885 | 6.507722 | 2.836241 | 0.963123 |
| NA | -3.81644 | NA | 2.478636 | NA | 3.81173 | 5.917295 | 6.708914 | 2.789897 | -1.65032 |
| NA | NA | NA | 4.577693 | NA | 4.972111 | 6.271006 | 5.467681 | 2.475161 | -1.84917 |
| NA | -3.7861 | NA | 5.091983 | NA | 6.438162 | 6.519687 | 6.33924 | 0.429154 | -1.23971 |
| NA | -5.83265 | NA | 3.299791 | NA | 3.389842 | 5.480971 | 4.453171 | 2.369477 | 0.03412 |
| NA | NA | 3.447573 | 3.697543 | NA | 2.904939 | 5.650784 | 5.749968 | NA | 0.475297 |
| NA | -3.03284 | NA | 3.389973 | NA | 6.016615 | 6.474698 | 5.801502 | 2.637697 | -1.56667 |
| NA | -6.3479 | NA | 4.617721 | NA | 4.185033 | 5.504046 | 5.300528 | 1.872893 | 0.198269 |
| NA | -4.03658 | NA | 2.860761 | NA | 3.580825 | 5.801099 | 4.535719 | 2.024811 | -1.5545 |
| 4.460106 | NA | NA | 2.920659 | 1.963196 | 3.688605 | 5.374736 | 3.726737 | NA | -0.46846 |
| NA | -3.53983 | NA | 6.017812 | NA | 5.693432 | 5.676827 | 5.695218 | 1.155008 | 0.48735 |
| NA | -5.84559 | NA | 3.267563 | NA | 4.785906 | 5.445772 | 3.555993 | NA | -1.66504 |
| NA | -0.65585 | NA | 1.749356 | NA | 4.095757 | 5.541955 | 6.290388 | 0.960738 | -0.76369 |
| 4.73835 | -2.87962 | 3.08147 | 1.960071 | 3.438236 | 4.081679 | 5.580102 | 5.004309 | 2.180548 | -0.86226 |
| NA | -3.62577 | NA | 6.144179 | NA | 5.919322 | 6.19679 | 4.854689 | 2.091429 | -3.61195 |
| NA | -4.6067 | NA | 5.41532 | NA | 5.118957 | 5.118979 | 5.118874 | 1.362736 | -0.17221 |
| NA | -3.00795 | NA | 3.573245 | NA | 5.409495 | 6.029703 | 5.987597 | 2.219717 | -5.19495 |
| NA | NA | NA | 2.653678 | NA | 2.527322 | 5.173898 | 4.118968 | 1.560589 | -3.3637 |
| NA | -2.87799 | NA | 3.232354 | NA | 5.284812 | 5.960145 | 5.380716 | 2.435718 | -0.88035 |
| NA | -4.13384 | NA | 2.325824 | NA | 4.954097 | 5.245469 | 4.601623 | 1.173017 | -4.8204 |
| NA | -2.83988 | NA | 2.594325 | NA | 4.048444 | 6.15373 | 5.786651 | 2.112926 | 0.001049 |
| NA | NA | NA | 4.202317 | NA | 5.467457 | 5.766371 | 4.926852 | 1.987551 | 0.212384 |
| NA | -3.45013 | NA | 2.407146 | NA | 2.58672 | 5.935437 | 5.935803 | 2.113935 | -2.95772 |
| NA | -3.71949 | NA | 1.937256 | NA | 3.874243 | 4.432563 | 2.983704 | 1.79213 | -2.29187 |
| NA | -4.82535 | NA | 6.485417 | NA | 5.866081 | 5.885901 | 5.125727 | 2.026948 | 0.467055 |
| NA | -3.88922 | NA | 5.425617 | NA | 5.660966 | 6.035158 | 4.32037 | 1.991393 | -2.03659 |
| NA | -3.41006 | NA | 3.462903 | NA | 4.120198 | 5.160848 | 5.590473 | 2.108744 | -0.55469 |
| NA | -3.68028 | NA | 3.786022 | NA | 5.483973 | 6.143297 | 4.633531 | 2.440176 | -0.02726 |
| 2.695354 | -5.10695 | 3.164922 | 3.867713 | -0.17869 | 4.758318 | 5.393478 | 2.940715 | 1.688181 | -4.5666 |
| NA | -5.57278 | NA | 4.443325 | NA | 5.68027 | 5.722527 | 5.337279 | 1.907449 | -2.30655 |
| NA | -4.66487 | NA | 3.519089 | NA | 4.567164 | 5.826116 | 3.144162 | 2.342216 | 0.192181 |
| NA | -4.40145 | NA | 2.069495 | NA | 5.177177 | 6.008373 | 4.281784 | 2.235395 | -0.1037 |
| NA | -2.61301 | NA | 2.931718 | NA | 5.574008 | 5.886541 | 5.881717 | 1.501017 | -1.1969 |
| NA | -3.64044 | NA | 3.598916 | NA | 5.257822 | 5.767868 | 5.516398 | 0.829684 | -1.58139 |
| NA | -3.26229 | NA | NA | NA | NA | NA | NA | 2.526132 | NA |
| 0.287637 | 0.293046 | 0.294356 | 0.29617 | 0.298925 | 0.309031 | 0.309726 | 0.312274 | 0.319643 | 0.319771 |
| 2.837129 | -3.25611 | 2.763382 | 3.220308 | 0.288467 | 4.414493 | 5.83588 | 5.188452 | 2.095549 | -0.98967 |

| | | | | | | | | | |
|---|---|---|---|---|---|---|---|---|---|
| 3.964603 | -3.74997 | 3.123196 | 3.599653 | 1.740914 | 4.787289 | 5.658503 | 4.829611 | 1.864307 | -1.52265 |

| Temsirolim | FMK | PIK-93 | CGP-60474 | TL-2-105 | GW 44175 | AZD6244 | QS11 | RO-3306 | NPK76-II-7 |
|---|---|---|---|---|---|---|---|---|---|
| -0.66333 | NA | NA | NA | NA | 1.257325 | 5.292161 | NA | 4.66422 | NA |
| NA | 5.756862 | -0.44521 | -3.12006 | 3.308421 | NA | NA | 3.243556 | NA | -0.70176 |
| -3.03609 | 4.077583 | 1.962386 | -2.45457 | 2.47762 | 2.585756 | -2.90618 | 0.514752 | 2.937764 | 2.327797 |
| -1.95486 | 5.766289 | 4.343402 | -2.05996 | 5.120244 | 0.724817 | 3.8571 | 2.138315 | 3.77723 | 5.124161 |
| 0.33945 | 5.579353 | 2.188729 | -1.65413 | 4.467157 | 4.003831 | 4.487842 | 2.932561 | 4.760393 | 5.4137 |
| -0.63993 | 5.859264 | 4.783856 | NA | 3.310774 | 3.368876 | 1.862607 | 5.181658 | 2.642548 | 4.758069 |
| -3.5052 | 5.143411 | 3.628162 | NA | 3.264108 | 3.519332 | 3.308619 | 5.045192 | 4.403747 | 4.3768 |
| -2.01249 | 7.649954 | 3.658328 | NA | 3.807214 | 5.282415 | 5.081798 | 4.506229 | 5.06763 | 6.449478 |
| -2.40115 | 5.109573 | 1.188466 | -2.1181 | 3.128565 | 2.643594 | 3.239885 | 3.089099 | 1.137279 | 1.252562 |
| -3.90115 | 5.764671 | 3.636544 | NA | 2.831046 | 1.801538 | 3.793812 | 4.717073 | 3.056537 | 5.310451 |
| -2.71689 | 5.392066 | -0.24522 | NA | 3.104154 | 1.770171 | 1.043118 | 5.613551 | 4.095113 | 4.493389 |
| NA | 5.99649 | 2.951134 | NA | 4.50284 | NA | 4.13688 | 1.779604 | NA | 4.437333 |
| -3.63292 | 6.543467 | 3.831286 | NA | 5.874036 | 1.735883 | 2.922617 | 5.653058 | 4.770312 | 5.283456 |
| -3.14757 | 6.635363 | 1.593651 | NA | 2.307464 | 2.181151 | 4.132338 | 5.794959 | NA | 3.430681 |
| NA | 5.642663 | 4.296697 | NA | 3.953506 | NA | NA | 5.080331 | NA | 4.461738 |
| 0.182921 | 5.98733 | 2.528927 | NA | 4.08115 | 2.555427 | 2.914169 | 4.964009 | 4.821509 | 3.989146 |
| -2.91094 | 5.412531 | 0.692643 | NA | 4.751168 | 2.584899 | 2.931737 | 4.673956 | 3.681359 | 3.407114 |
| -1.37664 | 4.892588 | 2.677492 | NA | 4.252357 | 3.400707 | 4.11747 | 5.099689 | 2.808236 | 1.143529 |
| NA | 5.376397 | 1.598525 | -2.19021 | 4.236555 | NA | NA | 3.095494 | NA | 1.60123 |
| -1.17279 | 5.695218 | 5.694704 | NA | 5.023082 | 1.606256 | 3.773306 | 1.257217 | 1.986008 | 5.011624 |
| -2.71503 | 5.434313 | 1.998905 | NA | 3.445965 | -0.56293 | NA | 2.123459 | 0.815109 | 2.987576 |
| 1.451619 | 5.397585 | 1.43406 | NA | 5.143514 | 1.501163 | 4.377894 | 4.877476 | 5.120292 | 3.252492 |
| -1.26157 | 5.99609 | 1.787953 | 1.000945 | 3.445317 | 2.608698 | 3.977048 | 4.369619 | 4.596781 | 4.202281 |
| -1.87176 | 5.560272 | 2.897042 | NA | 4.124367 | 3.703928 | 3.060544 | 5.491145 | 4.036209 | 4.711261 |
| -1.66823 | 4.893138 | 4.980043 | NA | 3.54665 | 2.816394 | 2.006685 | 0.911798 | 3.575257 | 4.449548 |
| -0.54965 | 4.424046 | 3.418082 | NA | 4.307147 | 2.722355 | 2.533938 | 2.60548 | 3.288238 | 5.358144 |
| NA | 5.325563 | 2.980525 | NA | 4.368901 | NA | 2.322328 | 2.626224 | 3.040724 | 0.522357 |
| 0.23388 | 5.58616 | 2.892201 | NA | 0.514826 | 3.804899 | 4.05995 | 3.066576 | 4.336577 | 3.366994 |
| -2.0215 | 5.093564 | 2.655555 | NA | 4.398449 | 1.331439 | 3.714446 | 4.149549 | 0.777502 | 2.14381 |
| 0.912745 | 5.786431 | 3.544806 | NA | 4.852904 | 3.271243 | 2.031187 | 4.946647 | 4.762758 | 4.299945 |
| NA | 5.734408 | 5.152336 | NA | 4.466841 | NA | 4.016343 | 4.308451 | 3.711874 | 3.223043 |
| -0.70349 | 5.341831 | 0.925279 | NA | 1.805872 | 2.489243 | 4.193024 | 2.865895 | 3.984089 | 1.763763 |
| NA | 5.516107 | 1.886694 | NA | 4.30947 | 3.12342 | 1.081176 | 4.886839 | NA | 2.580521 |
| -3.56406 | 5.884673 | 5.240062 | NA | 5.206092 | 2.081153 | 0.643169 | 5.085898 | 0.939966 | 5.193516 |
| -1.16763 | 5.268422 | 1.926467 | NA | 5.53656 | 3.666813 | 0.864448 | 4.243863 | 4.99019 | 1.200759 |
| -2.8684 | 5.769544 | 2.022096 | NA | 3.484741 | 3.076103 | 4.088137 | 1.412373 | 3.299456 | 3.030009 |
| -2.31174 | 6.278259 | 2.860575 | NA | 5.421339 | 1.560216 | 1.809146 | 5.372376 | 4.007945 | 3.634611 |
| -1.01961 | 5.022202 | 1.54371 | -1.77286 | 4.004764 | -2.38715 | 2.947685 | 3.690263 | 3.773231 | 1.243481 |
| -5.24035 | 5.710515 | 3.250471 | NA | 4.605581 | 3.062644 | 1.281802 | 2.945795 | 1.714658 | 4.88565 |
| -0.80127 | 5.388855 | 1.878782 | NA | 4.749104 | 3.737796 | 2.606744 | 5.519887 | 3.63867 | 1.823164 |
| -1.27784 | 5.810942 | 4.599461 | NA | 5.282279 | 1.815998 | 2.204233 | 3.522923 | 2.286979 | 4.074562 |
| -2.85984 | 5.851136 | 2.720755 | NA | 2.523865 | 1.5763 | 2.652523 | 5.385756 | 3.613729 | 3.370652 |
| -2.0546 | 5.63437 | 5.786472 | NA | 4.079344 | 3.219671 | 1.086891 | 3.187685 | 3.903866 | 5.126458 |
| -2.91974 | NA | NA | NA | NA | 1.119284 | 1.558716 | NA | 4.899767 | NA |
| 0.321666 | 0.322501 | 0.323288 | 0.324364 | 0.338304 | 0.341142 | 0.351239 | 0.35135 | 0.352391 | 0.355275 |
| -2.09179 | 5.718203 | 2.545369 | -2.28136 | 3.796578 | 2.627715 | 3.138499 | 4.11927 | 3.758848 | 3.821038 |

| | | | | | | | | | |
|---|---|---|---|---|---|---|---|---|---|
| -1.6114 | 5.511202 | 3.027022 | -0.98737 | 4.115341 | 2.214997 | 2.620473 | 3.677947 | 3.379161 | 3.322298 |

| Bexarotene | VNLG/124 | BMS-5369 | MP470 | UNC1215 | AC220 | Etoposide | SB52334 | AKT inhibit | NVP-BHG7 |
|---|---|---|---|---|---|---|---|---|---|
| NA | NA | NA | NA | 3.907907 | NA | NA | NA | NA | NA |
| 4.488646 | 4.196445 | 3.667138 | 3.628999 | NA | 2.050816 | -0.65514 | 5.192018 | 2.608522 | 2.029026 |
| 3.454544 | 3.531925 | 1.330491 | 3.700749 | 1.612709 | 1.808366 | -2.02607 | 3.746326 | 1.524838 | -0.58246 |
| 3.747 | 4.277858 | 2.868882 | 2.772455 | 1.826634 | -1.52938 | 2.533787 | 5.792229 | 1.318296 | 5.121094 |
| 5.390125 | 4.030197 | 4.005398 | -0.03798 | 1.706959 | 2.388327 | 4.882789 | 4.646549 | 1.851721 | 3.712697 |
| 2.671244 | 4.558418 | NA | 4.178006 | 2.856245 | 1.307684 | 4.678824 | 5.328949 | 2.745517 | 5.382695 |
| 3.594317 | 2.449036 | NA | 1.009318 | 2.702956 | 2.126099 | 3.64147 | 3.845517 | 3.763724 | 4.13208 |
| 6.159034 | 6.310809 | NA | 4.263585 | 4.476899 | 3.507311 | 5.782212 | 2.448345 | 3.999166 | 6.884215 |
| 4.157681 | 3.051371 | 2.488318 | 3.30345 | 2.054587 | 1.719129 | 3.913562 | 2.23811 | 2.466775 | 2.687177 |
| NA | 5.006886 | NA | 3.253413 | 3.529388 | 1.380642 | 6.205802 | 3.25459 | 1.195471 | 4.374156 |
| 4.954932 | 5.322316 | NA | 4.654486 | 3.266929 | 2.070497 | 1.938669 | 3.996267 | -0.2673 | 3.541403 |
| 1.434364 | 3.565927 | NA | 3.63341 | 3.159809 | 3.070224 | 1.57481 | 4.325648 | 2.364194 | 4.509324 |
| 4.311021 | 4.70592 | NA | 1.009223 | 3.27546 | 3.571451 | 4.580718 | 6.543057 | 0.506107 | 5.553706 |
| 2.650719 | 5.010128 | NA | 6.643099 | 3.443147 | 2.28761 | -0.01662 | 2.715647 | -0.76741 | 2.839884 |
| 4.833812 | 4.387865 | NA | 4.986151 | NA | 2.776343 | 3.431983 | 5.245391 | 3.678641 | 3.284368 |
| 2.897023 | 5.130921 | NA | 1.519412 | 3.330844 | 2.297813 | 5.259002 | 4.489401 | 1.332154 | 3.162889 |
| 3.080604 | 4.274017 | NA | 3.439654 | 2.559909 | 2.68121 | 2.679606 | 3.92865 | -0.60922 | 3.20922 |
| 4.523244 | 4.342779 | NA | 4.937793 | 2.744704 | 2.332469 | 0.930839 | 3.286912 | 0.73792 | 3.831386 |
| 4.443156 | 3.143874 | 3.720631 | 2.833735 | NA | 1.961378 | -0.36878 | 3.807174 | 1.550963 | 4.298997 |
| 2.63818 | 4.089716 | NA | 0.168494 | 2.543474 | 2.713604 | -0.03416 | 5.695218 | 2.162095 | 4.584757 |
| 2.722074 | 2.635926 | NA | 3.936964 | NA | 1.386408 | 1.054299 | 3.338529 | 0.062734 | 4.046381 |
| 3.9323 | 5.004187 | NA | NA | 3.101669 | NA | NA | 2.206094 | NA | 4.133831 |
| 5.085416 | 4.639129 | 4.254542 | 2.483447 | 2.873695 | 2.277451 | 4.347218 | 2.535702 | 2.245198 | 3.98226 |
| 4.79261 | 4.482732 | NA | 3.887654 | 3.11679 | 2.516425 | 1.730789 | 5.79629 | 1.313557 | 4.43838 |
| 2.768215 | 3.680474 | NA | 3.501542 | 1.380499 | 1.760656 | -0.30799 | 4.917831 | 0.986515 | 4.449146 |
| 4.996604 | 4.39519 | NA | 4.712744 | 2.902313 | 2.20536 | 2.161363 | 5.910914 | 1.926128 | 5.382223 |
| 3.311632 | 3.42588 | NA | 4.069147 | 2.230432 | 2.352552 | 1.659763 | 4.704466 | NA | 3.074353 |
| 4.519504 | 4.688566 | NA | 4.276808 | 3.128772 | 0.582989 | 1.215706 | 5.160104 | 1.014988 | 4.164289 |
| 3.523214 | 3.539835 | NA | 0.263571 | 2.232916 | 1.932594 | 0.520661 | 1.619906 | 1.012194 | 4.175917 |
| 3.936857 | 2.986982 | NA | 5.52113 | 3.036622 | 3.203947 | 2.109685 | 6.181127 | 2.763727 | 5.511166 |
| 2.340258 | 4.420544 | NA | 5.503802 | 2.439353 | 2.293161 | 2.524599 | 5.177773 | 2.503158 | 5.019205 |
| 2.965251 | 4.573815 | NA | 0.396503 | 2.807082 | 1.925989 | 5.199934 | 3.265284 | 1.553293 | 5.263452 |
| 3.812877 | 4.218079 | NA | 4.608368 | 2.27002 | 2.301204 | 4.358723 | 3.287338 | -0.26116 | 3.130156 |
| 3.61119 | 4.525828 | NA | 5.608435 | 2.707827 | 2.91639 | 5.665262 | 5.400838 | 2.92569 | 5.218487 |
| 1.541099 | 5.03725 | NA | 5.568585 | 3.293541 | 3.407465 | 4.388657 | 4.845832 | 2.503783 | 4.285696 |
| 2.591432 | 3.839372 | NA | 4.437244 | 2.813124 | 2.597891 | -0.136 | 4.797695 | 2.865243 | 4.115467 |
| 4.409371 | 4.760564 | NA | 4.668336 | 2.946333 | 3.323181 | 2.403427 | 3.777672 | 2.622249 | 4.288799 |
| 3.494196 | 3.98304 | 2.623426 | 3.507625 | 2.058853 | 1.946909 | 3.807016 | 3.825799 | 3.41178 | 2.480735 |
| 3.261584 | 4.064054 | NA | 3.611549 | 2.50867 | 2.798162 | 1.633112 | 5.518901 | 2.15094 | 3.129448 |
| 3.921264 | 4.140534 | NA | 4.726416 | 2.636325 | 2.801162 | 2.189807 | 4.617315 | 3.364304 | 2.556955 |
| 3.556244 | 4.709289 | NA | 5.555085 | 3.050737 | 3.101858 | 4.902427 | 5.815806 | 3.393058 | 4.556116 |
| 3.926388 | 4.689007 | NA | 4.692358 | 2.924543 | 2.169093 | 2.346022 | 4.990642 | 1.634026 | 4.106121 |
| 3.241756 | 3.455639 | NA | 3.288837 | 2.67875 | 2.790505 | 1.795578 | 5.750365 | 3.238039 | 2.83581 |
| NA | NA | NA | NA | 3.148795 | NA | NA | NA | NA | NA |
| 0.359192 | 0.361341 | 0.368896 | 0.369113 | 0.371821 | 0.372926 | 0.381455 | 0.382058 | 0.385268 | 0.386263 |
| 3.896769 | 4.36193 | 2.872045 | 3.346778 | 2.903443 | 2.108624 | 2.902132 | 4.177859 | 1.673478 | 3.745462 |

| | | | | | | | | | |
|---|---|---|---|---|---|---|---|---|---|
| 3.573707 | 4.12518 | 3.532866 | 3.826182 | 2.701297 | 2.386097 | 2.29863 | 4.517785 | 2.040978 | 4.129126 |

| CCT00709: | TGX221 | Salubrinal | Elesclomol | KIN001-05! | FTI-277 | Shikonin | Nutlin-3a | KIN001-26( | ABT-263 |
|---|---|---|---|---|---|---|---|---|---|
| 5.261029 | NA | NA | -4.93658 | NA | NA | NA | 5.66956 | NA | 3.736741 |
| NA | 1.572551 | 2.660693 | NA | 3.962879 | 2.186058 | -0.97112 | NA | 4.457526 | NA |
| 3.372004 | 3.861403 | 2.387091 | -2.21988 | 3.135032 | 0.658686 | -1.46186 | 1.739453 | 3.709197 | -0.3582 |
| 4.622066 | 3.111687 | 3.668324 | -3.8189 | 5.053755 | 2.26065 | -0.99509 | 4.681817 | 5.744255 | -0.62996 |
| 4.808456 | 5.03812 | 2.652784 | -1.92126 | 4.245144 | 2.361598 | -0.74619 | 4.062102 | 5.60562 | 3.711546 |
| 4.375996 | NA | NA | -2.4285 | 4.881402 | 2.70901 | 1.914123 | 4.9249 | 5.082073 | 1.748952 |
| 3.864668 | 4.394879 | NA | -0.83352 | 4.76167 | 2.848087 | 0.818452 | 3.630693 | 4.216587 | -2.9828 |
| 5.100963 | NA | NA | -3.79966 | 5.498508 | 4.69415 | 2.220945 | 5.179769 | 7.698295 | 2.812405 |
| 4.000911 | 1.753487 | 2.542246 | -3.29635 | 3.862979 | 2.119668 | -0.88004 | 4.189357 | 0.226695 | 1.501455 |
| 4.541219 | NA | NA | -3.8748 | 4.76211 | 3.535943 | 0.601555 | 5.029112 | 5.153451 | 0.876359 |
| 2.985812 | NA | NA | -4.09404 | 5.777519 | 3.003991 | -0.86056 | 4.103107 | 5.523094 | 3.998207 |
| 3.376344 | NA | NA | NA | 5.280409 | 1.985355 | 0.440224 | NA | 5.404479 | NA |
| 4.895409 | NA | NA | -1.87579 | 5.65082 | 3.02923 | 0.325221 | 5.591296 | 5.725365 | 0.66757 |
| 5.401506 | NA | NA | -2.76236 | 5.892941 | 1.38952 | -1.18172 | NA | 4.378436 | NA |
| NA | 5.056769 | 5.114508 | NA | 3.109213 | 2.668952 | 1.454986 | NA | 5.746471 | NA |
| 3.451142 | NA | NA | -1.0616 | 3.342492 | 2.398882 | 4.302222 | 4.023235 | 5.451715 | 2.816226 |
| 3.107172 | NA | NA | -4.18343 | 4.367275 | 1.13229 | 0.163481 | 3.976236 | 5.103702 | 2.831912 |
| 4.538729 | NA | NA | -1.40217 | 2.95872 | 1.703352 | 0.002422 | 4.916019 | 4.522003 | -0.22845 |
| NA | 4.164803 | 3.266688 | NA | 4.142362 | 2.275191 | -0.36696 | NA | 4.957289 | NA |
| 1.284488 | NA | NA | -3.99368 | 5.016867 | 2.713541 | -0.29822 | 1.72173 | 5.694759 | 2.474808 |
| 0.933741 | NA | NA | -4.39345 | 4.555363 | 1.253322 | -0.31001 | 4.738715 | 4.688951 | 1.981488 |
| 5.405674 | NA | NA | -2.12326 | 3.835332 | NA | NA | 5.58674 | 5.693196 | 2.948336 |
| 3.335818 | 5.192476 | 4.264305 | -3.98457 | 4.742564 | 2.723399 | -1.4552 | 4.808941 | 4.142448 | 3.385941 |
| 3.850709 | NA | NA | -3.2287 | 4.438205 | 2.351204 | 0.779973 | 5.34652 | 6.085715 | 1.097944 |
| 3.985403 | NA | NA | -6.25783 | 4.448659 | 2.036973 | -2.48943 | 1.971822 | 5.118954 | 2.786963 |
| 4.333461 | NA | NA | 1.337266 | 4.83727 | 2.570699 | -0.51343 | 2.64881 | 6.040302 | 2.598219 |
| 4.222042 | NA | NA | NA | 3.113953 | 1.954384 | 0.049446 | 3.624556 | 4.319504 | 0.957843 |
| 3.378034 | NA | NA | 1.043775 | 4.395722 | 2.944081 | -0.28589 | 3.627363 | 5.44393 | -1.44701 |
| 3.265218 | NA | NA | -6.51526 | 2.691854 | 2.079484 | -0.01553 | 4.513282 | 4.657807 | 1.165848 |
| 4.912065 | NA | NA | -0.14943 | 4.435224 | 3.16162 | -0.03699 | 5.182024 | 6.165986 | 1.391036 |
| 4.384072 | NA | NA | NA | 4.833598 | 2.312631 | -0.29613 | 3.098957 | 5.780269 | 1.170084 |
| 3.787893 | NA | NA | -3.55693 | 4.62237 | 1.881089 | 0.801748 | 4.973614 | 5.19102 | 3.310014 |
| 3.737656 | NA | NA | -1.99063 | 4.772858 | 2.388631 | 0.648158 | NA | 4.972826 | NA |
| 4.49653 | NA | NA | -4.27539 | 5.218579 | 2.897271 | 1.097585 | 4.913082 | 5.886459 | 3.157293 |
| 5.333821 | NA | NA | -2.50206 | 2.973241 | 2.929444 | -0.43469 | 4.851803 | 4.063064 | 2.802647 |
| 3.897894 | NA | NA | -1.01624 | 4.42561 | -0.37596 | -0.10578 | 3.448789 | 4.845398 | 1.00321 |
| 4.409235 | NA | NA | -4.50444 | 4.733058 | 1.374744 | 0.496627 | 4.203331 | 5.17324 | 2.438942 |
| 2.561016 | 3.342611 | 3.399407 | -2.96058 | 3.861231 | 2.179127 | 0.162024 | 3.818731 | 5.139346 | 2.523995 |
| 3.822911 | NA | NA | -4.10942 | 3.954098 | 1.144078 | -0.62433 | 3.760119 | 5.620482 | 1.14982 |
| 5.059295 | NA | NA | -4.15703 | 3.339621 | 1.539517 | 0.529752 | 3.600698 | 5.288163 | 0.22996 |
| 4.795227 | NA | NA | -3.89697 | 5.050817 | 2.316632 | 1.526029 | 4.909215 | 5.107634 | 1.626645 |
| 4.644993 | NA | NA | -5.65225 | 3.202167 | 2.924206 | 0.178694 | 4.077041 | 4.900073 | 0.571156 |
| 4.37089 | NA | NA | -3.89679 | 5.087122 | 2.298487 | 0.294562 | 4.410775 | 5.718502 | 3.074841 |
| 5.138422 | NA | NA | -4.85816 | NA | NA | NA | 5.50337 | NA | 4.005554 |
| 0.390582 | 0.400391 | 0.404607 | 0.404925 | 0.40812 | 0.409604 | 0.420719 | 0.43724 | 0.439191 | 0.443492 |
| 4.231464 | 3.541271 | 3.170941 | -2.83392 | 4.502522 | 2.39326 | 0.302768 | 4.408333 | 4.92641 | 1.464426 |

| | | | | | | | | | |
|---|---|---|---|---|---|---|---|---|---|
| 3.97386 | 4.233296 | 3.643467 | -3.28878 | 4.26911 | 2.161408 | -0.02783 | 4.139168 | 5.227813 | 1.933566 |

| ZG-10 | Lenalidomi | CP724714 | JQ1 | BMS-7081 | PLX4720 (r | GDC0941 ( | AZ628 | GSK42928 | STF-62247 |
|---|---|---|---|---|---|---|---|---|---|
| 2.536418 | 5.7039 | NA | 3.907907 | NA | 6.038634 | NA | NA | NA | NA |
| NA | NA | 4.717331 | NA | 3.431391 | NA | NA | 1.251045 | 4.730134 | 4.404087 |
| NA | 3.453011 | 3.057325 | 1.32682 | 4.212875 | -0.10673 | 2.191147 | -3.59171 | 3.101455 | 3.211406 |
| 0.296913 | 3.899539 | 4.738385 | 2.537122 | 5.124274 | 4.770603 | 0.582808 | 2.518996 | 5.054248 | 2.824578 |
| 1.334246 | 4.489811 | -1.83278 | 1.285683 | 3.553578 | 4.810721 | 0.637291 | 4.655011 | 5.387271 | 4.705935 |
| 2.098555 | 4.311696 | 4.731574 | 2.802999 | 4.457781 | 5.242439 | 3.836475 | NA | 5.850624 | 5.339387 |
| NA | 4.20393 | 4.242353 | 0.719182 | 5.075171 | 4.859221 | 2.595971 | 2.344141 | 5.152487 | 5.152302 |
| 5.500267 | 6.331979 | 4.581197 | 4.083035 | 4.916935 | 6.779484 | 2.560442 | NA | 7.723007 | 6.980813 |
| 0.69809 | 3.723279 | 4.245814 | 2.054036 | 4.439284 | 4.35662 | -0.95701 | 3.433354 | 5.109573 | 3.583209 |
| 3.453158 | 3.909066 | 1.358056 | 3.529388 | 6.043972 | 5.775022 | 1.9163 | NA | 5.787928 | 4.452351 |
| NA | 5.294231 | 4.011174 | 1.486243 | 5.565553 | 5.056766 | -1.11862 | NA | 6.59684 | 6.03006 |
| NA | NA | 4.465689 | 3.165482 | 5.654675 | 5.187032 | 2.725953 | NA | 5.30979 | 4.170106 |
| 1.807407 | 5.157172 | 5.687126 | 2.472916 | 5.851293 | 5.625812 | 0.389611 | NA | 6.543467 | 5.781882 |
| NA | 5.240451 | 5.309812 | 2.906585 | 5.982602 | 5.721053 | -0.28509 | NA | 6.327846 | 5.999238 |
| NA | NA | 3.279656 | NA | 4.940958 | NA | NA | 4.140672 | 5.747595 | 4.41916 |
| 3.261995 | 3.924706 | 2.267182 | 3.330122 | 5.786655 | 4.847308 | 0.641938 | NA | 5.607737 | 4.626097 |
| NA | 2.923454 | 3.327487 | 2.437415 | 4.983142 | 4.650654 | -0.5867 | NA | 5.583172 | 4.978076 |
| 1.940313 | 4.456831 | 4.617184 | 2.698855 | 4.603024 | 4.453763 | 2.267856 | NA | 4.555994 | 5.181757 |
| NA | NA | 4.445575 | NA | 4.698191 | NA | NA | 3.76198 | 5.325322 | 4.663154 |
| NA | 4.306668 | 4.608959 | 2.072562 | 5.009414 | 4.884355 | -0.7698 | NA | 5.694053 | 5.025787 |
| 1.089226 | 4.444755 | 1.376192 | NA | 5.161619 | 4.99592 | NA | NA | 5.281371 | 4.91648 |
| 2.364385 | 5.185184 | NA | 2.572672 | NA | 5.674557 | 1.12244 | NA | 3.578112 | 2.976872 |
| 1.059341 | 3.994355 | -0.65612 | 2.873695 | 5.327414 | 4.408168 | 0.516667 | 3.778453 | 6.000631 | 5.329266 |
| 5.069711 | 3.739158 | 5.07138 | 3.06625 | 4.077648 | 5.419375 | 1.412131 | NA | 6.270715 | 5.429825 |
| NA | 3.732684 | 3.991571 | 1.395178 | 4.390961 | 3.853204 | 0.32947 | NA | 4.633517 | 4.435858 |
| 2.060377 | 4.212928 | 2.322885 | 2.314562 | 5.229912 | 3.848858 | 0.850796 | NA | 6.055491 | 5.378402 |
| NA | NA | 4.400471 | 2.249383 | 4.139268 | 4.458524 | 1.904805 | NA | 5.254532 | 2.724919 |
| 0.8786 | 4.782161 | 4.138263 | 2.202587 | 5.410125 | 4.309195 | -0.02787 | NA | 5.065431 | 4.582734 |
| 0.625563 | 4.06707 | 3.468523 | 2.360908 | 4.764401 | 4.663493 | 1.315404 | NA | 5.012599 | 3.8063 |
| NA | 4.755848 | 5.240555 | 2.709757 | 5.509458 | 4.008151 | 1.116339 | NA | 6.033226 | 4.394409 |
| NA | NA | 5.115127 | 2.383394 | 5.112713 | 4.98457 | 1.983145 | NA | 5.577461 | 5.024447 |
| NA | 4.550098 | 1.784899 | 2.805822 | 3.718668 | 5.036222 | 0.066973 | NA | 5.501089 | 4.473592 |
| NA | 3.903715 | 4.834446 | 2.421412 | 4.766817 | 4.18406 | 0.830304 | NA | 5.548994 | 3.834253 |
| 1.51376 | 4.467415 | 5.218975 | 2.442934 | 5.2174 | 5.074242 | 0.886926 | NA | 5.579547 | 5.200789 |
| NA | 4.805975 | 4.247101 | 2.010018 | 5.811698 | 4.731131 | 0.51153 | NA | 6.401142 | 5.269962 |
| 0.976056 | 4.518637 | 4.743104 | 2.760001 | 5.144797 | 5.0609 | -0.04827 | NA | 5.941885 | 5.073456 |
| NA | 4.856242 | 5.213792 | 0.649173 | 5.590569 | 4.390585 | 1.389918 | NA | 6.240997 | 5.632067 |
| NA | 3.838311 | 3.751538 | 2.140987 | 4.59736 | 4.683913 | 0.918419 | 1.511605 | 3.449596 | 2.964436 |
| 2.944343 | 3.751108 | 4.754471 | 1.835488 | 4.83775 | 3.774477 | -0.48611 | NA | 5.636745 | 4.023003 |
| NA | 4.272166 | 5.473571 | 2.809466 | 3.908963 | 3.648978 | 0.13232 | NA | 4.104334 | 5.2309 |
| 1.208446 | 4.737316 | 5.288879 | 2.749035 | 3.955445 | 4.285989 | 1.229271 | NA | 5.53055 | 5.269724 |
| 2.224619 | 3.196057 | 4.913912 | 2.075706 | 4.927908 | 5.027678 | 2.754775 | NA | 6.061867 | 5.17267 |
| NA | 3.638746 | 4.249843 | 2.620953 | 4.316316 | 4.963929 | 1.893711 | NA | 3.694939 | 4.4562 |
| NA | 4.543497 | NA | 2.906068 | NA | 4.800859 | 0.837463 | NA | NA | NA |
| 0.452704 | 0.460105 | 0.468968 | 0.471545 | 0.484368 | 0.487226 | 0.495609 | 0.500663 | 0.502421 | 0.511912 |
| 2.292736 | 4.468204 | 3.694386 | 2.546487 | 4.977833 | 4.879275 | 1.15989 | 2.107358 | 5.539363 | 4.814144 |

| | | | | | | | | | |
|---|---|---|---|---|---|---|---|---|---|
| 1.834536 | 4.273917 | 4.083246 | 2.351167 | 4.817701 | 4.606853 | 0.861282 | 3.017346 | 5.338966 | 4.61158 |

| Bicalutami | Parthenoli | PLX4720 | A-770041 | XMD13-2 | XMD8-92 | EHT 1864 | OSI-930 | QL-XII-47 | Bortezomi |
|---|---|---|---|---|---|---|---|---|---|
| NA | NA | 6.397047 | NA | NA | 4.919018 | 3.884064 | NA | NA | NA |
| 2.547852 | 3.450705 | NA | 3.75037 | 2.410155 | NA | NA | 3.361369 | 0.497197 | -4.89517 |
| 1.359969 | 1.263522 | -0.82879 | 2.346526 | 2.327331 | NA | 2.659983 | 3.169406 | 0.101164 | -6.60958 |
| 2.761825 | 3.695274 | 4.872425 | 4.365073 | 2.956711 | 1.645437 | 3.170025 | 3.422082 | 3.059967 | -5.49881 |
| 3.013135 | 4.910317 | 4.407884 | 1.313175 | 2.631952 | 5.366554 | 3.908555 | 3.445541 | 2.041085 | -3.91153 |
| 3.02271 | NA | 4.952882 | NA | 5.145228 | 3.778843 | 4.836505 | 4.683782 | 2.694426 | NA |
| 2.765501 | 3.902965 | 4.603157 | NA | 3.09006 | NA | 2.53975 | 3.869288 | 0.661739 | -1.5711 |
| 4.660955 | NA | 6.27458 | NA | 6.237196 | 6.779484 | 3.580429 | 4.481285 | 1.729932 | NA |
| 2.090639 | 1.393362 | 3.701884 | 1.793397 | 3.307754 | 2.461859 | 2.519045 | 3.362401 | 1.269192 | -5.79111 |
| 3.369495 | NA | 4.896998 | NA | 4.610572 | 4.313489 | 2.73373 | 4.26594 | 2.530827 | NA |
| 3.720552 | NA | 6.015767 | NA | 3.136708 | NA | 3.146433 | 4.664118 | 0.509376 | NA |
| 1.193971 | NA | NA | NA | 3.836758 | NA | 3.386843 | 3.869287 | 1.355229 | NA |
| 3.483886 | NA | 5.821083 | NA | 5.619348 | 3.653134 | 5.514207 | 3.206428 | 5.179114 | NA |
| 3.531319 | NA | NA | NA | 4.892874 | NA | 4.736156 | 5.874878 | -0.45774 | NA |
| 2.769638 | 4.360326 | NA | NA | 4.873058 | NA | NA | 4.819275 | 1.667683 | -4.33138 |
| 3.532049 | NA | 5.556501 | NA | 4.061357 | 5.445907 | 1.89673 | 4.170232 | 0.380933 | NA |
| 2.530586 | NA | 4.956707 | NA | 3.527161 | NA | 3.253737 | 3.87932 | 2.173296 | NA |
| 2.334285 | NA | 5.163732 | NA | 2.616523 | 3.266049 | 4.3568 | 5.194021 | 2.168709 | NA |
| 2.380827 | 3.923159 | NA | 4.003597 | 2.58319 | NA | NA | 3.117867 | 2.833587 | -3.2566 |
| 1.996173 | NA | 2.651349 | NA | 5.025787 | NA | 2.453324 | 2.981734 | 5.025787 | NA |
| 2.105332 | NA | 4.499728 | NA | 3.71545 | 3.280698 | 3.455017 | 4.146008 | 1.695869 | NA |
| 2.790273 | NA | 3.785553 | NA | NA | 5.087666 | 4.622714 | NA | -1.19029 | NA |
| 3.007436 | 4.588734 | 5.257707 | 2.699687 | 2.709658 | 1.395726 | 2.676416 | 3.287362 | -0.10266 | -3.20881 |
| 3.208567 | NA | 4.43554 | NA | 3.759721 | 4.583927 | 3.61501 | 4.548996 | 0.00109 | NA |
| 2.08193 | NA | 4.025146 | NA | 3.971622 | NA | 2.974649 | 2.813081 | 2.484041 | NA |
| 2.489802 | NA | 5.314039 | NA | 2.191016 | 5.181378 | 3.602586 | 5.092307 | -1.36003 | NA |
| 2.06921 | NA | 3.269433 | NA | 1.973287 | NA | 3.957975 | 3.782555 | 2.709855 | NA |
| 3.331215 | NA | 5.565194 | NA | 4.347888 | 4.019416 | 3.436916 | 4.823234 | 1.166001 | NA |
| 2.463749 | NA | 4.731559 | NA | 2.199233 | 3.710992 | 2.645402 | 2.717346 | 1.406114 | NA |
| 3.111884 | NA | 4.10451 | NA | 4.226668 | NA | 4.510105 | 3.393082 | 3.211278 | NA |
| 2.792447 | NA | 2.612053 | NA | 4.510584 | NA | 3.624734 | 5.061338 | 2.361017 | NA |
| 2.010334 | NA | 4.629329 | NA | 2.128697 | NA | 2.225785 | 2.340742 | 0.23306 | NA |
| 2.364574 | NA | NA | NA | 2.81879 | NA | 3.834305 | 4.579537 | 2.230799 | NA |
| 2.903521 | NA | 4.581134 | NA | 5.050354 | 3.156761 | 3.851321 | 4.925875 | 3.277316 | NA |
| 3.384238 | NA | 5.748161 | NA | 4.662144 | NA | 5.133982 | 5.013009 | 2.740189 | NA |
| 2.96304 | NA | 4.535049 | NA | 4.876847 | 4.39151 | 3.782066 | 4.568893 | 0.994588 | NA |
| 3.317989 | NA | 5.039287 | NA | 4.893704 | NA | 4.329962 | 4.025479 | 3.885258 | NA |
| 2.373062 | 2.84956 | 4.631648 | 2.864723 | 0.505555 | NA | 3.392886 | 2.7381 | 1.906359 | -5.54907 |
| 2.813631 | NA | 2.858282 | NA | 4.787218 | 2.522916 | 3.97913 | 3.379624 | 2.407415 | NA |
| 3.170626 | NA | 3.616509 | NA | 3.911494 | NA | 3.738559 | 4.207668 | 2.347384 | NA |
| 2.863855 | NA | 4.979141 | NA | 4.003554 | 4.406961 | 4.396638 | 5.527926 | 4.834537 | NA |
| 3.083289 | NA | 4.875281 | NA | 3.021977 | 3.814914 | 3.801887 | 4.438201 | 0.681711 | NA |
| 2.758395 | NA | 4.955335 | NA | 4.480436 | NA | 4.368792 | 2.940217 | 1.935472 | NA |
| NA | NA | 5.726514 | NA | NA | NA | 3.728166 | NA | NA | NA |
| 0.512214 | 0.517737 | 0.518614 | 0.52831 | 0.530868 | 0.533303 | 0.53507 | 0.535274 | 0.536951 | 0.539575 |
| 2.864022 | 3.282353 | 4.770847 | 2.713708 | 3.840044 | 4.162977 | 3.507687 | 4.102274 | 1.621302 | -4.65838 |

| | | | | | | | | | |
|---|---|---|---|---|---|---|---|---|---|
| 2.713416 | 3.787151 | 4.434478 | 3.189336 | 3.59812 | 3.796072 | 3.685533 | 3.935424 | 1.90863 | -4.00483 |

| CP466722 | AZD6482 | MK-2206 | LY317615 | WH-4-023 | Masitinib | Vorinostat | AZD6482 | BIRB 0796 | AV-951 |
|---|---|---|---|---|---|---|---|---|---|
| NA | NA | 1.282224 | NA | NA | NA | 3.057939 | NA | 4.326489 | NA |
| 1.809596 | 3.266918 | NA | 2.120363 | 2.97198 | 3.042514 | NA | NA | NA | 0.660119 |
| 1.831072 | 2.593086 | 2.132672 | 1.546247 | 2.721515 | 2.76691 | -0.27121 | 2.50616 | 3.803144 | -0.66395 |
| 4.382642 | 1.067217 | -1.12298 | 4.762911 | 4.194659 | 5.755504 | -0.35071 | -0.35346 | 5.09832 | -1.5444 |
| 2.696403 | 3.855292 | 1.689402 | 0.774774 | 4.317484 | 2.518304 | 0.770289 | 3.977831 | 5.455853 | 0.750395 |
| 2.598053 | 3.405041 | 3.151197 | 4.380253 | NA | 3.635445 | -0.61346 | 3.376359 | 4.884352 | 0.833489 |
| 0.815787 | 3.001049 | 4.212479 | 3.037342 | NA | 3.524046 | 0.245134 | 2.976909 | 5.12877 | 0.056506 |
| 2.355445 | 4.790718 | 4.056795 | 1.814685 | NA | 5.869393 | 2.872684 | NA | 5.103329 | 2.43777 |
| 3.058476 | 0.970491 | -2.15263 | 4.250948 | 2.117207 | 2.798484 | 0.404656 | -1.56439 | 2.574152 | 0.057577 |
| 2.559375 | 3.501467 | -0.56645 | 2.881569 | NA | 3.123291 | 0.531647 | 2.962684 | 3.530547 | 1.526932 |
| 2.137029 | -1.02002 | -1.18987 | 5.422849 | NA | 3.361768 | 1.58096 | 0.062751 | 5.135078 | 1.298026 |
| 2.619128 | -0.262 | NA | 3.51177 | NA | 3.75485 | NA | 2.508922 | NA | 1.1969 |
| 5.656698 | -1.56232 | -0.29536 | 6.463307 | NA | 6.543467 | 2.270808 | 0.722758 | 4.566562 | 1.492009 |
| 1.967771 | 1.456643 | NA | -0.86735 | NA | 3.991385 | 0.37748 | 3.306296 | NA | 1.617048 |
| 2.346386 | 3.521895 | NA | 3.097317 | NA | 3.302417 | NA | NA | NA | 0.0082 |
| 2.453153 | 0.99252 | NA | 3.081983 | NA | 3.657788 | 3.274382 | 0.360315 | 4.277054 | 1.293333 |
| 3.280163 | 1.154864 | -0.54533 | 4.770307 | NA | 4.059649 | 0.654569 | 1.378839 | 3.489578 | 0.391731 |
| 2.533736 | 1.159535 | 4.23485 | 3.314854 | NA | 5.068172 | 2.57675 | 3.215879 | 5.060236 | 0.225215 |
| 3.640095 | 1.217932 | NA | 5.376397 | 3.928534 | 4.190243 | NA | NA | NA | 0.004591 |
| 5.133508 | 4.144686 | 0.889458 | 5.654872 | NA | 4.9238 | 1.746986 | 0.948264 | 4.54923 | 0.64376 |
| 2.502977 | 2.327714 | -0.97156 | 2.117381 | NA | 3.266141 | 1.055134 | 2.251859 | 2.053797 | 0.762997 |
| 1.59116 | NA | 1.695474 | 4.196788 | NA | NA | 1.510364 | 4.932229 | 5.879653 | NA |
| 2.39283 | 2.415071 | 1.111107 | 1.778612 | 3.995365 | 2.87422 | 1.531145 | 2.182119 | 5.222958 | 0.571603 |
| 2.865817 | -0.51512 | 1.507246 | 2.61535 | NA | 4.220723 | 1.92473 | 0.568571 | 5.554381 | 0.822676 |
| 2.835293 | 2.552237 | 1.870747 | 4.890529 | NA | 4.262819 | 0.832658 | 0.205237 | 4.407907 | -0.03471 |
| 1.855697 | 1.298057 | 2.086217 | 0.673253 | NA | 2.686664 | 3.132985 | 1.271514 | 4.004366 | 0.760411 |
| 2.907257 | 2.607442 | 1.888485 | 3.792568 | NA | 2.943107 | NA | 3.901376 | 3.497808 | -0.22011 |
| 1.415561 | 1.364304 | 2.290544 | 1.45181 | NA | 3.278451 | 2.019343 | 1.594952 | 4.798581 | 0.925073 |
| 1.585841 | 0.289815 | 1.207177 | 3.818043 | NA | 3.027398 | 1.41749 | 1.417825 | 2.424403 | 0.364919 |
| 2.513798 | 1.937468 | 3.140281 | 3.45946 | NA | 5.264764 | 2.762906 | 2.139195 | 5.332518 | 0.981994 |
| 4.014095 | 2.56943 | 1.73659 | 4.549691 | NA | 4.131686 | NA | 4.165975 | 2.894018 | -0.27381 |
| 2.44435 | 1.658741 | 1.406476 | 0.922633 | NA | 3.032298 | 0.880936 | 0.004631 | 4.340727 | 0.855036 |
| 2.488727 | 0.165117 | NA | 3.669367 | NA | 3.15478 | 1.111738 | 1.175637 | NA | 0.360054 |
| 5.665262 | 3.402159 | 0.451733 | 5.454497 | NA | 5.888405 | 1.804904 | 1.270688 | 4.997014 | 0.836948 |
| 2.132497 | 2.881527 | 2.986587 | 1.887349 | NA | 3.825475 | 1.392027 | 3.43441 | 5.79429 | 0.646876 |
| 2.399835 | 1.852727 | -0.35733 | 3.338874 | NA | 5.586053 | -0.20738 | 3.250467 | 5.248738 | 0.67162 |
| 4.340351 | 1.472255 | 1.730667 | 5.01076 | NA | 5.563478 | 1.50696 | 1.195866 | 3.470843 | 1.241609 |
| 1.669206 | 3.723655 | 1.356198 | 3.32506 | 2.895908 | 3.885358 | -2.07454 | 3.657103 | 4.484989 | 0.099643 |
| 2.722144 | 3.842583 | 0.229609 | 3.337098 | NA | 5.224575 | 0.547236 | 2.10474 | 3.803429 | 0.734051 |
| 2.263693 | 3.47415 | 2.257674 | 5.594773 | NA | 6.090088 | 1.738239 | 2.124705 | 4.781338 | 0.755106 |
| 4.741604 | 3.138918 | 0.75138 | 3.327535 | NA | 4.681056 | 2.393082 | 2.777563 | 3.953325 | 0.457773 |
| 2.692994 | 1.67077 | 1.291771 | 1.803478 | NA | 3.561842 | 1.372358 | 2.654405 | 3.341751 | 0.717539 |
| 2.761228 | 2.77772 | 3.909751 | 5.775294 | NA | 3.636406 | 1.774272 | 2.541677 | 4.191653 | 0.453914 |
| NA | NA | 2.783834 | NA | NA | NA | 1.749984 | 0.987601 | 3.915872 | NA |
| 0.544794 | 0.552166 | 0.552366 | 0.559759 | 0.562223 | 0.569628 | 0.576464 | 0.579134 | 0.580286 | 0.582886 |
| 2.652995 | 1.876023 | 1.145154 | 3.19789 | 3.264569 | 3.927846 | 1.158795 | 1.81699 | 4.459533 | 0.684523 |

| | | | | | | | | | |
|---|---|---|---|---|---|---|---|---|---|
| 2.863033 | 2.17789 | 1.552088 | 3.512859 | 3.606602 | 4.133326 | 1.387981 | 2.110344 | 4.289316 | 0.547481 |

| Roscovitine | TW 37 | SB590885 | Sorafenib | PFI-1 | Camptothe | KIN001-27( | TL-1-85 | GSK69069: | JNK Inhibit |
|---|---|---|---|---|---|---|---|---|---|
| NA | 0.557385 | 5.7039 | NA | 6.210492 | 1.055615 | NA | NA | NA | 5.852651 |
| 3.27616 | NA | NA | 3.080339 | NA | NA | 5.069975 | 3.312115 | 1.037543 | NA |
| 3.338482 | -0.45385 | -0.45705 | 1.009039 | 2.891704 | -5.19213 | 3.478834 | 2.013457 | 3.573142 | 3.551988 |
| 4.706751 | -0.6148 | 4.404427 | 2.013602 | 2.543173 | -4.98581 | 4.424778 | 4.288123 | 1.673503 | 4.984626 |
| 3.175656 | 0.080146 | 4.406264 | 0.781878 | 2.933533 | -0.21506 | 4.120687 | 2.613269 | 2.065227 | 5.538863 |
| NA | -0.86152 | 4.562742 | NA | 2.333122 | -2.07517 | 6.047739 | 5.114546 | 5.384145 | 4.422683 |
| NA | 1.670145 | 3.766548 | 2.257135 | 2.536255 | -3.40426 | 4.881639 | 3.909791 | 3.755246 | 5.128294 |
| NA | 2.380536 | 6.333998 | NA | 4.241949 | -1.78222 | 6.966142 | 6.598674 | 6.532658 | 6.934865 |
| 4.198839 | -1.11618 | 3.252403 | 1.51937 | 1.987699 | -3.40958 | 3.710079 | 1.254627 | 1.747699 | 3.595685 |
| NA | -0.15588 | 3.338306 | NA | 3.693113 | 0.814402 | 6.000957 | 5.05758 | 3.094846 | 3.17322 |
| NA | -0.32603 | 5.322619 | NA | 3.285948 | -3.5408 | 5.471722 | 4.810184 | 0.037393 | 5.512723 |
| NA | -0.427 | NA | NA | 2.985186 | NA | 4.814877 | 4.018219 | 2.273206 | NA |
| NA | NA | 5.157172 | NA | 3.520147 | -1.79777 | 6.245414 | 5.591246 | 4.722095 | 4.697947 |
| NA | 0.571375 | NA | NA | 2.03751 | -2.82498 | 6.103648 | 3.819755 | 2.79399 | NA |
| 5.018348 | NA | NA | 2.530897 | NA | NA | 5.26519 | 4.210774 | 3.511428 | NA |
| NA | 0.398461 | NA | NA | 4.684152 | -1.39284 | 5.357479 | 4.163412 | 0.126306 | 5.18372 |
| NA | -1.16494 | 4.274598 | NA | 2.143095 | -3.50528 | 5.298006 | 4.302706 | 2.60973 | 4.95163 |
| NA | -0.57692 | 4.477383 | NA | 2.867722 | -4.545 | 5.082693 | 3.166806 | 4.630587 | 4.86988 |
| 3.652613 | NA | NA | 1.806033 | NA | NA | 4.452175 | 3.644313 | -0.6669 | NA |
| NA | -0.93576 | 3.118583 | NA | 1.830745 | -4.10177 | 5.495633 | 5.025787 | 5.025787 | 3.957086 |
| NA | -0.93705 | 3.524918 | NA | 3.047626 | -4.82835 | 4.791862 | 4.982797 | 3.624801 | 3.404031 |
| NA | 4.060943 | 3.251779 | NA | 4.646871 | 0.546217 | 4.540005 | 2.832179 | NA | 5.716037 |
| 5.276474 | 1.309729 | 4.590285 | 2.838109 | 2.932597 | -0.18095 | 4.481004 | 4.667829 | 0.303778 | 5.254305 |
| NA | -0.33223 | 4.885745 | NA | 3.840643 | -3.60373 | 5.68972 | 5.535279 | 2.036629 | 5.004598 |
| NA | -1.27174 | 3.732168 | NA | 2.988342 | -5.07166 | 5.082781 | 4.395485 | 2.988638 | 4.425832 |
| NA | 0.115799 | 4.330614 | NA | 5.084214 | -0.21876 | 6.048241 | 5.384624 | 0.626398 | 4.433649 |
| NA | 0.598479 | 3.90813 | NA | 2.457232 | NA | 4.674406 | 3.679303 | 3.685511 | 3.797944 |
| NA | 1.478517 | 3.40388 | NA | 2.502596 | -2.07749 | 5.450982 | 4.288551 | 1.244428 | 5.589 |
| NA | -2.18179 | 3.346959 | NA | 2.883097 | -5.07787 | 4.277016 | 3.258604 | 1.942993 | 3.573454 |
| NA | -0.84704 | 4.566788 | NA | 4.202221 | -2.31132 | 5.581839 | 4.873936 | 4.577368 | 5.470192 |
| NA | -0.11513 | 4.398264 | NA | 2.55907 | NA | 4.946748 | 5.102798 | 5.082959 | 5.091411 |
| NA | NA | 3.968028 | NA | 2.100808 | -2.28135 | 4.759403 | 2.387096 | 0.724461 | 4.520177 |
| NA | -0.38057 | NA | NA | 3.874108 | -2.38295 | 5.489832 | 1.625604 | 2.227593 | NA |
| NA | -2.33301 | 4.502111 | NA | 2.576234 | -3.04869 | 5.640648 | 5.218569 | 5.218975 | 5.049878 |
| NA | 0.260653 | 5.104593 | NA | 3.548247 | -1.9367 | 5.115672 | 4.06472 | 5.070195 | 5.763925 |
| NA | 0.581329 | 3.012324 | NA | 2.874966 | -1.58087 | 5.344175 | 5.082247 | 5.186213 | 5.24474 |
| NA | -0.82457 | 4.916647 | NA | 3.233746 | -3.69785 | 4.67535 | 5.016264 | 5.63351 | 5.08956 |
| 3.98705 | -0.85147 | 3.415135 | 1.790895 | 2.668251 | -2.29173 | 4.662743 | 3.495012 | 2.324899 | 4.29094 |
| NA | -1.01786 | 2.371393 | NA | 2.41214 | -3.43303 | 5.114856 | 4.221505 | 3.84747 | 3.989551 |
| NA | -0.57276 | 3.050205 | NA | 1.855949 | -3.27093 | 4.964233 | 2.97882 | 4.505363 | 5.190215 |
| NA | 1.191745 | 4.811063 | NA | 3.644182 | -2.84862 | 5.049961 | 3.237745 | 3.927927 | 4.330181 |
| NA | 0.270975 | 4.238534 | NA | 3.344089 | -2.80743 | 4.822951 | 5.361672 | 2.515435 | 4.36657 |
| NA | -2.04639 | 2.882126 | NA | 1.777414 | -4.41227 | 5.631471 | 4.820757 | 5.125273 | 4.889857 |
| NA | -0.19667 | 5.033367 | NA | 2.43221 | -2.50186 | NA | NA | NA | 5.242196 |
| 0.585129 | 0.593007 | 0.59515 | 0.599521 | 0.610895 | 0.611135 | 0.612186 | 0.614983 | 0.623649 | 0.623933 |
| 3.952373 | -0.0026 | 4.195639 | 1.884609 | 3.180925 | -2.45339 | 5.196462 | 4.014428 | 2.915809 | 4.885627 |

| | | | | | | | | | |
|---|---|---|---|---|---|---|---|---|---|
| 4.305379 | -0.20733 | 3.931818 | 2.145012 | 3.012704 | -2.75739 | 5.071348 | 4.20726 | 3.199155 | 4.736889 |

| LFM-A13 | GDC0941 | Rapamycin | YM155 | BIX02189 | BMS-5369 | GSK26996 | Dabrafeni | PHA-6657 | VX-702 |
|---|---|---|---|---|---|---|---|---|---|
| NA | 1.249398 | NA | NA | NA | 3.357274 | NA | 5.278504 | NA | 4.787609 |
| 5.476242 | NA | -3.42599 | -5.34776 | 3.624346 | NA | 2.415258 | NA | 3.504858 | NA |
| 3.387497 | 2.133908 | -2.98639 | -2.54843 | 4.161157 | 2.132001 | 2.470136 | -4.24168 | 2.588793 | 2.590836 |
| 3.396823 | -1.62991 | -2.05333 | -3.54137 | 5.793705 | 2.257062 | 3.713936 | 4.795094 | 3.49112 | 2.900972 |
| 5.547961 | 0.484729 | -0.30514 | -5.52965 | 4.933493 | 4.534297 | 2.895937 | 5.432845 | 4.002914 | 2.731888 |
| 3.94792 | 4.165199 | NA | -4.49332 | 5.317982 | 4.054 | NA | 3.951821 | NA | 3.441131 |
| 5.434201 | 2.615637 | -1.23472 | -1.50913 | 5.439855 | 3.100371 | 2.310455 | 4.870824 | 3.051725 | 2.019426 |
| 7.442603 | 2.534675 | NA | -3.22885 | 6.858649 | 3.996896 | NA | 6.500473 | NA | 5.265397 |
| 4.750392 | -1.2203 | -2.75719 | -4.08266 | 3.841634 | 1.775889 | 2.871075 | 4.303997 | 2.693856 | 2.806988 |
| 4.852346 | 0.949836 | NA | -1.78819 | 5.696268 | 4.915682 | NA | 3.449199 | NA | 4.400454 |
| 5.517927 | -0.56694 | NA | -2.33413 | 4.059258 | 2.308633 | NA | 4.178035 | NA | 3.886784 |
| 5.599544 | NA | NA | -3.41179 | 5.188256 | 3.544132 | NA | 5.263554 | NA | NA |
| 4.158826 | 0.588649 | NA | -2.76066 | 6.526608 | 3.381149 | NA | 5.010538 | NA | 4.240882 |
| 4.655639 | NA | NA | -5.25087 | 3.892149 | 1.860462 | NA | 5.713451 | NA | 3.710282 |
| 5.260088 | NA | -2.08591 | -4.99198 | 5.634472 | NA | 3.37436 | NA | 2.4148 | NA |
| 3.663836 | 2.314464 | NA | -1.46534 | 6.114408 | 4.380634 | NA | 5.633429 | NA | 4.12698 |
| 4.778628 | -0.78428 | NA | -0.91842 | 5.367814 | 2.367489 | NA | 4.529021 | NA | 3.007791 |
| 5.633359 | 3.151547 | NA | -5.5209 | 4.185303 | 2.808325 | NA | 5.047289 | NA | 3.563687 |
| 4.684448 | NA | -1.52716 | -5.50781 | 4.554276 | NA | 2.214684 | NA | 3.073812 | NA |
| 3.975969 | -0.12415 | NA | -0.87846 | 5.695218 | 0.515368 | NA | 3.029935 | NA | 3.373204 |
| 5.458987 | -0.28426 | NA | -4.78242 | 5.795294 | 3.376638 | NA | NA | NA | 3.398733 |
| 3.12326 | 0.944726 | NA | -4.09147 | 4.785261 | 4.458911 | NA | 5.341524 | NA | 4.175906 |
| 5.101168 | 1.156335 | -1.50105 | -2.99841 | 5.167131 | 3.830732 | 3.610317 | 3.652748 | 3.497937 | 3.699121 |
| 5.420288 | 0.580981 | NA | -5.10409 | 6.196468 | 4.372039 | NA | 5.334565 | NA | 3.208965 |
| 4.797895 | 1.149131 | NA | -1.45124 | 5.118979 | 1.60151 | NA | 1.66722 | NA | 2.544538 |
| 5.825879 | 2.9389 | NA | -4.79934 | 6.055491 | 3.584897 | NA | 5.028654 | NA | 3.750092 |
| 5.091005 | 3.617452 | NA | -4.86959 | 3.691876 | 2.431166 | NA | 3.45486 | NA | NA |
| 3.913423 | 2.181634 | NA | -2.38589 | 5.293912 | 4.412101 | NA | 4.194485 | NA | 4.030817 |
| 4.776085 | 1.987233 | NA | -5.37678 | 4.153758 | 1.782869 | NA | 4.132821 | NA | 3.150779 |
| 5.936644 | 1.472859 | NA | -5.12463 | 6.119676 | 3.614445 | NA | 4.728852 | NA | 3.868855 |
| 5.447715 | 1.421294 | NA | -2.27686 | 5.44601 | 2.528531 | NA | 4.44324 | NA | NA |
| 4.681711 | 2.090643 | NA | -5.73057 | 4.687125 | 3.928087 | NA | 4.579079 | NA | 3.522475 |
| 3.306517 | NA | NA | -1.2139 | 3.050267 | 2.679726 | NA | 4.78331 | NA | 2.710512 |
| 5.574843 | 0.129865 | NA | 0.428894 | 5.350479 | 2.903844 | NA | 5.073451 | NA | 3.585821 |
| 4.361496 | 0.982115 | NA | -2.89027 | 5.981214 | 0.90483 | NA | 5.439417 | NA | 4.069941 |
| 3.609164 | -0.25825 | NA | -4.34924 | 4.955413 | 4.195284 | NA | 3.746763 | NA | 3.328178 |
| 4.789999 | 0.762204 | NA | -3.66418 | 5.486901 | 3.618682 | NA | 4.589388 | NA | 4.000356 |
| 4.956799 | 1.659058 | -2.55732 | -4.621 | 4.245416 | 1.952249 | 1.987512 | 4.187937 | 2.146295 | 2.42524 |
| 4.727149 | -0.7617 | NA | -4.11894 | 4.742537 | 2.703458 | NA | 2.106643 | NA | 3.106016 |
| 4.741457 | 2.496269 | NA | -4.9135 | 3.287472 | 1.348484 | NA | 1.431342 | NA | 3.842374 |
| 5.681666 | 1.536831 | NA | -1.76388 | 4.147705 | 3.186646 | NA | 4.155016 | NA | 3.716111 |
| 5.382655 | 0.903123 | NA | -4.31239 | 4.474958 | 4.143913 | NA | 5.198168 | NA | 3.156465 |
| 4.002146 | 2.418971 | NA | -5.44376 | 5.711313 | 2.962048 | NA | 3.236432 | NA | 2.997512 |
| NA | 4.469356 | NA | NA | NA | 4.586125 | NA | 4.612674 | NA | 3.950622 |
| 0.641583 | 0.642432 | 0.642854 | 0.646097 | 0.667223 | 0.667542 | 0.669501 | 0.682774 | 0.686679 | 0.687083 |
| 4.91199 | 1.1419 | -2.12124 | -3.45432 | 5.096197 | 3.173394 | 2.864451 | 4.357274 | 3.106867 | 3.565407 |

| | | | | | | | | | |
|---|---|---|---|---|---|---|---|---|---|
| 4.774735 | 1.394609 | -1.86184 | -3.68959 | 4.967766 | 3.024903 | 2.604171 | 4.089522 | 2.906014 | 3.461419 |

| NVP-TAE68 | AR-42 | SNX-2112 | QL-XI-92 | EKB-569 | GW-2580 | Pyrimethar | GNF-2 | PXD101, Be | NU-7441 |
|---|---|---|---|---|---|---|---|---|---|
| NA | NA | NA | NA | NA | NA | NA | NA | NA | 4.590103 |
| 2.016292 | -2.0436 | -2.61715 | 3.584383 | -0.46831 | 5.613968 | 2.611581 | 2.954181 | -1.21619 | NA |
| -0.17547 | -1.72365 | -3.59917 | 1.967222 | 1.29196 | 4.892228 | 2.717982 | 2.110086 | -1.12735 | 1.96307 |
| 0.774207 | 0.797842 | 3.717584 | 2.877819 | 3.714263 | 4.205086 | 4.499915 | 2.892242 | 0.751473 | 0.734275 |
| 3.203112 | 0.044571 | -0.08422 | 3.011099 | NA | 4.134641 | 5.34652 | 2.535645 | 1.774144 | 3.530983 |
| NA | -1.06688 | 2.603974 | 4.483002 | 2.796068 | 6.045111 | NA | NA | -0.37275 | 2.597355 |
| 1.748977 | -0.97336 | 1.443709 | 3.160549 | 1.666127 | 5.75684 | 4.409191 | NA | -0.17878 | 2.366464 |
| NA | 6.112678 | 2.271654 | 6.292572 | 5.39232 | 7.356 | NA | NA | 5.699988 | 4.975077 |
| 1.895576 | -1.6816 | -0.18107 | 4.076618 | 2.739871 | 4.084295 | 5.109573 | 2.360701 | -1.15036 | 1.024513 |
| NA | -0.25234 | 1.072175 | 4.988167 | -0.66079 | 6.155947 | NA | NA | 1.261325 | 4.341931 |
| NA | -1.3656 | 0.139512 | 5.48343 | -0.21696 | 6.702246 | NA | NA | -1.11671 | 2.96854 |
| NA | 0.586563 | 0.90932 | 4.245013 | 2.952663 | 5.363918 | NA | NA | 0.103696 | NA |
| NA | 3.113583 | 3.655188 | 5.873623 | 4.276903 | 5.111961 | NA | NA | 3.398082 | 2.260763 |
| NA | -0.70539 | 2.781106 | 5.999238 | 4.244634 | 6.092957 | NA | NA | 0.157256 | NA |
| 2.830111 | -0.10292 | 3.666616 | 4.95651 | 2.31176 | 5.465499 | 5.140448 | NA | 0.605446 | NA |
| NA | 1.511 | 0.858121 | 4.127296 | -1.79332 | 6.117622 | NA | NA | 1.479238 | 4.117287 |
| NA | -1.43612 | -1.87605 | 4.238667 | 0.145137 | 5.660893 | NA | NA | -0.43862 | 2.436125 |
| NA | -0.4402 | -1.96379 | 4.749353 | 1.376677 | 5.666044 | NA | NA | -0.09616 | 3.042122 |
| 2.655506 | -1.50102 | -2.52856 | 4.46885 | 2.647723 | 5.375217 | 4.912884 | 2.558277 | -1.19106 | NA |
| NA | 2.345573 | 3.639045 | 5.025787 | 3.611558 | 5.406631 | NA | NA | 3.358509 | 2.628234 |
| NA | -1.02321 | 0.528462 | 4.306227 | -1.54756 | 5.339713 | NA | NA | -0.56892 | 1.570052 |
| NA | -1.47601 | -0.85805 | NA | NA | NA | NA | NA | -1.06113 | 2.792911 |
| 3.39431 | 0.7743 | -1.54408 | 3.745498 | -2.36106 | 5.41719 | 5.633895 | 3.252834 | 1.243263 | 3.598144 |
| NA | 1.088658 | 1.429839 | 3.597343 | 1.5278 | 5.664283 | NA | NA | 1.270848 | 3.659285 |
| NA | 2.184248 | 2.175656 | 4.103933 | 2.191936 | 4.815327 | NA | NA | 3.132969 | 0.795621 |
| NA | 0.249005 | 1.706811 | 3.924499 | -0.01282 | 5.510698 | NA | NA | 1.202153 | 3.376526 |
| NA | -0.54507 | -1.83649 | 3.266205 | 0.324193 | 5.330705 | NA | NA | 0.011981 | 3.025745 |
| NA | -0.23256 | 0.670988 | 3.580099 | 1.009059 | 5.841591 | NA | NA | 0.474317 | 3.406957 |
| NA | 1.006845 | -1.8572 | 3.371741 | -0.78401 | 4.42031 | NA | NA | 1.973299 | 2.107736 |
| NA | -0.81627 | 1.164362 | 4.789621 | 2.60406 | 6.035552 | NA | NA | -0.37295 | 3.222104 |
| NA | 0.072311 | 3.364227 | 5.053021 | 3.699358 | 5.096219 | NA | NA | 0.526591 | 1.200891 |
| NA | -2.85217 | -3.05084 | 3.527745 | 1.274838 | 4.479081 | NA | NA | -2.61065 | 3.470698 |
| NA | -0.25173 | -0.10873 | 4.555921 | 2.7588 | 5.563969 | NA | NA | -0.68504 | NA |
| NA | 1.10849 | 2.618822 | 5.213156 | 3.808814 | 5.807856 | NA | NA | 1.806608 | 1.967815 |
| NA | 0.490569 | -1.99764 | 4.721867 | 1.107772 | 6.461014 | NA | NA | 0.671445 | 2.97412 |
| NA | -0.11781 | 0.923613 | 4.882709 | 2.845449 | 5.901089 | NA | NA | 0.883955 | 3.021574 |
| NA | 1.280145 | 0.845171 | 5.067304 | 4.212978 | 5.187076 | NA | NA | 1.803397 | 2.885423 |
| 0.503586 | 1.005223 | NA | 3.413542 | 0.900303 | 4.546036 | 3.268316 | 2.307708 | 2.034144 | 2.781791 |
| NA | 0.913075 | 0.739341 | 4.014137 | 1.270732 | 5.447754 | NA | NA | 1.132206 | 2.303377 |
| NA | 0.225071 | -1.36377 | 4.609918 | 0.40469 | 6.203621 | NA | NA | 0.463205 | 3.487001 |
| NA | 2.470732 | 2.555212 | 4.555687 | 3.543092 | 6.191274 | NA | NA | 2.459761 | 2.516624 |
| NA | 0.302319 | 0.525769 | 3.886436 | 1.25657 | 6.062049 | NA | NA | 0.367882 | 3.108078 |
| NA | -0.53843 | 3.714137 | 3.94367 | 2.490622 | 4.919361 | NA | NA | NA | 3.180741 |
| NA | NA | NA | NA | NA | NA | NA | NA | NA | 3.616793 |
| 0.688269 | 0.690206 | 0.690724 | 0.695111 | 0.697729 | 0.698592 | 0.700278 | 0.701521 | 0.702003 | 0.702838 |
| 1.756114 | 0.022034 | 0.752795 | 4.35968 | 1.860563 | 5.554427 | 4.262173 | 2.570571 | 0.560809 | 2.924901 |

| | | | | | | | | | |
|---|---|---|---|---|---|---|---|---|---|
| 2.184467 | 0.246492 | 0.477337 | 4.234372 | 1.616037 | 5.459317 | 4.605032 | 2.706273 | 0.763616 | 2.779093 |

| Nilotinib | AUY922 | MPS-1-IN-1 | CAY10603 | CH542480 | NSC-20789 | Afatinib | Afatinib (re | XL-184 | CI-1040 |
|---|---|---|---|---|---|---|---|---|---|
| 4.787609 | NA | NA | NA | NA | NA | -1.88671 | -0.3036 | NA | 3.972134 |
| NA | -2.78539 | 2.786613 | -0.72659 | 2.458105 | 4.17334 | NA | NA | 2.606874 | NA |
| 0.56163 | -4.54199 | 3.045482 | 0.335853 | 3.501301 | 4.069997 | 1.143399 | 3.819516 | 2.68384 | -1.99169 |
| 1.008982 | -3.01248 | 5.743041 | 1.476024 | 3.31457 | 5.570999 | -2.00007 | 2.620204 | 4.193994 | 3.164951 |
| 3.953522 | -2.57699 | 4.044058 | 0.664133 | 3.465527 | 5.346213 | -3.16462 | -1.40652 | 3.565894 | 5.166616 |
| 3.75099 | -1.81083 | 5.978935 | 0.980936 | 3.735914 | 5.941697 | 2.024299 | 5.238851 | 4.590783 | 3.395788 |
| 2.834243 | -1.42255 | 5.502872 | 1.043081 | 3.545173 | 4.552227 | 1.200269 | 4.542536 | 4.458095 | 3.29204 |
| 5.146388 | 3.268696 | 7.230106 | 3.474248 | 5.296054 | 3.541742 | 3.662703 | 6.779484 | 6.187347 | 4.021057 |
| 2.806671 | -2.13025 | 3.362133 | -0.194 | 1.357948 | 2.108958 | 0.623924 | 4.357172 | 2.542324 | 2.787536 |
| 2.95291 | -2.70842 | 6.456344 | 2.178219 | 4.386231 | 6.305868 | -1.12296 | 1.336209 | 5.312523 | 3.409918 |
| 4.358088 | -1.52468 | 3.194235 | 0.395272 | 3.524045 | 4.07233 | 0.675444 | 2.85587 | 3.183549 | 4.462912 |
| NA | -2.66888 | 4.032349 | 0.547227 | 2.590525 | 4.164566 | NA | 4.46011 | 3.435666 | NA |
| 4.240882 | -1.51007 | 6.462277 | 3.133664 | 4.330774 | 6.418955 | 2.590184 | 4.786839 | 5.034203 | 3.781467 |
| 4.212238 | NA | 5.111133 | 0.324902 | 4.124958 | 4.245491 | 2.660508 | 4.345098 | 2.818814 | 4.230546 |
| NA | -1.07979 | 4.781552 | 1.805396 | 4.359675 | 3.450768 | NA | NA | 2.745184 | NA |
| 3.499445 | -2.77732 | 6.00398 | 2.04492 | 3.28876 | 5.908364 | -2.23483 | -0.46333 | 4.260668 | 3.122363 |
| 2.588699 | -3.70537 | 5.006031 | 0.469085 | 4.289961 | 5.271973 | 0.423401 | 1.900493 | 3.188149 | 2.361759 |
| 3.556205 | -3.6823 | 5.758128 | 0.795697 | 4.369346 | 3.813343 | 1.934914 | 4.616276 | 2.76851 | 3.537881 |
| NA | -3.23394 | 3.764264 | 0.172634 | 3.479885 | 5.349756 | NA | NA | 3.84797 | NA |
| 3.006294 | -5.35485 | 5.694247 | 2.380025 | 4.19345 | 5.695218 | 1.139618 | 4.882209 | 4.331664 | 3.428073 |
| 3.199612 | -2.33016 | 5.336855 | 0.557899 | 4.032216 | 3.486565 | -3.12625 | -1.27523 | 3.328269 | 2.968502 |
| 4.207685 | -2.30323 | 6.571345 | 0.39852 | NA | 4.986046 | -1.03751 | 0.362722 | 4.841433 | 4.352849 |
| 3.699121 | -4.02412 | 5.012549 | 0.484938 | 4.624721 | 5.288635 | -2.98322 | 0.22293 | 2.747032 | 5.279098 |
| 3.658667 | -1.15198 | 5.54564 | 2.027757 | 4.662367 | 4.951853 | -0.38511 | 3.002986 | 4.235933 | 2.61742 |
| 2.363225 | -4.94243 | 5.11747 | 3.238281 | 3.652447 | 4.569701 | 1.02758 | 3.11338 | 3.753272 | 2.611736 |
| 3.281603 | -0.02729 | 6.044656 | 1.914514 | 4.474782 | 5.928987 | 1.289541 | 0.753197 | 4.558649 | 3.203268 |
| NA | -2.2824 | 4.125232 | 0.86548 | 3.080379 | 4.064545 | NA | 2.877776 | 2.536355 | NA |
| 3.763496 | -3.61863 | 5.680595 | 1.333354 | 3.988757 | 5.807761 | -0.38029 | 2.323559 | 4.313727 | 5.322402 |
| 2.525554 | -3.07919 | 3.241009 | 1.38337 | 1.863045 | 2.750155 | -1.08792 | 2.022896 | 3.394767 | 1.488465 |
| 3.849504 | -3.26979 | 4.193375 | 0.56772 | 2.666713 | 3.201261 | -1.20581 | 3.170063 | 4.115023 | 2.126584 |
| NA | -0.38714 | 5.096952 | 0.937424 | 4.25137 | 5.782732 | NA | 3.982533 | 4.205803 | NA |
| 3.627897 | -1.53489 | 4.4017 | -0.88109 | 3.236952 | 4.9104 | 1.249469 | 2.435385 | 3.734566 | 3.962318 |
| 1.515547 | -2.87048 | 4.758266 | -0.07433 | 2.330237 | 3.689142 | 0.749518 | 3.661102 | 3.463647 | 1.970986 |
| NA | 1.410102 | 5.882158 | 1.993231 | 2.518076 | 5.877909 | 1.994875 | 4.728739 | 4.510732 | 1.245701 |
| 4.150563 | -0.04934 | 2.855478 | 1.117743 | 2.356661 | 5.622175 | -0.28195 | 3.680258 | 1.393742 | 1.886175 |
| NA | -4.18408 | 5.051303 | 1.055878 | 4.365124 | 5.168436 | -3.34453 | 0.563277 | 2.601701 | 3.857466 |
| NA | -1.95152 | 3.645353 | 1.854792 | 3.447994 | 4.353161 | 1.797547 | 3.524212 | 4.213927 | 2.753834 |
| 2.853823 | -0.85689 | 3.728994 | 1.732658 | 3.071975 | 3.584056 | 0.496447 | 4.650091 | 2.831581 | 3.012349 |
| NA | -4.26883 | 4.652181 | 1.57092 | 4.197643 | 4.667619 | 0.067563 | 2.354498 | 3.476913 | 2.099167 |
| 3.47783 | -3.43144 | 4.441011 | 0.334517 | 4.699904 | 2.986218 | 1.471083 | 3.452226 | 2.50029 | 3.175033 |
| 3.190452 | -0.06927 | 5.826507 | 3.150546 | 4.521255 | 5.847135 | 2.508478 | 4.996183 | 3.540892 | 4.534155 |
| 2.230875 | -3.07698 | 4.346458 | 0.928982 | 3.74576 | 6.052186 | 0.744456 | 4.141739 | 3.958498 | 2.458413 |
| 2.682328 | -2.45218 | 5.793649 | 1.527167 | 1.514044 | 4.530666 | 1.603442 | 3.997606 | 4.35581 | 1.417166 |
| 3.69321 | NA | NA | NA | NA | NA | 2.560137 | 3.938038 | NA | 5.083925 |
| 0.705399 | 0.709396 | 0.717811 | 0.721082 | 0.728588 | 0.729988 | 0.730423 | 0.731899 | 0.731946 | 0.734907 |
| 3.350567 | -2.16679 | 4.970545 | 1.102828 | 3.643463 | 4.64452 | 0.435324 | 3.092825 | 3.739789 | 3.247685 |

| | | | | | | | | | |
|---|---|---|---|---|---|---|---|---|---|
| 3.209331 | -2.37364 | 4.83229 | 1.222917 | 3.540657 | 4.766093 | 0.211616 | 2.862495 | 3.631688 | 3.080656 |

| UNC0638 | AS605240 | PF-470867 | XMD11-85 | TPCA-1 | Lapatinib | PAC-1 | CX-5461 | Phenformi | BMS-3455 |
|---|---|---|---|---|---|---|---|---|---|
| 3.973989 | NA | 4.811262 | 5.563582 | NA | NA | NA | NA | NA | NA |
| NA | 1.649617 | NA | NA | 2.010406 | 3.073672 | 1.874246 | 0.026409 | 5.890749 | 1.480605 |
| 2.282952 | -3.94395 | 2.396599 | NA | 2.445204 | 2.338621 | 0.316781 | -0.62228 | 4.396785 | 1.469439 |
| 5.67708 | 4.805242 | 4.397888 | 2.321065 | 5.793705 | 1.534564 | 2.328757 | 5.124274 | 5.039379 | 3.360024 |
| 2.592839 | 2.469928 | 4.042528 | 5.372279 | 5.195759 | -2.71281 | 0.108281 | 3.108656 | 2.799805 | 1.555763 |
| 2.839404 | 4.369991 | 4.271074 | 4.834456 | 5.153102 | NA | 3.278738 | 5.271585 | 9.523396 | 4.29979 |
| 2.225711 | 3.322724 | 3.09396 | NA | 4.084329 | NA | 3.044591 | 4.646264 | NA | 2.761855 |
| 4.252769 | 3.823031 | 6.898326 | 6.779484 | 5.333795 | NA | 4.207461 | 3.891753 | 12.80228 | 4.909071 |
| 2.104484 | 3.299267 | 3.142456 | 2.432939 | 1.299151 | 1.725485 | 2.524979 | 3.918742 | 6.85799 | 2.808349 |
| 2.953395 | 3.739449 | 2.268801 | 5.216912 | 5.173597 | NA | 1.562052 | 5.967653 | 7.62336 | 4.365702 |
| 2.957455 | 2.09103 | 3.208953 | NA | 3.008104 | NA | 3.505975 | 4.7355 | 8.424156 | 4.352806 |
| 6.135854 | 4.784876 | 4.083671 | NA | 5.379395 | NA | 0.503795 | 4.470973 | 6.299376 | 4.025994 |
| 3.817666 | 5.372965 | NA | 5.331334 | 6.526209 | NA | 4.364473 | 5.874036 | 6.176591 | 5.931134 |
| 3.179069 | 2.860118 | 5.322649 | NA | 2.874421 | NA | 0.430411 | 5.402958 | 8.946114 | 3.260641 |
| NA | 3.586397 | NA | NA | 4.840019 | 1.38754 | 3.672455 | 4.372724 | 8.74355 | 3.777648 |
| 6.326576 | 3.056168 | 1.444594 | 5.471015 | 4.407337 | NA | 3.968167 | 5.553187 | 7.012571 | 3.681208 |
| 5.561211 | 3.143113 | 3.945184 | NA | 4.279429 | NA | 2.299133 | 4.990878 | 8.920272 | 3.705488 |
| 3.645934 | 3.828779 | 3.39283 | 4.178719 | 3.343429 | NA | 1.24158 | 2.201829 | 7.01628 | 3.623507 |
| NA | 4.034479 | NA | NA | 2.34715 | 3.073812 | 2.011677 | 0.312511 | 6.737507 | 2.970077 |
| 4.544073 | 4.505537 | 3.610594 | NA | 5.695218 | NA | 1.597691 | 5.025787 | 4.681212 | 5.148813 |
| 2.782754 | 3.291874 | 3.236377 | 4.055961 | 5.335442 | NA | 0.883912 | 5.11952 | 7.676069 | 3.038119 |
| 2.951342 | 1.397719 | 4.249694 | 5.616502 | 5.680834 | NA | -0.17149 | 5.041547 | 4.650003 | 3.003501 |
| 2.541155 | 2.68208 | 2.941455 | 3.974219 | 4.811866 | -1.02581 | 2.586631 | 3.981738 | 6.227628 | 3.102062 |
| 4.083339 | 2.656543 | 3.675307 | 5.419375 | 4.905916 | NA | 3.914117 | 5.2279 | 8.014971 | 3.617251 |
| 5.054392 | 3.776507 | 0.762725 | NA | 5.118979 | NA | 1.002627 | 3.999714 | 7.655451 | 3.30105 |
| 2.904968 | 1.204318 | 3.561802 | 5.193691 | 6.055491 | NA | 1.195103 | 5.132962 | 6.621552 | 1.608832 |
| 2.301796 | 3.56096 | 4.616394 | NA | 2.903063 | NA | 2.539795 | 2.048743 | 6.552038 | 2.734961 |
| 2.357154 | 3.505871 | 2.236717 | 3.899376 | 3.787161 | NA | 1.391588 | 4.939799 | 10.04673 | 3.300083 |
| 2.554708 | 2.910153 | 3.932243 | 3.329556 | 2.927419 | NA | 3.318765 | 2.602044 | 4.141853 | 2.919517 |
| 6.032354 | 3.30267 | 4.825497 | NA | 3.613531 | NA | 1.096621 | 0.310433 | 7.938055 | 4.386855 |
| 4.675894 | 2.972683 | 3.773707 | NA | 4.465968 | NA | 2.159913 | 5.070511 | 7.581208 | 4.01852 |
| 5.798967 | 0.249085 | NA | NA | 4.495839 | NA | 3.150096 | 2.567526 | 4.34625 | 2.57175 |
| 2.533597 | 2.768857 | 4.01291 | NA | 3.862828 | NA | 3.517746 | 3.386249 | 7.792536 | 2.820095 |
| 5.041958 | 5.88745 | 4.415633 | 4.715174 | 5.888406 | NA | 3.832681 | 5.218975 | 6.972361 | 5.412098 |
| 2.263947 | 2.572472 | 5.712422 | NA | 3.845419 | NA | 1.556364 | 5.761548 | 6.648494 | 3.272023 |
| 3.237746 | 3.841175 | 3.666143 | 4.693301 | 4.172357 | NA | 2.751707 | 3.306363 | 8.294665 | 3.67996 |
| 6.043373 | 5.219404 | 4.846005 | NA | 4.503016 | NA | 4.052577 | 3.12319 | 8.529266 | 5.427624 |
| 2.62395 | 2.471164 | 3.548741 | NA | 2.337012 | 3.163974 | 2.616189 | 4.305964 | 7.463908 | 2.334016 |
| 3.373456 | 2.807732 | 4.138736 | 4.415083 | 3.97072 | NA | 3.258269 | 4.051549 | 7.760395 | 3.940485 |
| 5.371517 | 3.281022 | 4.169336 | NA | 2.28914 | NA | 3.535843 | 3.428942 | 8.108964 | 5.092835 |
| 1.861754 | 6.09329 | 5.018172 | 5.336976 | 4.39139 | NA | 1.83288 | 5.024068 | 8.087767 | 4.704191 |
| 3.037918 | 3.480537 | 4.594681 | 4.472011 | 4.662283 | NA | 3.70612 | 4.044398 | 8.30301 | 3.888127 |
| 4.453841 | 3.063355 | 4.707349 | NA | 5.795889 | NA | 3.390919 | 4.566566 | 6.695354 | 3.361386 |
| 2.417718 | NA | 3.570355 | NA | NA | NA | NA | NA | NA | NA |
| 0.740255 | 0.741793 | 0.752393 | 0.754054 | 0.760964 | 0.768205 | 0.77282 | 0.784214 | 0.786837 | 0.793102 |
| 3.782899 | 3.074044 | 3.781385 | 4.750178 | 4.185141 | 1.224512 | 2.307757 | 4.055008 | 7.279541 | 3.492296 |

| | | | | | | | | | |
|---|---|---|---|---|---|---|---|---|---|
| 3.633747 | 3.261478 | 3.909291 | 4.593435 | 4.314493 | 1.737324 | 2.429134 | 3.903942 | 7.10109 | 3.586169 |

| Ruxolitinib | Dasatinib | AMG-706 | 17-AAG | Genentech | TG101348 | TAK-715 | NVP-BEZ23 | AICAR | Doxorubici |
|---|---|---|---|---|---|---|---|---|---|
| NA | NA | 3.042874 | -0.69282 | NA | NA | NA | -0.13898 | 8.844391 | NA |
| 4.156765 | 2.065998 | NA | NA | 0.74058 | 1.577777 | 3.469136 | NA | NA | -2.70972 |
| 3.492908 | 1.093992 | 2.358481 | -1.49997 | 0.322881 | 1.84472 | 2.78577 | -3.03883 | 6.712822 | -4.637 |
| 3.870857 | 2.577811 | 3.481203 | -2.31014 | 4.102536 | 1.857313 | 4.683752 | -3.1555 | 5.49695 | -2.75937 |
| 4.878211 | 3.368934 | 4.002534 | -1.55497 | 2.702504 | 4.540716 | 2.778277 | -0.8976 | 9.027364 | -1.80159 |
| 4.683448 | NA | 2.233271 | -0.14064 | 2.650736 | 3.173703 | 5.208619 | -1.75625 | 6.686801 | -0.01839 |
| 1.737125 | NA | 3.475066 | 0.84498 | 2.004868 | 3.125034 | 2.849556 | -2.80214 | 5.245158 | -1.16334 |
| 6.370024 | NA | 5.239431 | 0.356745 | 6.398223 | 6.75041 | 6.748859 | -1.56655 | 11.42798 | 2.159909 |
| 3.770163 | 0.612499 | 1.347216 | -3.14967 | 1.980013 | 2.255211 | 3.788854 | -3.94539 | 7.385097 | -0.24212 |
| 5.044947 | NA | 0.567118 | -1.60678 | 3.549596 | 4.235615 | 4.508573 | -3.12873 | 7.902158 | -0.12649 |
| 4.915884 | NA | 3.545683 | 2.547974 | 1.603634 | 2.414425 | 5.322748 | -2.59201 | 6.356385 | -0.65685 |
| 4.771254 | NA | NA | NA | 2.327923 | 4.18159 | 4.027173 | NA | NA | -1.88605 |
| 5.110702 | NA | 2.621117 | -0.69256 | 5.870966 | 5.87356 | 6.146787 | -2.2443 | 6.733672 | 1.467339 |
| 4.243979 | NA | NA | NA | 1.163952 | 2.155304 | 5.385805 | NA | 6.14438 | -0.75743 |
| 4.348434 | NA | NA | NA | 4.305281 | 3.30877 | 4.785963 | NA | NA | 1.249172 |
| 5.040616 | NA | 3.754268 | -0.82145 | 3.14313 | 4.132332 | 4.763337 | -0.86384 | 10.09475 | -0.37439 |
| 4.250756 | NA | 2.505685 | -3.52288 | 4.659639 | 3.39673 | 5.425187 | -2.72139 | 7.272557 | -1.04064 |
| 4.388529 | NA | 3.562689 | -1.17741 | 3.05105 | 0.909216 | 4.226195 | -1.49514 | 9.226187 | -3.10796 |
| 4.036688 | 2.607668 | NA | NA | 4.706926 | 2.744714 | 3.713131 | NA | NA | -3.66393 |
| 3.813217 | NA | 2.066931 | -3.18784 | 5.025787 | 5.025787 | 4.350795 | -3.04651 | 7.117109 | -4.61416 |
| 4.363225 | NA | 0.757001 | -2.23975 | 1.953714 | 4.545179 | 4.479637 | -2.78995 | 5.744526 | -2.0144 |
| 5.048796 | NA | 3.163943 | 0.805653 | 0.654839 | NA | 3.742945 | -1.63304 | 6.545857 | -0.35877 |
| 4.64228 | 3.032558 | 3.686576 | -3.32052 | 3.212415 | 3.453834 | 3.374528 | -1.54633 | 9.1791 | -0.35397 |
| 4.893044 | NA | 3.961929 | -0.86709 | 2.490732 | 3.321816 | 4.167677 | -2.22733 | 7.45244 | 0.098678 |
| 3.713302 | NA | 2.813544 | -3.70808 | 4.271332 | 2.862477 | 4.041953 | -3.70533 | 8.003472 | -3.85896 |
| 4.62652 | NA | 3.267463 | 0.157628 | 1.855317 | 3.739577 | 3.113664 | -0.89263 | 7.44694 | -1.00439 |
| 3.974559 | NA | 2.281679 | -2.00694 | 2.414317 | 2.279453 | 3.536747 | -2.24137 | NA | -2.42945 |
| 4.346308 | NA | 4.02362 | -0.14693 | 1.984319 | 1.5139 | 4.235209 | -1.71631 | 7.086926 | -2.20292 |
| 3.393942 | NA | 2.228723 | 1.854018 | 0.85601 | 2.973198 | 3.136524 | -2.40426 | 7.688379 | -2.55378 |
| 4.594451 | NA | 3.57672 | -0.32102 | 4.102572 | 4.434858 | 5.918941 | -1.52864 | 7.224606 | -0.92408 |
| 4.335495 | NA | 1.824882 | -2.03904 | 2.610475 | 2.467448 | 5.721273 | -3.08552 | NA | -0.56808 |
| 4.239109 | NA | 3.140235 | -1.16285 | 1.255102 | 2.677606 | 3.294152 | -0.23281 | 8.236142 | 0.924243 |
| 4.239378 | NA | NA | NA | 4.044023 | 2.811361 | 4.400231 | NA | 6.479598 | -0.51189 |
| 4.545511 | NA | 1.854794 | -2.50776 | 5.218975 | 3.770245 | 5.804281 | -3.65759 | 9.11021 | 0.74887 |
| 4.834002 | NA | 4.040403 | -1.70795 | 1.85609 | 1.980963 | 5.52161 | -1.64113 | 7.849978 | 1.961412 |
| 4.010384 | NA | 3.630403 | -2.59821 | 3.857246 | 5.221392 | 4.213611 | -2.72106 | 10.47609 | -3.36534 |
| 4.96353 | NA | 3.380492 | -1.89469 | 5.63351 | 5.63351 | 5.795632 | -2.42916 | 8.889763 | 0.447101 |
| 4.122487 | 0.870634 | 3.065631 | 2.244729 | 0.306483 | 1.884913 | 3.326798 | -1.36907 | 7.110512 | -0.37073 |
| 4.441179 | NA | 2.944958 | -2.8811 | 3.652924 | 2.797964 | 4.513919 | -4.05299 | 6.640379 | -1.55072 |
| 4.035931 | NA | 2.538937 | -0.5065 | 4.927853 | 1.68866 | 4.827939 | -3.26506 | 6.599183 | -0.20114 |
| 4.542293 | NA | 2.065277 | 0.718551 | 4.568514 | 3.192412 | 5.43264 | -1.86911 | 9.4234 | 1.087604 |
| 4.7138 | NA | 1.695116 | -1.57464 | 2.785127 | 2.693406 | 4.940371 | -1.48942 | 7.793849 | -1.99572 |
| 4.322205 | NA | 3.483187 | 2.330592 | 3.178081 | 2.464213 | 5.577659 | -2.42654 | 7.280164 | 0.214758 |
| NA | NA | 3.81883 | -1.85895 | NA | NA | NA | -1.91618 | 8.864872 | NA |
| 0.795646 | 0.797176 | 0.797838 | 0.798774 | 0.813687 | 0.814597 | 0.82122 | 0.823809 | 0.828188 | 0.830326 |
| 4.416153 | 1.943847 | 2.981188 | -0.95854 | 2.975148 | 3.278378 | 4.5238 | -2.16762 | 7.63711 | -0.965 |

| | | | | | | | | | |
|---|---|---|---|---|---|---|---|---|---|
| 4.351666 | 2.170286 | 2.88797 | -1.10078 | 3.096907 | 3.17412 | 4.447275 | -2.24531 | 7.749717 | -1.08239 |

| T0901317 | KIN001-13! | BI-2536 | SB-505124 | S-Trityl-L-c | Crizotinib | VX-680 | KIN001-26( | NG-25 | IOX2 |
|---|---|---|---|---|---|---|---|---|---|
| NA | NA | NA | 6.210492 | NA | NA | NA | NA | NA | 0.837354 |
| 3.877567 | 4.318811 | -3.00343 | NA | 0.397173 | 1.50137 | -0.77524 | 3.031617 | 2.46731 | NA |
| 4.080408 | 3.505941 | -2.25839 | 3.551734 | 0.011126 | 2.299579 | -0.18595 | 2.671271 | 2.080918 | -0.46497 |
| 3.947928 | 2.796554 | -0.19138 | 4.388215 | 2.123669 | 3.49112 | 2.52357 | 4.981075 | 4.630537 | 0.332729 |
| 3.773281 | 4.061593 | -0.57864 | 4.531512 | 1.453089 | 3.430817 | 3.339792 | 3.222526 | 2.633152 | 0.845417 |
| 4.949612 | NA | NA | 3.739866 | NA | NA | NA | 3.340558 | 5.115586 | -0.31847 |
| 3.966248 | 3.745624 | NA | 3.81957 | NA | 2.734849 | NA | 3.080497 | 3.416227 | 0.413508 |
| 6.460971 | NA | NA | 6.303439 | NA | NA | NA | 7.373833 | 5.992365 | 1.081498 |
| 3.504784 | 3.723279 | -1.79695 | 2.076782 | 0.603888 | 2.631489 | 2.082314 | 2.541251 | 0.73489 | -0.58079 |
| 5.96027 | NA | NA | 5.831973 | NA | NA | NA | 6.355309 | 4.751695 | 0.166672 |
| 4.87641 | NA | NA | 3.217225 | NA | NA | NA | 3.294206 | 2.454418 | 1.180459 |
| 4.764517 | NA | NA | 5.367025 | NA | NA | NA | 3.198355 | 3.611499 | -0.26392 |
| 4.996348 | NA | NA | 3.093944 | NA | NA | NA | 5.990943 | 5.64462 | -2.3375 |
| 5.991904 | NA | NA | 4.25745 | NA | NA | NA | 2.608791 | 1.674064 | 1.142796 |
| 4.922995 | 3.757308 | NA | NA | NA | 3.258954 | NA | 3.858381 | 2.20592 | NA |
| 5.344397 | NA | NA | 5.633429 | NA | NA | NA | 3.341443 | 2.365726 | 0.933743 |
| 4.144481 | NA | NA | 4.385598 | NA | NA | NA | 4.258131 | 3.614848 | 0.262359 |
| 5.005356 | NA | NA | 2.716691 | NA | NA | NA | 2.82399 | 2.242116 | 0.117506 |
| 4.029709 | 3.72874 | -2.17766 | NA | 0.66045 | 2.634566 | 1.77432 | 2.423276 | 3.430778 | NA |
| 4.947826 | NA | NA | 4.573303 | NA | NA | NA | 5.327426 | 4.549326 | 0.276654 |
| 4.568248 | NA | NA | 4.99592 | NA | NA | NA | 2.986423 | 2.882705 | -0.19219 |
| NA | NA | NA | 4.676264 | NA | NA | NA | 3.474304 | 2.032423 | 0.972894 |
| 5.164507 | 4.147424 | -0.67029 | 3.193834 | 2.771476 | 3.61805 | 3.150168 | 3.041838 | 2.790786 | -0.21277 |
| 5.308507 | NA | NA | 5.415348 | NA | NA | NA | 5.262457 | 2.908526 | 0.806538 |
| 3.498371 | NA | NA | 2.923443 | NA | NA | NA | 4.387924 | 2.843374 | -1.06929 |
| 5.040191 | NA | NA | 4.591907 | NA | NA | NA | 4.418658 | 5.378887 | 0.292999 |
| 3.753374 | NA | NA | 4.555963 | NA | NA | NA | 2.583761 | 3.012949 | -0.05831 |
| 5.654859 | NA | NA | 3.753626 | NA | NA | NA | 5.216967 | 4.159663 | 0.845592 |
| 3.897102 | NA | NA | 4.663493 | NA | NA | NA | 2.866057 | 3.215987 | 0.058046 |
| 5.339087 | NA | NA | 4.382918 | NA | NA | NA | 3.081551 | 5.271987 | 0.734037 |
| 4.833441 | NA | NA | 4.821685 | NA | NA | NA | 4.434105 | 4.026682 | -0.7038 |
| 3.756211 | NA | NA | 4.189064 | NA | NA | NA | 3.28883 | 2.459416 | 0.495293 |
| 4.868822 | NA | NA | 3.745679 | NA | NA | NA | 1.947214 | 1.683087 | -0.31652 |
| 5.218975 | NA | NA | 4.121788 | NA | NA | NA | 5.754562 | 4.946356 | -0.18535 |
| 5.797881 | NA | NA | 5.165894 | NA | NA | NA | 3.549546 | 2.41195 | 0.776063 |
| 5.264998 | NA | NA | 3.594791 | NA | NA | NA | 5.521311 | 3.798698 | 0.507529 |
| 5.40857 | NA | NA | 3.992766 | NA | NA | NA | 3.921764 | 3.554305 | 0.840667 |
| 3.800141 | 3.413704 | -2.30954 | 3.413822 | -0.11729 | 1.68148 | -0.09458 | 4.331873 | 2.633202 | 0.077844 |
| 4.588915 | NA | NA | 3.496922 | NA | NA | NA | 3.320192 | 2.569789 | -0.08431 |
| 5.483417 | NA | NA | 5.02234 | NA | NA | NA | 3.477402 | 2.214306 | 0.731928 |
| 5.048869 | NA | NA | 4.703808 | NA | NA | NA | 3.117976 | 2.111105 | 0.31819 |
| 5.315549 | NA | NA | 5.227128 | NA | NA | NA | 2.662542 | 4.341636 | 0.42183 |
| 4.451482 | NA | NA | 4.981335 | NA | NA | NA | 4.959171 | 4.264245 | 0.25765 |
| NA | NA | NA | 5.499229 | NA | NA | NA | NA | NA | 0.617747 |
| 0.83058 | 0.832886 | 0.843776 | 0.847495 | 0.85638 | 0.859863 | 0.870024 | 0.872833 | 0.874203 | 0.875597 |
| 4.739263 | 3.701302 | -1.56576 | 4.320309 | 0.917789 | 2.764025 | 1.396896 | 3.880716 | 3.272699 | 0.209275 |

| | | | | | | | | | |
|---|---|---|---|---|---|---|---|---|---|
| 4.793294 | 3.763289 | -1.71916 | 4.388091 | 1.104878 | 2.644699 | 1.609971 | 3.814285 | 3.339687 | 0.248358 |

| CUDC-101 | THZ-2-102- | KIN001-23( | XMD8-85 | PHA-79388 | Paclitaxel | AT-7519 | WZ-1-84 | A-443654 | CAL-101 |
|---|---|---|---|---|---|---|---|---|---|
| NA | NA | NA | NA | NA | NA | NA | NA | NA | NA |
| -1.98332 | -4.6611 | 3.70977 | 2.1667 | -0.17865 | -3.71086 | -1.45534 | 4.272731 | -1.80198 | 3.636004 |
| -1.62807 | -3.84322 | 1.970969 | 1.968148 | 0.281275 | -3.28914 | -1.17964 | 2.982 | -1.14793 | 4.627047 |
| 1.146509 | -0.21571 | 5.097952 | 2.576967 | 0.838846 | -2.30929 | 0.591528 | 3.736908 | -0.40677 | 5.6585 |
| -0.12856 | 1.650483 | 4.623281 | 4.865326 | 0.63857 | -3.03167 | -0.75152 | 2.915952 | -1.36386 | 4.554617 |
| -0.96447 | 1.523697 | 4.793952 | NA | 4.339685 | NA | 3.864877 | NA | NA | 5.331047 |
| -0.60677 | -3.7251 | 4.837872 | NA | 2.837928 | NA | -0.50674 | NA | NA | 5.151389 |
| 6.241743 | 1.438521 | 6.082187 | NA | 3.65151 | NA | 3.811822 | NA | NA | 5.3792 |
| -1.80365 | -2.65961 | 3.188591 | 2.50571 | 2.061036 | -3.5202 | 0.086521 | 3.351076 | -1.48606 | 2.789295 |
| 0.586051 | -0.30114 | 4.458671 | NA | 4.225734 | NA | 1.957556 | NA | NA | 5.744791 |
| -1.87024 | -1.7842 | 5.250875 | NA | 1.816496 | NA | 2.059761 | NA | NA | 2.442384 |
| -0.53838 | -2.73099 | 5.143582 | NA | 3.212265 | NA | 2.484548 | NA | NA | 5.315742 |
| 2.409749 | -1.50554 | 5.681041 | NA | 5.859442 | NA | 3.726413 | NA | NA | 4.42948 |
| -1.60425 | -4.7506 | 4.880431 | NA | 5.161846 | NA | 1.686588 | NA | NA | 4.996884 |
| 2.384753 | 3.663537 | 5.081012 | 4.831334 | 5.001499 | NA | 2.983026 | NA | NA | 4.196215 |
| 1.20715 | 0.831275 | 3.95165 | NA | 3.440303 | NA | 1.595702 | NA | NA | 6.499153 |
| -1.57441 | 1.057146 | 4.257654 | NA | 4.220698 | NA | 4.041995 | NA | NA | 4.985353 |
| -0.85281 | -2.36085 | 5.043774 | NA | 2.210171 | NA | 1.473544 | NA | NA | 4.436019 |
| -1.7585 | NA | 4.340733 | 3.192977 | 3.289196 | -3.24952 | 2.733509 | 4.295515 | -2.83555 | 5.192255 |
| 2.152213 | 3.016145 | 5.025787 | NA | 5.025787 | NA | 4.415688 | NA | NA | 5.687864 |
| -1.42083 | -3.7093 | 3.769136 | NA | 2.703554 | NA | -0.15966 | NA | NA | 5.169696 |
| -1.52628 | -0.45991 | 3.828461 | NA | NA | NA | -0.67045 | NA | NA | 2.900432 |
| 0.567281 | -1.42166 | 4.34981 | 4.339307 | 2.419239 | -1.86303 | 1.207616 | 2.515443 | -0.53924 | 4.069912 |
| 0.252424 | -1.36797 | 5.586055 | NA | 2.923298 | NA | 0.212099 | NA | NA | 6.000125 |
| 2.283707 | 3.036537 | 4.449548 | NA | 3.498695 | NA | 3.399148 | NA | NA | 4.716121 |
| -0.16961 | 3.280488 | 4.43433 | NA | 1.045993 | NA | -1.28585 | NA | NA | 4.911703 |
| -0.87339 | -3.85556 | 3.72014 | NA | -0.11869 | NA | -0.04216 | NA | NA | 5.320532 |
| -0.18999 | -0.97955 | 4.440287 | NA | 2.913415 | NA | 0.271671 | NA | NA | 3.379774 |
| 0.236623 | -3.80951 | 4.034549 | NA | 0.025923 | NA | 0.336323 | NA | NA | 2.757618 |
| -0.60779 | 0.766898 | 5.492175 | NA | 3.774692 | NA | 3.03351 | NA | NA | 5.506811 |
| -0.36927 | -2.82512 | 5.044448 | NA | 4.074012 | NA | 2.559784 | NA | NA | 5.48291 |
| -2.845 | 1.266304 | 3.849538 | NA | 1.662637 | NA | 1.193524 | NA | NA | 4.675324 |
| -0.68321 | -5.50399 | 4.911226 | NA | -0.95459 | NA | -1.35744 | NA | NA | 3.65486 |
| 0.411307 | -1.65748 | 5.216755 | NA | 5.218975 | NA | 5.20973 | NA | NA | 5.861397 |
| -1.2354 | -3.40686 | 5.695949 | NA | 2.654869 | NA | 0.975429 | NA | NA | 4.432598 |
| 0.057652 | -2.20209 | 3.745702 | NA | 5.12703 | NA | 2.376495 | NA | NA | 5.604367 |
| 0.944092 | -2.43107 | 5.440738 | NA | 4.841573 | NA | 4.480404 | NA | NA | 5.710773 |
| 0.277956 | -2.92459 | 4.43526 | 1.544326 | 0.045198 | -4.18014 | -1.3041 | 3.396886 | -0.13079 | 3.701149 |
| 0.874477 | 0.622752 | 4.514177 | NA | 3.382854 | NA | 2.789341 | NA | NA | 4.925331 |
| 0.298186 | -4.08781 | 4.448596 | NA | 4.180406 | NA | 3.979823 | NA | NA | 3.853685 |
| 2.2928 | -0.70463 | 5.299269 | NA | 4.504188 | NA | 3.350579 | NA | NA | 5.380704 |
| -0.28429 | -2.83244 | 4.543182 | NA | 2.190901 | NA | 1.407277 | NA | NA | 4.356412 |
| -0.25791 | 3.384665 | 5.032702 | NA | 4.262064 | NA | 1.133582 | NA | NA | 5.234808 |
| NA | NA | NA | NA | NA | NA | NA | NA | NA | NA |
| 0.880701 | 0.883114 | 0.89785 | 0.903118 | 0.921829 | 0.924624 | 0.929303 | 0.937041 | 0.940261 | 0.940812 |
| 0.024767 | -1.08079 | 4.591368 | 3.152364 | 2.918744 | -3.17223 | 1.557096 | 3.451733 | -1.24132 | 4.716066 |

| | | | | | | | | | |
|---|---|---|---|---|---|---|---|---|---|
| -0.06291 | -1.20024 | 4.625942 | 3.025537 | 2.862134 | -3.09757 | 1.609834 | 3.402615 | -1.16853 | 4.739486 |

| MS-275 | 5-Fluorour | YM201636 | VX-11e | Tamoxifen | GW843682 | KU-55933 |
|---|---|---|---|---|---|---|
| NA | NA | NA | NA | 5.517345 | NA | 5.784574 |
| -0.58813 | 3.230813 | 1.263637 | 3.122882 | NA | -3.88399 | NA |
| -0.44861 | 1.445262 | 1.305973 | -2.39865 | 3.142001 | -3.04946 | 3.53101 |
| 0.39849 | 2.687069 | 4.431127 | 4.431127 | 4.232997 | 0.156356 | 2.286584 |
| -0.09488 | 2.472024 | 1.79643 | 4.217888 | 3.671786 | -1.06692 | 4.983052 |
| NA | 5.789608 | 4.111627 | 4.598998 | 3.143101 | NA | 4.157245 |
| 2.469508 | 4.318062 | 3.69797 | 4.051752 | 3.284496 | NA | 4.034008 |
| NA | 2.86835 | 2.905473 | 5.977022 | 6.010764 | NA | 5.375938 |
| 0.294068 | 3.446957 | 1.267866 | 2.371256 | 2.762005 | -1.75004 | 2.291018 |
| NA | 6.318133 | 2.680829 | 5.138925 | 5.138826 | NA | 5.779971 |
| NA | 3.170151 | 1.625427 | 3.949949 | 0.989673 | NA | 4.51455 |
| NA | 4.17852 | 2.825129 | 4.073324 | 4.682365 | NA | NA |
| NA | 6.872441 | 4.171945 | 5.180822 | 3.814127 | NA | 3.577733 |
| NA | 4.899689 | 1.050895 | 5.092605 | 3.405997 | NA | NA |
| 4.279269 | 4.722323 | 2.741446 | 4.333219 | NA | NA | NA |
| NA | 4.456134 | 2.331987 | 3.101301 | 4.881859 | NA | 4.494996 |
| NA | 5.48657 | 1.305967 | 3.73276 | 2.526963 | NA | 4.147221 |
| NA | 2.189233 | 1.87854 | 4.063854 | 4.290192 | NA | 3.854204 |
| 0.269085 | -0.03334 | 1.972173 | 3.91339 | NA | -2.76998 | NA |
| NA | 6.165221 | 4.33264 | 4.33264 | 1.95347 | NA | 2.99346 |
| NA | 4.687255 | 1.160961 | 4.380709 | 4.28756 | NA | 3.364089 |
| NA | 4.308261 | 0.704161 | 3.611555 | 4.941355 | NA | 5.718079 |
| 0.556679 | 3.677685 | 1.751956 | 3.014921 | 3.681607 | -0.01293 | 5.302693 |
| NA | 5.330484 | 2.885133 | 4.891654 | 4.708914 | NA | 5.496251 |
| NA | 3.747422 | 3.756401 | 3.756401 | 3.659164 | NA | 2.706084 |
| NA | 4.798478 | 3.853334 | 4.692914 | 3.129231 | NA | 5.329521 |
| NA | 1.579634 | 2.451254 | 1.966738 | 3.819746 | NA | 4.53498 |
| NA | 5.15441 | 1.923532 | 4.680014 | 3.080212 | NA | 3.337899 |
| NA | 3.009851 | 2.330554 | 2.970942 | 3.970346 | NA | 2.809349 |
| NA | 2.463229 | 3.0298 | 4.77021 | 4.403192 | NA | 4.857135 |
| NA | 3.787246 | 2.577845 | 4.131678 | 4.241445 | NA | 3.797854 |
| NA | 1.898559 | 1.061551 | 3.912365 | 4.41588 | NA | 4.797944 |
| NA | 1.289946 | 2.484973 | 2.173967 | 4.092118 | NA | NA |
| NA | 6.346772 | 3.461603 | 4.513653 | 4.35494 | NA | 2.639337 |
| NA | 6.913547 | 2.553338 | 1.432403 | 4.628307 | NA | 4.155952 |
| NA | 4.479224 | 1.121821 | 3.438532 | 3.342459 | NA | 4.114792 |
| NA | 3.959331 | 2.566549 | 4.905852 | 4.734617 | NA | 4.601226 |
| 1.713433 | 4.497567 | 2.063947 | 3.473847 | 1.96404 | -3.01707 | 3.435883 |
| NA | 4.326438 | 3.031186 | 3.323847 | 3.103332 | NA | 4.160861 |
| NA | 3.580095 | 2.386597 | 4.128718 | 3.619871 | NA | 4.576612 |
| NA | 5.883768 | 2.209145 | 4.499301 | 4.629835 | NA | 4.333134 |
| NA | 4.569461 | 1.031991 | 4.666768 | 4.533981 | NA | 4.88549 |
| NA | 4.791823 | 4.422052 | 4.345528 | 3.83439 | NA | 4.729443 |
| NA | NA | NA | NA | 3.093078 | NA | NA |
| 0.947702 | 0.974393 | 0.976488 | 0.981433 | 0.987697 | 0.990895 | 0.994282 |
| 0.901388 | 4.032432 | 2.434839 | 3.825825 | 3.843406 | -1.91881 | 4.200865 |

| | | | | | | |
|---|---|---|---|---|---|---|
| 0.846399 | 4.048495 | 2.44498 | 3.837142 | 3.848923 | -1.93333 | 4.203394 |

| Cell Line | E2F2_Activity | **BMN-673** | **CHIR-9902** | **Olaparib (r** | **Bleomycin** | **AG-01469‹** | **AS601245** | **AZD7762** |
|---|---|---|---|---|---|---|---|---|
| ZR-75-30 | 0.1726741 | 4.831413 | 5.778228 | 6.210492 | 7.017756 | 5.7039 | NA | 2.933205 |
| HCC1599 | 0.1958902 | NA | NA | NA | 1.878536 | NA | 2.196232 | NA |
| DU-4475 | 0.2215439 | 3.482276 | 3.170432 | NA | 2.756884 | 2.377256 | 1.766916 | -0.63141 |
| HCC2157 | 0.2293765 | 4.984005 | 4.616089 | 4.727899 | 4.325277 | 4.084896 | 2.114977 | -1.05479 |
| HCC2218 | 0.2329256 | 4.670262 | 4.710469 | 5.207943 | 5.927842 | 4.706033 | 3.635666 | 3.684096 |
| HCC1428 | 0.2469709 | 3.686001 | 4.629534 | 4.351636 | 5.377754 | 4.489309 | 2.667916 | 2.058364 |
| HCC1500 | 0.2481241 | 5.018679 | 5.018679 | 5.018679 | 4.823976 | 4.435623 | 2.30015 | 2.047044 |
| BT-483 | 0.2722625 | 2.731514 | 5.131225 | 6.734945 | 6.274419 | 5.731183 | 6.079238 | 3.818782 |
| EVSA-T | 0.3215148 | 2.232709 | 4.080782 | 2.882333 | 1.464536 | 3.685837 | 2.761081 | 0.661062 |
| HCC1419 | 0.3479547 | 5.288988 | 4.903418 | 5.831973 | 7.094178 | 5.305562 | 3.305839 | 3.967624 |
| T47D | 0.3554464 | 5.782737 | 5.785629 | 5.240223 | 4.551292 | 3.033465 | 3.222415 | 4.40608 |
| CAMA-1 | 0.390238 | 5.470893 | 5.170566 | 3.859481 | 6.161734 | 4.429987 | 3.329246 | NA |
| MDA-MB-4 | 0.4064518 | 4.212353 | 4.61121 | 4.670301 | 6.221267 | NA | 2.631692 | 1.950864 |
| EFM-19 | 0.4245251 | 4.102543 | 5.747966 | 3.750004 | 4.780249 | 4.575343 | 2.222148 | 0.939586 |
| UACC-812 | 0.448281 | NA | NA | NA | NA | NA | 4.695599 | NA |
| UACC-893 | 0.4638305 | 4.635757 | 5.628592 | 2.569415 | 2.986994 | 4.622383 | 4.012544 | 4.016693 |
| MDA-MB-3 | 0.4881617 | 4.868625 | 4.629582 | 4.173797 | 5.304552 | 4.274598 | 1.421386 | -0.52255 |
| HCC38 | 0.495164 | 2.195339 | 3.449611 | 4.276386 | 2.142495 | 3.958404 | 1.248765 | -0.22068 |
| CAL-148 | 0.5070106 | NA | NA | NA | NA | NA | 4.346315 | NA |
| MCF7 | 0.5080066 | 1.110639 | 2.17295 | 3.252239 | 2.138198 | 1.907525 | 0.222062 | 1.620276 |
| AU565 | 0.5171166 | 1.372234 | NA | NA | 4.926217 | 2.38322 | 0.784133 | -0.52949 |
| HCC202 | 0.5425118 | 3.28477 | 5.447247 | 4.425001 | 5.610512 | 5.155564 | 2.41998 | 2.048066 |
| BT-474 | 0.5427423 | 4.035084 | 4.379491 | 3.851281 | 5.65449 | 2.769537 | 2.254004 | 2.73895 |
| BT-20 | 0.5678861 | 3.065859 | 4.468621 | 4.550445 | 2.543184 | 3.66445 | 3.688836 | 0.937712 |
| CAL-51 | 0.5767389 | -2.25266 | 3.24197 | 1.353893 | -1.58008 | 1.000823 | 1.041737 | -0.58322 |
| HCC1954 | 0.5794048 | 3.933652 | 4.129473 | 4.352374 | 1.67788 | 3.908693 | 1.821509 | 1.056772 |
| HCC1806 | 0.5897348 | NA | 3.861043 | 2.222954 | -1.20819 | 2.819255 | 0.217407 | NA |
| HCC1569 | 0.6107715 | 1.3908 | 3.642967 | 3.282759 | 4.254627 | 2.775353 | 0.675198 | 1.005155 |
| MDA-MB-4 | 0.6144312 | NA | 4.663064 | 2.356764 | 1.379961 | 4.01002 | 2.581057 | -0.25101 |
| HDQ-P1 | 0.6155078 | 4.927595 | 3.345958 | 4.057441 | 3.251652 | 4.747196 | 2.062223 | 0.703034 |
| MDA-MB-1 | 0.6234777 | 2.620001 | 3.495571 | 4.076912 | 3.507223 | 2.970848 | 1.289365 | NA |
| MDA-MB-4 | 0.6311505 | 3.053612 | 4.839225 | 3.954624 | 4.836065 | NA | 3.032996 | 0.656869 |
| HCC70 | 0.6535518 | 3.851926 | 4.373748 | 4.273914 | 4.045316 | 3.939267 | 1.718007 | -0.89108 |
| HS-578-T | 0.6792037 | 2.806696 | 2.465456 | 4.121294 | 2.319519 | 3.87678 | 2.086645 | -0.68102 |
| HCC1143 | 0.687603 | 3.379658 | 5.592451 | 5.301073 | 3.545506 | 4.57136 | 1.652607 | -1.28675 |
| EFM-192A | 0.6900504 | 4.310257 | 3.485003 | 4.69476 | 6.57658 | 4.546893 | -0.52607 | 2.378049 |
| MDA-MB-4 | 0.6952879 | 1.522867 | 4.925697 | 1.881495 | 2.175016 | 4.062379 | 2.751514 | 0.266813 |
| HCC1187 | 0.6972928 | 3.904177 | 1.72901 | 2.655109 | 0.829458 | 2.419654 | 0.720429 | -1.73671 |
| JIMT-1 | 0.7101335 | 1.130782 | 3.771768 | 3.236867 | 2.982429 | 4.277253 | 1.322525 | -0.92191 |
| CAL-120 | 0.7121776 | 2.006479 | 3.791532 | 2.152588 | 1.949369 | 3.548974 | 3.435589 | 1.506538 |
| HCC1395 | 0.7151043 | 0.849773 | 3.265475 | 3.79661 | 1.950418 | 4.494484 | 2.794891 | -1.32203 |
| HCC1937 | 0.7724891 | 3.428751 | 3.29749 | 3.955583 | 3.280989 | 3.429646 | 1.561914 | -1.1185 |
| MDA-MB-2 | 0.7794799 | 1.585491 | 3.961346 | 2.807325 | 3.400769 | 2.318364 | 2.113155 | 0.740676 |
| BT-549 | 0.7988771 | 4.257894 | 3.302338 | 3.494637 | 2.215068 | 3.743025 | NA | -0.81413 |
| | **Significance** | 0.000473 | 0.000803 | 0.004025 | 0.004676 | 0.007299 | 0.007331 | 0.008144 |

| CHIR-9902 | JNK-9L | PD-173074 | piperlongu | YK 4-279 | AP-24534 | PI-103 | Cetuximab | OSU-03012 | BX-795 |
|---|---|---|---|---|---|---|---|---|---|
| NA | NA | 4.787609 | 3.703994 | NA | NA | NA | NA | NA | 5.7039 |
| 3.44923 | -0.938 | NA | NA | NA | 1.821199 | -0.95642 | NA | 1.899682 | NA |
| 2.838778 | -0.55346 | 2.521195 | 1.688269 | 2.111051 | -0.77018 | 0.884285 | 5.96462 | 1.215319 | 0.681465 |
| 3.73798 | -0.5311 | 2.172291 | 1.326911 | 4.451779 | 2.126142 | -2.14396 | 6.86804 | 2.777007 | 1.918954 |
| 3.388665 | 0.376655 | 3.41178 | 1.8891 | 3.921407 | 2.312513 | -0.31806 | 7.334622 | 2.722441 | 4.801432 |
| 3.58686 | 0.181863 | 3.75099 | 2.474611 | 2.50065 | 1.193737 | 2.22469 | 6.686492 | 2.827933 | 3.61804 |
| 3.526322 | 0.299339 | 2.919339 | 1.93084 | 3.101283 | 2.148859 | 0.313379 | 6.902713 | 1.948533 | 3.757579 |
| 5.606226 | 1.895405 | 4.845739 | 3.805655 | 6.227111 | 3.663774 | 0.028027 | 6.695621 | 5.534807 | 4.730662 |
| 3.024469 | 0.327754 | 2.766194 | 1.515154 | 0.848933 | 1.4444 | -1.21686 | 6.229232 | 2.741296 | 1.182538 |
| 4.507715 | 0.031596 | 4.31411 | 4.545497 | 4.120854 | 2.982722 | 1.221674 | 6.969105 | 3.253287 | 3.970879 |
| 4.593205 | 2.162903 | 4.381591 | 1.646189 | 3.019059 | 2.89162 | -0.22888 | 7.526436 | 2.193505 | 4.306391 |
| 4.202409 | 2.736632 | NA | 1.768772 | 1.864271 | 2.355593 | 0.499753 | 7.335912 | 1.456718 | NA |
| 4.425192 | 0.446427 | 3.512774 | 2.23172 | NA | 1.90298 | 4.102549 | 5.448285 | 2.569253 | 5.157172 |
| 1.94289 | -0.3513 | NA | 1.852814 | 1.725506 | 1.984582 | 0.785622 | 7.545996 | 1.840563 | NA |
| 3.694253 | 0.381588 | NA | NA | NA | 2.076867 | -0.04077 | NA | 4.809792 | NA |
| 4.4424 | 2.055885 | 3.332611 | 1.052854 | NA | 2.836663 | 0.400145 | 7.036776 | 4.270798 | 4.783339 |
| 3.491228 | -0.17418 | 3.358308 | 1.133875 | 1.172244 | 1.99573 | 3.361363 | 6.179747 | 2.327666 | 3.504123 |
| 3.711218 | 0.560344 | 2.657742 | 2.303666 | 0.613259 | 0.221772 | -0.02184 | 5.092236 | 2.503469 | 2.161797 |
| 3.319813 | -0.30614 | NA | NA | NA | 1.313892 | 1.90321 | NA | 2.408915 | NA |
| 2.275748 | -1.15627 | 1.992114 | 1.469146 | 1.498999 | -0.08893 | 5.025787 | 6.509872 | 0.166764 | 2.450228 |
| 3.587735 | -0.71806 | 3.528464 | 1.546895 | NA | 1.930314 | 3.156637 | NA | 1.879458 | 3.210517 |
| NA | NA | 3.50545 | 2.003444 | 2.760535 | NA | NA | 6.707138 | 1.762575 | 5.022792 |
| 3.939417 | 0.046684 | 3.60418 | 1.803576 | NA | 2.262848 | -1.0665 | 6.934045 | 1.389069 | 3.881383 |
| 4.061699 | -0.46732 | 3.969455 | 1.212435 | 2.036492 | 2.152389 | 0.011318 | 4.343809 | 2.973814 | 4.157475 |
| 3.03941 | -1.58597 | 2.555578 | -0.01934 | 0.411372 | -0.21384 | 1.818811 | 6.011129 | -1.14196 | 0.277188 |
| 3.951843 | -0.64908 | 2.859551 | 1.248529 | 1.570377 | 1.998639 | 3.61239 | 6.176905 | 2.165298 | 3.03619 |
| 2.498027 | -1.08867 | 2.754734 | 1.249743 | 0.548604 | 0.127155 | 3.404958 | 5.374206 | 1.473585 | 2.328539 |
| 1.064706 | -0.85706 | 1.315301 | 2.786822 | 1.534797 | 2.63963 | -1.68361 | 6.504863 | 1.865003 | 2.985002 |
| 3.39764 | -0.24506 | 2.8531 | 0.899973 | 1.150922 | 1.21062 | 0.560949 | 3.262926 | 2.38835 | 0.482869 |
| 2.763785 | 0.693783 | 3.872027 | 2.241135 | 2.858375 | 0.556506 | 1.671566 | 5.447712 | 1.885759 | 3.381025 |
| 2.738685 | -0.32713 | 2.375876 | 1.508482 | 2.014975 | 0.122561 | 2.714923 | 6.662682 | 2.583569 | 0.440382 |
| 3.785517 | 1.116874 | 2.10435 | 1.587741 | 1.390712 | 2.05369 | -0.85255 | 6.430256 | 2.214842 | 3.785884 |
| 3.524932 | -1.13379 | NA | 1.334229 | 2.991041 | 1.47399 | 0.934097 | 5.621678 | 2.969636 | NA |
| 3.137159 | 2.401047 | 0.804292 | 1.777242 | 1.049827 | 0.579837 | 5.215496 | 5.431279 | 2.877632 | 1.176654 |
| 2.165547 | -0.02957 | 2.292319 | 1.754166 | 1.961077 | 1.813774 | 1.670133 | 7.180265 | 3.203218 | 2.298404 |
| 1.550724 | -2.63471 | 3.036574 | 2.233502 | 1.36467 | 2.276722 | 5.218006 | 5.978362 | 0.138459 | 2.734774 |
| 3.907934 | 0.026148 | 3.990774 | 0.691005 | 1.964103 | 2.596665 | 3.139002 | 6.902495 | 3.159428 | 3.210166 |
| 2.519043 | -0.34014 | 2.278415 | 1.419962 | -0.19666 | 1.773636 | 0.8498 | 6.507622 | 3.851266 | 1.169795 |
| 3.158416 | -0.68402 | 2.666726 | 0.826698 | 0.286883 | 0.26435 | 0.3568 | 5.325417 | 1.829899 | 1.301006 |
| 3.558759 | 0.535509 | 1.65718 | 0.907023 | 1.655311 | -0.70274 | -0.79875 | 6.588385 | 2.575128 | 1.608756 |
| 1.500492 | -0.51175 | 0.995574 | 2.809949 | 1.801346 | -0.47606 | 2.544552 | 6.3332 | 1.810206 | 3.299755 |
| 3.8994 | 0.14384 | 3.759464 | 1.891547 | 2.707559 | 2.390322 | 0.329766 | 6.127254 | 1.551405 | 3.936617 |
| 3.715641 | 0.059729 | 2.594906 | 0.608196 | 1.825202 | 0.964871 | 2.369128 | 5.995082 | 1.1604 | 1.772682 |
| NA | NA | 3.997544 | 0.65667 | 1.35037 | NA | NA | 7.162445 | NA | 2.102697 |
| 0.009225 | 0.013687 | 0.016344 | 0.018518 | 0.028952 | 0.031318 | 0.031927 | 0.032056 | 0.032295 | 0.035333 |

| XAV 939 | PF-562271 | Cisplatin | (5Z)-7-Oxo | Gefitinib | SN-38 | I-BET | Bryostatin | BX-912 | Olaparib |
|---|---|---|---|---|---|---|---|---|---|
| 5.509821 | NA | 5.231838 | 3.453564 | 3.178167 | 0.912175 | NA | NA | NA | 5.7039 |
| NA | 2.199525 | NA | NA | NA | -6.16386 | 2.557942 | -2.30597 | 1.680101 | NA |
| 3.253538 | 0.782466 | 2.516856 | -2.08125 | 0.12721 | -4.55076 | 1.990062 | -4.48863 | 1.36964 | 2.964822 |
| 4.265257 | 1.313927 | 3.2593 | 1.60199 | 0.57964 | -1.26079 | 2.966649 | -2.19357 | 4.673059 | 4.110472 |
| 4.700037 | 3.355113 | 5.053287 | 2.599752 | 1.891081 | -2.03051 | 3.51772 | -1.57484 | 5.618227 | 4.344174 |
| 4.524115 | NA | 5.323404 | 2.554607 | 1.798277 | -2.46545 | 3.582308 | -2.98932 | 4.393342 | 3.926668 |
| 4.03214 | 3.003786 | 5.037177 | 2.181913 | 1.103776 | -3.69554 | 2.420157 | -2.07751 | 5.099389 | 3.223817 |
| 6.086337 | 4.932546 | 6.579369 | 1.831044 | 3.050475 | -0.39304 | 4.350356 | -0.41632 | 7.033593 | 6.336712 |
| 3.011549 | 2.826884 | 2.300026 | 1.033165 | 1.420694 | -4.65128 | 0.968587 | -2.89165 | 3.900131 | 3.723279 |
| 5.13618 | 4.509163 | 2.081667 | 3.328108 | 1.821528 | 0.466268 | 4.864732 | -1.60356 | 5.903836 | 5.288459 |
| 2.622104 | 2.795079 | 4.831335 | 2.078189 | 3.020034 | -2.98958 | 3.787999 | -3.01962 | 4.095058 | 4.8246 |
| 3.877476 | 1.440215 | NA | 2.199292 | NA | -2.14565 | 3.953649 | -2.1003 | 4.241084 | NA |
| 3.893437 | 3.207342 | 4.03393 | 1.896552 | 2.854587 | -2.91778 | 6.537287 | -2.01413 | 5.742345 | 5.157172 |
| 5.020224 | NA | 4.836621 | 2.028932 | 2.85788 | -2.92163 | 3.322145 | -4.41835 | 3.309159 | 4.048679 |
| NA | 3.464285 | NA | NA | NA | NA | 3.655914 | -2.71786 | 5.0797 | NA |
| 3.535975 | 4.295862 | 3.335112 | 1.298917 | 2.516015 | -3.29192 | 5.286219 | -1.72313 | 5.827983 | 4.939071 |
| 3.191741 | 2.372267 | 2.696525 | 1.537264 | 0.124397 | -1.90032 | 2.611188 | -2.88658 | 4.972763 | 4.057999 |
| 3.045332 | 3.237375 | 4.321389 | 2.375457 | 1.553316 | -5.03191 | 3.508861 | -2.3785 | 3.366132 | 4.461596 |
| NA | 2.015123 | NA | NA | NA | NA | 2.753456 | -2.44765 | 4.291339 | NA |
| 3.67738 | 1.137707 | 3.119642 | 1.187821 | 1.02863 | -6.23815 | 5.69418 | -3.90286 | 4.994247 | 3.193926 |
| 4.167042 | 3.05057 | 2.135266 | NA | -0.33061 | -4.75278 | 4.39388 | -2.68825 | 4.530978 | 3.856696 |
| 3.302903 | NA | 3.462991 | 2.737893 | 1.83246 | -1.76723 | 3.076802 | -2.96003 | 3.290842 | 3.656634 |
| 3.715876 | 3.856008 | 5.308559 | 0.913706 | -0.17784 | -1.54471 | 4.661485 | -1.91598 | 4.19899 | 4.47975 |
| 3.85098 | 2.227746 | 3.406734 | 1.131463 | 0.127861 | -3.36746 | 6.040444 | -2.30809 | 4.514487 | 4.873831 |
| 2.265792 | 0.459596 | 2.106029 | -0.26207 | 0.595923 | -6.08969 | 4.549189 | -4.28304 | 3.408025 | 2.378359 |
| 3.963975 | 3.39982 | 4.211371 | 1.50594 | 1.257925 | -3.51418 | 5.97823 | -3.88769 | 5.256772 | 3.877398 |
| 3.763769 | 1.719302 | NA | 1.315145 | NA | -6.24572 | 4.268179 | -3.78104 | 3.369416 | NA |
| 3.320823 | 3.099463 | 4.963899 | 1.750524 | 2.550116 | -2.81545 | 3.673603 | -3.32617 | 2.440822 | 4.947107 |
| 3.45601 | 0.850336 | 1.831789 | 0.227915 | -0.51033 | -5.90304 | 4.375705 | -4.94708 | 0.085619 | 3.539518 |
| 3.434491 | 2.208718 | 3.688831 | 0.439281 | -0.21888 | -2.41156 | 6.144067 | -3.24923 | 4.284119 | 4.792145 |
| 3.962528 | 3.639437 | NA | 1.587809 | NA | -4.44819 | 4.899137 | -3.38254 | 2.795835 | NA |
| 4.41652 | 2.362586 | 4.880537 | 1.886043 | 2.155044 | -4.35585 | 1.244565 | -1.99456 | 2.246323 | 4.497219 |
| 4.095263 | 0.508189 | 2.288884 | 1.57784 | -0.04509 | -2.70774 | 2.481503 | -4.05936 | 1.912853 | 4.080932 |
| 3.844619 | 3.81394 | 2.531229 | 1.063642 | 1.822326 | -3.2724 | 5.737388 | -2.01086 | 3.89586 | 3.840579 |
| 4.259126 | 1.417619 | 3.926546 | 1.382298 | 0.940189 | -2.82331 | 6.153162 | -3.59976 | 4.402599 | 4.545143 |
| 4.418756 | -0.50974 | 4.105288 | 1.529494 | 1.665995 | -1.74368 | 3.111689 | -5.36102 | 3.983597 | 4.458873 |
| 2.666365 | 3.490238 | 1.533546 | 0.53645 | 1.940762 | -5.20307 | 4.844691 | -2.17725 | 5.514689 | 3.694957 |
| 0.766339 | 1.937517 | 2.441117 | 1.543672 | 1.669912 | -3.33733 | 3.128707 | -2.82832 | -0.07339 | 2.148416 |
| 2.298762 | 1.981048 | 2.615458 | 0.545187 | 0.223566 | -2.86477 | 5.570133 | -3.59914 | 4.371779 | 2.945602 |
| 3.296968 | 0.645555 | 3.880586 | 0.055861 | 1.244093 | -4.36203 | 2.660354 | -1.90879 | 2.479855 | 4.544478 |
| 4.026918 | 3.412051 | 2.30756 | 2.059545 | 2.501775 | -3.20869 | 5.42906 | -2.56531 | 4.249172 | 2.905373 |
| 3.385279 | 2.30426 | 3.027183 | 1.963418 | 1.36901 | -3.04026 | 3.779169 | -2.27671 | 3.440672 | 3.566916 |
| 3.065537 | 0.357546 | 2.928112 | -0.22819 | 1.845289 | -5.00218 | 4.987353 | -2.10909 | 4.626004 | 3.3429 |
| 2.973059 | NA | 3.390256 | 0.443387 | 2.712876 | -3.72654 | NA | NA | NA | 4.860366 |
| 0.037209 | 0.03829 | 0.041537 | 0.041559 | 0.044115 | 0.044413 | 0.044888 | 0.044946 | 0.045763 | 0.046044 |

| **GSK-19045** | Midostauri | Vinorelbin | Epothilone | IPA-3 | Trametinib | Pazopanib | BMS-7548 | BAY 61-36 | KIN001-10 |
|---|---|---|---|---|---|---|---|---|---|
| NA | NA | NA | NA | NA | 3.907907 | NA | NA | NA | NA |
| 2.851873 | -0.14243 | -4.99671 | -4.73721 | 4.212156 | NA | 3.932494 | 0.647959 | 1.148779 | 0.456112 |
| 1.672763 | -0.89089 | -3.15388 | -6.12673 | 3.070265 | -6.58334 | 3.421375 | -0.01638 | 0.74988 | 1.915194 |
| 0.607605 | -0.24411 | -3.88879 | -6.44712 | 4.391752 | 2.681126 | 4.543178 | -1.14051 | 2.240086 | 2.58928 |
| 3.178425 | 2.403972 | -3.20976 | -4.26745 | 5.309546 | 3.148003 | 2.51542 | 3.562362 | 2.563088 | 0.278426 |
| 1.794169 | 0.630848 | -1.73096 | -3.11524 | 6.483796 | 2.4843 | 2.680371 | 2.787622 | 5.132913 | 2.602823 |
| 2.740959 | 1.609742 | -3.84247 | -1.81519 | 4.754986 | 1.085616 | 3.166231 | 2.307829 | 3.821011 | 2.644524 |
| 4.643651 | 2.452228 | 0.294838 | 0.64183 | 7.200604 | 4.427726 | 6.414244 | 4.805036 | 6.050956 | 2.688934 |
| 2.08341 | 1.249444 | -2.50253 | -3.80399 | 4.978838 | 1.468879 | 3.850818 | 2.788086 | 3.496604 | 0.403005 |
| 3.71926 | 2.915077 | -2.49072 | -2.45743 | 4.56018 | 2.474246 | 2.729854 | 4.526719 | 4.905304 | 1.326838 |
| 3.114878 | 1.480253 | -1.51449 | -2.07905 | 3.732737 | 1.987017 | 4.315208 | 1.132843 | 3.560336 | -0.37954 |
| 1.964253 | 1.413943 | -4.41947 | -4.0142 | 4.524951 | 2.540606 | 5.407815 | 2.752889 | 4.245394 | 2.484314 |
| 2.394196 | 2.758034 | -3.93377 | -3.624 | 6.301144 | 0.48846 | 2.526502 | 3.101285 | 4.866008 | 2.628592 |
| 1.661685 | -1.9097 | -5.15749 | -5.02544 | 3.075905 | 2.439089 | 1.306493 | 2.150106 | 4.844345 | 0.210545 |
| 2.778427 | 2.075821 | -0.05708 | -1.35676 | 6.185236 | NA | 4.512387 | 3.69415 | 4.830704 | 3.115092 |
| 2.802522 | 2.756789 | -0.84771 | -3.29694 | 6.78664 | -1.23688 | 5.081244 | 4.448339 | 5.197304 | 0.761124 |
| 2.140955 | 1.839627 | -4.68376 | -4.25682 | NA | 1.476383 | 2.264301 | 1.551989 | 1.450701 | 0.810148 |
| 2.293921 | -0.77899 | -5.24977 | -5.67 | 5.70463 | 1.14597 | 4.306564 | 1.32688 | 3.974566 | 2.509642 |
| 2.271076 | -0.14554 | -4.38193 | -5.15059 | 5.418199 | NA | 3.993312 | 1.249043 | 3.18646 | 2.444577 |
| 2.17257 | -0.89767 | -5.04208 | -7.94039 | 1.202687 | 2.37559 | 2.811065 | -2.33244 | 0.150967 | 5.025787 |
| 0.631467 | 0.78833 | -5.54595 | -5.65883 | 2.774839 | NA | 2.429474 | 2.722045 | 1.961254 | 1.288038 |
| 2.186926 | NA | NA | -3.62612 | 4.910921 | 2.688847 | 1.958305 | 2.858412 | 1.957558 | 1.47201 |
| 1.945236 | 2.284232 | -3.32961 | -4.25465 | 2.152238 | 2.865382 | 3.655207 | 3.190994 | 2.413563 | -0.05838 |
| 2.166279 | 0.844491 | -4.1076 | -5.2181 | 6.67437 | -0.75056 | 3.818017 | 1.904965 | 4.922946 | 2.286607 |
| 2.057661 | -2.30819 | -4.73884 | -6.53411 | 1.252308 | 0.546741 | 2.017476 | -0.85717 | 1.247448 | 3.131692 |
| 2.910936 | 2.075844 | -4.73424 | -4.69911 | 3.285981 | -1.80299 | 3.310153 | 3.647489 | 3.711561 | 1.210711 |
| 1.242122 | 0.141963 | -5.42332 | -6.5739 | 3.419227 | 0.398291 | 3.863923 | 2.361713 | 2.420504 | 2.55085 |
| 2.020151 | 0.199283 | -5.15239 | -4.63472 | 4.97628 | -0.67939 | 2.449736 | 1.54669 | 1.961914 | 0.370328 |
| 0.887086 | -1.24321 | -5.37744 | -6.23354 | 5.829653 | 0.100559 | 3.189537 | 0.248932 | 4.44898 | 2.217929 |
| 3.05562 | 0.268912 | -3.40812 | -4.59206 | 3.766438 | 0.390292 | 4.30568 | 1.774849 | 2.504765 | 4.516814 |
| 2.174391 | -1.33725 | -2.76805 | -1.75164 | 5.449754 | 2.681984 | 3.203466 | 1.478456 | 4.840384 | 2.987451 |
| 1.843668 | 2.128894 | -2.75133 | -2.82661 | 4.146647 | 0.740011 | 3.735937 | 2.184215 | 5.020102 | -0.19831 |
| 2.537613 | 0.327094 | -4.59125 | -5.39732 | 6.05066 | -0.52183 | 2.882352 | -0.33781 | 0.898401 | 0.333848 |
| 2.910971 | 0.530427 | -0.38199 | -2.99062 | 4.587478 | -0.59095 | 2.345194 | 1.829729 | 3.636253 | 5.218975 |
| 2.571314 | 2.493454 | -1.96981 | -3.03242 | 3.638469 | -2.6093 | 3.400361 | 0.926384 | 4.252522 | 2.737528 |
| 0.649039 | -1.22195 | -5.15673 | -5.77797 | 2.630944 | 1.54984 | 0.436813 | -1.85391 | -0.64234 | 1.757487 |
| 0.307983 | 2.478773 | -3.3037 | -4.04053 | 5.981389 | -0.67969 | 4.369952 | 3.267049 | 3.975247 | 4.020539 |
| 2.463241 | -0.09466 | -5.04887 | -4.31801 | 3.94661 | 1.795029 | 3.01523 | 0.589311 | 2.346433 | 1.568754 |
| 1.576236 | 0.361242 | -5.34738 | -7.10612 | 3.332216 | -1.60126 | 2.124655 | 0.932177 | 1.506473 | 3.287061 |
| 2.499183 | -1.41118 | -3.42406 | -4.37887 | 6.426475 | -1.49568 | 3.028693 | 0.982828 | 2.325654 | 2.969804 |
| 1.223475 | -1.30941 | -4.98083 | -2.74682 | 5.516876 | -0.12214 | 0.873905 | 2.930613 | 1.863746 | 2.819224 |
| 2.859113 | 1.803888 | -2.42298 | -3.45671 | 3.104516 | 0.024956 | 3.850392 | 3.109904 | 3.765882 | 1.869097 |
| 1.177055 | -1.38864 | -3.51626 | -4.1886 | 4.732643 | -2.80238 | 3.577239 | 0.955915 | 4.836068 | 2.9665 |
| NA | NA | NA | NA | NA | -0.38889 | NA | NA | NA | NA |
| 0.046192 | 0.051139 | 0.05385 | 0.054633 | 0.056934 | 0.061153 | 0.064404 | 0.065554 | 0.065939 | 0.066009 |

| JW-7-24-1 | Thapsigarg | Obatoclax | LAQ824 | ZM-447439 | XL-880 | GSK212645 | AZD6244 | CGP-08299 | Bosutinib |
|---|---|---|---|---|---|---|---|---|---|
| NA | NA | NA | NA | 5.480756 | NA | NA | 5.480756 | NA | 2.656999 |
| 0.355777 | -2.13324 | -1.17849 | -3.91763 | NA | 0.217238 | -4.00231 | NA | 3.060186 | NA |
| 1.1853 | -5.11575 | -1.78434 | -4.14376 | 1.562851 | -0.49023 | -3.1499 | 2.454812 | 2.128681 | 1.72868 |
| 0.445482 | -5.31169 | 0.135899 | -3.88531 | 1.009134 | 4.431127 | -3.63716 | 3.44023 | 3.564874 | 2.590433 |
| 1.304272 | -2.8268 | 0.871306 | -4.06745 | 4.568157 | 1.857114 | -4.49834 | 4.394458 | 3.238629 | 2.892677 |
| 2.004268 | -4.89544 | -1.0485 | -4.34486 | 4.311142 | 4.518124 | -1.3743 | 3.871579 | NA | 2.722894 |
| 1.340893 | -5.08611 | -0.60592 | -3.13865 | 3.763563 | 2.279205 | -3.06323 | 3.484443 | NA | 1.9985 |
| 2.438359 | 2.683596 | 1.248586 | -1.96697 | 5.36306 | 5.268187 | -2.72276 | 4.217242 | NA | 4.967891 |
| 1.50878 | -2.46336 | -0.26642 | -2.81172 | 1.359806 | 0.713231 | -4.01734 | 3.464542 | 1.775457 | 2.472247 |
| 2.586034 | -0.13376 | 1.253129 | -3.32931 | 3.55059 | 5.109895 | -3.52109 | 2.808105 | NA | 4.193449 |
| -0.4002 | 0.073659 | 0.185653 | -3.5759 | 3.072209 | 2.047729 | -6.37929 | 4.425886 | NA | 3.743738 |
| 2.000756 | -4.27621 | -0.9229 | -3.14377 | NA | 2.735684 | -2.72756 | NA | NA | NA |
| 2.984989 | -2.03654 | 0.908128 | -1.98248 | 4.934029 | 5.180889 | -1.27341 | 4.934029 | NA | 4.240882 |
| 1.684403 | NA | -2.23457 | -3.47334 | NA | 0.69211 | -3.21534 | NA | NA | 2.898111 |
| 1.887878 | 1.931399 | 2.597472 | 0.492632 | NA | 1.63411 | -2.59163 | NA | NA | NA |
| 0.804296 | -2.33625 | 1.115656 | -2.1538 | 3.290221 | 3.018819 | -4.40351 | NA | NA | 2.58059 |
| 2.070056 | -5.72532 | -0.26727 | -3.75082 | 2.686139 | 2.635083 | -5.31429 | 3.102471 | NA | 1.399532 |
| 1.523724 | -5.03841 | 0.811372 | -3.6602 | 3.158159 | -0.30959 | -2.81245 | 4.159882 | NA | 3.5632 |
| 2.086841 | -0.42917 | 0.116684 | -3.17141 | NA | 0.57152 | -2.70315 | NA | 3.227284 | NA |
| 5.002746 | -7.54894 | -4.45886 | -3.7321 | 2.883385 | 4.33264 | 0.64376 | 3.736568 | NA | 3.165394 |
| 1.862823 | -4.74173 | -0.88955 | -3.94866 | 2.760714 | 1.677758 | -3.24442 | 4.221612 | NA | 2.632323 |
| NA | -3.71359 | -0.30854 | -3.8313 | 4.93871 | 1.791367 | NA | 4.845031 | NA | 4.270215 |
| 2.261426 | -1.91729 | -1.3882 | -3.11027 | 4.027359 | 1.228759 | -5.01855 | 4.224651 | 3.682369 | 3.210853 |
| 1.005325 | -1.82405 | 1.282379 | -1.16918 | 2.754761 | 2.752267 | -3.9266 | 2.870975 | NA | 2.792236 |
| 2.129099 | -5.631 | -2.59965 | -2.767 | 0.542423 | 1.351984 | -2.30461 | 3.02632 | NA | 2.816245 |
| 0.551362 | -4.19703 | 0.809731 | -2.27477 | 2.545258 | 4.098559 | -5.1764 | 1.991434 | NA | 1.829475 |
| 1.317001 | -5.05049 | -2.14404 | -2.41035 | 1.472728 | 0.592483 | -2.50046 | 1.762746 | NA | NA |
| 1.081228 | -5.28083 | -2.79105 | -3.12194 | 3.206764 | 0.599672 | -4.81334 | 4.010753 | NA | 3.80076 |
| 1.365689 | -5.77074 | -0.06454 | -1.71324 | 1.641224 | 0.584112 | -3.80542 | 2.700076 | NA | 1.547244 |
| 2.917543 | -6.12274 | 0.366343 | -1.21646 | 3.143651 | 1.006063 | -2.3313 | 1.377238 | NA | 1.415012 |
| 2.717674 | -0.32256 | -0.12634 | -4.0994 | 0.924759 | 0.865793 | -0.81243 | 3.176425 | NA | NA |
| 1.509786 | -0.60006 | 0.466845 | -3.02268 | 3.729705 | 1.941375 | -5.32867 | 4.282518 | NA | 2.850938 |
| 1.629414 | -3.71768 | 0.066243 | -2.76868 | NA | 0.966941 | -3.79764 | NA | NA | 3.194034 |
| 5.217906 | NA | -1.15925 | 2.768001 | NA | 0.724067 | 0.743892 | 4.174237 | NA | 2.562327 |
| 1.113575 | NA | -1.51049 | -3.20572 | 2.275664 | -0.12677 | -2.452 | 4.240978 | NA | 3.016 |
| 4.999977 | -5.8761 | -2.04845 | -3.47357 | NA | 2.277107 | -1.57383 | 2.603273 | NA | 2.403321 |
| 3.095327 | -4.99304 | 0.058235 | -0.825 | NA | 2.019785 | -0.45294 | 4.693503 | NA | 2.392762 |
| 1.45652 | -1.52222 | 0.76214 | 0.652347 | 0.81349 | -0.23737 | -3.48295 | 2.608428 | 3.610325 | 1.865993 |
| 1.833053 | -5.08034 | -0.94566 | -2.95894 | NA | 1.4883 | -3.28904 | 3.257606 | NA | 0.57157 |
| 1.166043 | -3.36416 | -1.48423 | -2.39339 | 1.39531 | 0.243846 | -3.8235 | 2.23605 | NA | 2.711429 |
| 2.166094 | -5.02913 | 0.837056 | -1.97923 | 2.942808 | 0.845742 | 0.933966 | 3.847504 | NA | 2.196235 |
| 1.001303 | -7.2067 | -0.46614 | -2.64445 | 4.148706 | 1.487981 | -2.74082 | 4.14023 | NA | 0.644639 |
| 1.971958 | -6.00022 | -0.13306 | -2.62992 | 2.450001 | 4.101236 | -1.27819 | 1.144875 | NA | 2.615692 |
| NA | NA | NA | NA | 3.268785 | NA | NA | 4.180838 | NA | 1.305018 |
| 0.068976 | 0.070695 | 0.071947 | 0.082062 | 0.082706 | 0.083374 | 0.089742 | 0.093223 | 0.09526 | 0.09545 |

| rTRAIL | CCT01815( | HG-5-113-( | HG-6-64-1 | FH535 | KIN001-24( | OSI-906 | Docetaxel | AZD-0530 | SB 216763 |
|---|---|---|---|---|---|---|---|---|---|
| 1.500469 | 3.918395 | 3.542845 | NA | NA | NA | NA | -3.83588 | NA | 4.404266 |
| NA | NA | NA | 1.79406 | 2.55494 | 1.695807 | 3.661304 | NA | 1.851124 | NA |
| -1.15679 | 2.572675 | NA | 0.314659 | 1.863958 | 2.646711 | 1.366059 | -4.09655 | 2.540592 | 4.201356 |
| -2.04108 | 1.770152 | 1.472246 | 1.608242 | 2.309332 | 4.265847 | 1.689439 | -5.48222 | 3.166796 | 5.099654 |
| -1.19175 | 2.453471 | 2.130509 | 3.478511 | 1.384061 | 2.615076 | 4.250691 | -2.17493 | 2.44106 | 4.562455 |
| 0.443602 | 3.283986 | 3.276492 | 3.801642 | 1.673764 | 4.303559 | 3.135243 | -3.65992 | NA | 5.261353 |
| -0.06873 | 5.053628 | NA | 1.350589 | 0.228277 | 3.667181 | 3.396382 | -2.40918 | 3.259376 | 4.733403 |
| 2.174314 | 6.816611 | 5.932732 | 6.056788 | 6.648677 | 4.224653 | 4.305548 | 0.252336 | NA | 7.029859 |
| -0.28626 | 2.440111 | 1.59133 | 3.209683 | 2.639138 | 1.497584 | 3.053821 | -6.13827 | 1.95984 | NA |
| 1.226803 | 5.210782 | 4.24451 | 3.54544 | 1.462474 | 4.628305 | 4.401204 | -2.19718 | NA | 6.020256 |
| 1.134091 | 3.664499 | NA | 3.461654 | 3.062713 | 2.172093 | 3.794077 | -5.46532 | NA | 5.088006 |
| -0.8235 | 4.295374 | NA | -0.38787 | 0.086225 | 3.291395 | 3.558849 | NA | NA | NA |
| 0.646657 | NA | 3.852632 | 1.914388 | 1.611409 | 3.903246 | 4.136907 | -5.62306 | NA | NA |
| 1.131631 | 4.903527 | NA | 0.668381 | 1.74551 | 2.601435 | 2.132381 | -6.62905 | NA | 5.965988 |
| NA | NA | NA | 3.949226 | 3.371274 | 3.635612 | 3.694717 | NA | 1.244721 | NA |
| 0.361799 | 4.092588 | 4.328073 | 4.991147 | 0.869543 | 3.687774 | 4.052268 | -3.42996 | NA | 5.524027 |
| -0.36964 | 2.775558 | NA | 1.49365 | 0.993528 | 3.558056 | 3.032584 | -5.49929 | NA | 4.804017 |
| -0.1625 | 3.516382 | 2.270182 | 1.501389 | 2.374012 | 3.339724 | 2.973036 | -4.80044 | NA | 5.169576 |
| NA | NA | NA | 2.829438 | 2.147825 | 2.937106 | 3.320672 | NA | 2.906149 | NA |
| 0.279185 | 2.128981 | NA | 0.949836 | 1.021694 | 4.894026 | 0.448191 | -6.40977 | NA | 2.906915 |
| 0.341073 | 1.70806 | 1.671553 | 1.99253 | 0.104195 | 2.917469 | 2.789401 | -6.02519 | NA | 4.18593 |
| 0.692748 | 5.671195 | 3.039057 | NA | 0.610859 | 3.471968 | 2.326066 | -1.47089 | NA | 5.879653 |
| 0.57111 | 2.213715 | 2.771177 | 2.737666 | 1.669397 | 3.117458 | 3.911251 | -3.83553 | 2.888081 | 5.00926 |
| -0.09795 | 2.93279 | 2.728544 | 2.751498 | 2.101645 | 4.391499 | 3.816043 | -5.56278 | NA | 5.55528 |
| -0.31175 | 1.400475 | NA | -1.13169 | 1.208764 | 4.963725 | 2.739712 | -5.6823 | NA | 3.070165 |
| 0.48992 | 2.564539 | 2.708636 | 4.047373 | 0.778678 | 5.92171 | 3.929662 | -4.26491 | NA | 5.362344 |
| -4.16736 | 2.641337 | NA | 0.473206 | 2.977856 | 3.479389 | 2.051525 | NA | NA | NA |
| 0.643499 | 2.58948 | 2.633306 | 0.686756 | 1.407174 | 3.992307 | 3.752135 | -2.19152 | NA | 4.909582 |
| -1.19018 | 3.232116 | 0.551386 | 2.528249 | 0.30526 | 2.475908 | 3.325888 | -5.16536 | NA | 4.760217 |
| -0.858 | 3.028873 | NA | 0.992723 | 2.238043 | 4.24065 | 0.772633 | -4.65038 | NA | 5.48798 |
| -1.74202 | 3.015463 | NA | 0.993576 | 1.945519 | 4.760607 | 3.342162 | NA | NA | NA |
| 0.502662 | NA | NA | 3.631269 | 2.206704 | 2.82643 | 3.691203 | -5.378 | NA | 4.974592 |
| -1.84244 | 3.338919 | NA | 2.160284 | 1.701771 | 2.489552 | 3.52281 | -6.07003 | NA | 4.840127 |
| 0.473654 | 3.745105 | 2.693497 | 2.121468 | 2.069653 | 5.020028 | 3.112799 | -5.61662 | NA | 4.684878 |
| 0.943282 | 4.443201 | NA | NA | 0.378993 | 4.172925 | 0.641903 | -5.46326 | NA | NA |
| 0.510539 | 3.60148 | 2.117514 | 1.333259 | -0.9453 | 3.715569 | 1.213337 | -4.00633 | NA | 4.858717 |
| -2.14159 | 4.994376 | NA | 1.898484 | 0.09397 | 3.145345 | 3.862595 | -4.77482 | NA | NA |
| -1.68965 | 2.22274 | NA | 3.703008 | 2.545915 | 3.147812 | 2.095443 | -6.60204 | 2.797581 | 4.665233 |
| -0.43037 | 3.221702 | 2.089646 | -0.63374 | 1.127993 | 3.542556 | 3.232956 | -6.96885 | NA | 4.36378 |
| -1.57557 | 3.00285 | NA | -0.00976 | 2.036691 | 2.507093 | 3.183724 | -5.54507 | NA | 5.234254 |
| -2.06341 | 4.725347 | 2.798055 | 0.299582 | -0.58874 | 3.415816 | 3.872751 | -3.89421 | NA | 5.245707 |
| -0.07163 | 3.338595 | 2.996711 | 3.318293 | 2.044354 | 3.663978 | 3.32569 | -4.40804 | NA | NA |
| -0.47689 | 1.236454 | NA | 0.373792 | 2.595457 | 4.809667 | 2.378074 | -5.16325 | NA | 3.495015 |
| 0.900967 | 3.016594 | NA | NA | NA | NA | NA | -5.83927 | NA | 5.726514 |
| 0.09648 | 0.099364 | 0.102334 | 0.103794 | 0.104776 | 0.107541 | 0.107727 | 0.111103 | 0.112012 | 0.114303 |

| JNJ-268541 | MLN4924 | BMS-70816 | Tipifarnib | ABT-888 | CEP-701 | RDEA119 (r | CMK | QL-XII-61 | Cyclopamir |
|---|---|---|---|---|---|---|---|---|---|
| 4.681742 | NA | 5.599381 | NA | 5.695832 | 1.946481 | NA | NA | 6.210492 | NA |
| NA | NA | NA | 2.104874 | NA | NA | NA | 0.592935 | NA | 4.033746 |
| 1.909191 | -2.613 | 3.361825 | 0.694831 | 3.508208 | -1.67239 | -2.66685 | 2.579957 | NA | 4.667482 |
| 2.772485 | 2.481303 | 3.679676 | -0.78586 | 4.407411 | -0.96178 | 4.278093 | 3.066445 | NA | 4.767863 |
| 4.504446 | 2.910458 | 4.670892 | 2.314874 | 4.777435 | 2.905273 | 4.51629 | 2.24144 | NA | 4.486272 |
| 3.598651 | -2.88626 | 4.466691 | 2.800532 | 4.60205 | 2.959007 | 4.522936 | NA | 4.65419 | NA |
| 3.753397 | 2.716094 | 4.057597 | 1.744822 | 4.003889 | 2.337575 | 2.830235 | NA | NA | NA |
| 3.897835 | 4.476899 | 6.179653 | 6.89163 | 6.336712 | 2.796737 | 5.863028 | NA | 6.174516 | NA |
| 2.781531 | NA | 3.172789 | 2.281768 | 3.723257 | -0.60253 | 3.233239 | 0.720767 | 2.077513 | 2.555326 |
| 2.838202 | NA | 3.138265 | 1.692912 | 4.143298 | 3.582932 | 4.329572 | NA | 5.117352 | NA |
| 2.539164 | 2.199484 | 3.208302 | 2.478212 | 5.270697 | 1.535883 | 4.376772 | NA | NA | NA |
| 2.84018 | 0.423758 | 4.168237 | 3.148643 | NA | NA | 4.531278 | NA | NA | NA |
| NA | NA | 3.368235 | 1.511306 | 5.156107 | 1.167005 | 2.725961 | NA | 5.374866 | NA |
| 4.513746 | 3.040382 | 4.906138 | 0.423744 | 5.257239 | -1.78106 | 5.054819 | NA | NA | NA |
| NA | NA | NA | 4.900214 | NA | NA | NA | NA | NA | 5.526151 |
| 2.282531 | NA | 5.071027 | 1.886254 | 4.951282 | 2.841741 | 2.130337 | NA | 5.633429 | NA |
| 1.83097 | -0.29369 | 2.593747 | 0.690137 | 3.742485 | -0.49785 | 2.717748 | NA | NA | NA |
| 2.43479 | -0.37193 | 2.032903 | 1.803873 | 4.44312 | 0.740594 | 3.661828 | NA | 4.046917 | NA |
| NA | NA | NA | 0.170803 | NA | NA | NA | 2.39832 | NA | 5.021899 |
| 2.704935 | -1.29545 | 3.214564 | -1.62865 | 4.308923 | 0.151008 | 3.775071 | NA | NA | NA |
| 1.846832 | NA | 1.790222 | 0.490158 | 4.444755 | 0.22069 | NA | NA | NA | NA |
| 4.465345 | 1.386132 | 4.469956 | 0.793659 | 4.84285 | 3.676188 | 4.407459 | NA | 4.810359 | NA |
| 3.061318 | NA | 2.41159 | 3.07137 | 4.615412 | 0.508437 | 4.118347 | 4.292005 | 0.570398 | 5.568235 |
| 4.02703 | 2.305944 | 3.197845 | 1.958622 | 4.220982 | 0.490009 | 3.398638 | NA | 5.414475 | NA |
| 2.247961 | -1.54572 | 3.429015 | 0.739831 | 3.279957 | -1.34564 | 2.754388 | NA | NA | NA |
| 2.222434 | NA | 3.336689 | 1.686713 | 4.591148 | 1.673511 | 1.685757 | NA | 4.867473 | NA |
| 1.711343 | -3.50973 | 3.908938 | 2.265627 | NA | NA | 2.48813 | NA | NA | NA |
| 2.504819 | -1.58073 | 2.724011 | 0.86475 | 4.895682 | 2.420272 | 2.179113 | NA | 4.413981 | NA |
| 2.68385 | -0.99383 | 3.020614 | 2.035276 | 4.06707 | -0.703 | 2.320777 | NA | 4.540751 | NA |
| 4.061459 | 1.550549 | 4.001189 | 1.986907 | 4.789686 | -0.15531 | 2.97531 | NA | NA | NA |
| 2.614476 | -0.49993 | 1.805924 | 2.069291 | NA | NA | 4.123087 | NA | NA | NA |
| NA | 0.865289 | 2.435318 | 3.649794 | 4.550098 | 0.473524 | 3.081593 | NA | NA | NA |
| 2.666907 | -1.34044 | 2.747917 | 0.086372 | 3.080514 | 0.484247 | 1.623372 | NA | NA | NA |
| 2.515175 | -0.02142 | 4.502111 | 2.447402 | 4.481857 | -0.13238 | 1.402852 | NA | 4.81176 | NA |
| 3.198552 | 0.982418 | 3.879385 | 2.225496 | 5.076394 | 0.342908 | 0.70186 | NA | NA | NA |
| 3.108527 | 1.329763 | 3.828285 | -0.59248 | 3.928542 | 1.570387 | 4.231164 | NA | 2.593285 | NA |
| 2.629314 | 0.133281 | 3.774845 | 1.620049 | 4.916647 | 0.228733 | 2.302148 | NA | NA | NA |
| 2.768269 | -1.50888 | 2.41319 | 1.644382 | 3.723075 | -1.51661 | NA | 2.902331 | NA | 4.785403 |
| 2.317601 | NA | 4.217604 | -0.61262 | 3.722447 | 0.265137 | 1.337081 | NA | 2.898506 | NA |
| 2.535295 | -0.44625 | 4.744778 | 1.960682 | 4.805004 | -1.31539 | 2.302913 | NA | NA | NA |
| 2.605861 | NA | 4.289936 | 2.024524 | 3.97146 | 0.824123 | 2.270231 | NA | 4.545949 | NA |
| 2.755637 | 0.193685 | 3.295456 | 1.723601 | 3.751069 | 0.658549 | 2.608789 | NA | 3.429358 | NA |
| 1.411001 | 0.087059 | 4.100888 | 1.776776 | 3.69588 | -0.52073 | 0.731456 | NA | NA | NA |
| 1.566562 | -0.39846 | 4.814345 | NA | 4.962313 | -1.28433 | 2.9555 | NA | NA | NA |
| 0.115563 | 0.119966 | 0.12303 | 0.123973 | 0.125532 | 0.129035 | 0.133285 | 0.134198 | 0.13524 | 0.138428 |

| JW-7-52-1 | JQ12 | 681640 | DMOG | SL 0101-1 | ABT-869 | ATRA | Gemcitabir | Vismodegil | OSI-027 |
|---|---|---|---|---|---|---|---|---|---|
| NA | NA | 4.787609 | NA | 6.326907 | NA | 5.913542 | NA | 6.397047 | NA |
| -2.89888 | 0.658535 | NA | 4.800609 | NA | 2.853722 | NA | -3.09324 | NA | -0.71624 |
| -2.3126 | 1.157995 | 1.509024 | 6.805193 | 3.001991 | 1.711145 | 3.413325 | -5.01156 | 3.734374 | 1.086309 |
| -1.51082 | 0.582376 | 3.49112 | 8.302976 | 4.701431 | 2.821689 | 2.335241 | 0.195478 | 3.242529 | 1.052841 |
| -1.3157 | 2.017919 | 3.552097 | 9.81835 | 5.563018 | 2.953605 | 4.596423 | 1.311052 | 5.407425 | 0.680917 |
| NA | -0.04119 | 2.591429 | 9.89317 | 4.979725 | 2.594997 | 2.489689 | 2.550824 | 5.36002 | 2.014411 |
| NA | 1.671681 | 3.08015 | 5.078331 | 4.694202 | 2.024091 | 2.615885 | 0.037159 | 4.20701 | -1.29442 |
| NA | 3.424973 | 3.291616 | 10.67158 | 4.779451 | 3.99531 | 4.663379 | 4.509703 | 6.319405 | 0.340746 |
| -4.45874 | 2.160317 | 2.192616 | 7.755202 | 4.416426 | 2.137558 | 2.814561 | 0.895603 | 4.416426 | 1.419357 |
| NA | -0.24063 | 1.80064 | 8.948995 | 3.669713 | 3.196481 | 2.011709 | 2.528603 | 4.785616 | 0.941729 |
| NA | 1.219361 | 4.220548 | 9.792483 | 5.90854 | 3.498132 | 4.67825 | 0.794021 | 5.744801 | -2.17417 |
| NA | 0.32941 | NA | 7.870519 | NA | 2.743201 | NA | -1.11814 | NA | 1.422614 |
| NA | 1.465682 | NA | 7.810074 | 5.819851 | 3.571451 | 5.035765 | 1.726412 | 5.85032 | 4.818514 |
| NA | -1.43618 | NA | 6.804465 | NA | 2.559363 | 3.821927 | 2.806349 | 5.524393 | 2.290799 |
| NA | 4.365008 | NA | 7.90334 | NA | 2.770801 | NA | 1.534162 | NA | 3.627549 |
| NA | 2.370684 | 3.351804 | 9.141169 | 5.810917 | 3.390467 | 3.708794 | 1.340783 | 5.360204 | -0.16867 |
| NA | -1.11214 | 2.152506 | 7.296818 | 4.903945 | 2.681425 | 3.792885 | -1.17001 | 4.585691 | 1.688642 |
| NA | 1.891398 | 1.417337 | 6.837124 | 4.734139 | 2.844467 | 5.130739 | -3.63992 | 4.62168 | 0.182161 |
| -2.10056 | 1.965852 | NA | 7.855056 | NA | 2.246354 | NA | -3.36089 | NA | -0.09429 |
| NA | -1.12287 | 1.181031 | 7.941586 | 4.270219 | 2.385033 | 2.808433 | -5.95855 | 4.99363 | 5.607905 |
| NA | 0.588562 | 3.295951 | NA | 5.137902 | 2.367553 | 2.453498 | -3.49658 | 5.137902 | 2.014428 |
| NA | NA | 4.212394 | NA | 5.444927 | NA | 5.879653 | 1.477012 | 4.041832 | NA |
| -1.14216 | 2.655405 | 3.432221 | 9.676272 | 5.308559 | 2.864646 | 5.063419 | 2.384115 | 5.173205 | -0.14762 |
| NA | 4.486033 | 2.167905 | 6.67233 | 5.300326 | 2.906387 | 4.608966 | -1.68455 | 3.65304 | 1.426915 |
| NA | 0.524545 | 1.394004 | 5.39711 | 2.777539 | 2.146779 | 2.877054 | -1.56469 | 3.017803 | 2.678976 |
| NA | 0.609689 | 0.857607 | 6.231704 | 1.904346 | 2.031507 | 4.507808 | 0.432497 | 4.653538 | -0.06372 |
| NA | 1.424879 | 0.525068 | 7.320565 | 3.112376 | 2.347617 | NA | -5.32407 | NA | 1.925877 |
| NA | 2.678611 | 0.797343 | 7.728156 | 4.724005 | 2.642446 | 4.229809 | -2.4248 | 4.368371 | -0.31203 |
| NA | 2.816709 | NA | 8.497674 | 4.116967 | 2.453384 | 4.394096 | -4.16905 | 4.348647 | -0.47723 |
| NA | 4.575116 | 2.409198 | 6.404601 | 4.598145 | 2.685941 | 5.48798 | 0.022364 | 5.429744 | 2.79892 |
| NA | -0.21974 | 0.741186 | 6.249964 | 3.216069 | 2.658272 | NA | 2.265695 | NA | 1.442087 |
| NA | 3.137281 | 3.375071 | 10.16491 | 4.648378 | 2.277055 | 5.243246 | 1.846299 | 5.173619 | -1.29671 |
| NA | 0.897811 | NA | 5.957088 | NA | 2.296925 | 4.35777 | NA | 3.338456 | 1.873987 |
| NA | 4.388568 | NA | 7.584461 | 3.875313 | 2.912099 | 2.591386 | 2.620321 | 4.614078 | 5.888406 |
| NA | 0.518362 | 4.188302 | 9.305967 | 5.552988 | 3.278764 | 5.759617 | 1.71759 | 4.359712 | 2.020064 |
| NA | -0.10398 | NA | 4.421241 | 4.647387 | 2.691097 | 1.75536 | -5.36974 | 4.815721 | 3.728964 |
| NA | 2.397083 | NA | 6.964447 | 5.609794 | 3.193624 | 5.609794 | -0.30064 | 5.609794 | 4.743981 |
| -0.84077 | 3.340449 | 2.504495 | 8.490384 | 4.052662 | 1.973638 | 4.319965 | 2.305829 | 4.453697 | 1.181945 |
| NA | 0.875417 | NA | 6.532645 | 4.437302 | 2.795194 | 4.195956 | -3.52194 | 4.410526 | 1.288736 |
| NA | 2.870613 | 1.559796 | 6.577786 | 3.723394 | 3.199276 | 4.588209 | -0.82042 | 5.515466 | -1.18433 |
| NA | 2.937933 | 2.31913 | 8.209999 | 5.50421 | 3.147182 | 5.476998 | 2.685291 | 5.378882 | 4.406863 |
| NA | 3.709914 | 3.75321 | 6.703118 | 5.261733 | 2.690671 | 5.159597 | 3.08421 | 4.691495 | 2.079166 |
| NA | 0.335558 | 1.558842 | 5.854509 | 4.7704 | 2.467907 | 3.403775 | -3.52702 | 3.719444 | 2.244415 |
| NA | NA | 3.207555 | NA | 5.726514 | NA | 5.711773 | NA | 5.67846 | NA |
| 0.146197 | 0.150386 | 0.151827 | 0.153487 | 0.153778 | 0.157032 | 0.159559 | 0.159672 | 0.163034 | 0.169818 |

| Embelin | JQ1 | MG-132 | BMS-5097 | EX-527 | THZ-2-49 | GSK-65039 | GSK26996 | ZSTK474 | Axitinib |
|---|---|---|---|---|---|---|---|---|---|
| NA | NA | NA | NA | NA | NA | NA | 3.725545 | NA | 4.337563 |
| 2.236298 | 0.355273 | 2.539951 | 3.803622 | 5.766681 | 1.181387 | 3.468804 | NA | 0.018692 | NA |
| 2.245511 | -1.05208 | -1.87229 | 2.739073 | 4.314186 | 2.626697 | 2.790786 | 1.991969 | 1.524103 | 1.163854 |
| 3.772285 | -1.38014 | -1.28936 | 4.078939 | 5.752085 | 2.918373 | 2.388889 | 4.207025 | 1.444755 | 1.163471 |
| 2.845983 | 0.475235 | 1.819109 | 3.523075 | 5.982479 | 2.134374 | 6.009931 | 1.351668 | 0.245591 | 3.545302 |
| 3.488467 | -1.589 | NA | NA | 6.030451 | 3.076633 | 5.306985 | 4.549292 | 2.927051 | 2.909833 |
| 3.859959 | 0.390088 | 1.297333 | NA | 5.633608 | 1.863802 | 5.421689 | 1.106155 | -0.35298 | 2.738465 |
| 5.373178 | 1.713639 | NA | NA | 6.974184 | 5.020603 | 7.281311 | 6.086337 | -1.03472 | 5.105473 |
| 3.922125 | -0.52487 | -1.41645 | 2.80703 | 4.799248 | 0.398588 | 4.244858 | 1.939491 | -0.60044 | 1.367242 |
| 3.117759 | -0.40986 | NA | NA | 6.713403 | 3.961187 | 6.48338 | 4.436697 | 0.881687 | 1.338609 |
| 5.508252 | 1.336326 | NA | NA | 6.705712 | 1.364992 | 5.990336 | 1.725044 | -1.6085 | 4.336429 |
| 1.840056 | -2.20126 | NA | NA | 5.130602 | 2.008503 | 2.857823 | 3.32553 | 1.307269 | NA |
| 3.42389 | 0.837796 | NA | NA | 6.261377 | 2.405436 | 3.897212 | 4.113742 | 2.49146 | 2.325264 |
| 1.828539 | -0.21511 | NA | NA | 6.643156 | -0.74553 | 2.859891 | 2.838657 | 0.120672 | 2.167109 |
| 3.620326 | 3.655574 | 0.766552 | NA | 5.75023 | 2.610738 | 5.520821 | NA | 0.73024 | NA |
| 3.575904 | 1.035515 | NA | NA | 6.503316 | 1.39849 | 6.235456 | 2.009388 | 0.206986 | 3.656045 |
| 2.40523 | -1.26942 | NA | NA | 5.327621 | 3.158388 | 2.694132 | 2.762548 | 0.061543 | 2.82464 |
| 2.927231 | -1.36519 | NA | NA | 5.151719 | 0.475768 | 4.10694 | 2.918294 | 1.493841 | 2.601906 |
| 3.884631 | -0.21002 | 0.719469 | 3.781098 | 4.858882 | 0.204901 | 3.772884 | NA | 1.797164 | NA |
| 1.989126 | -0.15027 | NA | NA | 5.36174 | 3.544424 | 0.351968 | 1.334062 | 4.325176 | 1.850646 |
| 2.512362 | 0.859036 | NA | NA | 5.830233 | 0.209811 | 3.7665 | 2.475084 | 0.967897 | 2.079958 |
| 2.559318 | NA | NA | NA | 6.216284 | 0.689641 | 3.268569 | 2.936197 | -0.56331 | 1.545527 |
| 2.82829 | 2.235082 | 1.76006 | 4.977363 | 4.900594 | 1.710247 | 5.754847 | 1.957923 | -0.66441 | 3.481672 |
| 2.820834 | 1.968401 | NA | NA | 5.869215 | 0.984528 | 4.891124 | 4.686301 | 0.587281 | 3.338919 |
| 1.604846 | 1.15553 | NA | NA | 4.724464 | 2.668392 | 1.41858 | 1.183188 | 1.692435 | 1.259479 |
| 2.73102 | 2.978862 | NA | NA | 5.907159 | 3.268233 | 3.908926 | 3.525441 | -0.48497 | 3.030144 |
| 3.060597 | 1.016607 | NA | NA | 4.890028 | 0.659282 | 4.094732 | 2.211053 | 1.876722 | NA |
| 2.504025 | -1.3984 | NA | NA | 6.13477 | 5.254831 | 3.916075 | 3.179923 | -0.5317 | 1.758864 |
| 3.049108 | -0.55859 | NA | NA | 4.523331 | 1.062751 | NA | 2.214895 | -0.37632 | 0.766138 |
| 2.064274 | 2.675998 | NA | NA | 5.553381 | 1.735295 | 4.050058 | 3.655111 | 1.481488 | 3.878542 |
| 2.144695 | 0.270872 | NA | NA | 5.566114 | 1.003842 | 5.214038 | 2.535865 | 3.749291 | NA |
| 5.7155 | -0.71409 | NA | NA | 5.165988 | 1.879239 | 2.794432 | 2.566607 | -0.8028 | 3.346557 |
| 4.425117 | -0.80312 | NA | NA | 5.468512 | -0.16681 | 5.060895 | 4.095263 | -0.1253 | 1.632337 |
| 2.537302 | 1.398676 | NA | NA | 5.452937 | 2.395207 | 4.356067 | 1.488263 | 4.477394 | 1.126848 |
| 2.240429 | 1.4439 | NA | NA | 6.337303 | 0.835363 | 6.204293 | 4.46801 | 1.122412 | 2.522587 |
| 1.848054 | -0.43224 | NA | NA | 5.94181 | 2.242307 | -0.04494 | 2.21651 | 0.045434 | 3.500746 |
| 2.13356 | 1.212375 | NA | NA | 6.049689 | 0.699771 | 5.550956 | 3.160079 | 3.081926 | 3.553313 |
| 3.698391 | 0.434376 | 1.687883 | 3.801558 | 5.1964 | 0.514671 | 4.344466 | NA | -0.6076 | 1.061314 |
| 2.63515 | 0.280793 | NA | NA | 5.538092 | 2.215089 | 3.401338 | 2.407959 | 0.801386 | 1.138446 |
| 3.415464 | 0.231339 | NA | NA | 6.213909 | 0.622118 | 3.284125 | 0.127621 | 0.483033 | 3.504139 |
| 3.397197 | -1.50429 | NA | NA | 6.185189 | 0.91476 | 5.802253 | 2.15289 | 3.1166 | 1.163125 |
| 2.486121 | 1.325434 | NA | NA | 6.050897 | 0.014127 | 4.396969 | 3.845945 | 1.009152 | 1.355 |
| 3.051471 | 0.611986 | NA | NA | 5.114497 | 3.069361 | 3.861695 | 2.063798 | 2.489333 | 2.23036 |
| NA | NA | NA | NA | NA | NA | NA | -0.8741 | NA | 2.962346 |
| 0.170384 | 0.17044 | 0.170829 | 0.173391 | 0.181008 | 0.183002 | 0.186421 | 0.187742 | 0.192555 | 0.198653 |

| RDEA119 | FK866 | QL-VIII-58 | Imatinib | XMD15-27 | XMD14-99 | Cytarabine | GSK107091 | QL-X-138 | PD-032590 |
|---|---|---|---|---|---|---|---|---|---|
| 5.176746 | 3.902735 | 1.113515 | NA | NA | NA | 1.207208 | NA | NA | 0.174232 |
| NA | NA | NA | 2.864849 | 4.562693 | 4.402772 | NA | -2.19886 | -0.28236 | NA |
| -3.06867 | -6.74183 | NA | 2.590154 | 4.213962 | 2.303815 | -1.25668 | 0.719763 | NA | -5.70496 |
| 2.904582 | -4.11779 | -1.74632 | 3.49112 | 5.124274 | 5.124147 | 1.475061 | 2.68102 | 1.51637 | 0.80473 |
| 4.020655 | -2.26477 | -0.18452 | 2.655552 | 4.619305 | 5.542231 | 4.003831 | 3.966103 | 0.024132 | 1.924389 |
| 4.580163 | 1.817009 | 0.082566 | NA | 5.210188 | 5.384145 | 1.948403 | 3.765461 | 2.340197 | 0.013377 |
| 2.286524 | -3.1132 | NA | 2.684335 | 4.590386 | 5.133677 | 3.519332 | 3.497832 | 0.14051 | 1.370302 |
| 4.584542 | 3.416614 | -1.49821 | NA | 6.968761 | 6.643063 | 5.239041 | 5.441079 | 0.516792 | 2.408612 |
| 3.229515 | 0.722868 | -2.90103 | 2.806988 | 4.440143 | 4.424356 | -0.53475 | 2.989147 | 1.560961 | -0.53651 |
| 4.655122 | 3.326869 | -1.5966 | NA | 5.906383 | 5.905589 | 0.756129 | 3.807521 | 0.731533 | -1.98837 |
| 4.015701 | -2.00293 | NA | NA | 6.007905 | 5.177829 | 1.770175 | 2.883008 | -0.24484 | 2.083251 |
| NA | 0.487673 | NA | NA | 5.28892 | 5.092454 | NA | 2.36635 | 1.411114 | NA |
| 2.064652 | 1.975284 | -1.86136 | NA | 5.874036 | 5.766867 | 0.631588 | 4.485362 | 4.275551 | -0.53763 |
| NA | 0.122172 | NA | NA | 5.999238 | 5.958738 | 0.49257 | 2.185492 | 1.64194 | NA |
| NA | NA | NA | 1.814045 | 5.038193 | 5.074121 | NA | 2.970151 | 3.430962 | NA |
| 2.729722 | NA | -2.19507 | NA | 5.497662 | 5.575379 | 2.5027 | 4.351595 | -0.2528 | 1.48238 |
| 1.710493 | 1.61474 | NA | NA | 4.677538 | 4.981215 | 0.919788 | 3.527086 | 1.292157 | -0.9677 |
| 3.510112 | 2.744704 | -0.50775 | NA | 3.739851 | 5.189259 | 0.369613 | 2.334635 | 0.360595 | 1.169274 |
| NA | NA | NA | 2.97865 | 4.683591 | 4.70035 | NA | 2.945014 | 0.905476 | NA |
| 3.73536 | -2.523 | NA | NA | 5.025787 | 4.99735 | 0.443234 | 3.639493 | 4.877221 | -0.30279 |
| 3.690994 | NA | -1.79593 | NA | 4.649195 | 5.086311 | -1.64006 | 3.397651 | 1.109333 | -1.76983 |
| 5.023067 | 1.193695 | -0.50244 | NA | NA | NA | 1.157034 | 2.247011 | NA | 1.893057 |
| 2.579432 | 0.779299 | -1.93882 | 3.597029 | 5.332276 | 5.329311 | 2.850993 | 3.088846 | -0.87651 | 0.698121 |
| 2.246649 | 1.742538 | -0.93705 | NA | 5.456034 | 5.602609 | 3.963279 | 1.120644 | 1.303384 | -0.47801 |
| 1.295966 | -6.39981 | NA | NA | 4.449373 | 4.057369 | -1.266 | 2.462361 | 2.33895 | 0.110679 |
| 2.629019 | 1.447775 | -2.99589 | NA | 4.50865 | 3.781951 | 2.514348 | 2.713509 | -0.41866 | -1.08199 |
| 2.113863 | -3.85864 | NA | NA | 4.101478 | 3.93738 | NA | 3.051867 | 1.457783 | -1.05913 |
| 2.781486 | -2.40382 | -1.88247 | NA | 4.79076 | 4.148068 | 2.307443 | 1.060117 | 1.059062 | -0.41308 |
| 1.637065 | NA | -2.97429 | NA | 4.248815 | 3.823861 | -0.02301 | -3.81546 | -0.02068 | -2.61747 |
| 0.757705 | 1.277525 | NA | NA | 5.285857 | 5.511696 | 1.096563 | 2.939328 | 2.80606 | -1.29595 |
| 2.025495 | -1.30431 | NA | NA | 5.111679 | 5.114657 | NA | 2.747315 | 2.174989 | 1.059197 |
| 4.100374 | -1.82686 | NA | NA | 3.419607 | 3.312157 | 1.494372 | 3.153652 | -0.81352 | 1.324433 |
| NA | -1.2801 | NA | NA | 4.911226 | 4.662537 | 1.54916 | 2.302746 | 1.51135 | NA |
| 0.324655 | 2.51683 | -1.94874 | NA | 5.218975 | 5.218975 | -0.72473 | 3.824409 | 3.666889 | -4.17225 |
| 0.226827 | -3.79856 | NA | NA | 5.821456 | 5.787208 | 0.680976 | 2.347221 | 1.51321 | -1.56392 |
| 3.300162 | 0.349167 | -2.96355 | NA | 5.238793 | 4.952131 | 2.961319 | 3.156224 | 5.240533 | -0.24643 |
| 1.208173 | 0.002246 | NA | NA | 5.63351 | 5.62802 | 0.24575 | 4.125788 | 3.719665 | -2.4866 |
| 2.457572 | -0.88341 | NA | 2.827324 | 4.574921 | 4.259398 | 1.813334 | -3.67316 | 1.927467 | -0.22256 |
| 1.683349 | -3.81938 | -2.11897 | NA | 5.090405 | 5.116216 | -0.50147 | 2.687508 | 2.202634 | -1.3466 |
| 1.676126 | -4.41431 | NA | NA | 4.412676 | 5.222021 | 0.352009 | 1.764457 | 0.192957 | 0.751203 |
| 2.881705 | -0.08192 | -0.63006 | NA | 4.782848 | 5.412217 | 0.748612 | 2.953668 | 3.432527 | -0.57896 |
| 2.308769 | 1.477101 | -0.27164 | NA | 5.146472 | 5.388273 | 0.23781 | 2.54614 | 0.504071 | -1.31261 |
| 0.159994 | -2.50216 | NA | NA | 5.126458 | 4.77851 | -0.87887 | 2.914967 | 1.830378 | -2.05855 |
| 2.586053 | -1.33732 | NA | NA | NA | NA | 0.486525 | NA | NA | 1.134145 |
| 0.200084 | 0.201115 | 0.212269 | 0.212566 | 0.214351 | 0.217104 | 0.219763 | 0.224041 | 0.224969 | 0.227662 |

| Methotrex | NSC-87877 | Temozolon | PD-033299 | Mitomycin | Bicalutami | FR-180204 | WZ3105 | HG-5-88-0 | AZD8055 |
|---|---|---|---|---|---|---|---|---|---|
| 2.284752 | NA | 7.309104 | 5.480756 | NA | NA | NA | NA | 5.323976 | 2.03403 |
| NA | 5.344281 | NA | NA | -0.99998 | NA | 4.753504 | -2.30257 | NA | NA |
| -1.63267 | 3.889081 | 4.739896 | 3.285065 | -2.96603 | 3.617614 | 3.676106 | -0.96485 | NA | -0.24915 |
| 0.306654 | 3.914125 | 4.931195 | 4.078938 | -0.2558 | 4.984005 | 5.793705 | -3.61515 | 3.447998 | -1.40454 |
| -0.04077 | 5.459469 | 6.5492 | 4.608763 | -0.57493 | 5.319904 | 5.529537 | -2.3017 | 4.463998 | 0.636028 |
| 1.434191 | 5.765304 | 6.341052 | 2.125334 | 1.154755 | 5.241743 | 5.762941 | -0.73901 | 5.232121 | 0.54702 |
| -0.17385 | 5.586051 | 5.993915 | 4.188253 | -0.14411 | 4.487337 | 5.775328 | -2.06106 | NA | -2.06576 |
| 3.056676 | 7.488978 | 7.878096 | 4.399261 | 2.654632 | 6.779484 | 7.065682 | 1.036819 | 6.742257 | 0.818342 |
| -0.27324 | 4.436442 | 5.453596 | 1.881867 | 0.402503 | 4.357172 | 4.743262 | -0.35185 | 4.357172 | -1.45924 |
| 1.945574 | 6.490259 | 6.930585 | 4.484216 | 1.586521 | 5.026686 | 5.7153 | -0.16091 | 5.826963 | 0.056672 |
| 2.044191 | 6.094744 | 6.627371 | 0.187503 | 2.050098 | 4.061591 | 5.765696 | -0.51578 | NA | -1.25367 |
| NA | 5.526472 | 4.857846 | NA | -0.66056 | 4.417839 | 6.249193 | -0.41802 | NA | NA |
| 1.672719 | 5.899666 | 6.614242 | 3.289634 | -0.36232 | 4.054 | 6.507319 | 1.817177 | 2.788709 | 0.257422 |
| 1.633584 | 6.146734 | 6.000286 | NA | 0.801457 | 4.810572 | 6.321762 | -0.59077 | NA | NA |
| NA | 5.370852 | NA | NA | 2.152897 | NA | 4.744038 | 2.980429 | NA | NA |
| -0.20442 | 4.541765 | 5.73113 | NA | -0.62175 | 4.059476 | 6.084106 | -0.6739 | 5.633429 | -0.16691 |
| 0.861523 | 4.092951 | 5.206669 | 1.849501 | -0.23608 | 4.656098 | 4.653694 | 0.24038 | NA | -0.77246 |
| 1.201637 | 3.942307 | 6.145901 | 3.26809 | -0.49267 | 5.047289 | 5.218955 | -0.95571 | 5.021629 | 1.34343 |
| NA | 4.363902 | NA | NA | -2.82526 | NA | 5.15138 | -0.6512 | NA | NA |
| 1.047754 | 4.895969 | 5.257403 | 0.398061 | -3.76768 | 4.56848 | 5.695218 | 2.528574 | NA | -0.72112 |
| -2.75464 | 3.522988 | NA | 2.24936 | -0.63979 | NA | 5.018204 | -1.09589 | 3.261301 | 0.273267 |
| 1.96763 | NA | 6.58615 | 4.94928 | 0.938724 | 4.969892 | 4.800626 | NA | 5.615143 | 2.293678 |
| 1.047751 | 4.772583 | 6.253399 | 3.940825 | 0.874882 | 4.714557 | 4.627764 | -0.83652 | 3.872247 | 0.463185 |
| -0.11872 | 5.636889 | 6.375606 | 4.278308 | -1.05304 | 5.418992 | 6.166347 | 0.519209 | 5.419375 | 0.084981 |
| -0.4262 | 1.778179 | 5.315239 | 0.33508 | -3.11384 | 3.642996 | 5.118979 | 1.543953 | NA | -1.57815 |
| 1.41465 | 4.715836 | 6.227005 | 2.768309 | 0.514634 | 3.932742 | 6.033484 | -1.0882 | 4.302881 | 1.381865 |
| NA | 2.614183 | 5.012518 | 1.437114 | -1.99549 | 4.334062 | 4.313825 | -1.70275 | NA | -0.1729 |
| 0.464364 | 5.639004 | 5.730452 | 4.155152 | -0.37965 | 4.057129 | 5.373552 | -1.62027 | 3.836154 | -1.04787 |
| 0.768507 | 5.173178 | 5.570383 | 3.672414 | -1.06322 | 4.623169 | 5.252213 | -1.56548 | 4.141987 | 1.034457 |
| 0.043792 | 5.950293 | 6.431368 | 4.42737 | 0.544436 | 4.631802 | 4.763617 | 1.855322 | NA | 1.014922 |
| NA | 5.465653 | 6.083182 | 2.921748 | 0.812804 | 4.692372 | 5.65828 | -0.61186 | NA | -0.38598 |
| 0.184527 | 5.604159 | 6.136512 | 4.010754 | -0.06234 | 4.464695 | 5.633746 | -1.24515 | NA | 0.99153 |
| -0.87673 | 5.349304 | 5.174188 | NA | -0.15427 | 3.989716 | 3.028609 | -2.58087 | NA | NA |
| 1.172321 | 5.5566 | 5.925798 | -0.0375 | 1.882155 | 4.221995 | 5.879915 | 3.058632 | 4.128592 | -0.45494 |
| 0.008914 | 4.258518 | 6.695671 | 0.563056 | 0.222682 | 4.699309 | 5.277314 | -1.34675 | NA | -0.1367 |
| -0.75091 | 5.700967 | 5.900039 | 2.454971 | -2.61249 | 4.637401 | 4.87216 | 2.869248 | 5.11571 | 0.312584 |
| 0.659132 | 6.079798 | 5.66016 | 3.470862 | 0.648636 | 4.385867 | 6.026169 | 2.465856 | NA | 0.25499 |
| -0.55783 | 5.009328 | 4.605749 | 3.150757 | 1.073797 | 4.032192 | 4.931318 | -2.47412 | NA | 0.880263 |
| 1.075981 | 4.882755 | 5.910104 | 2.61392 | -0.02317 | 4.135277 | 5.52331 | 2.488858 | 3.790833 | -0.67203 |
| 1.505908 | 3.659879 | 5.599893 | 1.732396 | 1.442508 | 4.116247 | 5.75948 | 0.265733 | NA | 0.052923 |
| 1.549797 | 5.931194 | 6.109092 | 2.165124 | -0.03205 | 4.974732 | 6.072383 | 2.783855 | 5.162744 | 0.704375 |
| 0.244426 | 5.406174 | 6.288177 | 3.600866 | -0.60003 | 5.227128 | 5.554408 | -1.33602 | 4.059077 | 0.110848 |
| 1.089929 | 5.420635 | 4.542579 | 0.736405 | 1.053544 | 4.777723 | 5.156306 | -0.78735 | NA | 0.666401 |
| 1.814491 | NA | 5.062426 | 4.808468 | NA | 4.156728 | NA | NA | NA | 1.471571 |
| 0.231789 | 0.232865 | 0.235166 | 0.237282 | 0.243345 | 0.250988 | 0.255111 | 0.255654 | 0.266665 | 0.274287 |

| Bleomycin | Sunitinib | Vinblastine | Erlotinib | UNC0638 | Z-LLNle-CH | Tubastatin | Zibotentan | Y-39983 | SGC0946 |
|---|---|---|---|---|---|---|---|---|---|
| NA | NA | -2.24725 | NA | NA | NA | NA | NA | NA | 3.21476 |
| 0.769875 | 2.283829 | NA | 3.130688 | 2.298006 | -0.70534 | 3.185976 | 5.73239 | 2.831123 | NA |
| 0.3717 | 1.119297 | -2.73519 | 2.574158 | 2.302143 | -0.16381 | 3.026766 | 4.889398 | 3.108626 | 1.157914 |
| 4.233593 | 3.802692 | -3.43696 | 1.420509 | 3.34356 | 1.126975 | 4.374183 | 5.783027 | 5.787427 | 0.828277 |
| 3.868778 | 5.111926 | -1.3089 | 3.35383 | 2.225149 | 1.227148 | 4.122428 | 5.05111 | 5.418727 | 2.363556 |
| 4.565741 | NA | -2.5992 | NA | 3.025252 | NA | 5.141228 | 5.916319 | 5.655247 | 2.223315 |
| -0.5925 | NA | -1.63691 | NA | 2.25796 | NA | 4.296697 | 5.377114 | 5.315948 | 1.85901 |
| 7.399005 | NA | -0.64984 | NA | 3.692354 | NA | 7.204119 | 7.231458 | 6.087685 | 3.160459 |
| 5.15896 | 1.867899 | -4.01776 | 2.653534 | 1.877952 | -0.04264 | 3.175661 | 5.080703 | 3.134453 | 1.286156 |
| 5.207652 | NA | -3.35705 | NA | 3.708755 | NA | 5.220067 | 6.52885 | 6.507722 | 2.836241 |
| 4.945506 | NA | -3.81644 | NA | 2.478636 | NA | 3.81173 | 5.917295 | 6.708914 | 2.789897 |
| 2.339499 | NA | NA | NA | 4.577693 | NA | 4.972111 | 6.271006 | 5.467681 | 2.475161 |
| 3.208609 | NA | -3.7861 | NA | 5.091983 | NA | 6.438162 | 6.519687 | 6.33924 | 0.429154 |
| -1.92782 | NA | -5.83265 | NA | 3.299791 | NA | 3.389842 | 5.480971 | 4.453171 | 2.369477 |
| 4.708179 | NA | NA | 3.447573 | 3.697543 | NA | 2.904939 | 5.650784 | 5.749968 | NA |
| 3.501145 | NA | -3.03284 | NA | 3.389973 | NA | 6.016615 | 6.474698 | 5.801502 | 2.637697 |
| -1.52022 | NA | -6.3479 | NA | 4.617721 | NA | 4.185033 | 5.504046 | 5.300528 | 1.872893 |
| -1.47818 | NA | -4.03658 | NA | 2.860761 | NA | 3.580825 | 5.801099 | 4.535719 | 2.024811 |
| 1.550322 | 4.460106 | NA | NA | 2.920659 | 1.963196 | 3.688605 | 5.374736 | 3.726737 | NA |
| 0.255555 | NA | -3.53983 | NA | 6.017812 | NA | 5.693432 | 5.676827 | 5.695218 | 1.155008 |
| 0.94687 | NA | -5.84559 | NA | 3.267563 | NA | 4.785906 | 5.445772 | 3.555993 | NA |
| 3.352587 | NA | -0.65585 | NA | 1.749356 | NA | 4.095757 | 5.541955 | 6.290388 | 0.960738 |
| 6.664339 | 4.73835 | -2.87962 | 3.08147 | 1.960071 | 3.438236 | 4.081679 | 5.580102 | 5.004309 | 2.180548 |
| 0.819087 | NA | -3.62577 | NA | 6.144179 | NA | 5.919322 | 6.19679 | 4.854689 | 2.091429 |
| -2.87135 | NA | -4.6067 | NA | 5.41532 | NA | 5.118957 | 5.118979 | 5.118874 | 1.362736 |
| 4.030649 | NA | -3.00795 | NA | 3.573245 | NA | 5.409495 | 6.029703 | 5.987597 | 2.219717 |
| -1.48852 | NA | NA | NA | 2.653678 | NA | 2.527322 | 5.173898 | 4.118968 | 1.560589 |
| -0.14984 | NA | -2.87799 | NA | 3.232354 | NA | 5.284812 | 5.960145 | 5.380716 | 2.435718 |
| 3.684741 | NA | -4.13384 | NA | 2.325824 | NA | 4.954097 | 5.245469 | 4.601623 | 1.173017 |
| 3.834665 | NA | -2.83988 | NA | 2.594325 | NA | 4.048444 | 6.15373 | 5.786651 | 2.112926 |
| 4.769807 | NA | NA | NA | 4.202317 | NA | 5.467457 | 5.766371 | 4.926852 | 1.987551 |
| 2.468775 | NA | -3.45013 | NA | 2.407146 | NA | 2.58672 | 5.935437 | 5.935803 | 2.113935 |
| 4.05367 | NA | -3.71949 | NA | 1.937256 | NA | 3.874243 | 4.432563 | 2.983704 | 1.79213 |
| 5.497327 | NA | -4.82535 | NA | 6.485417 | NA | 5.866081 | 5.885901 | 5.125727 | 2.026948 |
| 2.749578 | NA | -3.88922 | NA | 5.425617 | NA | 5.660966 | 6.035158 | 4.32037 | 1.991393 |
| -1.78645 | NA | -3.41006 | NA | 3.462903 | NA | 4.120198 | 5.160848 | 5.590473 | 2.108744 |
| 3.301251 | NA | -3.68028 | NA | 3.786022 | NA | 5.483973 | 6.143297 | 4.633531 | 2.440176 |
| 3.258369 | 2.695354 | -5.10695 | 3.164922 | 3.867713 | -0.17869 | 4.758318 | 5.393478 | 2.940715 | 1.688181 |
| 0.540316 | NA | -5.57278 | NA | 4.443325 | NA | 5.68027 | 5.722527 | 5.337279 | 1.907449 |
| 1.607026 | NA | -4.66487 | NA | 3.519089 | NA | 4.567164 | 5.826116 | 3.144162 | 2.342216 |
| -1.23755 | NA | -4.40145 | NA | 2.069495 | NA | 5.177177 | 6.008373 | 4.281784 | 2.235395 |
| -2.74336 | NA | -2.61301 | NA | 2.931718 | NA | 5.574008 | 5.886541 | 5.881717 | 1.501017 |
| -0.7747 | NA | -3.64044 | NA | 3.598916 | NA | 5.257822 | 5.767868 | 5.516398 | 0.829684 |
| NA | NA | -3.26229 | NA | NA | NA | NA | NA | NA | 2.526132 |
| 0.283994 | 0.287637 | 0.293046 | 0.294356 | 0.29617 | 0.298925 | 0.309031 | 0.309726 | 0.312274 | 0.319643 |

| SB-715992 | Temsirolim | FMK | PIK-93 | CGP-60474 | TL-2-105 | GW 44175 | AZD6244 | QS11 | RO-3306 |
|---|---|---|---|---|---|---|---|---|---|
| NA | -0.66333 | NA | NA | NA | NA | 1.257325 | 5.292161 | NA | 4.66422 |
| -3.61751 | NA | 5.756862 | -0.44521 | -3.12006 | 3.308421 | NA | NA | 3.243556 | NA |
| -3.17974 | -3.03609 | 4.077583 | 1.962386 | -2.45457 | 2.47762 | 2.585756 | -2.90618 | 0.514752 | 2.937764 |
| -1.29728 | -1.95486 | 5.766289 | 4.343402 | -2.05996 | 5.120244 | 0.724817 | 3.8571 | 2.138315 | 3.77723 |
| -2.37114 | 0.33945 | 5.579353 | 2.188729 | -1.65413 | 4.467157 | 4.003831 | 4.487842 | 2.932561 | 4.760393 |
| 0.936572 | -0.63993 | 5.859264 | 4.783856 | NA | 3.310774 | 3.368876 | 1.862607 | 5.181658 | 2.642548 |
| -1.06496 | -3.5052 | 5.143411 | 3.628162 | NA | 3.264108 | 3.519332 | 3.308619 | 5.045192 | 4.403747 |
| 2.639187 | -2.01249 | 7.649954 | 3.658328 | NA | 3.807214 | 5.282415 | 5.081798 | 4.506229 | 5.06763 |
| -2.67998 | -2.40115 | 5.109573 | 1.188466 | -2.1181 | 3.128565 | 2.643594 | 3.239885 | 3.089099 | 1.137279 |
| 0.963123 | -3.90115 | 5.764671 | 3.636544 | NA | 2.831046 | 1.801538 | 3.793812 | 4.717073 | 3.056537 |
| -1.65032 | -2.71689 | 5.392066 | -0.24522 | NA | 3.104154 | 1.770171 | 1.043118 | 5.613551 | 4.095113 |
| -1.84917 | NA | 5.99649 | 2.951134 | NA | 4.50284 | NA | 4.13688 | 1.779604 | NA |
| -1.23971 | -3.63292 | 6.543467 | 3.831286 | NA | 5.874036 | 1.735883 | 2.922617 | 5.653058 | 4.770312 |
| 0.03412 | -3.14757 | 6.635363 | 1.593651 | NA | 2.307464 | 2.181151 | 4.132338 | 5.794959 | NA |
| 0.475297 | NA | 5.642663 | 4.296697 | NA | 3.953506 | NA | NA | 5.080331 | NA |
| -1.56667 | 0.182921 | 5.98733 | 2.528927 | NA | 4.08115 | 2.555427 | 2.914169 | 4.964009 | 4.821509 |
| 0.198269 | -2.91094 | 5.412531 | 0.692643 | NA | 4.751168 | 2.584899 | 2.931737 | 4.673956 | 3.681359 |
| -1.5545 | -1.37664 | 4.892588 | 2.677492 | NA | 4.252357 | 3.400707 | 4.11747 | 5.099689 | 2.808236 |
| -0.46846 | NA | 5.376397 | 1.598525 | -2.19021 | 4.236555 | NA | NA | 3.095494 | NA |
| 0.48735 | -1.17279 | 5.695218 | 5.694704 | NA | 5.023082 | 1.606256 | 3.773306 | 1.257217 | 1.986008 |
| -1.66504 | -2.71503 | 5.434313 | 1.998905 | NA | 3.445965 | -0.56293 | NA | 2.123459 | 0.815109 |
| -0.76369 | 1.451619 | 5.397585 | 1.43406 | NA | 5.143514 | 1.501163 | 4.377894 | 4.877476 | 5.120292 |
| -0.86226 | -1.26157 | 5.99609 | 1.787953 | 1.000945 | 3.445317 | 2.608698 | 3.977048 | 4.369619 | 4.596781 |
| -3.61195 | -1.87176 | 5.560272 | 2.897042 | NA | 4.124367 | 3.703928 | 3.060544 | 5.491145 | 4.036209 |
| -0.17221 | -1.66823 | 4.893138 | 4.980043 | NA | 3.54665 | 2.816394 | 2.006685 | 0.911798 | 3.575257 |
| -5.19495 | -0.54965 | 4.424046 | 3.418082 | NA | 4.307147 | 2.722355 | 2.533938 | 2.60548 | 3.288238 |
| -3.3637 | NA | 5.325563 | 2.980525 | NA | 4.368901 | NA | 2.322328 | 2.626224 | 3.040724 |
| -0.88035 | 0.23388 | 5.58616 | 2.892201 | NA | 0.514826 | 3.804899 | 4.05995 | 3.066576 | 4.336577 |
| -4.8204 | -2.0215 | 5.093564 | 2.655555 | NA | 4.398449 | 1.331439 | 3.714446 | 4.149549 | 0.777502 |
| 0.001049 | 0.912745 | 5.786431 | 3.544806 | NA | 4.852904 | 3.271243 | 2.031187 | 4.946647 | 4.762758 |
| 0.212384 | NA | 5.734408 | 5.152336 | NA | 4.466841 | NA | 4.016343 | 4.308451 | 3.711874 |
| -2.95772 | -0.70349 | 5.341831 | 0.925279 | NA | 1.805872 | 2.489243 | 4.193024 | 2.865895 | 3.984089 |
| -2.29187 | NA | 5.516107 | 1.886694 | NA | 4.30947 | 3.12342 | 1.081176 | 4.886839 | NA |
| 0.467055 | -3.56406 | 5.884673 | 5.240062 | NA | 5.206092 | 2.081153 | 0.643169 | 5.085898 | 0.939966 |
| -2.03659 | -1.16763 | 5.268422 | 1.926467 | NA | 5.53656 | 3.666813 | 0.864448 | 4.243863 | 4.99019 |
| -0.55469 | -2.8684 | 5.769544 | 2.022096 | NA | 3.484741 | 3.076103 | 4.088137 | 1.412373 | 3.299456 |
| -0.02726 | -2.31174 | 6.278259 | 2.860575 | NA | 5.421339 | 1.560216 | 1.809146 | 5.372376 | 4.007945 |
| -4.5666 | -1.01961 | 5.022202 | 1.54371 | -1.77286 | 4.004764 | -2.38715 | 2.947685 | 3.690263 | 3.773231 |
| -2.30655 | -5.24035 | 5.710515 | 3.250471 | NA | 4.605581 | 3.062644 | 1.281802 | 2.945795 | 1.714658 |
| 0.192181 | -0.80127 | 5.388855 | 1.878782 | NA | 4.749104 | 3.737796 | 2.606744 | 5.519887 | 3.63867 |
| -0.1037 | -1.27784 | 5.810942 | 4.599461 | NA | 5.282279 | 1.815998 | 2.204233 | 3.522923 | 2.286979 |
| -1.1969 | -2.85984 | 5.851136 | 2.720755 | NA | 2.523865 | 1.5763 | 2.652523 | 5.385756 | 3.613729 |
| -1.58139 | -2.0546 | 5.63437 | 5.786472 | NA | 4.079344 | 3.219671 | 1.086891 | 3.187685 | 3.903866 |
| NA | -2.91974 | NA | NA | NA | NA | 1.119284 | 1.558716 | NA | 4.899767 |
| 0.319771 | 0.321666 | 0.322501 | 0.323288 | 0.324364 | 0.338304 | 0.341142 | 0.351239 | 0.35135 | 0.352391 |

| NPK76-II-7. | Bexarotene | VNLG/124 | BMS-53692 | MP470 | UNC1215 | AC220 | Etoposide | SB52334 | AKT inhibit |
|---|---|---|---|---|---|---|---|---|---|
| NA | NA | NA | NA | NA | 3.907907 | NA | NA | NA | NA |
| -0.70176 | 4.488646 | 4.196445 | 3.667138 | 3.628999 | NA | 2.050816 | -0.65514 | 5.192018 | 2.608522 |
| 2.327797 | 3.454544 | 3.531925 | 1.330491 | 3.700749 | 1.612709 | 1.808366 | -2.02607 | 3.746326 | 1.524838 |
| 5.124161 | 3.747 | 4.277858 | 2.868882 | 2.772455 | 1.826634 | -1.52938 | 2.533787 | 5.792229 | 1.318296 |
| 5.4137 | 5.390125 | 4.030197 | 4.005398 | -0.03798 | 1.706959 | 2.388327 | 4.882789 | 4.646549 | 1.851721 |
| 4.758069 | 2.671244 | 4.558418 | NA | 4.178006 | 2.856245 | 1.307684 | 4.678824 | 5.328949 | 2.745517 |
| 4.3768 | 3.594317 | 2.449036 | NA | 1.009318 | 2.702956 | 2.126099 | 3.64147 | 3.845517 | 3.763724 |
| 6.449478 | 6.159034 | 6.310809 | NA | 4.263585 | 4.476899 | 3.507311 | 5.782212 | 2.448345 | 3.999166 |
| 1.252562 | 4.157681 | 3.051371 | 2.488318 | 3.30345 | 2.054587 | 1.719129 | 3.913562 | 2.23811 | 2.466775 |
| 5.310451 | NA | 5.006886 | NA | 3.253413 | 3.529388 | 1.380642 | 6.205802 | 3.25459 | 1.195471 |
| 4.493389 | 4.954932 | 5.322316 | NA | 4.654486 | 3.266929 | 2.070497 | 1.938669 | 3.996267 | -0.2673 |
| 4.437333 | 1.434364 | 3.565927 | NA | 3.63341 | 3.159809 | 3.070224 | 1.57481 | 4.325648 | 2.364194 |
| 5.283456 | 4.311021 | 4.70592 | NA | 1.009223 | 3.27546 | 3.571451 | 4.580718 | 6.543057 | 0.506107 |
| 3.430681 | 2.650719 | 5.010128 | NA | 6.643099 | 3.443147 | 2.28761 | -0.01662 | 2.715647 | -0.76741 |
| 4.461738 | 4.833812 | 4.387865 | NA | 4.986151 | NA | 2.776343 | 3.431983 | 5.245391 | 3.678641 |
| 3.989146 | 2.897023 | 5.130921 | NA | 1.519412 | 3.330844 | 2.297813 | 5.259002 | 4.489401 | 1.332154 |
| 3.407114 | 3.080604 | 4.274017 | NA | 3.439654 | 2.559909 | 2.68121 | 2.679606 | 3.92865 | -0.60922 |
| 1.143529 | 4.523244 | 4.342779 | NA | 4.937793 | 2.744704 | 2.332469 | 0.930839 | 3.286912 | 0.73792 |
| 1.60123 | 4.443156 | 3.143874 | 3.720631 | 2.833735 | NA | 1.961378 | -0.36878 | 3.807174 | 1.550963 |
| 5.011624 | 2.63818 | 4.089716 | NA | 0.168494 | 2.543474 | 2.713604 | -0.03416 | 5.695218 | 2.162095 |
| 2.987576 | 2.722074 | 2.635926 | NA | 3.936964 | NA | 1.386408 | 1.054299 | 3.338529 | 0.062734 |
| 3.252492 | 3.9323 | 5.004187 | NA | NA | 3.101669 | NA | NA | 2.206094 | NA |
| 4.202281 | 5.085416 | 4.639129 | 4.254542 | 2.483447 | 2.873695 | 2.277451 | 4.347218 | 2.535702 | 2.245198 |
| 4.711261 | 4.79261 | 4.482732 | NA | 3.887654 | 3.11679 | 2.516425 | 1.730789 | 5.79629 | 1.313557 |
| 4.449548 | 2.768215 | 3.680474 | NA | 3.501542 | 1.380499 | 1.760656 | -0.30799 | 4.917831 | 0.986515 |
| 5.358144 | 4.996604 | 4.39519 | NA | 4.712744 | 2.902313 | 2.20536 | 2.161363 | 5.910914 | 1.926128 |
| 0.522357 | 3.311632 | 3.42588 | NA | 4.069147 | 2.230432 | 2.352552 | 1.659763 | 4.704466 | NA |
| 3.366994 | 4.519504 | 4.688566 | NA | 4.276808 | 3.128772 | 0.582989 | 1.215706 | 5.160104 | 1.014988 |
| 2.14381 | 3.523214 | 3.539835 | NA | 0.263571 | 2.232916 | 1.932594 | 0.520661 | 1.619906 | 1.012194 |
| 4.299945 | 3.936857 | 2.986982 | NA | 5.52113 | 3.036622 | 3.203947 | 2.109685 | 6.181127 | 2.763727 |
| 3.223043 | 2.340258 | 4.420544 | NA | 5.503802 | 2.439353 | 2.293161 | 2.524599 | 5.177773 | 2.503158 |
| 1.763763 | 2.965251 | 4.573815 | NA | 0.396503 | 2.807082 | 1.925989 | 5.199934 | 3.265284 | 1.553293 |
| 2.580521 | 3.812877 | 4.218079 | NA | 4.608368 | 2.27002 | 2.301204 | 4.358723 | 3.287338 | -0.26116 |
| 5.193516 | 3.61119 | 4.525828 | NA | 5.608435 | 2.707827 | 2.91639 | 5.665262 | 5.400838 | 2.92569 |
| 1.200759 | 1.541099 | 5.03725 | NA | 5.568585 | 3.293541 | 3.407465 | 4.388657 | 4.845832 | 2.503783 |
| 3.030009 | 2.591432 | 3.839372 | NA | 4.437244 | 2.813124 | 2.597891 | -0.136 | 4.797695 | 2.865243 |
| 3.634611 | 4.409371 | 4.760564 | NA | 4.668336 | 2.946333 | 3.323181 | 2.403427 | 3.777672 | 2.622249 |
| 1.243481 | 3.494196 | 3.98304 | 2.623426 | 3.507625 | 2.058853 | 1.946909 | 3.807016 | 3.825799 | 3.41178 |
| 4.88565 | 3.261584 | 4.064054 | NA | 3.611549 | 2.50867 | 2.798162 | 1.633112 | 5.518901 | 2.15094 |
| 1.823164 | 3.921264 | 4.140534 | NA | 4.726416 | 2.636325 | 2.801162 | 2.189807 | 4.617315 | 3.364304 |
| 4.074562 | 3.556244 | 4.709289 | NA | 5.555085 | 3.050737 | 3.101858 | 4.902427 | 5.815806 | 3.393058 |
| 3.370652 | 3.926388 | 4.689007 | NA | 4.692358 | 2.924543 | 2.169093 | 2.346022 | 4.990642 | 1.634026 |
| 5.126458 | 3.241756 | 3.455639 | NA | 3.288837 | 2.67875 | 2.790505 | 1.795578 | 5.750365 | 3.238039 |
| NA | NA | NA | NA | NA | 3.148795 | NA | NA | NA | NA |
| 0.355275 | 0.359192 | 0.361341 | 0.368896 | 0.369113 | 0.371821 | 0.372926 | 0.381455 | 0.382058 | 0.385268 |

| NVP-BHG7 | CCT00709: | TGX221 | Salubrinal | Elesclomol | KIN001-05! | FTI-277 | Shikonin | Nutlin-3a | KIN001-26( |
|---|---|---|---|---|---|---|---|---|---|
| NA | 5.261029 | NA | NA | -4.93658 | NA | NA | NA | 5.66956 | NA |
| 2.029026 | NA | 1.572551 | 2.660693 | NA | 3.962879 | 2.186058 | -0.97112 | NA | 4.457526 |
| -0.58246 | 3.372004 | 3.861403 | 2.387091 | -2.21988 | 3.135032 | 0.658686 | -1.46186 | 1.739453 | 3.709197 |
| 5.121094 | 4.622066 | 3.111687 | 3.668324 | -3.8189 | 5.053755 | 2.26065 | -0.99509 | 4.681817 | 5.744255 |
| 3.712697 | 4.808456 | 5.03812 | 2.652784 | -1.92126 | 4.245144 | 2.361598 | -0.74619 | 4.062102 | 5.60562 |
| 5.382695 | 4.375996 | NA | NA | -2.4285 | 4.881402 | 2.70901 | 1.914123 | 4.9249 | 5.082073 |
| 4.13208 | 3.864668 | 4.394879 | NA | -0.83352 | 4.76167 | 2.848087 | 0.818452 | 3.630693 | 4.216587 |
| 6.884215 | 5.100963 | NA | NA | -3.79966 | 5.498508 | 4.69415 | 2.220945 | 5.179769 | 7.698295 |
| 2.687177 | 4.000911 | 1.753487 | 2.542246 | -3.29635 | 3.862979 | 2.119668 | -0.88004 | 4.189357 | 0.226695 |
| 4.374156 | 4.541219 | NA | NA | -3.8748 | 4.76211 | 3.535943 | 0.601555 | 5.029112 | 5.153451 |
| 3.541403 | 2.985812 | NA | NA | -4.09404 | 5.777519 | 3.003991 | -0.86056 | 4.103107 | 5.523094 |
| 4.509324 | 3.376344 | NA | NA | NA | 5.280409 | 1.985355 | 0.440224 | NA | 5.404479 |
| 5.553706 | 4.895409 | NA | NA | -1.87579 | 5.65082 | 3.02923 | 0.325221 | 5.591296 | 5.725365 |
| 2.839884 | 5.401506 | NA | NA | -2.76236 | 5.892941 | 1.38952 | -1.18172 | NA | 4.378436 |
| 3.284368 | NA | 5.056769 | 5.114508 | NA | 3.109213 | 2.668952 | 1.454986 | NA | 5.746471 |
| 3.162889 | 3.451142 | NA | NA | -1.0616 | 3.342492 | 2.398882 | 4.302222 | 4.023235 | 5.451715 |
| 3.20922 | 3.107172 | NA | NA | -4.18343 | 4.367275 | 1.13229 | 0.163481 | 3.976236 | 5.103702 |
| 3.831386 | 4.538729 | NA | NA | -1.40217 | 2.95872 | 1.703352 | 0.002422 | 4.916019 | 4.522003 |
| 4.298997 | NA | 4.164803 | 3.266688 | NA | 4.142362 | 2.275191 | -0.36696 | NA | 4.957289 |
| 4.584757 | 1.284488 | NA | NA | -3.99368 | 5.016867 | 2.713541 | -0.29822 | 1.72173 | 5.694759 |
| 4.046381 | 0.933741 | NA | NA | -4.39345 | 4.555363 | 1.253322 | -0.31001 | 4.738715 | 4.688951 |
| 4.133831 | 5.405674 | NA | NA | -2.12326 | 3.835332 | NA | NA | 5.58674 | 5.693196 |
| 3.98226 | 3.335818 | 5.192476 | 4.264305 | -3.98457 | 4.742564 | 2.723399 | -1.4552 | 4.808941 | 4.142448 |
| 4.43838 | 3.850709 | NA | NA | -3.2287 | 4.438205 | 2.351204 | 0.779973 | 5.34652 | 6.085715 |
| 4.449146 | 3.985403 | NA | NA | -6.25783 | 4.448659 | 2.036973 | -2.48943 | 1.971822 | 5.118954 |
| 5.382223 | 4.333461 | NA | NA | 1.337266 | 4.83727 | 2.570699 | -0.51343 | 2.64881 | 6.040302 |
| 3.074353 | 4.222042 | NA | NA | NA | 3.113953 | 1.954384 | 0.049446 | 3.624556 | 4.319504 |
| 4.164289 | 3.378034 | NA | NA | 1.043775 | 4.395722 | 2.944081 | -0.28589 | 3.627363 | 5.44393 |
| 4.175917 | 3.265218 | NA | NA | -6.51526 | 2.691854 | 2.079484 | -0.01553 | 4.513282 | 4.657807 |
| 5.511166 | 4.912065 | NA | NA | -0.14943 | 4.435224 | 3.16162 | -0.03699 | 5.182024 | 6.165986 |
| 5.019205 | 4.384072 | NA | NA | NA | 4.833598 | 2.312631 | -0.29613 | 3.098957 | 5.780269 |
| 5.263452 | 3.787893 | NA | NA | -3.55693 | 4.62237 | 1.881089 | 0.801748 | 4.973614 | 5.19102 |
| 3.130156 | 3.737656 | NA | NA | -1.99063 | 4.772858 | 2.388631 | 0.648158 | NA | 4.972826 |
| 5.218487 | 4.49653 | NA | NA | -4.27539 | 5.218579 | 2.897271 | 1.097585 | 4.913082 | 5.886459 |
| 4.285696 | 5.333821 | NA | NA | -2.50206 | 2.973241 | 2.929444 | -0.43469 | 4.851803 | 4.063064 |
| 4.115467 | 3.897894 | NA | NA | -1.01624 | 4.42561 | -0.37596 | -0.10578 | 3.448789 | 4.845398 |
| 4.288799 | 4.409235 | NA | NA | -4.50444 | 4.733058 | 1.374744 | 0.496627 | 4.203331 | 5.17324 |
| 2.480735 | 2.561016 | 3.342611 | 3.399407 | -2.96058 | 3.861231 | 2.179127 | 0.162024 | 3.818731 | 5.139346 |
| 3.129448 | 3.822911 | NA | NA | -4.10942 | 3.954098 | 1.144078 | -0.62433 | 3.760119 | 5.620482 |
| 2.556955 | 5.059295 | NA | NA | -4.15703 | 3.339621 | 1.539517 | 0.529752 | 3.600698 | 5.288163 |
| 4.556116 | 4.795227 | NA | NA | -3.89697 | 5.050817 | 2.316632 | 1.526029 | 4.909215 | 5.107634 |
| 4.106121 | 4.644993 | NA | NA | -5.65225 | 3.202167 | 2.924206 | 0.178694 | 4.077041 | 4.900073 |
| 2.83581 | 4.37089 | NA | NA | -3.89679 | 5.087122 | 2.298487 | 0.294562 | 4.410775 | 5.718502 |
| NA | 5.138422 | NA | NA | -4.85816 | NA | NA | NA | 5.50337 | NA |
| 0.386263 | 0.390582 | 0.400391 | 0.404607 | 0.404925 | 0.40812 | 0.409604 | 0.420719 | 0.43724 | 0.439191 |

| | | | | | | | | | |
|---|---|---|---|---|---|---|---|---|---|
| ABT-263 | ZG-10 | Lenalidomi | CP724714 | JQ1 | BMS-7081( | PLX4720 (r | GDC0941 ( | AZ628 | GSK42928( |
| 3.736741 | 2.536418 | 5.7039 | NA | 3.907907 | NA | 6.038634 | NA | NA | NA |
| NA | NA | NA | 4.717331 | NA | 3.431391 | NA | NA | 1.251045 | 4.730134 |
| -0.3582 | NA | 3.453011 | 3.057325 | 1.32682 | 4.212875 | -0.10673 | 2.191147 | -3.59171 | 3.101455 |
| -0.62996 | 0.296913 | 3.899539 | 4.738385 | 2.537122 | 5.124274 | 4.770603 | 0.582808 | 2.518996 | 5.054248 |
| 3.711546 | 1.334246 | 4.489811 | -1.83278 | 1.285683 | 3.553578 | 4.810721 | 0.637291 | 4.655011 | 5.387271 |
| 1.748952 | 2.098555 | 4.311696 | 4.731574 | 2.802999 | 4.457781 | 5.242439 | 3.836475 | NA | 5.850624 |
| -2.9828 | NA | 4.20393 | 4.242353 | 0.719182 | 5.075171 | 4.859221 | 2.595971 | 2.344141 | 5.152487 |
| 2.812405 | 5.500267 | 6.331979 | 4.581197 | 4.083035 | 4.916935 | 6.779484 | 2.560442 | NA | 7.723007 |
| 1.501455 | 0.69809 | 3.723279 | 4.245814 | 2.054036 | 4.439284 | 4.35662 | -0.95701 | 3.433354 | 5.109573 |
| 0.876359 | 3.453158 | 3.909066 | 1.358056 | 3.529388 | 6.043972 | 5.775022 | 1.9163 | NA | 5.787928 |
| 3.998207 | NA | 5.294231 | 4.011174 | 1.486243 | 5.565553 | 5.056766 | -1.11862 | NA | 6.59684 |
| NA | NA | NA | 4.465689 | 3.165482 | 5.654675 | 5.187032 | 2.725953 | NA | 5.30979 |
| 0.66757 | 1.807407 | 5.157172 | 5.687126 | 2.472916 | 5.851293 | 5.625812 | 0.389611 | NA | 6.543467 |
| NA | NA | 5.240451 | 5.309812 | 2.906585 | 5.982602 | 5.721053 | -0.28509 | NA | 6.327846 |
| NA | NA | NA | 3.279656 | NA | 4.940958 | NA | NA | 4.140672 | 5.747595 |
| 2.816226 | 3.261995 | 3.924706 | 2.267182 | 3.330122 | 5.786655 | 4.847308 | 0.641938 | NA | 5.607737 |
| 2.831912 | NA | 2.923454 | 3.327487 | 2.437415 | 4.983142 | 4.650654 | -0.5867 | NA | 5.583172 |
| -0.22845 | 1.940313 | 4.456831 | 4.617184 | 2.698855 | 4.603024 | 4.453763 | 2.267856 | NA | 4.555994 |
| NA | NA | NA | 4.445575 | NA | 4.698191 | NA | NA | 3.76198 | 5.325322 |
| 2.474808 | NA | 4.306668 | 4.608959 | 2.072562 | 5.009414 | 4.884355 | -0.7698 | NA | 5.694053 |
| 1.981488 | 1.089226 | 4.444755 | 1.376192 | NA | 5.161619 | 4.99592 | NA | NA | 5.281371 |
| 2.948336 | 2.364385 | 5.185184 | NA | 2.572672 | NA | 5.674557 | 1.12244 | NA | 3.578112 |
| 3.385941 | 1.059341 | 3.994355 | -0.65612 | 2.873695 | 5.327414 | 4.408168 | 0.516667 | 3.778453 | 6.000631 |
| 1.097944 | 5.069711 | 3.739158 | 5.07138 | 3.06625 | 4.077648 | 5.419375 | 1.412131 | NA | 6.270715 |
| 2.786963 | NA | 3.732684 | 3.991571 | 1.395178 | 4.390961 | 3.853204 | 0.32947 | NA | 4.633517 |
| 2.598219 | 2.060377 | 4.212928 | 2.322885 | 2.314562 | 5.229912 | 3.848858 | 0.850796 | NA | 6.055491 |
| 0.957843 | NA | NA | 4.400471 | 2.249383 | 4.139268 | 4.458524 | 1.904805 | NA | 5.254532 |
| -1.44701 | 0.8786 | 4.782161 | 4.138263 | 2.202587 | 5.410125 | 4.309195 | -0.02787 | NA | 5.065431 |
| 1.165848 | 0.625563 | 4.06707 | 3.468523 | 2.360908 | 4.764401 | 4.663493 | 1.315404 | NA | 5.012599 |
| 1.391036 | NA | 4.755848 | 5.240555 | 2.709757 | 5.509458 | 4.008151 | 1.116339 | NA | 6.033226 |
| 1.170084 | NA | NA | 5.115127 | 2.383394 | 5.112713 | 4.98457 | 1.983145 | NA | 5.577461 |
| 3.310014 | NA | 4.550098 | 1.784899 | 2.805822 | 3.718668 | 5.036222 | 0.066973 | NA | 5.501089 |
| NA | NA | 3.903715 | 4.834446 | 2.421412 | 4.766817 | 4.18406 | 0.830304 | NA | 5.548994 |
| 3.157293 | 1.51376 | 4.467415 | 5.218975 | 2.442934 | 5.2174 | 5.074242 | 0.886926 | NA | 5.579547 |
| 2.802647 | NA | 4.805975 | 4.247101 | 2.010018 | 5.811698 | 4.731131 | 0.51153 | NA | 6.401142 |
| 1.00321 | 0.976056 | 4.518637 | 4.743104 | 2.760001 | 5.144797 | 5.0609 | -0.04827 | NA | 5.941885 |
| 2.438942 | NA | 4.856242 | 5.213792 | 0.649173 | 5.590569 | 4.390585 | 1.389918 | NA | 6.240997 |
| 2.523995 | NA | 3.838311 | 3.751538 | 2.140987 | 4.59736 | 4.683913 | 0.918419 | 1.511605 | 3.449596 |
| 1.14982 | 2.944343 | 3.751108 | 4.754471 | 1.835488 | 4.83775 | 3.774477 | -0.48611 | NA | 5.636745 |
| 0.22996 | NA | 4.272166 | 5.473571 | 2.809466 | 3.908963 | 3.648978 | 0.13232 | NA | 4.104334 |
| 1.626645 | 1.208446 | 4.737316 | 5.288879 | 2.749035 | 3.955445 | 4.285989 | 1.229271 | NA | 5.53055 |
| 0.571156 | 2.224619 | 3.196057 | 4.913912 | 2.075706 | 4.927908 | 5.027678 | 2.754775 | NA | 6.061867 |
| 3.074841 | NA | 3.638746 | 4.249843 | 2.620953 | 4.316316 | 4.963929 | 1.893711 | NA | 3.694939 |
| 4.005554 | NA | 4.543497 | NA | 2.906068 | NA | 4.800859 | 0.837463 | NA | NA |
| 0.443492 | 0.452704 | 0.460105 | 0.468968 | 0.471545 | 0.484368 | 0.487226 | 0.495609 | 0.500663 | 0.502421 |

| STF-62247 | Bicalutami | Parthenolid | PLX4720 | A-770041 | XMD13-2 | XMD8-92 | EHT 1864 | OSI-930 | QL-XII-47 |
|---|---|---|---|---|---|---|---|---|---|
| NA | NA | NA | 6.397047 | NA | NA | 4.919018 | 3.884064 | NA | NA |
| 4.404087 | 2.547852 | 3.450705 | NA | 3.75037 | 2.410155 | NA | NA | 3.361369 | 0.497197 |
| 3.211406 | 1.359969 | 1.263522 | -0.82879 | 2.346526 | 2.327331 | NA | 2.659983 | 3.169406 | 0.101164 |
| 2.824578 | 2.761825 | 3.695274 | 4.872425 | 4.365073 | 2.956711 | 1.645437 | 3.170025 | 3.422082 | 3.059967 |
| 4.705935 | 3.013135 | 4.910317 | 4.407884 | 1.313175 | 2.631952 | 5.366554 | 3.908555 | 3.445541 | 2.041085 |
| 5.339387 | 3.02271 | NA | 4.952882 | NA | 5.145228 | 3.778843 | 4.836505 | 4.683782 | 2.694426 |
| 5.152302 | 2.765501 | 3.902965 | 4.603157 | NA | 3.09006 | NA | 2.53975 | 3.869288 | 0.661739 |
| 6.980813 | 4.660955 | NA | 6.27458 | NA | 6.237196 | 6.779484 | 3.580429 | 4.481285 | 1.729932 |
| 3.583209 | 2.090639 | 1.393362 | 3.701884 | 1.793397 | 3.307754 | 2.461859 | 2.519045 | 3.362401 | 1.269192 |
| 4.452351 | 3.369495 | NA | 4.896998 | NA | 4.610572 | 4.313489 | 2.73373 | 4.26594 | 2.530827 |
| 6.03006 | 3.720552 | NA | 6.015767 | NA | 3.136708 | NA | 3.146433 | 4.664118 | 0.509376 |
| 4.170106 | 1.193971 | NA | NA | NA | 3.836758 | NA | 3.386843 | 3.869287 | 1.355229 |
| 5.781882 | 3.483886 | NA | 5.821083 | NA | 5.619348 | 3.653134 | 5.514207 | 3.206428 | 5.179114 |
| 5.999238 | 3.531319 | NA | NA | NA | 4.892874 | NA | 4.736156 | 5.874878 | -0.45774 |
| 4.41916 | 2.769638 | 4.360326 | NA | NA | 4.873058 | NA | NA | 4.819275 | 1.667683 |
| 4.626097 | 3.532049 | NA | 5.556501 | NA | 4.061357 | 5.445907 | 1.89673 | 4.170232 | 0.380933 |
| 4.978076 | 2.530586 | NA | 4.956707 | NA | 3.527161 | NA | 3.253737 | 3.87932 | 2.173296 |
| 5.181757 | 2.334285 | NA | 5.163732 | NA | 2.616523 | 3.266049 | 4.3568 | 5.194021 | 2.168709 |
| 4.663154 | 2.380827 | 3.923159 | NA | 4.003597 | 2.58319 | NA | NA | 3.117867 | 2.833587 |
| 5.025787 | 1.996173 | NA | 2.651349 | NA | 5.025787 | NA | 2.453324 | 2.981734 | 5.025787 |
| 4.91648 | 2.105332 | NA | 4.499728 | NA | 3.71545 | 3.280698 | 3.455017 | 4.146008 | 1.695869 |
| 2.976872 | 2.790273 | NA | 3.785553 | NA | NA | 5.087666 | 4.622714 | NA | -1.19029 |
| 5.329266 | 3.007436 | 4.588734 | 5.257707 | 2.699687 | 2.709658 | 1.395726 | 2.676416 | 3.287362 | -0.10266 |
| 5.429825 | 3.208567 | NA | 4.43554 | NA | 3.759721 | 4.583927 | 3.61501 | 4.548996 | 0.00109 |
| 4.435858 | 2.08193 | NA | 4.025146 | NA | 3.971622 | NA | 2.974649 | 2.813081 | 2.484041 |
| 5.378402 | 2.489802 | NA | 5.314039 | NA | 2.191016 | 5.181378 | 3.602586 | 5.092307 | -1.36003 |
| 2.724919 | 2.06921 | NA | 3.269433 | NA | 1.973287 | NA | 3.957975 | 3.782555 | 2.709855 |
| 4.582734 | 3.331215 | NA | 5.565194 | NA | 4.347888 | 4.019416 | 3.436916 | 4.823234 | 1.166001 |
| 3.8063 | 2.463749 | NA | 4.731559 | NA | 2.199233 | 3.710992 | 2.645402 | 2.717346 | 1.406114 |
| 4.394409 | 3.111884 | NA | 4.10451 | NA | 4.226668 | NA | 4.510105 | 3.393082 | 3.211278 |
| 5.024447 | 2.792447 | NA | 2.612053 | NA | 4.510584 | NA | 3.624734 | 5.061338 | 2.361017 |
| 4.473592 | 2.010334 | NA | 4.629329 | NA | 2.128697 | NA | 2.225785 | 2.340742 | 0.23306 |
| 3.834253 | 2.364574 | NA | NA | NA | 2.81879 | NA | 3.834305 | 4.579537 | 2.230799 |
| 5.200789 | 2.903521 | NA | 4.581134 | NA | 5.050354 | 3.156761 | 3.851321 | 4.925875 | 3.277316 |
| 5.269962 | 3.384238 | NA | 5.748161 | NA | 4.662144 | NA | 5.133982 | 5.013009 | 2.740189 |
| 5.073456 | 2.96304 | NA | 4.535049 | NA | 4.876847 | 4.39151 | 3.782066 | 4.568893 | 0.994588 |
| 5.632067 | 3.317989 | NA | 5.039287 | NA | 4.893704 | NA | 4.329962 | 4.025479 | 3.885258 |
| 2.964436 | 2.373062 | 2.84956 | 4.631648 | 2.864723 | 0.505555 | NA | 3.392886 | 2.7381 | 1.906359 |
| 4.023003 | 2.813631 | NA | 2.858282 | NA | 4.787218 | 2.522916 | 3.97913 | 3.379624 | 2.407415 |
| 5.2309 | 3.170626 | NA | 3.616509 | NA | 3.911494 | NA | 3.738559 | 4.207668 | 2.347384 |
| 5.269724 | 2.863855 | NA | 4.979141 | NA | 4.003554 | 4.406961 | 4.396638 | 5.527926 | 4.834537 |
| 5.17267 | 3.083289 | NA | 4.875281 | NA | 3.021977 | 3.814914 | 3.801887 | 4.438201 | 0.681711 |
| 4.4562 | 2.758395 | NA | 4.955335 | NA | 4.480436 | NA | 4.368792 | 2.940217 | 1.935472 |
| NA | NA | NA | 5.726514 | NA | NA | NA | 3.728166 | NA | NA |
| 0.511912 | 0.512214 | 0.517737 | 0.518614 | 0.52831 | 0.530868 | 0.533303 | 0.53507 | 0.535274 | 0.536951 |

| Bortezomib | CP466722 | AZD6482 | MK-2206 | LY317615 | WH-4-023 | Masitinib | Vorinostat | AZD6482 | BIRB 0796 |
|---|---|---|---|---|---|---|---|---|---|
| NA | NA | NA | 1.282224 | NA | NA | NA | 3.057939 | NA | 4.326489 |
| -4.89517 | 1.809596 | 3.266918 | NA | 2.120363 | 2.97198 | 3.042514 | NA | NA | NA |
| -6.60958 | 1.831072 | 2.593086 | 2.132672 | 1.546247 | 2.721515 | 2.76691 | -0.27121 | 2.50616 | 3.803144 |
| -5.49881 | 4.382642 | 1.067217 | -1.12298 | 4.762911 | 4.194659 | 5.755504 | -0.35071 | -0.35346 | 5.09832 |
| -3.91153 | 2.696403 | 3.855292 | 1.689402 | 0.774774 | 4.317484 | 2.518304 | 0.770289 | 3.977831 | 5.455853 |
| NA | 2.598053 | 3.405041 | 3.151197 | 4.380253 | NA | 3.635445 | -0.61346 | 3.376359 | 4.884352 |
| -1.5711 | 0.815787 | 3.001049 | 4.212479 | 3.037342 | NA | 3.524046 | 0.245134 | 2.976909 | 5.12877 |
| NA | 2.355445 | 4.790718 | 4.056795 | 1.814685 | NA | 5.869393 | 2.872684 | NA | 5.103329 |
| -5.79111 | 3.058476 | 0.970491 | -2.15263 | 4.250948 | 2.117207 | 2.798484 | 0.404656 | -1.56439 | 2.574152 |
| NA | 2.559375 | 3.501467 | -0.56645 | 2.881569 | NA | 3.123291 | 0.531647 | 2.962684 | 3.530547 |
| NA | 2.137029 | -1.02002 | -1.18987 | 5.422849 | NA | 3.361768 | 1.58096 | 0.062751 | 5.135078 |
| NA | 2.619128 | -0.262 | NA | 3.51177 | NA | 3.75485 | NA | 2.508922 | NA |
| NA | 5.656698 | -1.56232 | -0.29536 | 6.463307 | NA | 6.543467 | 2.270808 | 0.722758 | 4.566562 |
| NA | 1.967771 | 1.456643 | NA | -0.86735 | NA | 3.991385 | 0.37748 | 3.306296 | NA |
| -4.33138 | 2.346386 | 3.521895 | NA | 3.097317 | NA | 3.302417 | NA | NA | NA |
| NA | 2.453153 | 0.99252 | NA | 3.081983 | NA | 3.657788 | 3.274382 | 0.360315 | 4.277054 |
| NA | 3.280163 | 1.154864 | -0.54533 | 4.770307 | NA | 4.059649 | 0.654569 | 1.378839 | 3.489578 |
| NA | 2.533736 | 1.159535 | 4.23485 | 3.314854 | NA | 5.068172 | 2.57675 | 3.215879 | 5.060236 |
| -3.2566 | 3.640095 | 1.217932 | NA | 5.376397 | 3.928534 | 4.190243 | NA | NA | NA |
| NA | 5.133508 | 4.144686 | 0.889458 | 5.654872 | NA | 4.9238 | 1.746986 | 0.948264 | 4.54923 |
| NA | 2.502977 | 2.327714 | -0.97156 | 2.117381 | NA | 3.266141 | 1.055134 | 2.251859 | 2.053797 |
| NA | 1.59116 | NA | 1.695474 | 4.196788 | NA | NA | 1.510364 | 4.932229 | 5.879653 |
| -3.20881 | 2.39283 | 2.415071 | 1.111107 | 1.778612 | 3.995365 | 2.87422 | 1.531145 | 2.182119 | 5.222958 |
| NA | 2.865817 | -0.51512 | 1.507246 | 2.61535 | NA | 4.220723 | 1.92473 | 0.568571 | 5.554381 |
| NA | 2.835293 | 2.552237 | 1.870747 | 4.890529 | NA | 4.262819 | 0.832658 | 0.205237 | 4.407907 |
| NA | 1.855697 | 1.298057 | 2.086217 | 0.673253 | NA | 2.686664 | 3.132985 | 1.271514 | 4.004366 |
| NA | 2.907257 | 2.607442 | 1.888485 | 3.792568 | NA | 2.943107 | NA | 3.901376 | 3.497808 |
| NA | 1.415561 | 1.364304 | 2.290544 | 1.45181 | NA | 3.278451 | 2.019343 | 1.594952 | 4.798581 |
| NA | 1.585841 | 0.289815 | 1.207177 | 3.818043 | NA | 3.027398 | 1.41749 | 1.417825 | 2.424403 |
| NA | 2.513798 | 1.937468 | 3.140281 | 3.45946 | NA | 5.264764 | 2.762906 | 2.139195 | 5.332518 |
| NA | 4.014095 | 2.56943 | 1.73659 | 4.549691 | NA | 4.131686 | NA | 4.165975 | 2.894018 |
| NA | 2.44435 | 1.658741 | 1.406476 | 0.922633 | NA | 3.032298 | 0.880936 | 0.004631 | 4.340727 |
| NA | 2.488727 | 0.165117 | NA | 3.669367 | NA | 3.15478 | 1.111738 | 1.175637 | NA |
| NA | 5.665262 | 3.402159 | 0.451733 | 5.454497 | NA | 5.888405 | 1.804904 | 1.270688 | 4.997014 |
| NA | 2.132497 | 2.881527 | 2.986587 | 1.887349 | NA | 3.825475 | 1.392027 | 3.43441 | 5.79429 |
| NA | 2.399835 | 1.852727 | -0.35733 | 3.338874 | NA | 5.586053 | -0.20738 | 3.250467 | 5.248738 |
| NA | 4.340351 | 1.472255 | 1.730667 | 5.01076 | NA | 5.563478 | 1.50696 | 1.195866 | 3.470843 |
| -5.54907 | 1.669206 | 3.723655 | 1.356198 | 3.32506 | 2.895908 | 3.885358 | -2.07454 | 3.657103 | 4.484989 |
| NA | 2.722144 | 3.842583 | 0.229609 | 3.337098 | NA | 5.224575 | 0.547236 | 2.10474 | 3.803429 |
| NA | 2.263693 | 3.47415 | 2.257674 | 5.594773 | NA | 6.090088 | 1.738239 | 2.124705 | 4.781338 |
| NA | 4.741604 | 3.138918 | 0.75138 | 3.327535 | NA | 4.681056 | 2.393082 | 2.777563 | 3.953325 |
| NA | 2.692994 | 1.67077 | 1.291771 | 1.803478 | NA | 3.561842 | 1.372358 | 2.654405 | 3.341751 |
| NA | 2.761228 | 2.77772 | 3.909751 | 5.775294 | NA | 3.636406 | 1.774272 | 2.541677 | 4.191653 |
| NA | NA | NA | 2.783834 | NA | NA | NA | 1.749984 | 0.987601 | 3.915872 |
| 0.539575 | 0.544794 | 0.552166 | 0.552366 | 0.559759 | 0.562223 | 0.569628 | 0.576464 | 0.579134 | 0.580286 |

| AV-951 | Roscovitine | TW 37 | SB590885 | Sorafenib | PFI-1 | Camptothe | KIN001-27 | TL-1-85 | GSK69069 |
|---|---|---|---|---|---|---|---|---|---|
| NA | NA | 0.557385 | 5.7039 | NA | 6.210492 | 1.055615 | NA | NA | NA |
| 0.660119 | 3.27616 | NA | NA | 3.080339 | NA | NA | 5.069975 | 3.312115 | 1.037543 |
| -0.66395 | 3.338482 | -0.45385 | -0.45705 | 1.009039 | 2.891704 | -5.19213 | 3.478834 | 2.013457 | 3.573142 |
| -1.5444 | 4.706751 | -0.6148 | 4.404427 | 2.013602 | 2.543173 | -4.98581 | 4.424778 | 4.288123 | 1.673503 |
| 0.750395 | 3.175656 | 0.080146 | 4.406264 | 0.781878 | 2.933533 | -0.21506 | 4.120687 | 2.613269 | 2.065227 |
| 0.833489 | NA | -0.86152 | 4.562742 | NA | 2.333122 | -2.07517 | 6.047739 | 5.114546 | 5.384145 |
| 0.056506 | NA | 1.670145 | 3.766548 | 2.257135 | 2.536255 | -3.40426 | 4.881639 | 3.909791 | 3.755246 |
| 2.43777 | NA | 2.380536 | 6.333998 | NA | 4.241949 | -1.78222 | 6.966142 | 6.598674 | 6.532658 |
| 0.057577 | 4.198839 | -1.11618 | 3.252403 | 1.51937 | 1.987699 | -3.40958 | 3.710079 | 1.254627 | 1.747699 |
| 1.526932 | NA | -0.15588 | 3.338306 | NA | 3.693113 | 0.814402 | 6.000957 | 5.05758 | 3.094846 |
| 1.298026 | NA | -0.32603 | 5.322619 | NA | 3.285948 | -3.5408 | 5.471722 | 4.810184 | 0.037393 |
| 1.1969 | NA | -0.427 | NA | NA | 2.985186 | NA | 4.814877 | 4.018219 | 2.273206 |
| 1.492009 | NA | NA | 5.157172 | NA | 3.520147 | -1.79777 | 6.245414 | 5.591246 | 4.722095 |
| 1.617048 | NA | 0.571375 | NA | NA | 2.03751 | -2.82498 | 6.103648 | 3.819755 | 2.79399 |
| 0.0082 | 5.018348 | NA | NA | 2.530897 | NA | NA | 5.26519 | 4.210774 | 3.511428 |
| 1.293333 | NA | 0.398461 | NA | NA | 4.684152 | -1.39284 | 5.357479 | 4.163412 | 0.126306 |
| 0.391731 | NA | -1.16494 | 4.274598 | NA | 2.143095 | -3.50528 | 5.298006 | 4.302706 | 2.60973 |
| 0.225215 | NA | -0.57692 | 4.477383 | NA | 2.867722 | -4.545 | 5.082693 | 3.166806 | 4.630587 |
| 0.004591 | 3.652613 | NA | NA | 1.806033 | NA | NA | 4.452175 | 3.644313 | -0.6669 |
| 0.64376 | NA | -0.93576 | 3.118583 | NA | 1.830745 | -4.10177 | 5.495633 | 5.025787 | 5.025787 |
| 0.762997 | NA | -0.93705 | 3.524918 | NA | 3.047626 | -4.82835 | 4.791862 | 4.982797 | 3.624801 |
| NA | NA | 4.060943 | 3.251779 | NA | 4.646871 | 0.546217 | 4.540005 | 2.832179 | NA |
| 0.571603 | 5.276474 | 1.309729 | 4.590285 | 2.838109 | 2.932597 | -0.18095 | 4.481004 | 4.667829 | 0.303778 |
| 0.822676 | NA | -0.33223 | 4.885745 | NA | 3.840643 | -3.60373 | 5.68972 | 5.535279 | 2.036629 |
| -0.03471 | NA | -1.27174 | 3.732168 | NA | 2.988342 | -5.07166 | 5.082781 | 4.395485 | 2.988638 |
| 0.760411 | NA | 0.115799 | 4.330614 | NA | 5.084214 | -0.21876 | 6.048241 | 5.384624 | 0.626398 |
| -0.22011 | NA | 0.598479 | 3.90813 | NA | 2.457232 | NA | 4.674406 | 3.679303 | 3.685511 |
| 0.925073 | NA | 1.478517 | 3.40388 | NA | 2.502596 | -2.07749 | 5.450982 | 4.288551 | 1.244428 |
| 0.364919 | NA | -2.18179 | 3.346959 | NA | 2.883097 | -5.07787 | 4.277016 | 3.258604 | 1.942993 |
| 0.981994 | NA | -0.84704 | 4.566788 | NA | 4.202221 | -2.31132 | 5.581839 | 4.873936 | 4.577368 |
| -0.27381 | NA | -0.11513 | 4.398264 | NA | 2.55907 | NA | 4.946748 | 5.102798 | 5.082959 |
| 0.855036 | NA | NA | 3.968028 | NA | 2.100808 | -2.28135 | 4.759403 | 2.387096 | 0.724461 |
| 0.360054 | NA | -0.38057 | NA | NA | 3.874108 | -2.38295 | 5.489832 | 1.625604 | 2.227593 |
| 0.836948 | NA | -2.33301 | 4.502111 | NA | 2.576234 | -3.04869 | 5.640648 | 5.218569 | 5.218975 |
| 0.646876 | NA | 0.260653 | 5.104593 | NA | 3.548247 | -1.9367 | 5.115672 | 4.06472 | 5.070195 |
| 0.67162 | NA | 0.581329 | 3.012324 | NA | 2.874966 | -1.58087 | 5.344175 | 5.082247 | 5.186213 |
| 1.241609 | NA | -0.82457 | 4.916647 | NA | 3.233746 | -3.69785 | 4.67535 | 5.016264 | 5.63351 |
| 0.099643 | 3.98705 | -0.85147 | 3.415135 | 1.790895 | 2.668251 | -2.29173 | 4.662743 | 3.495012 | 2.324899 |
| 0.734051 | NA | -1.01786 | 2.371393 | NA | 2.41214 | -3.43303 | 5.114856 | 4.221505 | 3.84747 |
| 0.755106 | NA | -0.57276 | 3.050205 | NA | 1.855949 | -3.27093 | 4.964233 | 2.97882 | 4.505363 |
| 0.457773 | NA | 1.191745 | 4.811063 | NA | 3.644182 | -2.84862 | 5.049961 | 3.237745 | 3.927927 |
| 0.717539 | NA | 0.270975 | 4.238534 | NA | 3.344089 | -2.80743 | 4.822951 | 5.361672 | 2.515435 |
| 0.453914 | NA | -2.04639 | 2.882126 | NA | 1.777414 | -4.41227 | 5.631471 | 4.820757 | 5.125273 |
| NA | NA | -0.19667 | 5.033367 | NA | 2.43221 | -2.50186 | NA | NA | NA |
| 0.582886 | 0.585129 | 0.593007 | 0.59515 | 0.599521 | 0.610895 | 0.611135 | 0.612186 | 0.614983 | 0.623649 |

| JNK Inhibit | LFM-A13 | GDC0941 | Rapamycin | YM155 | BIX02189 | BMS-5369 | GSK26996 | Dabrafenib | PHA-6657 |
|---|---|---|---|---|---|---|---|---|---|
| 5.852651 | NA | 1.249398 | NA | NA | NA | 3.357274 | NA | 5.278504 | NA |
| NA | 5.476242 | NA | -3.42599 | -5.34776 | 3.624346 | NA | 2.415258 | NA | 3.504858 |
| 3.551988 | 3.387497 | 2.133908 | -2.98639 | -2.54843 | 4.161157 | 2.132001 | 2.470136 | -4.24168 | 2.588793 |
| 4.984626 | 3.396823 | -1.62991 | -2.05333 | -3.54137 | 5.793705 | 2.257062 | 3.713936 | 4.795094 | 3.49112 |
| 5.538863 | 5.547961 | 0.484729 | -0.30514 | -5.52965 | 4.933493 | 4.534297 | 2.895937 | 5.432845 | 4.002914 |
| 4.422683 | 3.94792 | 4.165199 | NA | -4.49332 | 5.317982 | 4.054 | NA | 3.951821 | NA |
| 5.128294 | 5.434201 | 2.615637 | -1.23472 | -1.50913 | 5.439855 | 3.100371 | 2.310455 | 4.870824 | 3.051725 |
| 6.934865 | 7.442603 | 2.534675 | NA | -3.22885 | 6.858649 | 3.996896 | NA | 6.500473 | NA |
| 3.595685 | 4.750392 | -1.2203 | -2.75719 | -4.08266 | 3.841634 | 1.775889 | 2.871075 | 4.303997 | 2.693856 |
| 3.17322 | 4.852346 | 0.949836 | NA | -1.78819 | 5.696268 | 4.915682 | NA | 3.449199 | NA |
| 5.512723 | 5.517927 | -0.56694 | NA | -2.33413 | 4.059258 | 2.308633 | NA | 4.178035 | NA |
| NA | 5.599544 | NA | NA | -3.41179 | 5.188256 | 3.544132 | NA | 5.263554 | NA |
| 4.697947 | 4.158826 | 0.588649 | NA | -2.76066 | 6.526608 | 3.381149 | NA | 5.010538 | NA |
| NA | 4.655639 | NA | NA | -5.25087 | 3.892149 | 1.860462 | NA | 5.713451 | NA |
| NA | 5.260088 | NA | -2.08591 | -4.99198 | 5.634472 | NA | 3.37436 | NA | 2.4148 |
| 5.18372 | 3.663836 | 2.314464 | NA | -1.46534 | 6.114408 | 4.380634 | NA | 5.633429 | NA |
| 4.95163 | 4.778628 | -0.78428 | NA | -0.91842 | 5.367814 | 2.367489 | NA | 4.529021 | NA |
| 4.86988 | 5.633359 | 3.151547 | NA | -5.5209 | 4.185303 | 2.808325 | NA | 5.047289 | NA |
| NA | 4.684448 | NA | -1.52716 | -5.50781 | 4.554276 | NA | 2.214684 | NA | 3.073812 |
| 3.957086 | 3.975969 | -0.12415 | NA | -0.87846 | 5.695218 | 0.515368 | NA | 3.029935 | NA |
| 3.404031 | 5.458987 | -0.28426 | NA | -4.78242 | 5.795294 | 3.376638 | NA | NA | NA |
| 5.716037 | 3.12326 | 0.944726 | NA | -4.09147 | 4.785261 | 4.458911 | NA | 5.341524 | NA |
| 5.254305 | 5.101168 | 1.156335 | -1.50105 | -2.99841 | 5.167131 | 3.830732 | 3.610317 | 3.652748 | 3.497937 |
| 5.004598 | 5.420288 | 0.580981 | NA | -5.10409 | 6.196468 | 4.372039 | NA | 5.334565 | NA |
| 4.425832 | 4.797895 | 1.149131 | NA | -1.45124 | 5.118979 | 1.60151 | NA | 1.66722 | NA |
| 4.433649 | 5.825879 | 2.9389 | NA | -4.79934 | 6.055491 | 3.584897 | NA | 5.028654 | NA |
| 3.797944 | 5.091005 | 3.617452 | NA | -4.86959 | 3.691876 | 2.431166 | NA | 3.45486 | NA |
| 5.589 | 3.913423 | 2.181634 | NA | -2.38589 | 5.293912 | 4.412101 | NA | 4.194485 | NA |
| 3.573454 | 4.776085 | 1.987233 | NA | -5.37678 | 4.153758 | 1.782869 | NA | 4.132821 | NA |
| 5.470192 | 5.936644 | 1.472859 | NA | -5.12463 | 6.119676 | 3.614445 | NA | 4.728852 | NA |
| 5.091411 | 5.447715 | 1.421294 | NA | -2.27686 | 5.44601 | 2.528531 | NA | 4.44324 | NA |
| 4.520177 | 4.681711 | 2.090643 | NA | -5.73057 | 4.687125 | 3.928087 | NA | 4.579079 | NA |
| NA | 3.306517 | NA | NA | -1.2139 | 3.050267 | 2.679726 | NA | 4.78331 | NA |
| 5.049878 | 5.574843 | 0.129865 | NA | 0.428894 | 5.350479 | 2.903844 | NA | 5.073451 | NA |
| 5.763925 | 4.361496 | 0.982115 | NA | -2.89027 | 5.981214 | 0.90483 | NA | 5.439417 | NA |
| 5.24474 | 3.609164 | -0.25825 | NA | -4.34924 | 4.955413 | 4.195284 | NA | 3.746763 | NA |
| 5.08956 | 4.789999 | 0.762204 | NA | -3.66418 | 5.486901 | 3.618682 | NA | 4.589388 | NA |
| 4.29094 | 4.956799 | 1.659058 | -2.55732 | -4.621 | 4.245416 | 1.952249 | 1.987512 | 4.187937 | 2.146295 |
| 3.989551 | 4.727149 | -0.7617 | NA | -4.11894 | 4.742537 | 2.703458 | NA | 2.106643 | NA |
| 5.190215 | 4.741457 | 2.496269 | NA | -4.9135 | 3.287472 | 1.348484 | NA | 1.431342 | NA |
| 4.330181 | 5.681666 | 1.536831 | NA | -1.76388 | 4.147705 | 3.186646 | NA | 4.155016 | NA |
| 4.36657 | 5.382655 | 0.903123 | NA | -4.31239 | 4.474958 | 4.143913 | NA | 5.198168 | NA |
| 4.889857 | 4.002146 | 2.418971 | NA | -5.44376 | 5.711313 | 2.962048 | NA | 3.236432 | NA |
| 5.242196 | NA | 4.469356 | NA | NA | NA | 4.586125 | NA | 4.612674 | NA |
| 0.623933 | 0.641583 | 0.642432 | 0.642854 | 0.646097 | 0.667223 | 0.667542 | 0.669501 | 0.682774 | 0.686679 |

| VX-702 | NVP-TAE68 | AR-42 | SNX-2112 | QL-XI-92 | EKB-569 | GW-2580 | Pyrimethar | GNF-2 | PXD101, Be |
|---|---|---|---|---|---|---|---|---|---|
| 4.787609 | NA | NA | NA | NA | NA | NA | NA | NA | NA |
| NA | 2.016292 | -2.0436 | -2.61715 | 3.584383 | -0.46831 | 5.613968 | 2.611581 | 2.954181 | -1.21619 |
| 2.590836 | -0.17547 | -1.72365 | -3.59917 | 1.967222 | 1.29196 | 4.892228 | 2.717982 | 2.110086 | -1.12735 |
| 2.900972 | 0.774207 | 0.797842 | 3.717584 | 2.877819 | 3.714263 | 4.205086 | 4.499915 | 2.892242 | 0.751473 |
| 2.731888 | 3.203112 | 0.044571 | -0.08422 | 3.011099 | NA | 4.134641 | 5.34652 | 2.535645 | 1.774144 |
| 3.441131 | NA | -1.06688 | 2.603974 | 4.483002 | 2.796068 | 6.045111 | NA | NA | -0.37275 |
| 2.019426 | 1.748977 | -0.97336 | 1.443709 | 3.160549 | 1.666127 | 5.75684 | 4.409191 | NA | -0.17878 |
| 5.265397 | NA | 6.112678 | 2.271654 | 6.292572 | 5.39232 | 7.356 | NA | NA | 5.699988 |
| 2.806988 | 1.895576 | -1.6816 | -0.18107 | 4.076618 | 2.739871 | 4.084295 | 5.109573 | 2.360701 | -1.15036 |
| 4.400454 | NA | -0.25234 | 1.072175 | 4.988167 | -0.66079 | 6.155947 | NA | NA | 1.261325 |
| 3.886784 | NA | -1.3656 | 0.139512 | 5.48343 | -0.21696 | 6.702246 | NA | NA | -1.11671 |
| NA | NA | 0.586563 | 0.90932 | 4.245013 | 2.952663 | 5.363918 | NA | NA | 0.103696 |
| 4.240882 | NA | 3.113583 | 3.655188 | 5.873623 | 4.276903 | 5.111961 | NA | NA | 3.398082 |
| 3.710282 | NA | -0.70539 | 2.781106 | 5.999238 | 4.244634 | 6.092957 | NA | NA | 0.157256 |
| NA | 2.830111 | -0.10292 | 3.666616 | 4.95651 | 2.31176 | 5.465499 | 5.140448 | NA | 0.605446 |
| 4.12698 | NA | 1.511 | 0.858121 | 4.127296 | -1.79332 | 6.117622 | NA | NA | 1.479238 |
| 3.007791 | NA | -1.43612 | -1.87605 | 4.238667 | 0.145137 | 5.660893 | NA | NA | -0.43862 |
| 3.563687 | NA | -0.4402 | -1.96379 | 4.749353 | 1.376677 | 5.666044 | NA | NA | -0.09616 |
| NA | 2.655506 | -1.50102 | -2.52856 | 4.46885 | 2.647723 | 5.375217 | 4.912884 | 2.558277 | -1.19106 |
| 3.373204 | NA | 2.345573 | 3.639045 | 5.025787 | 3.611558 | 5.406631 | NA | NA | 3.358509 |
| 3.398733 | NA | -1.02321 | 0.528462 | 4.306227 | -1.54756 | 5.339713 | NA | NA | -0.56892 |
| 4.175906 | NA | -1.47601 | -0.85805 | NA | NA | NA | NA | NA | -1.06113 |
| 3.699121 | 3.39431 | 0.7743 | -1.54408 | 3.745498 | -2.36106 | 5.41719 | 5.633895 | 3.252834 | 1.243263 |
| 3.208965 | NA | 1.088658 | 1.429839 | 3.597343 | 1.5278 | 5.664283 | NA | NA | 1.270848 |
| 2.544538 | NA | 2.184248 | 2.175656 | 4.103933 | 2.191936 | 4.815327 | NA | NA | 3.132969 |
| 3.750092 | NA | 0.249005 | 1.706811 | 3.924499 | -0.01282 | 5.510698 | NA | NA | 1.202153 |
| NA | NA | -0.54507 | -1.83649 | 3.266205 | 0.324193 | 5.330705 | NA | NA | 0.011981 |
| 4.030817 | NA | -0.23256 | 0.670988 | 3.580099 | 1.009059 | 5.841591 | NA | NA | 0.474317 |
| 3.150779 | NA | 1.006845 | -1.8572 | 3.371741 | -0.78401 | 4.42031 | NA | NA | 1.973299 |
| 3.868855 | NA | -0.81627 | 1.164362 | 4.789621 | 2.60406 | 6.035552 | NA | NA | -0.37295 |
| NA | NA | 0.072311 | 3.364227 | 5.053021 | 3.699358 | 5.096219 | NA | NA | 0.526591 |
| 3.522475 | NA | -2.85217 | -3.05084 | 3.527745 | 1.274838 | 4.479081 | NA | NA | -2.61065 |
| 2.710512 | NA | -0.25173 | -0.10873 | 4.555921 | 2.7588 | 5.563969 | NA | NA | -0.68504 |
| 3.585821 | NA | 1.10849 | 2.618822 | 5.213156 | 3.808814 | 5.807856 | NA | NA | 1.806608 |
| 4.069941 | NA | 0.490569 | -1.99764 | 4.721867 | 1.107772 | 6.461014 | NA | NA | 0.671445 |
| 3.328178 | NA | -0.11781 | 0.923613 | 4.882709 | 2.845449 | 5.901089 | NA | NA | 0.883955 |
| 4.000356 | NA | 1.280145 | 0.845171 | 5.067304 | 4.212978 | 5.187076 | NA | NA | 1.803397 |
| 2.42524 | 0.503586 | 1.005223 | NA | 3.413542 | 0.900303 | 4.546036 | 3.268316 | 2.307708 | 2.034144 |
| 3.106016 | NA | 0.913075 | 0.739341 | 4.014137 | 1.270732 | 5.447754 | NA | NA | 1.132206 |
| 3.842374 | NA | 0.225071 | -1.36377 | 4.609918 | 0.40469 | 6.203621 | NA | NA | 0.463205 |
| 3.716111 | NA | 2.470732 | 2.555212 | 4.555687 | 3.543092 | 6.191274 | NA | NA | 2.459761 |
| 3.156465 | NA | 0.302319 | 0.525769 | 3.886436 | 1.25657 | 6.062049 | NA | NA | 0.367882 |
| 2.997512 | NA | -0.53843 | 3.714137 | 3.94367 | 2.490622 | 4.919361 | NA | NA | NA |
| 3.950622 | NA | NA | NA | NA | NA | NA | NA | NA | NA |
| 0.687083 | 0.688269 | 0.690206 | 0.690724 | 0.695111 | 0.697729 | 0.698592 | 0.700278 | 0.701521 | 0.702003 |

| NU-7441 | Nilotinib | AUY922 | MPS-1-IN-: | CAY10603 | CH542480: | NSC-20789 | Afatinib | Afatinib (re | XL-184 |
|---|---|---|---|---|---|---|---|---|---|
| 4.590103 | 4.787609 | NA | NA | NA | NA | NA | -1.88671 | -0.3036 | NA |
| NA | NA | -2.78539 | 2.786613 | -0.72659 | 2.458105 | 4.17334 | NA | NA | 2.606874 |
| 1.96307 | 0.56163 | -4.54199 | 3.045482 | 0.335853 | 3.501301 | 4.069997 | 1.143399 | 3.819516 | 2.68384 |
| 0.734275 | 1.008982 | -3.01248 | 5.743041 | 1.476024 | 3.31457 | 5.570999 | -2.00007 | 2.620204 | 4.193994 |
| 3.530983 | 3.953522 | -2.57699 | 4.044058 | 0.664133 | 3.465527 | 5.346213 | -3.16462 | -1.40652 | 3.565894 |
| 2.597355 | 3.75099 | -1.81083 | 5.978935 | 0.980936 | 3.735914 | 5.941697 | 2.024299 | 5.238851 | 4.590783 |
| 2.366464 | 2.834243 | -1.42255 | 5.502872 | 1.043081 | 3.545173 | 4.552227 | 1.200269 | 4.542536 | 4.458095 |
| 4.975077 | 5.146388 | 3.268696 | 7.230106 | 3.474248 | 5.296054 | 3.541742 | 3.662703 | 6.779484 | 6.187347 |
| 1.024513 | 2.806671 | -2.13025 | 3.362133 | -0.194 | 1.357948 | 2.108958 | 0.623924 | 4.357172 | 2.542324 |
| 4.341931 | 2.95291 | -2.70842 | 6.456344 | 2.178219 | 4.386231 | 6.305868 | -1.12296 | 1.336209 | 5.312523 |
| 2.96854 | 4.358088 | -1.52468 | 3.194235 | 0.395272 | 3.524045 | 4.07233 | 0.675444 | 2.85587 | 3.183549 |
| NA | NA | -2.66888 | 4.032349 | 0.547227 | 2.590525 | 4.164566 | NA | 4.46011 | 3.435666 |
| 2.260763 | 4.240882 | -1.51007 | 6.462277 | 3.133664 | 4.330774 | 6.418955 | 2.590184 | 4.786839 | 5.034203 |
| NA | 4.212238 | NA | 5.111133 | 0.324902 | 4.124958 | 4.245491 | 2.660508 | 4.345098 | 2.818814 |
| NA | NA | -1.07979 | 4.781552 | 1.805396 | 4.359675 | 3.450768 | NA | NA | 2.745184 |
| 4.117287 | 3.499445 | -2.77732 | 6.00398 | 2.04492 | 3.28876 | 5.908364 | -2.23483 | -0.46333 | 4.260668 |
| 2.436125 | 2.588699 | -3.70537 | 5.006031 | 0.469085 | 4.289961 | 5.271973 | 0.423401 | 1.900493 | 3.188149 |
| 3.042122 | 3.556205 | -3.6823 | 5.758128 | 0.795697 | 4.369346 | 3.813343 | 1.934914 | 4.616276 | 2.76851 |
| NA | NA | -3.23394 | 3.764264 | 0.172634 | 3.479885 | 5.349756 | NA | NA | 3.84797 |
| 2.628234 | 3.006294 | -5.35485 | 5.694247 | 2.380025 | 4.19345 | 5.695218 | 1.139618 | 4.882209 | 4.331664 |
| 1.570052 | 3.199612 | -2.33016 | 5.336855 | 0.557899 | 4.032216 | 3.486565 | -3.12625 | -1.27523 | 3.328269 |
| 2.792911 | 4.207685 | -2.30323 | 6.571345 | 0.39852 | NA | 4.986046 | -1.03751 | 0.362722 | 4.841433 |
| 3.598144 | 3.699121 | -4.02412 | 5.012549 | 0.484938 | 4.624721 | 5.288635 | -2.98322 | 0.22293 | 2.747032 |
| 3.659285 | 3.658667 | -1.15198 | 5.54564 | 2.027757 | 4.662367 | 4.951853 | -0.38511 | 3.002986 | 4.235933 |
| 0.795621 | 2.363225 | -4.94243 | 5.11747 | 3.238281 | 3.652447 | 4.569701 | 1.02758 | 3.11338 | 3.753272 |
| 3.376526 | 3.281603 | -0.02729 | 6.044656 | 1.914514 | 4.474782 | 5.928987 | 1.289541 | 0.753197 | 4.558649 |
| 3.025745 | NA | -2.2824 | 4.125232 | 0.86548 | 3.080379 | 4.064545 | NA | 2.877776 | 2.536355 |
| 3.406957 | 3.763496 | -3.61863 | 5.680595 | 1.333354 | 3.988757 | 5.807761 | -0.38029 | 2.323559 | 4.313727 |
| 2.107736 | 2.525554 | -3.07919 | 3.241009 | 1.38337 | 1.863045 | 2.750155 | -1.08792 | 2.022896 | 3.394767 |
| 3.222104 | 3.849504 | -3.26979 | 4.193375 | 0.56772 | 2.666713 | 3.201261 | -1.20581 | 3.170063 | 4.115023 |
| 1.200891 | NA | -0.38714 | 5.096952 | 0.937424 | 4.25137 | 5.782732 | NA | 3.982533 | 4.205803 |
| 3.470698 | 3.627897 | -1.53489 | 4.4017 | -0.88109 | 3.236952 | 4.9104 | 1.249469 | 2.435385 | 3.734566 |
| NA | 1.515547 | -2.87048 | 4.758266 | -0.07433 | 2.330237 | 3.689142 | 0.749518 | 3.661102 | 3.463647 |
| 1.967815 | NA | 1.410102 | 5.882158 | 1.993231 | 2.518076 | 5.877909 | 1.994875 | 4.728739 | 4.510732 |
| 2.97412 | 4.150563 | -0.04934 | 2.855478 | 1.117743 | 2.356661 | 5.622175 | -0.28195 | 3.680258 | 1.393742 |
| 3.021574 | NA | -4.18408 | 5.051303 | 1.055878 | 4.365124 | 5.168436 | -3.34453 | 0.563277 | 2.601701 |
| 2.885423 | NA | -1.95152 | 3.645353 | 1.854792 | 3.447994 | 4.353161 | 1.797547 | 3.524212 | 4.213927 |
| 2.781791 | 2.853823 | -0.85689 | 3.728994 | 1.732658 | 3.071975 | 3.584056 | 0.496447 | 4.650091 | 2.831581 |
| 2.303377 | NA | -4.26883 | 4.652181 | 1.57092 | 4.197643 | 4.667619 | 0.067563 | 2.354498 | 3.476913 |
| 3.487001 | 3.47783 | -3.43144 | 4.441011 | 0.334517 | 4.699904 | 2.986218 | 1.471083 | 3.452226 | 2.50029 |
| 2.516624 | 3.190452 | -0.06927 | 5.826507 | 3.150546 | 4.521255 | 5.847135 | 2.508478 | 4.996183 | 3.540892 |
| 3.108078 | 2.230875 | -3.07698 | 4.346458 | 0.928982 | 3.74576 | 6.052186 | 0.744456 | 4.141739 | 3.958498 |
| 3.180741 | 2.682328 | -2.45218 | 5.793649 | 1.527167 | 1.514044 | 4.530666 | 1.603442 | 3.997606 | 4.35581 |
| 3.616793 | 3.69321 | NA | NA | NA | NA | NA | 2.560137 | 3.938038 | NA |
| 0.702838 | 0.705399 | 0.709396 | 0.717811 | 0.721082 | 0.728588 | 0.729988 | 0.730423 | 0.731899 | 0.731946 |

| CI-1040 | UNC0638 | AS605240 | PF-470867 | XMD11-85 | TPCA-1 | Lapatinib | PAC-1 | CX-5461 | Phenformi |
|---|---|---|---|---|---|---|---|---|---|
| 3.972134 | 3.973989 | NA | 4.811262 | 5.563582 | NA | NA | NA | NA | NA |
| NA | NA | 1.649617 | NA | NA | 2.010406 | 3.073672 | 1.874246 | 0.026409 | 5.890749 |
| -1.99169 | 2.282952 | -3.94395 | 2.396599 | NA | 2.445204 | 2.338621 | 0.316781 | -0.62228 | 4.396785 |
| 3.164951 | 5.67708 | 4.805242 | 4.397888 | 2.321065 | 5.793705 | 1.534564 | 2.328757 | 5.124274 | 5.039379 |
| 5.166616 | 2.592839 | 2.469928 | 4.042528 | 5.372279 | 5.195759 | -2.71281 | 0.108281 | 3.108656 | 2.799805 |
| 3.395788 | 2.839404 | 4.369991 | 4.271074 | 4.834456 | 5.153102 | NA | 3.278738 | 5.271585 | 9.523396 |
| 3.29204 | 2.225711 | 3.322724 | 3.09396 | NA | 4.084329 | NA | 3.044591 | 4.646264 | NA |
| 4.021057 | 4.252769 | 3.823031 | 6.898326 | 6.779484 | 5.333795 | NA | 4.207461 | 3.891753 | 12.80228 |
| 2.787536 | 2.104484 | 3.299267 | 3.142456 | 2.432939 | 1.299151 | 1.725485 | 2.524979 | 3.918742 | 6.85799 |
| 3.409918 | 2.953395 | 3.739449 | 2.268801 | 5.216912 | 5.173597 | NA | 1.562052 | 5.967653 | 7.62336 |
| 4.462912 | 2.957455 | 2.09103 | 3.208953 | NA | 3.008104 | NA | 3.505975 | 4.7355 | 8.424156 |
| NA | 6.135854 | 4.784876 | 4.083671 | NA | 5.379395 | NA | 0.503795 | 4.470973 | 6.299376 |
| 3.781467 | 3.817666 | 5.372965 | NA | 5.331334 | 6.526209 | NA | 4.364473 | 5.874036 | 6.176591 |
| 4.230546 | 3.179069 | 2.860118 | 5.322649 | NA | 2.874421 | NA | 0.430411 | 5.402958 | 8.946114 |
| NA | NA | 3.586397 | NA | NA | 4.840019 | 1.38754 | 3.672455 | 4.372724 | 8.74355 |
| 3.122363 | 6.326576 | 3.056168 | 1.444594 | 5.471015 | 4.407337 | NA | 3.968167 | 5.553187 | 7.012571 |
| 2.361759 | 5.561211 | 3.143113 | 3.945184 | NA | 4.279429 | NA | 2.299133 | 4.990878 | 8.920272 |
| 3.537881 | 3.645934 | 3.828779 | 3.39283 | 4.178719 | 3.343429 | NA | 1.24158 | 2.201829 | 7.01628 |
| NA | NA | 4.034479 | NA | NA | 2.34715 | 3.073812 | 2.011677 | 0.312511 | 6.737507 |
| 3.428073 | 4.544073 | 4.505537 | 3.610594 | NA | 5.695218 | NA | 1.597691 | 5.025787 | 4.681212 |
| 2.968502 | 2.782754 | 3.291874 | 3.236377 | 4.055961 | 5.335442 | NA | 0.883912 | 5.11952 | 7.676069 |
| 4.352849 | 2.951342 | 1.397719 | 4.249694 | 5.616502 | 5.680834 | NA | -0.17149 | 5.041547 | 4.650003 |
| 5.279098 | 2.541155 | 2.68208 | 2.941455 | 3.974219 | 4.811866 | -1.02581 | 2.586631 | 3.981738 | 6.227628 |
| 2.61742 | 4.083339 | 2.656543 | 3.675307 | 5.419375 | 4.905916 | NA | 3.914117 | 5.2279 | 8.014971 |
| 2.611736 | 5.054392 | 3.776507 | 0.762725 | NA | 5.118979 | NA | 1.002627 | 3.999714 | 7.655451 |
| 3.203268 | 2.904968 | 1.204318 | 3.561802 | 5.193691 | 6.055491 | NA | 1.195103 | 5.132962 | 6.621552 |
| NA | 2.301796 | 3.56096 | 4.616394 | NA | 2.903063 | NA | 2.539795 | 2.048743 | 6.552038 |
| 5.322402 | 2.357154 | 3.505871 | 2.236717 | 3.899376 | 3.787161 | NA | 1.391588 | 4.939799 | 10.04673 |
| 1.488465 | 2.554708 | 2.910153 | 3.932243 | 3.329556 | 2.927419 | NA | 3.318765 | 2.602044 | 4.141853 |
| 2.126584 | 6.032354 | 3.30267 | 4.825497 | NA | 3.613531 | NA | 1.096621 | 0.310433 | 7.938055 |
| NA | 4.675894 | 2.972683 | 3.773707 | NA | 4.465968 | NA | 2.159913 | 5.070511 | 7.581208 |
| 3.962318 | 5.798967 | 0.249085 | NA | NA | 4.495839 | NA | 3.150096 | 2.567526 | 4.34625 |
| 1.970986 | 2.533597 | 2.768857 | 4.01291 | NA | 3.862828 | NA | 3.517746 | 3.386249 | 7.792536 |
| 1.245701 | 5.041958 | 5.88745 | 4.415633 | 4.715174 | 5.888406 | NA | 3.832681 | 5.218975 | 6.972361 |
| 1.886175 | 2.263947 | 2.572472 | 5.712422 | NA | 3.845419 | NA | 1.556364 | 5.761548 | 6.648494 |
| 3.857466 | 3.237746 | 3.841175 | 3.666143 | 4.693301 | 4.172357 | NA | 2.751707 | 3.306363 | 8.294665 |
| 2.753834 | 6.043373 | 5.219404 | 4.846005 | NA | 4.503016 | NA | 4.052577 | 3.12319 | 8.529266 |
| 3.012349 | 2.62395 | 2.471164 | 3.548741 | NA | 2.337012 | 3.163974 | 2.616189 | 4.305964 | 7.463908 |
| 2.099167 | 3.373456 | 2.807732 | 4.138736 | 4.415083 | 3.97072 | NA | 3.258269 | 4.051549 | 7.760395 |
| 3.175033 | 5.371517 | 3.281022 | 4.169336 | NA | 2.28914 | NA | 3.535843 | 3.428942 | 8.108964 |
| 4.534155 | 1.861754 | 6.09329 | 5.018172 | 5.336976 | 4.39139 | NA | 1.83288 | 5.024068 | 8.087767 |
| 2.458413 | 3.037918 | 3.480537 | 4.594681 | 4.472011 | 4.662283 | NA | 3.70612 | 4.044398 | 8.30301 |
| 1.417166 | 4.453841 | 3.063355 | 4.707349 | NA | 5.795889 | NA | 3.390919 | 4.566566 | 6.695354 |
| 5.083925 | 2.417718 | NA | 3.570355 | NA | NA | NA | NA | NA | NA |
| 0.734907 | 0.740255 | 0.741793 | 0.752393 | 0.754054 | 0.760964 | 0.768205 | 0.77282 | 0.784214 | 0.786837 |

| BMS-3455 | Ruxolitinib | Dasatinib | AMG-706 | 17-AAG | Genentech | TG101348 | TAK-715 | NVP-BEZ23 | AICAR |
|---|---|---|---|---|---|---|---|---|---|
| NA | NA | NA | 3.042874 | -0.69282 | NA | NA | NA | -0.13898 | 8.844391 |
| 1.480605 | 4.156765 | 2.065998 | NA | NA | 0.74058 | 1.577777 | 3.469136 | NA | NA |
| 1.469439 | 3.492908 | 1.093992 | 2.358481 | -1.49997 | 0.322881 | 1.84472 | 2.78577 | -3.03883 | 6.712822 |
| 3.360024 | 3.870857 | 2.577811 | 3.481203 | -2.31014 | 4.102536 | 1.857313 | 4.683752 | -3.1555 | 5.49695 |
| 1.555763 | 4.878211 | 3.368934 | 4.002534 | -1.55497 | 2.702504 | 4.540716 | 2.778277 | -0.8976 | 9.027364 |
| 4.29979 | 4.683448 | NA | 2.233271 | -0.14064 | 2.650736 | 3.173703 | 5.208619 | -1.75625 | 6.686801 |
| 2.761855 | 1.737125 | NA | 3.475066 | 0.84498 | 2.004868 | 3.125034 | 2.849556 | -2.80214 | 5.245158 |
| 4.909071 | 6.370024 | NA | 5.239431 | 0.356745 | 6.398223 | 6.75041 | 6.748859 | -1.56655 | 11.42798 |
| 2.808349 | 3.770163 | 0.612499 | 1.347216 | -3.14967 | 1.980013 | 2.255211 | 3.788854 | -3.94539 | 7.385097 |
| 4.365702 | 5.044947 | NA | 0.567118 | -1.60678 | 3.549596 | 4.235615 | 4.508573 | -3.12873 | 7.902158 |
| 4.352806 | 4.915884 | NA | 3.545683 | 2.547974 | 1.603634 | 2.414425 | 5.322748 | -2.59201 | 6.356385 |
| 4.025994 | 4.771254 | NA | NA | NA | 2.327923 | 4.18159 | 4.027173 | NA | NA |
| 5.931134 | 5.110702 | NA | 2.621117 | -0.69256 | 5.870966 | 5.87356 | 6.146787 | -2.2443 | 6.733672 |
| 3.260641 | 4.243979 | NA | NA | NA | 1.163952 | 2.155304 | 5.385805 | NA | 6.14438 |
| 3.777648 | 4.348434 | NA | NA | NA | 4.305281 | 3.30877 | 4.785963 | NA | NA |
| 3.681208 | 5.040616 | NA | 3.754268 | -0.82145 | 3.14313 | 4.132332 | 4.763337 | -0.86384 | 10.09475 |
| 3.705488 | 4.250756 | NA | 2.505685 | -3.52288 | 4.659639 | 3.39673 | 5.425187 | -2.72139 | 7.272557 |
| 3.623507 | 4.388529 | NA | 3.562689 | -1.17741 | 3.05105 | 0.909216 | 4.226195 | -1.49514 | 9.226187 |
| 2.970077 | 4.036688 | 2.607668 | NA | NA | 4.706926 | 2.744714 | 3.713131 | NA | NA |
| 5.148813 | 3.813217 | NA | 2.066931 | -3.18784 | 5.025787 | 5.025787 | 4.350795 | -3.04651 | 7.117109 |
| 3.038119 | 4.363225 | NA | 0.757001 | -2.23975 | 1.953714 | 4.545179 | 4.479637 | -2.78995 | 5.744526 |
| 3.003501 | 5.048796 | NA | 3.163943 | 0.805653 | 0.654839 | NA | 3.742945 | -1.63304 | 6.545857 |
| 3.102062 | 4.64228 | 3.032558 | 3.686576 | -3.32052 | 3.212415 | 3.453834 | 3.374528 | -1.54633 | 9.1791 |
| 3.617251 | 4.893044 | NA | 3.961929 | -0.86709 | 2.490732 | 3.321816 | 4.167677 | -2.22733 | 7.45244 |
| 3.30105 | 3.713302 | NA | 2.813544 | -3.70808 | 4.271332 | 2.862477 | 4.041953 | -3.70533 | 8.003472 |
| 1.608832 | 4.62652 | NA | 3.267463 | 0.157628 | 1.855317 | 3.739577 | 3.113664 | -0.89263 | 7.44694 |
| 2.734961 | 3.974559 | NA | 2.281679 | -2.00694 | 2.414317 | 2.279453 | 3.536747 | -2.24137 | NA |
| 3.300083 | 4.346308 | NA | 4.02362 | -0.14693 | 1.984319 | 1.5139 | 4.235209 | -1.71631 | 7.086926 |
| 2.919517 | 3.393942 | NA | 2.228723 | 1.854018 | 0.85601 | 2.973198 | 3.136524 | -2.40426 | 7.688379 |
| 4.386855 | 4.594451 | NA | 3.57672 | -0.32102 | 4.102572 | 4.434858 | 5.918941 | -1.52864 | 7.224606 |
| 4.01852 | 4.335495 | NA | 1.824882 | -2.03904 | 2.610475 | 2.467448 | 5.721273 | -3.08552 | NA |
| 2.57175 | 4.239109 | NA | 3.140235 | -1.16285 | 1.255102 | 2.677606 | 3.294152 | -0.23281 | 8.236142 |
| 2.820095 | 4.239378 | NA | NA | NA | 4.044023 | 2.811361 | 4.400231 | NA | 6.479598 |
| 5.412098 | 4.545511 | NA | 1.854794 | -2.50776 | 5.218975 | 3.770245 | 5.804281 | -3.65759 | 9.11021 |
| 3.272023 | 4.834002 | NA | 4.040403 | -1.70795 | 1.85609 | 1.980963 | 5.52161 | -1.64113 | 7.849978 |
| 3.67996 | 4.010384 | NA | 3.630403 | -2.59821 | 3.857246 | 5.221392 | 4.213611 | -2.72106 | 10.47609 |
| 5.427624 | 4.96353 | NA | 3.380492 | -1.89469 | 5.63351 | 5.63351 | 5.795632 | -2.42916 | 8.889763 |
| 2.334016 | 4.122487 | 0.870634 | 3.065631 | 2.244729 | 0.306483 | 1.884913 | 3.326798 | -1.36907 | 7.110512 |
| 3.940485 | 4.441179 | NA | 2.944958 | -2.8811 | 3.652924 | 2.797964 | 4.513919 | -4.05299 | 6.640379 |
| 5.092835 | 4.035931 | NA | 2.538937 | -0.5065 | 4.927853 | 1.68866 | 4.827939 | -3.26506 | 6.599183 |
| 4.704191 | 4.542293 | NA | 2.065277 | 0.718551 | 4.568514 | 3.192412 | 5.43264 | -1.86911 | 9.4234 |
| 3.888127 | 4.7138 | NA | 1.695116 | -1.57464 | 2.785127 | 2.693406 | 4.940371 | -1.48942 | 7.793849 |
| 3.361386 | 4.322205 | NA | 3.483187 | 2.330592 | 3.178081 | 2.464213 | 5.577659 | -2.42654 | 7.280164 |
| NA | NA | NA | 3.81883 | -1.85895 | NA | NA | NA | -1.91618 | 8.864872 |
| 0.793102 | 0.795646 | 0.797176 | 0.797838 | 0.798774 | 0.813687 | 0.814597 | 0.82122 | 0.823809 | 0.828188 |

| Doxorubici | T0901317 | KIN001-13! | BI-2536 | SB-505124 | S-Trityl-L-c | Crizotinib | VX-680 | KIN001-26( | NG-25 |
|---|---|---|---|---|---|---|---|---|---|
| NA | NA | NA | NA | 6.210492 | NA | NA | NA | NA | NA |
| -2.70972 | 3.877567 | 4.318811 | -3.00343 | NA | 0.397173 | 1.50137 | -0.77524 | 3.031617 | 2.46731 |
| -4.637 | 4.080408 | 3.505941 | -2.25839 | 3.551734 | 0.011126 | 2.299579 | -0.18595 | 2.671271 | 2.080918 |
| -2.75937 | 3.947928 | 2.796554 | -0.19138 | 4.388215 | 2.123669 | 3.49112 | 2.52357 | 4.981075 | 4.630537 |
| -1.80159 | 3.773281 | 4.061593 | -0.57864 | 4.531512 | 1.453089 | 3.430817 | 3.339792 | 3.222526 | 2.633152 |
| -0.01839 | 4.949612 | NA | NA | 3.739866 | NA | NA | NA | 3.340558 | 5.115586 |
| -1.16334 | 3.966248 | 3.745624 | NA | 3.81957 | NA | 2.734849 | NA | 3.080497 | 3.416227 |
| 2.159909 | 6.460971 | NA | NA | 6.303439 | NA | NA | NA | 7.373833 | 5.992365 |
| -0.24212 | 3.504784 | 3.723279 | -1.79695 | 2.076782 | 0.603888 | 2.631489 | 2.082314 | 2.541251 | 0.73489 |
| -0.12649 | 5.96027 | NA | NA | 5.831973 | NA | NA | NA | 6.355309 | 4.751695 |
| -0.65685 | 4.87641 | NA | NA | 3.217225 | NA | NA | NA | 3.294206 | 2.454418 |
| -1.88605 | 4.764517 | NA | NA | 5.367025 | NA | NA | NA | 3.198355 | 3.611499 |
| 1.467339 | 4.996348 | NA | NA | 3.093944 | NA | NA | NA | 5.990943 | 5.64462 |
| -0.75743 | 5.991904 | NA | NA | 4.25745 | NA | NA | NA | 2.608791 | 1.674064 |
| 1.249172 | 4.922995 | 3.757308 | NA | NA | NA | 3.258954 | NA | 3.858381 | 2.20592 |
| -0.37439 | 5.344397 | NA | NA | 5.633429 | NA | NA | NA | 3.341443 | 2.365726 |
| -1.04064 | 4.144481 | NA | NA | 4.385598 | NA | NA | NA | 4.258131 | 3.614848 |
| -3.10796 | 5.005356 | NA | NA | 2.716691 | NA | NA | NA | 2.82399 | 2.242116 |
| -3.66393 | 4.029709 | 3.72874 | -2.17766 | NA | 0.66045 | 2.634566 | 1.77432 | 2.423276 | 3.430778 |
| -4.61416 | 4.947826 | NA | NA | 4.573303 | NA | NA | NA | 5.327426 | 4.549326 |
| -2.0144 | 4.568248 | NA | NA | 4.99592 | NA | NA | NA | 2.986423 | 2.882705 |
| -0.35877 | NA | NA | NA | 4.676264 | NA | NA | NA | 3.474304 | 2.032423 |
| -0.35397 | 5.164507 | 4.147424 | -0.67029 | 3.193834 | 2.771476 | 3.61805 | 3.150168 | 3.041838 | 2.790786 |
| 0.098678 | 5.308507 | NA | NA | 5.415348 | NA | NA | NA | 5.262457 | 2.908526 |
| -3.85896 | 3.498371 | NA | NA | 2.923443 | NA | NA | NA | 4.387924 | 2.843374 |
| -1.00439 | 5.040191 | NA | NA | 4.591907 | NA | NA | NA | 4.418658 | 5.378887 |
| -2.42945 | 3.753374 | NA | NA | 4.555963 | NA | NA | NA | 2.583761 | 3.012949 |
| -2.20292 | 5.654859 | NA | NA | 3.753626 | NA | NA | NA | 5.216967 | 4.159663 |
| -2.55378 | 3.897102 | NA | NA | 4.663493 | NA | NA | NA | 2.866057 | 3.215987 |
| -0.92408 | 5.339087 | NA | NA | 4.382918 | NA | NA | NA | 3.081551 | 5.271987 |
| -0.56808 | 4.833441 | NA | NA | 4.821685 | NA | NA | NA | 4.434105 | 4.026682 |
| 0.924243 | 3.756211 | NA | NA | 4.189064 | NA | NA | NA | 3.28883 | 2.459416 |
| -0.51189 | 4.868822 | NA | NA | 3.745679 | NA | NA | NA | 1.947214 | 1.683087 |
| 0.74887 | 5.218975 | NA | NA | 4.121788 | NA | NA | NA | 5.754562 | 4.946356 |
| 1.961412 | 5.797881 | NA | NA | 5.165894 | NA | NA | NA | 3.549546 | 2.41195 |
| -3.36534 | 5.264998 | NA | NA | 3.594791 | NA | NA | NA | 5.521311 | 3.798698 |
| 0.447101 | 5.40857 | NA | NA | 3.992766 | NA | NA | NA | 3.921764 | 3.554305 |
| -0.37073 | 3.800141 | 3.413704 | -2.30954 | 3.413822 | -0.11729 | 1.68148 | -0.09458 | 4.331873 | 2.633202 |
| -1.55072 | 4.588915 | NA | NA | 3.496922 | NA | NA | NA | 3.320192 | 2.569789 |
| -0.20114 | 5.483417 | NA | NA | 5.02234 | NA | NA | NA | 3.477402 | 2.214306 |
| 1.087604 | 5.048869 | NA | NA | 4.703808 | NA | NA | NA | 3.117976 | 2.111105 |
| -1.99572 | 5.315549 | NA | NA | 5.227128 | NA | NA | NA | 2.662542 | 4.341636 |
| 0.214758 | 4.451482 | NA | NA | 4.981335 | NA | NA | NA | 4.959171 | 4.264245 |
| NA | NA | NA | NA | 5.499229 | NA | NA | NA | NA | NA |
| 0.830326 | 0.83058 | 0.832886 | 0.843776 | 0.847495 | 0.85638 | 0.859863 | 0.870024 | 0.872833 | 0.874203 |

| IOX2 | CUDC-101 | THZ-2-102- | KIN001-23( | XMD8-85 | PHA-79388 | Paclitaxel | AT-7519 | WZ-1-84 | A-443654 |
|---|---|---|---|---|---|---|---|---|---|
| 0.837354 | NA | NA | NA | NA | NA | NA | NA | NA | NA |
| NA | -1.98332 | -4.6611 | 3.70977 | 2.1667 | -0.17865 | -3.71086 | -1.45534 | 4.272731 | -1.80198 |
| -0.46497 | -1.62807 | -3.84322 | 1.970969 | 1.968148 | 0.281275 | -3.28914 | -1.17964 | 2.982 | -1.14793 |
| 0.332729 | 1.146509 | -0.21571 | 5.097952 | 2.576967 | 0.838846 | -2.30929 | 0.591528 | 3.736908 | -0.40677 |
| 0.845417 | -0.12856 | 1.650483 | 4.623281 | 4.865326 | 0.63857 | -3.03167 | -0.75152 | 2.915952 | -1.36386 |
| -0.31847 | -0.96447 | 1.523697 | 4.793952 | NA | 4.339685 | NA | 3.864877 | NA | NA |
| 0.413508 | -0.60677 | -3.7251 | 4.837872 | NA | 2.837928 | NA | -0.50674 | NA | NA |
| 1.081498 | 6.241743 | 1.438521 | 6.082187 | NA | 3.65151 | NA | 3.811822 | NA | NA |
| -0.58079 | -1.80365 | -2.65961 | 3.188591 | 2.50571 | 2.061036 | -3.5202 | 0.086521 | 3.351076 | -1.48606 |
| 0.166672 | 0.586051 | -0.30114 | 4.458671 | NA | 4.225734 | NA | 1.957556 | NA | NA |
| 1.180459 | -1.87024 | -1.7842 | 5.250875 | NA | 1.816496 | NA | 2.059761 | NA | NA |
| -0.26392 | -0.53838 | -2.73099 | 5.143582 | NA | 3.212265 | NA | 2.484548 | NA | NA |
| -2.3375 | 2.409749 | -1.50554 | 5.681041 | NA | 5.859442 | NA | 3.726413 | NA | NA |
| 1.142796 | -1.60425 | -4.7506 | 4.880431 | NA | 5.161846 | NA | 1.686588 | NA | NA |
| NA | 2.384753 | 3.663537 | 5.081012 | 4.831334 | 5.001499 | NA | 2.983026 | NA | NA |
| 0.933743 | 1.20715 | 0.831275 | 3.95165 | NA | 3.440303 | NA | 1.595702 | NA | NA |
| 0.262359 | -1.57441 | 1.057146 | 4.257654 | NA | 4.220698 | NA | 4.041995 | NA | NA |
| 0.117506 | -0.85281 | -2.36085 | 5.043774 | NA | 2.210171 | NA | 1.473544 | NA | NA |
| NA | -1.7585 | NA | 4.340733 | 3.192977 | 3.289196 | -3.24952 | 2.733509 | 4.295515 | -2.83555 |
| 0.276654 | 2.152213 | 3.016145 | 5.025787 | NA | 5.025787 | NA | 4.415688 | NA | NA |
| -0.19219 | -1.42083 | -3.7093 | 3.769136 | NA | 2.703554 | NA | -0.15966 | NA | NA |
| 0.972894 | -1.52628 | -0.45991 | 3.828461 | NA | NA | NA | -0.67045 | NA | NA |
| -0.21277 | 0.567281 | -1.42166 | 4.34981 | 4.339307 | 2.419239 | -1.86303 | 1.207616 | 2.515443 | -0.53924 |
| 0.806538 | 0.252424 | -1.36797 | 5.586055 | NA | 2.923298 | NA | 0.212099 | NA | NA |
| -1.06929 | 2.283707 | 3.036537 | 4.449548 | NA | 3.498695 | NA | 3.399148 | NA | NA |
| 0.292999 | -0.16961 | 3.280488 | 4.43433 | NA | 1.045993 | NA | -1.28585 | NA | NA |
| -0.05831 | -0.87339 | -3.85556 | 3.72014 | NA | -0.11869 | NA | -0.04216 | NA | NA |
| 0.845592 | -0.18999 | -0.97955 | 4.440287 | NA | 2.913415 | NA | 0.271671 | NA | NA |
| 0.058046 | 0.236623 | -3.80951 | 4.034549 | NA | 0.025923 | NA | 0.336323 | NA | NA |
| 0.734037 | -0.60779 | 0.766898 | 5.492175 | NA | 3.774692 | NA | 3.03351 | NA | NA |
| -0.7038 | -0.36927 | -2.82512 | 5.044448 | NA | 4.074012 | NA | 2.559784 | NA | NA |
| 0.495293 | -2.845 | 1.266304 | 3.849538 | NA | 1.662637 | NA | 1.193524 | NA | NA |
| -0.31652 | -0.68321 | -5.50399 | 4.911226 | NA | -0.95459 | NA | -1.35744 | NA | NA |
| -0.18535 | 0.411307 | -1.65748 | 5.216755 | NA | 5.218975 | NA | 5.20973 | NA | NA |
| 0.776063 | -1.2354 | -3.40686 | 5.695949 | NA | 2.654869 | NA | 0.975429 | NA | NA |
| 0.507529 | 0.057652 | -2.20209 | 3.745702 | NA | 5.12703 | NA | 2.376495 | NA | NA |
| 0.840667 | 0.944092 | -2.43107 | 5.440738 | NA | 4.841573 | NA | 4.480404 | NA | NA |
| 0.077844 | 0.277956 | -2.92459 | 4.43526 | 1.544326 | 0.045198 | -4.18014 | -1.3041 | 3.396886 | -0.13079 |
| -0.08431 | 0.874477 | 0.622752 | 4.514177 | NA | 3.382854 | NA | 2.789341 | NA | NA |
| 0.731928 | 0.298186 | -4.08781 | 4.448596 | NA | 4.180406 | NA | 3.979823 | NA | NA |
| 0.31819 | 2.2928 | -0.70463 | 5.299269 | NA | 4.504188 | NA | 3.350579 | NA | NA |
| 0.42183 | -0.28429 | -2.83244 | 4.543182 | NA | 2.190901 | NA | 1.407277 | NA | NA |
| 0.25765 | -0.25791 | 3.384665 | 5.032702 | NA | 4.262064 | NA | 1.133582 | NA | NA |
| 0.617747 | NA | NA | NA | NA | NA | NA | NA | NA | NA |
| 0.875597 | 0.880701 | 0.883114 | 0.89785 | 0.903118 | 0.921829 | 0.924624 | 0.929303 | 0.937041 | 0.940261 |

| CAL-101 | MS-275 | 5-Fluorour | YM201636 | VX-11e | Tamoxifen | GW843682 | KU-55933 |
|---|---|---|---|---|---|---|---|
| NA | NA | NA | NA | NA | 5.517345 | NA | 5.784574 |
| 3.636004 | -0.58813 | 3.230813 | 1.263637 | 3.122882 | NA | -3.88399 | NA |
| 4.627047 | -0.44861 | 1.445262 | 1.305973 | -2.39865 | 3.142001 | -3.04946 | 3.53101 |
| 5.6585 | 0.39849 | 2.687069 | 4.431127 | 4.431127 | 4.232997 | 0.156356 | 2.286584 |
| 4.554617 | -0.09488 | 2.472024 | 1.79643 | 4.217888 | 3.671786 | -1.06692 | 4.983052 |
| 5.331047 | NA | 5.789608 | 4.111627 | 4.598998 | 3.143101 | NA | 4.157245 |
| 5.151389 | 2.469508 | 4.318062 | 3.69797 | 4.051752 | 3.284496 | NA | 4.034008 |
| 5.3792 | NA | 2.86835 | 2.905473 | 5.977022 | 6.010764 | NA | 5.375938 |
| 2.789295 | 0.294068 | 3.446957 | 1.267866 | 2.371256 | 2.762005 | -1.75004 | 2.291018 |
| 5.744791 | NA | 6.318133 | 2.680829 | 5.138925 | 5.138826 | NA | 5.779971 |
| 2.442384 | NA | 3.170151 | 1.625427 | 3.949949 | 0.989673 | NA | 4.51455 |
| 5.315742 | NA | 4.17852 | 2.825129 | 4.073324 | 4.682365 | NA | NA |
| 4.42948 | NA | 6.872441 | 4.171945 | 5.180822 | 3.814127 | NA | 3.577733 |
| 4.996884 | NA | 4.899689 | 1.050895 | 5.092605 | 3.405997 | NA | NA |
| 4.196215 | 4.279269 | 4.722323 | 2.741446 | 4.333219 | NA | NA | NA |
| 6.499153 | NA | 4.456134 | 2.331987 | 3.101301 | 4.881859 | NA | 4.494996 |
| 4.985353 | NA | 5.48657 | 1.305967 | 3.73276 | 2.526963 | NA | 4.147221 |
| 4.436019 | NA | 2.189233 | 1.87854 | 4.063854 | 4.290192 | NA | 3.854204 |
| 5.192255 | 0.269085 | -0.03334 | 1.972173 | 3.91339 | NA | -2.76998 | NA |
| 5.687864 | NA | 6.165221 | 4.33264 | 4.33264 | 1.95347 | NA | 2.99346 |
| 5.169696 | NA | 4.687255 | 1.160961 | 4.380709 | 4.28756 | NA | 3.364089 |
| 2.900432 | NA | 4.308261 | 0.704161 | 3.611555 | 4.941355 | NA | 5.718079 |
| 4.069912 | 0.556679 | 3.677685 | 1.751956 | 3.014921 | 3.681607 | -0.01293 | 5.302693 |
| 6.000125 | NA | 5.330484 | 2.885133 | 4.891654 | 4.708914 | NA | 5.496251 |
| 4.716121 | NA | 3.747422 | 3.756401 | 3.756401 | 3.659164 | NA | 2.706084 |
| 4.911703 | NA | 4.798478 | 3.853334 | 4.692914 | 3.129231 | NA | 5.329521 |
| 5.320532 | NA | 1.579634 | 2.451254 | 1.966738 | 3.819746 | NA | 4.53498 |
| 3.379774 | NA | 5.15441 | 1.923532 | 4.680014 | 3.080212 | NA | 3.337899 |
| 2.757618 | NA | 3.009851 | 2.330554 | 2.970942 | 3.970346 | NA | 2.809349 |
| 5.506811 | NA | 2.463229 | 3.0298 | 4.77021 | 4.403192 | NA | 4.857135 |
| 5.48291 | NA | 3.787246 | 2.577845 | 4.131678 | 4.241445 | NA | 3.797854 |
| 4.675324 | NA | 1.898559 | 1.061551 | 3.912365 | 4.41588 | NA | 4.797944 |
| 3.65486 | NA | 1.289946 | 2.484973 | 2.173967 | 4.092118 | NA | NA |
| 5.861397 | NA | 6.346772 | 3.461603 | 4.513653 | 4.35494 | NA | 2.639337 |
| 4.432598 | NA | 6.913547 | 2.553338 | 1.432403 | 4.628307 | NA | 4.155952 |
| 5.604367 | NA | 4.479224 | 1.121821 | 3.438532 | 3.342459 | NA | 4.114792 |
| 5.710773 | NA | 3.959331 | 2.566549 | 4.905852 | 4.734617 | NA | 4.601226 |
| 3.701149 | 1.713433 | 4.497567 | 2.063947 | 3.473847 | 1.96404 | -3.01707 | 3.435883 |
| 4.925331 | NA | 4.326438 | 3.031186 | 3.323847 | 3.103332 | NA | 4.160861 |
| 3.853685 | NA | 3.580095 | 2.386597 | 4.128718 | 3.619871 | NA | 4.576612 |
| 5.380704 | NA | 5.883768 | 2.209145 | 4.499301 | 4.629835 | NA | 4.333134 |
| 4.356412 | NA | 4.569461 | 1.031991 | 4.666768 | 4.533981 | NA | 4.88549 |
| 5.234808 | NA | 4.791823 | 4.422052 | 4.345528 | 3.83439 | NA | 4.729443 |
| NA | NA | NA | NA | NA | 3.093078 | NA | NA |
| 0.940812 | 0.947702 | 0.974393 | 0.976488 | 0.981433 | 0.987697 | 0.990895 | 0.994282 |
